# Precipitation is the main axis of tropical phylogenetic turnover across space and time

**DOI:** 10.1101/2022.05.27.493777

**Authors:** Jens J. Ringelberg, Erik J.M. Koenen, Benjamin Sauter, Anahita Aebli, Juliana G. Rando, João R. Iganci, Luciano P. de Queiroz, Daniel J. Murphy, Myriam Gaudeul, Anne Bruneau, Melissa Luckow, Gwilym P. Lewis, Joseph T. Miller, Marcelo F. Simon, Lucas S.B. Jordão, Matías Morales, C. Donovan Bailey, Madhugiri Nageswara-Rao, Oriane Loiseau, R. Toby Pennington, Kyle G. Dexter, Niklaus E. Zimmermann, Colin E. Hughes

## Abstract

Early natural historians – Compte de Buffon, von Humboldt and De Candolle – established ecology and geography as two principal axes determining the distribution of groups of organisms, laying the foundations for biogeography over the subsequent 200 years, yet the relative importance of these two axes remains unresolved. Leveraging phylogenomic and global species distribution data for Mimosoid legumes, an pantropical plant clade of 3,400 species, we show that the water availability gradient from deserts to rainforests dictates turnover of lineages within continents across the tropics. We demonstrate that 95% of speciation occurs within a precipitation niche, showing profound phylogenetic niche conservatism, and that lineage turnover boundaries coincide with isohyets of precipitation. We reveal similar patterns on different continents, implying that evolution and dispersal follow universal processes.

## Introduction

Ever since natural historians first observed that groups of organisms are geographically restricted – palms to the tropics, pines to north temperate climates, extant lemurs to Madagascar, hummingbirds to the Americas (*1*) – understanding the factors that shape turnover of evolutionary lineages has been a central question in biogeography and macroevolution (*2–4*). While the importance of geography and ecology as the main axes determining the distribution of lineages, i.e. phylogenetic turnover or beta diversity, was already established >200 years ago (*1*), their relative contributions are still debated (*4–8*). The key processes determining turnover, phylogenetic niche conservatism (PNC) and geographic dispersal limitation (DL) (*3, 6*), have both spatial and temporal dimensions, reflecting the spatial distribution of lineages and the evolutionary time scales over which barriers to dispersal, whether environmental (i.e., PNC) or spatial (i.e., DL), are overcome (*6, 9*). To understand the contemporary spatial structure of diversity, turnover across space and time need to be considered together (*4, 6, 9, 10*), but this is rarely achieved in empirical studies.

For most terrestrial organisms, oceans and areas experiencing frost pose major barriers to dispersal and adaptation, resulting in high phylogenetic turnover amongst continents (*11, 12*) and across the tropical-temperate divide, manifest in tropical niche conservatism (*3, 13*). However, more general insights remain elusive. Precipitation appears to be a major axis of tropical turnover for some lineages (*3, 14*) but not others (*6*), and the relative importance of PNC versus DL within continents is disputed (*3, 6*). Whether phylogenetic turnover shows congruent patterns, driven by similar processes on different continents, remains unclear. Likewise, uncertainty remains about temporal turnover of lineages and the influence of PNC and DL through time (*2, 8, 15*). This lack of understanding about spatial and temporal turnover is aggravated by: 1) some reports of lineage- and region-specific idiosyncrasies of phylogenetic turnover (*11, 16*); 2) primary focus on spatial rather than temporal turnover (*6, 11*); 3) strong focus on understanding the latitudinal diversity gradient (*17*), rather than turnover longitudinally across continents within latitudinal zones, where water availability gradients are just as prominent as latitudinal temperature gradients; and 4) lack of high resolution species occurrence data and robust phylogenies for species-rich clades.

This lack of understanding applies especially to turnover of tropical plant lineages, since almost all studies of tropical plant species turnover are geographically restricted and/or lack precise species distribution data or phylogenies (*3, 6*). This gap in the tropics is critical given that plants are the primary producers and structural elements of terrestrial ecosystems (*17*), and the tropics are the most species-rich region of the planet (*3, 17*). To gain a more general understanding of adaptation and turnover of woody plants across the tropics, standardised analyses of species-rich and geographically and ecologically widespread tropical lineages across continents are needed to disentangle regional and taxon effects on the drivers of phylogenetic turnover across space and time.

We investigate phylogenetic turnover through the last c. 45 Myr across the global lowland tropics using the Mimosoid clade of legumes. Mimosoids, originating in the Eocene (*18*), are functionally diverse, comprise c. 3,400 species of trees, shrubs, geoxyles, and lianas, and make up ecologically important elements of all major lowland tropical biomes (Fig. 1e), including deserts, seasonally dry forests, savannas and rainforests (*18, 19*): across tropical biomes and continents, they constitute 5-17% of all species (*20*). Mimosoids contain numerous iconic tropical tree radiations, such as wattles (*Acacia*) with > 1000 species across Australia, umbrella thorn ‘Acacias’ (*Senegalia* and *Vachellia*) with > 150 species dominating African savannas, and ice-cream bean trees (*Inga*) with c. 300 species, a model system for Amazonian rainforest diversification (*19*).

**Figure 1.**
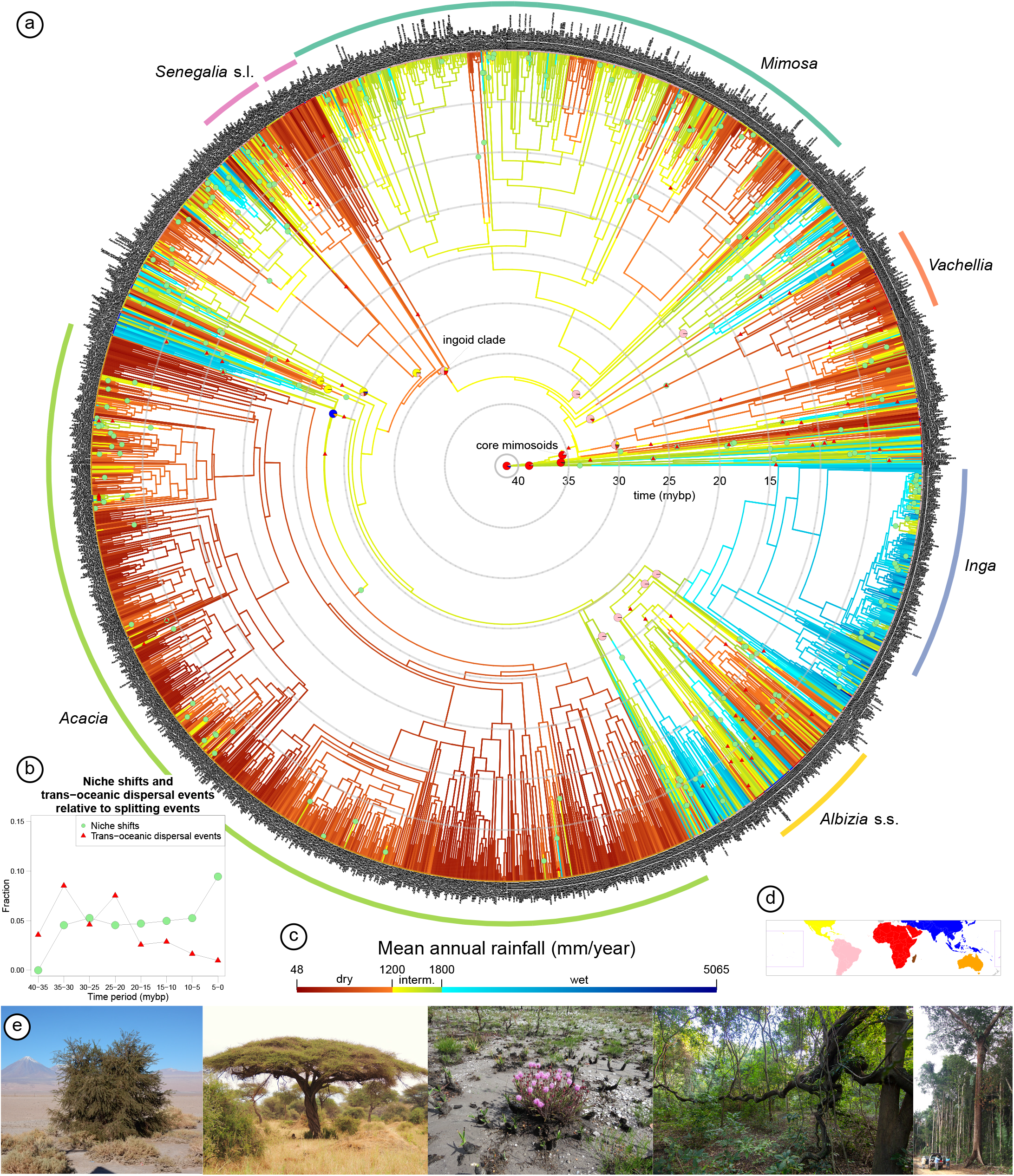
a) Phylogeny of Mimosoid legumes showing the evolution of precipitation niches and transcontinental dispersal events through time. Branch colours correspond to MAP estimates (see c for scale). Pie charts at tips and nodes of named clades (sensu (*18*)) represent observed and estimated spatial distributions (based on area definitions in d). Ancestral niches and areas were estimated using a complete metachronogram for Caesalpinioideae, including non-Mimosoid Caesalpinioideae taxa, but only the Mimosoid clade is shown here. Green circles on branches indicate shifts between precipitation categories (following (*14*)) that encompass a difference of at least 250 mm MAP; red triangles indicate postulated trans-continental dispersals according to the best-supported model. The six most species-rich genera are labelled. b) Fractions of niche shifts and trans-continental dispersal events, averaged across multiple optimisations (*20*), relative to total splitting events plotted through time for 5 Myr bins. e) Mimosoid growth form diversity across the tropical precipitation gradient, from deserts (left) through savannas to rainforests (right). See SI for species names and acknowledgements for photographers.

Using newly-generated phylogenomic, taxonomic, and occurrence datasets, we assess 1) whether turnover is primarily structured by dispersal limitation (DL) or phylogenetic niche conservatism (PNC), and which environmental factors determine PNC; 2) what levels of PNC vs niche shifting and DL vs long-distance dispersal through time characterise a clade that has fully colonized the lowland tropics; 3) whether patterns and drivers of turnover are similar across continents, implying general processes shaping turnover, or whether phylogenetic turnover is primarily driven by taxon and area-specific idiosyncrasies; and 4) whether the tempo of lineage diversification and turnover through time are associated with global Cenozoic climate change.

### Phylogeny and distribution of Mimosoids

We generated a robust, time-calibrated backbone phylogeny based on sequences of 997 nuclear genes (*18*) for 89 of 90 Mimosoid genera and 420 species (*20*)(Figs. S1-S33). We combined this well-resolved backbone with 15 species phylogenies derived from diverse DNA sequence data types to generate a species-level meta-chronogram for 1863 species (54% of Mimosoids) (Figs. 1, S34). Using a new taxonomic checklist, we assembled 423,333 quality-controlled species occurrence records (Fig. S35) to derive climate niches for 93% of Mimosoid species (*20*). We show that Mimosoids span a 100-fold range in mean annual precipitation (MAP) from < 50 mm in Somalia (*Vachellia qandalensis*), the Atacama of Chile (*Prosopis tamarugo*), and the Namib desert (*Senegalia montis-usti*), to > 5000 mm in the hyper-wet rainforests of the Colombian Chocó (*Zygia dissitiflora*) and the Indian western Ghats (*Archidendron monadelphum* var. *gracile*).

### Dispersal limitation or phylogenetic niche conservatism?

Pantropically, geographic distance explains a much greater fraction of Mimosoid phylogenetic turnover than climatic distance (Figs. 2, S45), in line with DL caused by oceanic barriers as the most important factor shaping the global distribution of lineages (*11, 12*). Nevertheless, despite high DL, trans-oceanic dispersal has been important in shaping the distribution of Mimosoids: trans-continentally distributed Mimosoid clades are common, including five pantropical genera (*Entada, Vachellia, Neptunia, Senegalia* and *Parkia*), with 61 (±27) trans-oceanic disjunctions inferred across the phylogeny (Fig. 1, Table S17). Given the Early to Mid-Eocene crown age of Mimosoids (Fig. 1), these disjunctions and the strong geographical structuring of global Mimosoid phylogenetic turnover are explained not by vicariance (*21*), but by stochastic sweepstakes long-distance dispersal (*22*) followed by in-situ diversification within continents, against a backdrop of DL. The percentage of phylogenetic nodes associated with trans-oceanic dispersal, a measure of DL, varies from 6.9% (±3.5%) between 20 and 35 Mya to 2% (±0.9%) in the last 20 Myr (Fig. 1, Table S17), suggesting an early burst of Mimosoid dispersal establishing their pantropical distribution.

**Figure 2.**
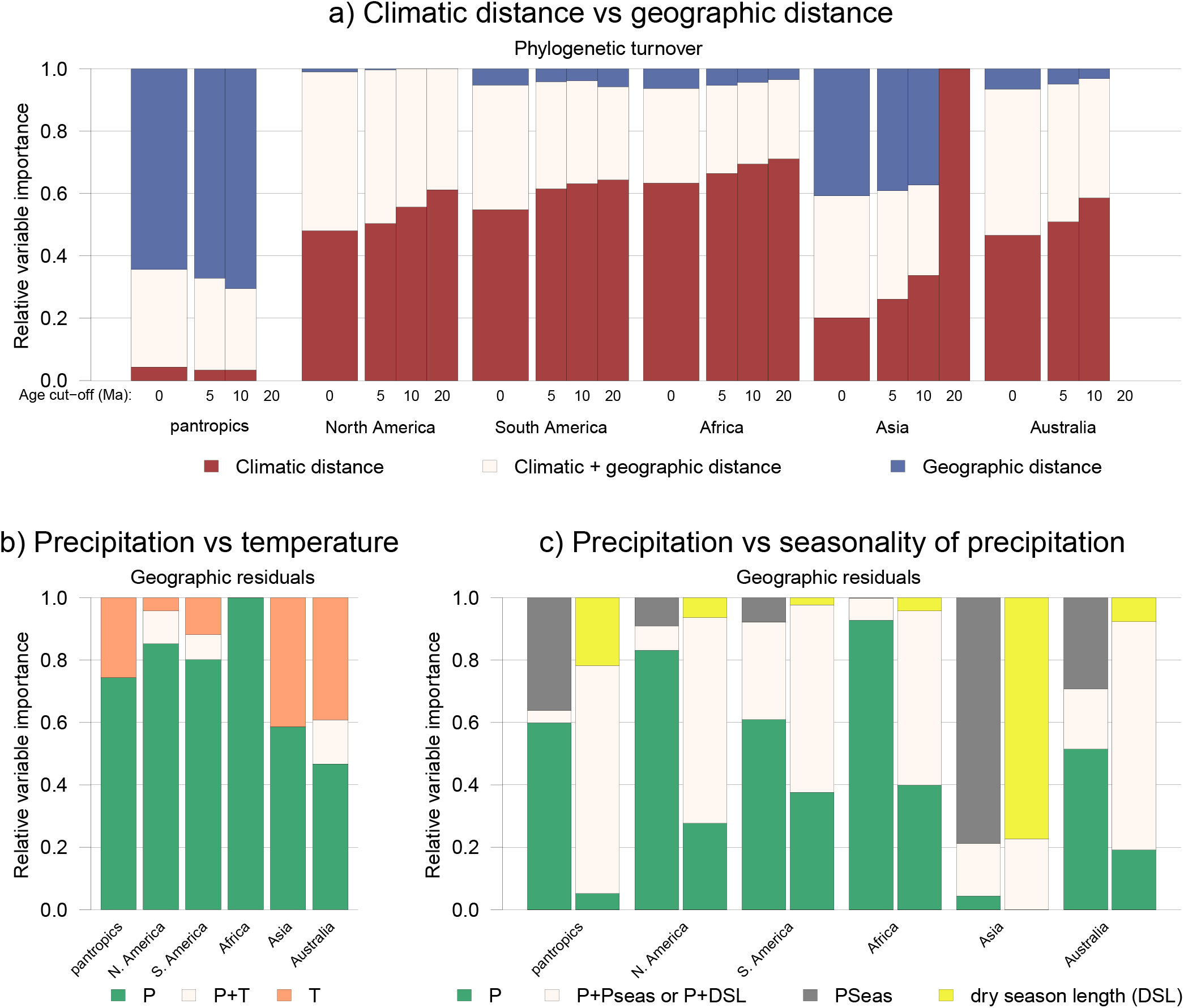
Drivers of phylogenetic turnover of Mimosoid legumes across the global lowland tropics. Bars show fractions of phylogenetic turnover explained by predictors (rescaled to add up to 1). a) Phylogenetic turnover explained by climatic distance (maroon), geographic distance (blue), or their interaction (cream). Turnover is assessed across four depths in the phylogeny: with the full metachronogram (age cut-off of 0), and with all clades younger than 5, 10, and 20 million years collapsed. Note that it was not possible to fit a model to the phylogeny collapsed at 20 million years for the pantropical and Australian models. b) Phylogenetic turnover explained by mean annual precipitation (MAP; green) and/or annual mean temperature (orange). Turnover is expressed as phylogenetic turnover not explained by geographic distance (‘geographic residuals’). c) Geographic residuals of phylogenetic turnover explained by MAP (green) and/or precipitation seasonality (grey; left) or DSL (i.e., the number of consecutive months with precipitation < 100 mm/month; yellow; right). See Fig. S45 for results obtained with an alternative, genus-level Mimosoid phylogeny. Abbreviations: P = MAP, T = annual mean temperature, Pseas = precipitation seasonality, and DSL = dry season length.

Contrary to the high phylogenetic turnover among continents, within four of five continents spatial distance plays a minor role in explaining turnover (Fig. 2a). The exception is within Asia, likely because the greater climatic homogeneity and numerous island archipelagos across SE Asia favour DL-driven turnover. On all other continents climatic distance has a much larger effect than spatial distance (Fig. 2a), suggesting relatively infrequent adaptation to novel climatic conditions compared to within-continent dispersal. In other words, for Mimosoids within continents, it has been *‘easier* [for lineages] *to move than to evolve’* (*9*), implying strong PNC (*3, 6*) and little DL. The impact of climatic distance is most pronounced deeper in the phylogeny (Fig. 2a) (*23*), suggesting that in the past climatic tolerances evolved even less frequently relative to lineage diversification and within-continent dispersal (*6*), although uncertainty increases for reconstructions further back in time. Furthermore, even when covariation between climatic and geographic distance is accounted for by analysing the residuals of a linear regression of phylogenetic turnover with geographic distance (*24, 25*), climatic distance still explains a considerable fraction of phylogenetic turnover (Fig. 2; Table S15). This contrasts with patterns of global mammalian phylogenetic turnover (*24*), indicating a stronger impact of climate on the distributions of vascular plants than mammals. Drivers of turnover are remarkably similar across continents: compared to precipitation, temperature explains only a small fraction of phylogenetic turnover (Fig. 2b), because Mimosoids are largely confined to <2,500 m elevation in the tropics. Whether total annual precipitation or precipitation seasonality is more important in driving turnover is less clear, depending on how seasonality is quantified (Fig. 2c, Table S15). Although the distribution of Mimosoids across precipitation niches varies across continents (Fig. 3), reflecting available niche space, our results (Fig. 2) strongly suggest that evolution and dispersal of Mimosoids across continents followed universal processes and drivers associated with water availability.

**Figure 3.**
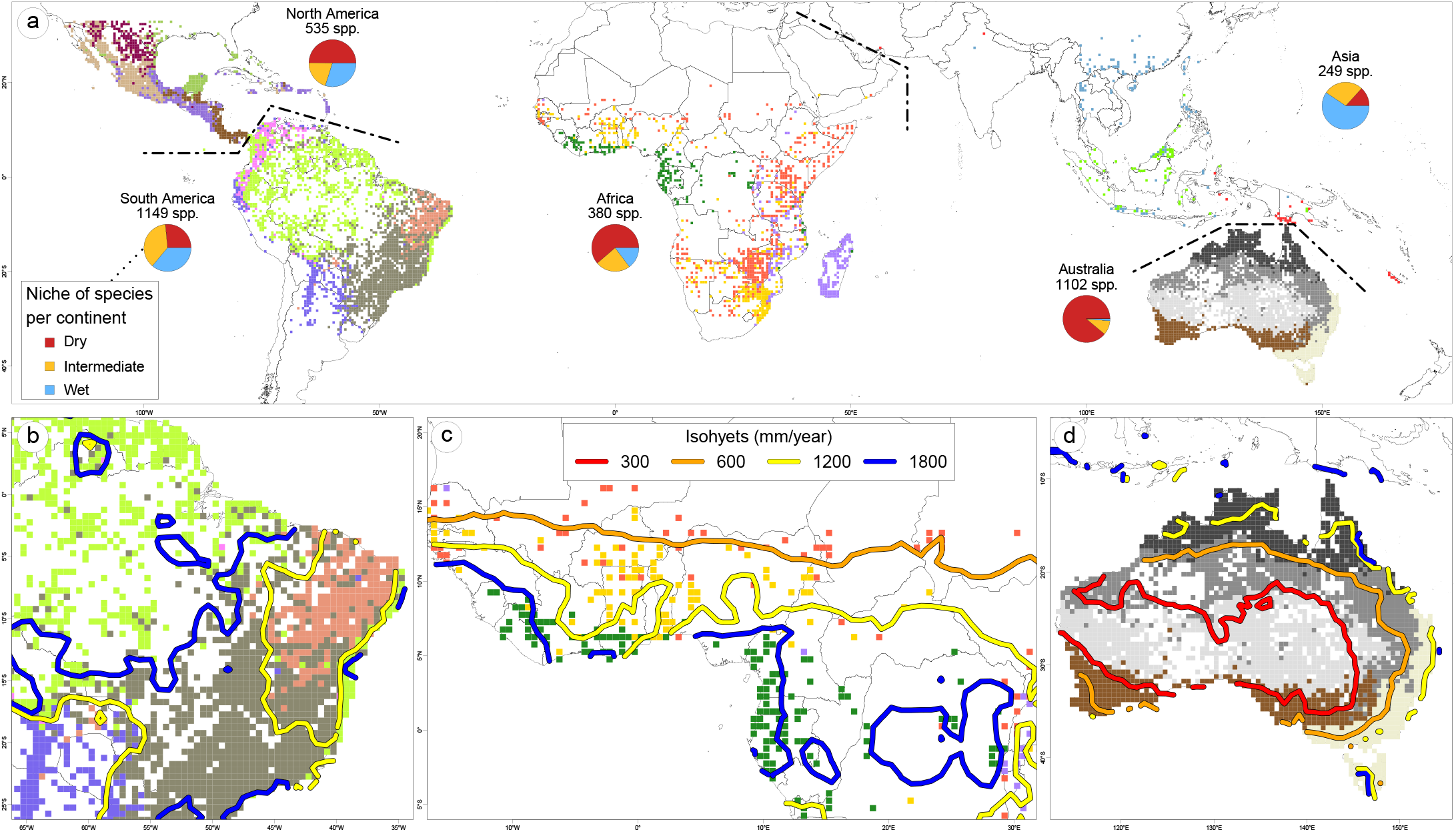
a) Phylogenetic regionalization of Mimosoid legumes within continents, and correspondence between phyloregion boundaries and isohyets in b) eastern South America, c) west Africa, and d) Australia. Coloured map cells denote phyloregions but are not indicative of phylogenetic proximity between regions; dashed lines indicate borders between continents and regionalizations. Pie charts (a) show niche distribution of species (categories follow Fig. 1). See Figs. S36-S41, S44 for results of pantropical clustering, different numbers of phyloregions, and clusters obtained using ancient turnover, geographic residuals, and a genus-level Mimosoid phylogeny.

Two additional lines of evidence suggest that the evolution and distribution of Mimosoids are structured by precipitation. First, we observe high phylogenetic signal of MAP and dry season length (DSL) (Pagel’s lambda (*26*) of 0.88 and 0.84, respectively, with 1 being the maximum), both considerably higher than the phylogenetic signal of MAP measured across 1,100 lowland tropical tree genera in South America (i.e., lambda of 0.5) (*14*). Second, spatial clustering of phylogenetic turnover via bioregionalization analyses (*23*), estimated independently of climate, reveals that on most continents higher-level clusters, representing the deepest phylogenetic divergences between areas, involve phyloregions characterised by large water availability differences (Figs. 3, S36-S44). Many lower-level clusters, i.e., geographic areas that are less phylogenetically divergent, are also characterised by different precipitation regimes (Fig. S43) and their boundaries broadly coincide with isohyets of precipitation (and major vegetation types) (Fig. 3). For example, in eastern South America phyloregions defined solely by Mimosoid turnover closely match the seasonally dry Caatinga, Cerrado (savanna), Chaco dry woodland, and Amazonian and Mata Atlântica rainforests (*27*) (Fig. 3), supporting regional analyses (*28*). This correspondence between isohyets and phyloregional boundaries is not perfect, nor equally compelling in all regions (Figure S42), indicating that other factors such as fire, soil, or herbivory also shape vegetation patterns (*29, 30*). Nevertheless, the overall similarity between precipitation regimes and phyloregions is striking, providing independent evidence that precipitation is the main driver of evolutionary turnover within the tropics.

### Phylogenetic niche conservatism through time

The prevalence of PNC related to precipitation demonstrates that precipitation gradients strongly influence the evolution and distribution of diversity within continents (*3, 13, 31*), yet Mimosoids were able to overcome adaptive barriers to occupy all tropical lowland climates and diversify as ubiquitous components of dry and wet biomes (Figs. 1, 3) (*18, 19, 28*), showing considerable climatic niche evolution. This apparent paradox between high PNC and niche evolvability, suggests the balance between PNC and niche evolution is central to explain turnover. We show that throughout the evolutionary history of Mimosoids circa 5% of speciation events involved a shift in precipitation regime, regardless of whether MAP (Fig. 1) or DSL are used (Fig. S46). This rate remained remarkably constant (Fig. 1), until apparently increasing in the last five Myr, although this may be due to undersampling of taxa within precipitation regimes towards the tips of the phylogeny. Thus, while many more species resulted from diversification within a precipitation niche (i.e., 95%) than via adaptation across precipitation regimes (i.e., 5%), we document 263 shifts in precipitation niche over the last 45 Myr. Many of these shifts spawned species-rich clades confined to wet or dry niches: for example, rapid diversification of the c. 350 species in *Inga* and allies followed a shift into wet Neotropical rainforests c. 17 Mya; similarly, *Acacia*, with > 1000 species largely in dry areas of Australia followed dispersal and a niche shift to drier climates c. 23 Mya (Fig. 1). This Mimosoid-wide 5% average of niche shifts masks considerable heterogeneity in niche specificity within subclades, with some genera like *Mimosa* and *Albizia* more labile than others (Fig. 1). Lack of adaptation to different temperature regimes contrasts with findings for precipitation. Only 7.7% of Mimosoids occur outside the tropics (just 320 taxa in nine genera have > half their occurrences north or south of 33 degrees latitude, with 288 of these in *Acacia*) and no diverse extra-tropical radiations are evident, indicating strong tropical niche conservatism (*3*) in Mimosoids, perhaps because they have never evolved an annual life history, that would be beneficial in temperate zones.

Just how prevalent PNC is across the tree of life, and how it influences diversity patterns, are heavily debated (*3, 6, 9, 32*). An important factor is phylogenetic scale which clearly influences assessment of PNC (*32*), yet almost all estimates of PNC are from studies of small lineages or geographically confined floras (*30, 31, 33*). Here, using a large pantropical clade of c. 3,400 species, we show that the evolution and distribution of Mimosoids were profoundly shaped by PNC, demonstrating that conservatism does indeed structure larger, older clades that are ecologically diverse, even when their constituent lineages vary substantially in how niche-conserved they are (Fig. 1). Remarkably, our 5% estimate for niche shifts through time closely matches estimates from continental floras, such as 4% shifts in the Southern Hemisphere (*33*) and 7% in tropical Africa (*31*). Theoretical models of diversification suggest that intermediate levels of PNC and niche evolution result in the highest species richness (*15, 34*). Under these models, overly high PNC limits distributions resulting in high extinction; conversely, if niche evolution is too prevalent this results in smaller numbers of more widespread species (*15, 34*). We suggest that our observed 20:1 PNC to niche evolution ratio, i.e., repeated individual niche colonizations followed by in-situ diversification of niche-conserved and often species-rich clades, ultimately resulting in c. 3,400 species across the global tropics, provides a compelling empirical estimate of optimal PNC, corroborating theoretical model predictions (*15, 34*) and other empirical observations (*31, 33*). Thus, while potent examples of PNC or niche evolution can be derived from studies of small clades (*30, 33, 35*), PNC to niche evolution ratios need to be assessed across species-rich, ecologically-diverse clades to test the ubiquity of this 20:1 ratio.

### Macro-evolutionary species turnover

Quantifying turnover of lineages through time and assessing whether, like spatial turnover, episodes of extinction and speciation are also associated with climate, are more challenging than quantifying spatial turnover because of the difficulties of estimating extinction rates. To account for these difficulties, we estimated speciation rates through time across various levels of temporal turnover implied by a wide range of fixed extinction rates. These analyses show that while the centre of gravity of speciation moves from the tips to deeper in the phylogeny under increasing extinction rates (Fig. S48), there is significant diversification rate heterogeneity among lineages and through time across all extinction rates (Fig. S49). Notably, across almost all potential extinction rates we find a marked increase in speciation rate in the Late Eocene or at the Eocene-Oligocene transition (c. 34 Mya; Fig. S49). This acceleration in speciation coincides with the transition from ‘Warmhouse’ to ‘Coolhouse’ conditions, the most prominent Cenozoic climate transition which coincided with formation of ice sheets in Antarctica (*36*). This acceleration in speciation also coincides with the transition from Mimosoid lineages of typically large tree and liana genera of African and Asian humid forests to dry-adapted lineages of the core Mimosoid clade originating in the Oligocene (34 – 23 Mya; Fig. 1). This prominent transition from slower to faster diversification thus appears to be associated with cooling and global expansion of dry habitats in the Oligocene and Miocene (*37*), confirming the adaptability of Mimosoids across precipitation regimes as seen from patterns of spatial turnover. Elevated speciation rates were sustained through this Coolhouse period of global aridification (*37*) until the mid- to late-Miocene (c. 13 – 8 Mya; Fig. S49), notably by a set of arid radiations, especially of *Acacia* with > 1000 species in Australia, and diversification along the mainly dry-adapted backbones of the core mimosoid and ingoid clades (Figs. 1, S48). More recent radiations are ecologically diverse and geographically scattered, including both wet forest lineages such as *Archidendron* (SE-Asian), and the Jupunba and Inga clades (both Neotropical) (Figs. 1, S48), as well as the mostly arid synchronous radiations of three clades of Madagascan Mimosoids in the last c. 15-10 Myr (Figs. 1, S31).

Radiations in the Mimosoid clade, characterized by nodes showing elevated gene tree conflict and often very short branches (Fig. S31), are temporally and geographically scattered and episodically nested across the phylogeny (Fig. S48). This pattern is consistent with a model of episodic species turnover (*2*), and suggests a molecular signature of macroevolutionary processes. The regionally heterogeneous opportunities provided by global climate change (*8*) and long-distance dispersal, in combination with the ability of Mimosoids to diversify within and across precipitation niches (Fig. 1e), have repeatedly resulted in phylogenetically and geographically scattered rapid radiations across water availability gradients, ultimately leading to the current pantropical prevalence of Mimosoids.

## Supporting information

Supplementary methods, results, tables, and figures

## Acknowledgements

The authors thank B. Adhikari, D. Lorence, B. Marazzi, É. de Souza, the G, K, JRAU, L, MEL, NY, FHO, P, RB, WAG and Z herbaria, the Millennium Seed Bank, Kew, the South African National Botanical Institute, the Direction de Environment, New Caledonia, and the National Tropical Botanical Garden, Hawaii, U.S.A. for provision of leaf or DNA samples; P. Ribeiro, É. de Souza, M. Morim, and F. Bonadeu in Brazil, and staff of the Kew Madagascar Conservation Centre and Botanical and Zoological Garden of Tsimbazaza, Madagascar for fieldwork support; the national regulatory authorities of Brazil for access to DNA of Brazilian plants (authorized through SISGEN n° A464716), and Madagascar (research permit 006/14/MEF/SG/DGF/DCB.SAP/SCB); U. Grossniklaus, V. Gagliardini, Dept Plant & Microbial Biology, Univ. Zurich for use of their TapeStation; the S3IT, Univ. Zurich for use of the ScienceCloud computational infrastructure; D. Filer for assistance with taxonomic databasing, C. Graham and M. Kessler for advice, and O. Whaley, Robur.q (Wikimedia Commons), and BT Wursten (www.mozambiqueflora.com) for permission to use photos in Fig. 1.

## Author contributions

Conceptualization: JJR, EJMK, NEZ, CEH

Investigation: JJR, EJMK, BS, AA, JGR, OL

Resources: EJMK, JRI, LPdQ, DJM, MG, AB, ML, GPL, JTM, MFS, LSBJ, MM, CDB, MNR, RTP, KGD, CEH

Funding acquisition: CEH, EJMK, RTP, CDB, AB, LPdQ, JGR, JRI, MM, LSBJ, MFS

Writing – original draft: JJR, EJMK, CEH

Writing – review and editing: all authors

## References

1. A. von Humboldt, A. Bonpland, Essai sur la géographie des plantes; accompagné d’un table physique des régions équinoxales (Levrault, Schoell et Compagnie, Libraires, Paris, France, 1805).

2. E. J. M. Koenen, J. J. Clarkson, T. D. Pennington, L. W. Chatrou, Recently evolved diversity and convergent radiations of rainforest mahoganies (Meliaceae) shed new light on the origins of rainforest hyperdiversity. New Phytol. 207, 327–339 (2015).

3. R. A. Segovia, R. T. Pennington, T. R. Baker, F. C. de Souza, D. M. Neves, C. C. Davis, J. J. Armesto, A. T. Olivera-Filho, K. G. Dexter, Freezing and water availability structure the evolutionary diversity of trees across the Americas. Sci. Adv. 6, eaaz5373 (2020).

4. W. D. Kissling, W. L. Eiserhardt, W. J. Baker, F. Borchsenius, T. L. P. Couvreur, H. Balslev, J.-C. Svenning, Cenozoic imprints on the phylogenetic structure of palm species assemblages worldwide. Proc. Natl. Acad. Sci. U.S.A. 109, 7379–7384 (2012).

5. H. Qian, N. G. Swenson, J. Zhang, Phylogenetic beta diversity of angiosperms in North America. Glob. Ecol. Biogeogr. 22, 1152–1161 (2013).

6. W. L. Eiserhardt, J.-C. Svenning, W. J. Baker, T. L. P. Couvreur, H. Balslev, Dispersal and niche evolution jointly shape the geographic turnover of phylogenetic clades across continents. Sci. Rep. 3, 1164 (2013).

7. M. R. Carvalho, C. A. Jaramillo, F. de la Parra, D. Caballero-Rodríguez, F. Herrera, S. L. Wing, B. L. Turner, C. D⍰Apolito, M. Romero-Báez, P. Narváez, C. Martínez, M. Gutierrez, C. Labandeira, G. Bayona, M. J. Rueda, M. Paez-Reyes, D. Cárdenas, Á. Duque, J. L. Crowley, C. Santos, D. Silvestro, Extinction at the end-Cretaceous and the origin of modern Neotropical rainforests. Science. 372, 63–68 (2021).

8. Y. Xing, R. E. Onstein, R. J. Carter, T. Stadler, H. P. Linder, Fossils and a large molecular phylogeny show that the evolution of species richness, generic diversity, and turnover rates are disconnected. Evolution. 68, 2821–2832 (2014).

9. M. J. Donoghue, A phylogenetic perspective on the distribution of plant diversity. Proc. Natl. Acad. Sci. U.S.A. 105, 11549–11555 (2008).

10. C. H. Graham, P. V. A. Fine, Phylogenetic beta diversity: Linking ecological and evolutionary processes across space in time. Ecol. Lett. 11, 1265–1277 (2008).

11. C. König, P. Weigelt, H. Kreft, Dissecting global turnover in vascular plants. Glob. Ecol. Biogeogr. 26, 228–242 (2017).

12. F. Mazel, R. O. Wüest, J.-P. Lessard, J. Renaud, G. F. Ficetola, S. Lavergne, W. Thuiller, Global patterns of β-diversity along the phylogenetic time-scale: The role of climate and plate tectonics. Glob. Ecol. Biogeogr. 26, 1211–1221 (2017).

13. D. M. Neves, A. J. Kerkhoff, S. Echeverría-Londoño, C. Merow, N. Morueta-holme, R. K. Peet, B. Sandel, J.-C. Svenning, S. K. Wiser, B. J. Enquist, The adaptive challenge of extreme conditions shapes evolutionary diversity of plant assemblages at continental scales. Proc. Natl. Acad. Sci. U.S.A. 118 (2021), doi:10.1073/pnas.2021132118.

14. D. M. Neves, K. G. Dexter, T. R. Baker, F. Coelho de Souza, A. T. Oliveira-Filho, L. P. Queiroz, H. C. Lima, M. F. Simon, G. P. Lewis, R. A. Segovia, L. Arroyo, C. Reynel, J. L. Marcelo-Peña, I. Huamantupa-Chuquimaco, D. Villarroel, G. A. Parada, A. Daza, R. Linares-Palomino, L. V. Ferreira, R. P. Salomão, G. S. Siqueira, M. T. Nascimento, C. N. Fraga, R. T. Pennington, Evolutionary diversity in tropical tree communities peaks at intermediate precipitation. Sci. Rep. 10, 1–7 (2020).

15. T. F. Rangel, N. R. Edwards, P. B. Holden, J. A. F. Diniz-Filho, W. D. Gosling, M. T. P. Coelho, F. A. S. Cassemiro, C. Rahbek, R. K. Colwell, Modeling the ecology and evolution of biodiversity: Biogeographical cradles, museums, and graves. Science. 361 (2018), doi:10.1126/science.aar5452.

16. F. P. Peixoto, F. Villalobos, A. S. Melo, J. A. F. Diniz-Filho, R. Loyola, T. F. Rangel, M. V. Cianciaruso, Geographical patterns of phylogenetic beta-diversity components in terrestrial mammals. Glob. Ecol. Biogeogr. 26, 573–583 (2017).

17. H. Kreft, W. Jetz, Global patterns and determinants of vascular plant diversity. Proc. Natl. Acad. Sci. U.S.A. 104, 5925–5930 (2007).

18. E. J. M. Koenen, C. A. Kidner, É. R. de Souza, M. F. Simon, J. R. V. Iganci, J. A. Nicholls, G. K. Brown, L. P. de Queiroz, M. A. Luckow, G. P. Lewis, R. T. Pennington, C. E. Hughes, Hybrid capture of 964 nuclear genes resolves evolutionary relationships in the mimosoid legumes and reveals the polytomous origins of a large pantropical radiation. Am. J. Bot. 107, 1710–1735 (2020).

19. G. P. Lewis, B. Schrire, B. Mackinder, M. Lock, Legumes of the world (Royal Botanic Gardens, Kew, United Kingdom, 2005).

20. See supplementary materials and methods.

21. M. Lavin, B. P. Schrire, G. P. Lewis, R. T. Pennington, A. Delgado-Salinas, M. Thulin, C. E. Hughes, A. B. Matos, M. F. Wojciechowski, Metacommunity process rather than continental tectonic history better explains geographically structured phylogenies in legumes. Philos. Trans. R. Soc. Lond. B. Biol. Sci. 359, 1509–1522 (2004).

22. G. G. Simpson, Mammals and land bridges. J. Washingt. Acad. Sci. 30, 137–163 (1940).

23. B. H. Daru, M. van der Bank, T. J. Davies, Unravelling the evolutionary origins of biogeographic assemblages. Divers. Distrib. 24, 313–324 (2018).

24. C. Penone, G. C. Costa, B. G. Weinstein, C. H. Graham, A. D. Davidson, T. M. Brooks, C. Rondinini, S. B. Hedges, Global mammal beta diversity shows parallel assemblage structure in similar but isolated environments. Proc. R. Soc. B Biol. Sci. 283, 1–9 (2016).

25. B. Saladin, W. Thuiller, C. H. Graham, S. Lavergne, L. Maiorano, N. Salamin, N. E. Zimmermann, Environment and evolutionary history shape phylogenetic turnover in European tetrapods. Nat. Commun. 10, 1–9 (2019).

26. M. Pagel, Inferring the historical patterns of biological evolution. Nature. 401, 877–884 (1999).

27. R. T. Pennington, M. Lavin, The contrasting nature of woody plant species in different neotropical forest biomes reflects differences in ecological stability. New Phytol. 210, 25–37 (2016).

28. A. T. Oliveira-Filho, D. Cardoso, B. D. Schrire, G. P. Lewis, R. T. Pennington, T. J. Brummer, J. Rotella, M. Lavin, Stability structures tropical woody plant diversity more than seasonality: Insights into the ecology of high legume-succulent-plant biodiversity. South African J. Bot. 89, 42–57 (2013).

29. G. R. Moncrieff, W. J. Bond, S. I. Higgins, Revising the biome concept for understanding and predicting global change impacts. J. Biogeogr. 43, 863–873 (2016).

30. J. J. Ringelberg, N. E. Zimmermann, A. Weeks, M. Lavin, C. E. Hughes, Biomes as evolutionary arenas: Convergence and conservatism in the trans-continental succulent biome. Glob. Ecol. Biogeogr. 29, 1100–1113 (2020).

31. A.-P. Gorel, O. J. Hardy, G. Dauby, K. G. Dexter, R. A. Segovia, K. Steppe, A. Fayolle, Climatic niche lability but growth form conservatism in the African woody flora. Ecol. Lett. (2022), doi:10.1111/ele.13985.

32. C. H. Graham, D. Storch, A. Machac, Phylogenetic scale in ecology and evolution. Glob. Ecol. Biogeogr. 27, 175–187 (2018).

33. M. D. Crisp, M. T. K. Arroyo, L. G. Cook, M. A. Gandolfo, G. J. Jordan, M. S. McGlone, P. H. Weston, M. Westoby, P. Wilf, H. P. Linder, Phylogenetic biome conservatism on a global scale. Nature. 458, 754–756 (2009).

34. O. Hagen, A. Skeels, R. Onstein, W. Jetz, L. Pellissier, Earth history events shaped the evolution of biodiversity across tropical rainforests. Proc. Natl. Acad. Sci. U.S.A. 118 (2021), doi:10.1073/pnas.2026347118/-/DCSupplemental.y.

35. M. J. Donoghue, E. J. Edwards, Model clades are vital for comparative biology, and ascertainment bias is not a problem in practice: a response to Beaulieu and O’Meara (2018). Am. J. Bot. 106, 1–4 (2019).

36. T. Westerhold, N. Marwan, A. J. Drury, D. Liebrand, C. Agnini, E. Anagnostou, J. S. K. Barnet, S. M. Bohaty, D. De Vleeschouwer, F. Florindo, T. Frederichs, D. A. Hodell, A. E. Holbourn, D. Kroon, V. Lauretano, K. Littler, L. J. Lourens, M. Lyle, H. Pälike, U. Röhl, J. Tian, R. H. Wilkens, P. A. Wilson, J. C. Zachos, An astronomically dated record of Earth’s climate and its predictability over the last 66 million years. Science. 369, 1383–1388 (2020).

37. M. Arakaki, P.-A. Christin, R. Nyffeler, A. Lendel, U. Eggli, R. M. Ogburn, E. Spriggs, M. J. Moore, E. J. Edwards, Contemporaneous and recent radiations of the world’s major succulent plant lineages. Proc. Natl. Acad. Sci. U.S.A. 108, 8379–8384 (2011).

