## Supplementary methods, results, tables, and figures for "Precipitation is the main axis of tropical phylogenetic turnover across space and time"

#### Index

1. Methods page 2
    1.1 Phylogenetic inference page 2
        1.1.1 Backbone topology page 2
        1.1.2 Time calibration page 6
        1.1.3 Metachronogram page 7
    1.2 Taxonomic checklist and occurrence dataset page 10
    1.3 Phylogenetic turnover across space page 11
    1.4 Optimisations of climate and geography page 14
    1.5 Phylogenetic turnover across time page 15
2. Results page 17
    2.1 Phylogenetic inference page 17
        2.1.1 Backbone topology page 17
        2.1.2 Time calibration page 19
        2.1.2 Metachronogram page 19
    2.2 Taxonomic checklist and occurrence dataset page 19
    2.3 Phylogenetic turnover across space page 20
    2.4 Optimisations of climate and geography page 21
    2.5 Phylogenetic turnover across time page 22
3. Discussion page 23
4. References page 26
5. Supplementary tables page 37
6. Supplementary figures page 73
Appendix 1 – Information about the checklist and occurrence data page 99

### 1. Methods

This study deploys large, global-scale, high-resolution taxonomic, phylogenetic and geographic distribution data for the pantropical Mimosoid clade of legumes with 3,469 species. These data are based on plant material collected during targeted fieldwork in key tropical biodiversity hotpots over the last two decades and leaf tissue sampled from specimens across the world's largest museum collections of plants. Large scale quality-controlled species occurrence data were also derived from museum collections worldwide, in the form of digitised herbarium specimen records. This study is therefore a prime example of the importance of museums for biodiversity research (*1*, *2*).

Unless stated otherwise, all data handling and analyses were performed in R (*3*).

#### 1.1 Phylogenetic inference

To construct a phylogeny that is both robustly supported and densely sampled in terms of taxa we combined a newly generated Hybseq backbone phylogeny with a set of 15 newly generated, taxonomically enhanced, or previously published species-level phylogenies for particular subclades. The backbone phylogeny was constructed via targeted enrichment of 997 nuclear genes selected specifically for phylogenomics of Mimosoid legumes (Mimobaits: (*4*), https://github.com/erikkoenen/mimobaits). For the backbone, data were generated for 420 species representing 147 of 152 genera across subfamily Caesalpinioideae and sampling taxa spanning the root nodes of each of the species-level subclades. This large dataset ensures that the backbone phylogeny is as robustly supported as possible and facilitates high-precision time-calibration using a subset of informative and clock-like genes. The 15 species-level phylogenies were constructed using a range of DNA sequence data types (Hybseq, RADseq and traditional Sanger sequenced loci), either newly generated or from published data, and appropriately rooted. Densely sampled ultrametric subtrees for these 15 subclades were grafted onto the time-constrained backbone phylogeny. We refer to the resulting time-calibrated phylogeny as a *meta-chronogram*, based on the earlier idea of meta-trees (*5*, *6*), as implemented by Spriggs et al. (*7*) to build a phylogeny for grasses. This approach means that diverse data types can be combined to build a single phylogeny, thereby tapping into the full wealth of available DNA sequence data and maximising the number of taxa that can be sampled with molecular data, and provides a computationally tractable way to build a large phylogeny with many taxa.

##### 1.1.1 Backbone topology

###### Taxon sampling, hybrid capture, and sequencing

We extracted DNA from herbarium specimens and silica-dried leaf samples of 284 taxa (Table S1) using the DNeasy Plant Mini Kit (Qiagen, Venlo, the Netherlands). DNA integrity and concentration of a subset of extractions from older herbarium specimens were checked on a 4200 TapeStation System using D5000 ScreenTape (Agilent Technologies, Santa Clara, United States of America). Library preparation, hybrid capture, enrichment, and sequencing were performed by Arbor Biosciences (previously Mycroarray; Ann Arbor, USA). For the capture, a custom bait set of 1Mb, targeting 997 nuclear genes, was used. This bait set was specifically designed for phylogenomic analyses of the Mimosoid clade of subfamily Caesalpinioideae (sensu (*8*)) of the legumes by Koenen et al. (*4*). To generate the bait set, a custom pipeline was used to select putatively single-copy genes from transcriptomes of four Mimosoid genera (*Albizia julibrissin*, *Entada abyssinica*, *Microlobius foetidus*, and three species of *Inga*) (*4*). During the capture reactions, 25 libraries were pooled based on approximate evolutionary distances between taxa. Sequencing was performed on Illumina HiSeq 4000, resulting in 150 bp paired-end reads. In addition to these newly-generated sequences, we also used the raw reads of the same genes from 122 Caesalpinioideae taxa previously generated by Koenen et al. (*4*) and 18 *Inga* species generated by Nicholls et al. (*9*). Together, this resulted in a dataset of 424 taxa, covering 150 of the 152 genera of subfamily Caesalpinioideae, and 89 of 90 Mimosoid genera (Table S1). The two missing genera are *Stenodrepanum*, the monospecific sister genus of *Hoffmannseggia* (*10*), and the Mimosoid *Microlobius,* which is also monospecific, and closely related to, or potentially nested within, *Stryphnodendron* (*11*, *12*). Finally, sequences for as many of the 997 genes as possible were extracted from published genomes of five taxa from subfamilies Cercidoideae and Papilionoideae (Table S2) using BLAST (*13*) and BLAT (*14*) to serve as outgroups for phylogenetic analyses.

#### Data cleaning and target assembly

Figure S1 presents an overview of this step. Duplicated reads were removed with FastUniq 1.1 (*15*). Reads were filtered with Trimmomatic 0.36 (*16*), to remove adapter sequences, leading and trailing bases with a quality score < 20, sliding windows of five bp with an average quality score < 20, and reads shorter than 51 bp. Finally, we used HybPiper 1.3.1 (*17*) to generate sequences for the 997 target genes. HybPiper uses BWA (*18*) to map the cleaned paired and unpaired reads to their targets (i.e., the DNA sequences of the bait set used for the capture reaction), SPAdes (*19*) for the *de novo* assembly of the mapped reads into contigs, and Exonerate (*20*) to extract coding sequences of the target genes. We ran HybPiper using default settings, except that the coverage cut-off of SPAdes was lowered to two, which increased the number and length of recovered genes for several samples with degraded DNA derived from old herbarium specimens. HybPiper yields two types of output. The main output consists of one DNA sequence for each taxon/gene combination. In addition, in cases where multiple sequences are recovered for particular taxon/gene combinations, these extra, potentially paralogous sequences are also stored.

#### Trimming outlier sequences and orthology assessment

Figure S2 presents an overview of this step. To remove outlier sequences and assess orthology of potentially paralogous sequences of genes recovered by HybPiper we used slightly edited versions of steps five and six of the Yang and Smith (*21*) pipeline. For this we divided the sequences of the 997 genes into two groups: single-copy and multi-copy genes. The multi-copy genes are those genes for which HybPiper recovered potentially paralogous sequences in at least 5% of the taxa; genes for which fewer or no potential paralogs were found were presumed to be single-copy. We applied Yang and Smith's (*21*) pipeline in different ways to these two subgroups (Figure S2).

To trim outlier sequences from the single-copy genes, we first split the sequences for each gene into two roughly equally-sized groups of taxa: a subclade of the ingoid clade (*4*) here referred to as the 'core ingoid' (CI) clade, i.e. all taxa descended from the most recent common ancestor of *Zapoteca* and *Zygia*, and the 'early-branching Caesalpinioideae' (EBC), i.e. all taxa not in the CI clade. Splitting the data in this way was done because, based on Koenen et al. (*4*) and preliminary analyses of our data, branch lengths in the CI clade tend to be significantly shorter than those outside this clade across the EBC. Removal of outlier sequences based on branch length therefore requires different settings and is more accurate if the CI and EBC taxa are analysed separately. For each group, codon alignments were produced with MACSE 2.01 (*22*), all sites with a column occupancy < 0.3 were trimmed using pxclsq (*23*), and gene trees inferred with RaxML 8.2.12 (*24*) using the GTRCAT model. Next, Yang and Smith's (*21*) trim_tips.py script was used to trim outlier long tips with a relative cut-off of 0.1 and absolute cut-offs of 0.15 for the CI sequences and 0.3 for the EBC sequences. An additional round of trimming outlier taxa was performed by running TreeShrink 1.3.3 (*25*) with a quantile of 0.1. Finally, long internal branches were cut using Yang and Smith's (*21*) cut_long_internal_branches.py script, cutting all internal branches with a cut-off of 0.3 and only keeping clusters with at least 22 taxa. At this point, the CI and EBC sequences were reunited into a single dataset for a second round of alignment with MACSE, tree-building with RaxML, removal of outlier long tips with trim_tips.py (with relative and absolute cut-offs of 0.1 and 0.3, respectively), and further removal of outliers using TreeShrink. Finally, outgroup sequences (if available) were added to the codon alignments with MACSE's enrichAlignment option, and codons with > 95% missing data were removed with BMGE 1.1 (*26*), resulting in final codon alignments with outgroups (if available) for all single-copy genes.

The multi-copy genes were analysed in two different ways, resulting in two different sets of final alignments for the same multi-copy genes (Figure S2). For the first set, potentially paralogous sequences were not included, and the multi-copy genes were analysed in exactly the same way as the single-copy genes (described above), resulting in codon alignments with outgroups (if available).

For the second set, potentially paralogous sequences were included, and a full orthology assessment following Yang and Smith (*21*) was performed (Figure S2). For this, the sequences of each gene were first split between the CI and the EBC. Then sequences for each of these two groups were aligned using MACSE, sites with < 0.3 column occupancy trimmed using pxclsq, gene trees inferred using RaxML, and outlier tips removed using Yang and Smith's (*21*) trim_tips.py (with absolute cut-offs of 0.15 and 0.3 for the CI and EBC, respectively) and TreeShrink (quantile 0.1). At this point, Yang and Smith's (*21*) mask_tips_by_taxonID_transcripts.py script was used to mask mono- and paraphyletic tips, after which internal branches > 0.3 were cut with cut_long_internal_branches.py, only keeping clusters with at least 22 taxa. This round of aligning, tree building, and trimming outliers was repeated for the CI and EBC separately, after which sequences of these two groups of taxa were reunited into a single dataset, for an additional round of aligning, tree building, and trimming outliers. However, before trimming outliers in this last round, all sequences that were flagged as potential paralogs by HybPiper for eight putative recent polyploids were removed. These putative polyploids, *Vachellia erioloba*, *V. farnesiana*, *V. nilotica*, *Sympetalandra schmutzii*, *S. unijugum*, *Mimosa tricephala*, *Schleinitzia insularum*, and *Leucaena trichandra*, were identified based on total numbers of potential paralogs recovered by HybPiper and analysis of preliminary gene trees, as well as, in some cases, earlier reports of polyploidy (*27*, *28*). Potential paralogs were removed because presence of these homeologous polyploid sequences would negatively affect the next step of orthology assessment following the maximum inclusion method (*21*) with the prune_paralogs_MI.py script, using relative and absolute cut-offs of 0.1 and 0.3 respectively, and only keeping clusters with at least 22 taxa. The resulting ortholog clusters were subjected to one final round of aligning with MACSE, trimming codons with > 95% missing data with BMGE, tree inference with RaxML, and removing outliers with trim_tips.py (relative and absolute cut-off 0.1 and 0.3) and TreeShrink (quantile 0.1), resulting in trimmed, final ortholog alignments.

At this point three groups of gene alignments were available for phylogenetic analyses (second-bottom row in Figure S2): the single-copy gene alignments (with outgroups, if available), alignments of the multi-copy genes treated as single-copy (with outgroups, if available), and ortholog alignments resulting from full orthology assessment of the multi-copy genes (without outgroups).

#### Phylogenetic analyses

We applied coalescent and concatenation approaches to infer several species trees using all three sets of gene alignments.

For the coalescent approach (Figure S3), all sequences shorter than 300 bp and at the same time shorter than half the total alignment length of the nucleotide alignments were trimmed. Using these cleaned alignments, final gene trees were inferred using RaxML with the GTRGAMMA model and 200 rapid bootstrap replicates. Based on Koenen et al. (*4*) and visual inspection of the resulting gene trees, all trees with a root-to-tip length variance over 0.009 were excluded from the dataset, in order to remove outlier gene trees that may be affected by alignment or orthology inference issues. Following Mirarab (*29*), branches with bootstrap support lower than 10% were collapsed. Gene trees were used to infer three separate species trees with ASTRAL-III 5.7.3 (*30*) (Figure S3): one using just the single-copy genes, one using the single-copy genes plus the multi-copy genes treated as single-copy (i.e., without paralogs), and one using the single-copy genes plus the multi-copy genes with paralogs.

We also used both maximum likelihood and Bayesian approaches to infer species trees based on gene alignments concatenated with pxcat (*23*). For the maximum likelihood analyses (Figure S4), we excluded alignments of loci of gene trees with a root-to-tip length variance higher than 0.009, and trimmed codons with > 90% missing data using BMGE. We produced six concatenated alignments: i.e. separate nucleotide and amino acid alignments of just the single-copy gene loci, of all gene loci including paralogs, and of all gene loci excluding paralogs. Six species trees were inferred with RaxML, using 200 rapid bootstrap replicates with the GTRCAT model (*31*) for the nucleotide alignments and the PROTGAMMALG4X model (*32*) for the amino acid alignments, two complex substitution models that allow for rate heterogeneity across sites. Finally, we used a gene jackknifing approach (following Koenen et al. (*4*)) to infer a Bayesian species tree (Figure S5). After excluding loci of gene trees with a root-to-tip length variance > 0.009 and trimming codons with > 90% missing data, we divided the amino acid single-copy gene alignments over 16 roughly equally sized concatenated alignments of circa 19,700 sites each. This gene jackknifing procedure was repeated four times, resulting in 48 unique jackknifes. These jackknifes were then used as input to PhyloBayes-MPI (*33*) using the CATGTR model (*34*, *35*) and 1000 cycles, the first 500 of which were discarded as burn-in. Convergence of the jackknife-runs was checked with the R (*3*) package CODA (*36*). Finally, all trees resulting from the 48 unique jackknifes were summarized as a majority-rule consensus tree.

#### Chloroplast phylogeny

A chloroplast phylogeny was inferred following the approach of Koenen et al. (*4*). Off-target reads were extracted using BLAST and three Caesalpinioideae reference chloroplast genomes: *Inga leiocalycina* ((*37*); GenBank accession KT428296), *Leucaena trichandra* ((*37*); GenBank accession KT428297), and *Erythrophleum fordii* ((*38*); GenBank accession MG644609). Coding sequences for each plastid gene were extracted using a custom Python script (*4*), aligned with MACSE, and concatenated with pxcat. The concatenated alignment contains 72 plastid genes, because two genes (*accD* and *clpP*) were removed following Koenen et al. (*4*). Finally, all taxa with > 95% missing data were trimmed from the alignment, and a species tree was inferred with RaxML using the GTRCAT model and 200 rapid bootstrap replicates.

#### Assessing gene tree conflict and topological congruence

Conflict among single-copy gene trees was assessed in several ways. Absolute numbers of the single-copy gene trees (with a root-to-tip length variance < 0.009) supporting and conflicting each bipartition in the single-copy genes ASTRAL tree were calculated using PhyParts (*39*). PhyParts was also used to calculate bipartition-based Internode Certainty All (ICA) values on the same tree. For both analyses, only gene tree nodes with > 50% bootstrap support were considered, following Smith et al. (*39*). Because bipartition-based calculations of support and conflict can be impacted by missing taxa in gene trees (*40*), quartet-based Extended Quadripartition Internode Certainty (EQP-IC) values were also calculated using QuartetScores (*40*).

Congruence between trees was expressed in Robinson-Foulds distances (*41*) calculated with phangorn's (*42*) 'treedist' function. Topological comparisons of all tree pair combinations were made using phytools' (*43*) 'cophylo' function.

ASTRAL's polytomy test (*30*) was used to assess the probability and locations of potential polytomies on the ASTRAL species tree based on the single-copy genes with a root-to-tip length variance < 0.009. Rogue taxa were identified by using RogueNaRok (*44*) to compare all six sets of nuclear RaxML bootstrap replicate trees to three reference trees (the relevant RaxML best tree, a strict consensus tree, and a majority rule consensus tree) with three dropset sizes (from one to three; analyses were repeated with each dropset size until no additional rogue taxa were identified), resulting in 54 unique RogueNaRok analyses.

##### 1.1.2 Time calibration

The ASTRAL single-copy genes topology of the Caesalpinioideae Hybseq backbone phylogeny was time-scaled in BEAST v. 1.8.4 (*45*, *46*) using a fixed local clock model with six local clocks to account for discrete branch length variation observed across the tree. From the Hybseq data set, 100 gene alignments were selected using SortaDate (*47*) as follows: (1) all genes with less bipartition correspondence to the ASTRAL topology than the median were discarded, (2) all genes with total tree length shorter than the median were discarded, and (3) the 100 most clock-like genes, i.e., those with the least root-to-tip length variance were selected. These 100 genes were concatenated and analysed in a single partition with a GTR + GAMMA model. To calibrate the tree, seven fossil calibrations were employed with minimum ages set as listed in Table S3, using uniform priors and maximum ages set at 66 Mya. Additionally, the root node (i.e. the crown node of the Leguminosae) was calibrated using a normal prior at 66 Mya, with a standard deviation of 0.1, as this was shown to be the likely approximate crown age of the family by Koenen et al. (*48*). The fossils and their minimum age calibrations are discussed in detail in Lavin et al. (*49*), Bruneau et al. (*50*), Simon et al. (*51*) (Supporting Information), and Koenen et al. (*48*) (Supplementary Appendix).

The analyses were carried out using two chains of 50 million generations each, from which subsequently 10,000 trees of each chain were sampled and used to estimate 95% HPD age intervals. A further subsample of 1,000 trees was used as the backbone set to create a set of 1,000 metachronograms, in combination with 1,000 post-burn-in trees of each subtree (see section 1.1.3).

##### 1.1.3 Metachronogram

As indicated above, in order to generate a densely sampled time-calibrated phylogeny we grafted a set of densely sampled phylogenies of subclades and genera (hereafter referred to as subtrees) onto the Hybseq time-calibrated backbone phylogeny described above. Because the backbone tree and the grafted subtrees were derived from data sets that employed different markers, both the backbone and the subtrees were estimated as ultrametric trees in order to make the branch lengths compatible. The subtrees were then re-scaled according to the corresponding node in the time-calibrated backbone onto which it was subsequently inserted. A python script to carry out this procedure is available at https://github.com/erikkoenen/metachronogram, and can be used to produce similar metachronograms for other clades where a backbone and subtree phylogenies with (potentially) non-overlapping markers are available.

Here we present a clade-by-clade description of the analyses to infer the subtrees. All analyses were carried out in MrBayes v. 3.2.1 (*52*) or BEAST v. 1.8.4 (*45*, *46*), using a GTR+GAMMA model, but with different clock models as specified below. For all analyses, convergence was assessed using Tracer v. 1.7 (*53*) and appropriate amounts of burn-in were discarded from the posterior samples. The subtree alignments and MrBayes execution files or BEAST xml files are available at https://github.com/erikkoenen/metachronogram, as well as example tree files that include 100 posterior trees for each subclade and the backbone.

*Acacia*

The *Acacia* subtrees were estimated from an alignment of 717 *Acacia* species and 7 outgroup taxa, using the nuclear internal and external transcribed spacers (ITS and ETS), and chloroplast markers *matK*, *psbA* and *trnL,* with a total aligned length of 6,407 nucleotide positions. This data set builds on that of Mishler et al. (*54*), but includes sequences for 210 *Acacia* species newly generated here, representing the largest phylogenetic tree of the genus to date. GenBank accession numbers for newly generated sequences are included in Table S4, for the other sequences see Mishler et al. (*54*). The analysis was carried out in BEAST, using the GTR+GAMMA model, with separate partitions for the nuclear and chloroplast data and an uncorrelated relaxed clock (lognormal). Four separate chains were run for 100 million generations each, after which the first 50 million generations were discarded as burn-in and 1,000 post-burn-in trees extracted. The outgroup taxa were pruned prior to grafting the trees onto the *Acacia* crown node in the backbone phylogeny.

Albizia clade

For the Albizia clade sensu Koenen et al. (*4*) we selected 32 informative and clock-like genes using the same SortaDate procedure described above for the backbone phylogeny, from an unpublished hybrid capture data set for the same Mimobaits gene set and probes used to construct the backbone tree. European Nucleotide Archive (ENA) accession numbers for these new data are included in Table S5. All accessions of the Albizia clade that were used in the backbone are also included here. A total of 79 accessions of *Albizia,* six accessions of *Enterolobium*, a single accession of *Leucochloron bolivianum* and seven outgroup accessions were sampled, for a total aligned length of 71,328 nucleotide positions. The ultrametric trees were inferred with MrBayes using an independent gamma rates (IGR) clock model with a uniform branch length prior, running for 50 million generations. The outgroup was pruned prior to grafting the posterior trees onto the most recent common ancestor (MRCA) node of *Albizia* + *Enterolobium* + *Leucochloron bolivianum* (i.e., the Albizia clade sensu Koenen et al. (*4*)) on the backbone phylogeny.

*Calliandra*

For *Calliandra* we used the alignment of Souza et al. (*55*) which includes 90 species of the genus plus 40 outgroup accessions, with a total aligned length of 2,170 nucleotide positions. Analyses were carried out using the same settings in MrBayes as for the Albizia clade, but running only 30 million generations as this was sufficient to reach convergence due to the shorter alignment length. The outgroup was pruned in the posterior trees prior to grafting these on the crown node of *Calliandra* on the backbone phylogeny.

Dichrostachys group

For *Alantsilodendron, Dichrostachys and Gagnebina* (i.e. the informal Dichrostachys group of Hughes et al. (*56*), excluding *Calliandropsis* for which the accession in the backbone was retained), we used a subset of a newly generated nextRAD RADseq data set, including six species of *Alantsilodendron*, 17 of *Dichrostachys* and five of *Gagnebina*, for a total aligned length of 34,030 nucleotide positions. The loci to be included were selected based on a minimum taxon occupancy of 75 % (i.e., a maximum of seven out of 28 taxa with only undetermined nucleotide positions). The analysis was carried out in MrBayes using the same settings as for the Albizia clade. An outgroup was not included but the subtrees were inferred with a constraint where the non-Madagascan species of *Dichrostachys* were enforced as the sister clade to the clade including the Madagascan species of *Dichrostachys* and the other two genera to make sure the trees were properly rooted. The subtrees were then grafted onto the MRCA node of the three genera on the backbone phylogeny.

*Inga*

We used a new Hybseq dataset (Nicholls et al. submitted) that builds on the data presented in Nicholls et al. (*9*) for the genus *Inga,* with additional taxon sampling to include a total of 162 accessions of the genus (including morphospecies and/or cryptic taxa that have not yet been formally described), with a total aligned length of 134,648 nucleotide positions. Accessions of *Zygia inundata* and *Z. sabatieri*, that are the sister taxa of *Inga*, from the backbone, were included in the subtree analysis to improve estimation of the branch length of the stem lineage and thus of the crown age of *Inga*. Analyses were carried out in BEAST using a fixed local clock model with a separate partition for a large clade within *Inga* which has discreetly higher substitution rates than the remainder of the genus. Three chains of each 50 million generations were run and the postburn-in posterior samples of these were combined. After pruning the single outgroup accession (*Zygia* sp. voucher Coley & Kursar *Tip917*), the subtrees were grafted onto the MRCA node of *Inga, Z. inundata and Z. sabatieri* in the backbone phylogeny.

Leucaena group

Subtrees for the clade composed of *Desmanthus, Kanaloa, Leucaena* and *Schleinitzia* (i.e., the informal Leucaena group of Hughes et al. (*56*)), were inferred from an alignment of 72 chloroplast genes extracted from off-target reads from the hybrid capture accessions used to build the backbone phylogeny, plus a new set of *Leucaena* transcriptomes (Table S6) and the *Desmanthus illinoensis* transcriptome of Cannon et al. (*57*). This dataset includes a total of five *Desmanthus*, one *Kanaloa*, 24 *Leucaena* and three *Schleinitzia* accessions, for a total aligned length of 53,788 nucleotide positions. The accessions from the backbone phylogeny of *Lemurodendron capuronensis* and *Neptunia oleracea* were used as the outgroup. The analysis was run in MrBayes with the same settings as for the Albizia clade, and after pruning the outgroup, the subtrees were grafted onto the MRCA of the four genera in the backbone phylogeny.

*Mimosa*

For the *Mimosa* subtrees we used an alignment of the chloroplast marker *trnD-trnT* that builds on the data sets of Simon et al. (*58*) and Vasconcelos et al. (*59*), plus 49 newly generated sequences, giving a total of 428 accessions from the genus plus 13 outgroup accessions, for a total aligned length of 2,704 nucleotide positions. GenBank accession numbers of the newly sequenced accessions are included in Table S7. The analysis was run in BEAST using the same settings as for the *Acacia* subtree analysis and after pruning the outgroup the subtrees were grafted onto the crown node of the genus in the backbone phylogeny.

Madagascan *Mimosa*

A separate set of posterior subtrees was inferred for the Old World clade of *Mimosa*, which is mostly confined to Madagascar, from a RADseq data set similar to that of *Alantsilodendron*, *Dichrostachys* and *Gagnebina*. However, since the Madagascan *Mimosa* data set has more missing data across the RAD loci, we used a cut-off of at least 40% taxon occupancy per locus to have sufficient total aligned length after concatenation. The alignment includes 29 accessions, with a total aligned length of 41,729 nucleotide positions. No outgroup was included, but instead the analysis was run with a constrained topology in which the non-Madagascan species were enforced as the sister clade to the clade of Madagascan species (previously shown to be monophyletic by Simon et al. (*58*)) to ensure correct rooting of the subtrees. The analysis was run in MrBayes using the same settings as for the Albizia clade and the subtrees were grafted onto the MRCA node of the Old World clade of *Mimosa* in the *trnD-trnT Mimosa* subtrees described above.

Parkia clade

To generate an alignment for the Parkia clade sensu Koenen et al. (*4*), we used the *Anadenanthera*, *Parkia* and *Vachellia* sequence data of Boatwright et al. (*60*) and added additional sequences from GenBank (see Table S8), including those of Comben et al. (*61*) for Australian *Vachellia*. In addition, sequences for *Albizia kalkora, Chloroleucon mangense, Lysiloma tergeminum, Neptunia monosperma* and *Piptadenia stipulacea* from Boatwright et al. (*60*) and *Senegalia senegal* of Terra et al. (*62*) were included as the outgroup. The alignment includes two species of *Anadenanthera,* 17 species of *Parkia* and 57 species of *Vachellia* and has a total aligned length of 4,444 nucleotide positions. The analysis was run in MrBayes using a strict clock, because using an IGR clock model caused the long stem lineages of the three genera in the Parkia clade to be substantially shortened, and with unrealistically high substitution rates inferred for these. The chain was run for 30 million generations. After pruning the outgroup the subtrees were grafted onto the crown node of the Parkia clade in the backbone phylogeny.

*Piptadenia*

We extracted the 13 accessions of *Piptadenia* from the alignment of Simon et al. (*12*), and *Mimosa myriadenia* to serve as the outgroup. These data include three chloroplast markers (*matK*/*trnK*, *trnD-trnT* and *trnL-trnF*) and nuclear ITS, with a total aligned length of 6,426 nucleotide positions when concatenated. The analysis was run with MrBayes using the same settings as for the Albizia clade, but running only 30 million generations. The outgroup was pruned and subtrees were grafted on the crown node of the genus in the backbone phylogeny.

*Senegalia*

We assembled an alignment of 70 *Senegalia* accessions plus *Parasenegalia vogeliana* as an outgroup accession by combining the data sets of Boatwright et al. (*60*) and Terra et al. (*62*) for the three chloroplast loci *trnK*/*matK*, *trnL-trnF* and *psbA-trnH*, with a total aligned length of 4,079. The analysis was run in MrBayes using the same settings as for the Albizia clade, but running only 30 million generations. Because *Senegalia* is non-monophyletic under its current circumscription (Ringelberg et al., submitted; Terra et al., in press) subtrees corresponding to the two separate clades of the genus were extracted and grafted separately onto their respective crown nodes in the backbone phylogeny.

Stryphnodendron clade

For the Stryphnodendron clade, we extracted sequence data for one accession of *Microlobius foetidus*, four accessions of *Parapiptadenia*, three of *Pityrocarpa*, five of *Pseudopiptadenia* and 23 of *Stryphnodendron* from the alignment of Simon et al. (*12*). *Inga edulis, Mimosa myriadenia, Piptadenia robusta, Lachesiodendron viridiflora* and *Senegalia nigrescens* were used as the outgroup. These data include three chloroplast markers (*matK*/*trnK*, *trnD-trnT* and *trnL-trnF*) and nuclear ITS, with a total aligned length of 6,426 nucleotide positions when concatenated. The analysis was run with MrBayes using the same settings as for the Albizia clade, but running only 30 million generations. The outgroup was pruned and subtrees were grafted on the MRCA node of *Parapiptadenia* and *Stryphnodendron* (i.e. the crown node of the Stryphnodendron clade sensu Koenen et al. (*4*)) in the backbone phylogeny.

Zapoteca clade

For the Zapoteca clade, sequence data for chloroplast locus *trnL-trnF* and nuclear ETS and ITS of Ferm (*63*) and Souza et al. (*64*) were combined to include *Faidherbia albida*, *Sanjappa cynometroides*, two species of *Thailentadopsis*, six species of *Viguieranthus* and 27 species and subspecies of *Zapoteca*, with a total aligned length of 2,759 nucleotide positions. Sequence data from Ferm (*63*) for *Calliandra dysantha, C. surinamensis, Havardia mexicana, H. pallens, Pithecellobium diversifolium* and *P. dulce* were included as the outgroup. The analysis was run in MrBayes using the same settings as for the Albizia clade, but running 30 million generations. After pruning the outgroup, the subtrees were grafted onto the crown node of the Zapoteca clade in the backbone phylogeny.

*Zygia*

For *Zygia*, we used data of Ferm et al. (*65*) to include 41 species and varieties of *Zygia* sensu stricto, and eight outgroup species (including *Z. inundata, Z. ocumarensis*, and *Z. sabatieri* that were shown by Ferm et al. (*65*) to be more closely related to *Inga* and *Macrosamanea*), with a total aligned length of 3,228 nucleotide positions. The analysis was run in MrBayes using the same settings as for the Albizia clade, but running only 30 million generations. After pruning the outgroup, the subtrees were grafted onto the crown node of *Zygia* in the backbone phylogeny.

#### 1.2 Taxonomic checklist and species occurrence dataset

##### Mimosoid clade

In order to assemble an accurate quality-controlled species occurrence dataset, we first compiled a comprehensive taxonomic checklist of accepted names with partial synonymy for all species and infraspecific taxa in the Mimosoid clade. For each genus or clade, the most recent taxonomic monograph or revision (when available) was used, and more recently described taxa were added using the International Plant Names Index (IPNI; www.ipni.org). This checklist was used to download species occurrence records from the Global Biodiversity Facility (GBIF; www.gbif.org), the Latin American Seasonally Dry Tropical Forest Floristic Network (DryFlor; www.dryflor.info), and the Southwestern Environmental Information Network (SEINet; http://swbiodiversity.org/seinet), as well as various other taxon- or region-specific sources. Extensive data cleaning was performed: we updated synonymous names using the checklist, removed records not based on vouchered herbarium specimens (except for DryFlor records based on plot data or checklists), records located in the sea or on country or major area centroids, cultivated records, and records located outside native distribution ranges as delimited in the primary taxonomic literature.

For some taxa, pre-compiled occurrence datasets were used. For example, for the vast majority of the largest Mimosoid genus, the predominantly Australian genus *Acacia* which comprises circa 1,000 species, we did not assemble a custom checklist and occurrence data but instead relied on the occurrence dataset of González-Orozco et al. (*66*) with minor updates (see Appendix 1 for details).

Literature used to assemble the taxonomic checklist and perform quality control of the occurrence data, as well as GBIF DOIs, taxon-specific notes, and other sources of occurrence data, are in Appendix 1.

###### Non-Mimosoid Caesalpinioideae

For the optimisations of climate and geography across the metachronogram (see section 1.4) the 77 outgroup taxa across Caesalpinioideae outside the Mimosoid clade in the phylogenomic backbone were included to improve the accuracy of ancestral area and climate reconstructions across the early nodes of the Mimosoid clade. We assembled an occurrence dataset of these 77 non-Mimosoid Caesalpinioideae taxa present in the phylogeny, partly by using published datasets (*67*, *68*) and partly by downloading and cleaning new occurrence data in the same way as for the Mimosoid clade (see above). For each outgroup taxon, literature, GBIF DOIs, data sources, and notes are presented in Appendix 1.

###### Abundance of Mimosoids across biomes

We estimated the fraction of Mimosoids across biomes and continents using several datasets.

The fraction of Mimosoid species among all Amazonian tree species was estimated in two ways: using Appendix S1 of ter Steege et al. (*69*) and dataset S01 of Cardoso et al. (*70*). For the fraction of Mimosoid species among Neotropical dry forest trees, the occurrence dataset of DRYFLOR (*71*) (downloaded from http://www.dryflor.info/data/datasets) was used. The fraction of Mimosoid species among African savanna trees was assessed using Appendix S1 of Fayolle et al. (*72*). Finally, the fraction of Mimosoid species among all native Australian Angiosperms was calculated using a species checklist downloaded from the Australasian Virtual Herbarium (https://avh.chah.org.au/). For this checklist, only Angiosperms native to Australia were considered.

#### 1.3 Phylogenetic turnover across space

###### Phylogenies

Phylogenetic turnover was calculated using the maximum clade credibility (MCC) tree of the new metachronogram. All non-Mimosoid taxa, multiple occurrences of the same taxon, and taxa lacking occurrence data were removed, leaving a total of 1,940 unique taxa in the tree, including infraspecific taxa.

As a robustness test, we repeated all spatial turnover analyses using a genus-level Mimosoid phylogeny. The ultrametric Caesalpinioideae phylogeny was used as a backbone topology, from which all non-Mimosoid taxa were removed and Mimosoid species added as genus-level polytomies, resulting in an ultrametric Mimosoid phylogeny with 3,165 species, i.e. almost all Mimosoid species (a few missing species could not be placed with confidence due to generic non-monophyly). This phylogeny does not include infraspecific taxa, and for analyses using this tree occurrence data for infraspecific taxa were amalgamated into their inclusive species. Thus, compared to the metachronogram, this genus-level tree used for the robustness test has a higher number of species but lacks resolution within genera.

##### Phylogenetic turnover

Phylogenetic turnover across the global tropics was quantified using the phylogenetic version of Simpson's pairwise dissimilarity index (*73*–*75*). Also known as (phylo)beta-sim or βsim (*73*, *76*), Simpson's dissimilarity index quantifies the 'true' turnover component of Sørensen's dissimilarity index (*73*, *74*), and has the important advantage of not being influenced by differences in species richness between sites (*73*, *74*, *76*, *77*).

The betapart R package (*78*) was used to calculate pairwise phylogenetic turnover between all half-by-half degree longitude/latitude grid cells with at least three (metachronogram) or five (genus-level tree) taxa located 33 degrees latitude north or south of the equator, i.e. restricting assessment of phylogenetic turnover patterns to the global tropics and subtropics. Temperate regions were excluded because Mimosoids show high tropical niche conservatism (*79*): only 31 taxa (i.e., 0.83% of all taxa in the occurrence dataset), belonging to *Prosopis* (15 taxa), *Mimosa* (six taxa), *Desmanthus* (three taxa), *Albizia*, *Prosopidastrum*, and *Senegalia* (each two taxa), and *Calliandra* (one taxon), have over half their occurrence points north or south of 33 degrees latitude in North or South America, Africa, the Middle East, or Asia, and these climatic and geographic outliers are likely to strongly bias explanations of patterns of phylogenetic turnover in these regions. The only exception to this is Australia, where 288 taxa of *Acacia* and one taxon of *Paraserianthes* (i.e., 7.70% of all Australian taxa) have over half of their occurrence points south of 33 degrees latitude, and phylogenetic turnover was therefore quantified across all of Australia, including Tasmania.

As a robustness test, phylogenetic turnover was also calculated using one-by-one degree longitude/latitude grid cells. The result was used in all spatial turnover analyses described below.

##### Predictor variables

Geographic (great circle) distances between centres of grid cells were calculated using the 'RdistEarth' function of the fields package (*80*), or internally in the 'gdm' function of the gdm package (*81*).

Climatic distances between grid cells were calculated using all 19 Bioclim variables of CHELSA (*82*), cloud cover and the intra-annual standard deviation of cloud cover downloaded from EarthEnv.org (*83*), plus dry season length, i.e. the number of consecutive months with rainfall < 100 mm/month (*67*). Predictor variables were aggregated to a half degree resolution using the 'aggregate' function of the raster package (*84*).

##### Explaining phylogenetic turnover

Generalised dissimilarity modelling (GDM) provides a powerful technique to analyse and explain patterns of spatial turnover (*85*, *86*). It offers several advantages over other approaches, including fitting non-linear relationships between turnover and predictor variables (*85*–*87*), which provides a more realistic way to assess complex biological patterns than strictly linear relationships (*88*). GDM also allows precise quantification of the effect of each individual predictor variable on turnover patterns (*87*, *89*). GDMs were run using the 'gdm' function of the gdm package (*81*) with default settings.

**Hypothesis testing**

Hypotheses were tested using variation partitioning (*89*–*92*), whereby, in order to tease apart the unique and combined effects of hypothetical predictors A and B, we compared the effects of GDMs run with predictors A and B together, just with A, and just with B.

To tease apart the influence of dispersal limitation and phylogenetic niche conservatism, we quantified the fraction of global phylogenetic turnover explained by geographic distance and by climatic predictors. This was done on a pantropical scale (excluding temperate areas) and at the level of individual continents: North America (including Central America and the Caribbean), South America, Africa (including Madagascar and Arabia), Asia, and Australia. Exploratory analyses show that using other definitions of continents, e.g. Africa without Madagascar and Arabia, and Australia without its temperate regions, does not have a large impact on the results (results not shown).

Next we ran a simple linear regression of phylogenetic turnover with geographic distance. The residuals of this model, hereafter referred to as geographic residuals, represent the fraction of phylogenetic turnover that is not explained by spatial distance (*93*, *94*). Using these geographic residuals ensures that we are not mistakenly assigning environmental explanations to phylogenetic turnover patterns that are actually driven purely by spatial distance (*95*). Geographic residuals were used for two additional rounds of variation partitioning, at both pantropical level and at the level of the five individual continents: quantifying the fraction of phylogenetic turnover explained by mean annual precipitation (Bio12) and annual mean temperature (Bio1), and quantifying the fraction explained by mean annual precipitation (Bio12) and seasonality of precipitation, quantified either as CHELSA's Bio15 ('precipitation seasonality'), EarthEnv.org's intra-annual standard deviation of cloud cover, or dry season length.

**Ancient turnover**

Although our aim is to assess turnover of lineages rather than turnover of taxa, taxonomic turnover and phylogenetic turnover are strongly correlated in Mimosoids (Tables S9 and S10) and other taxonomic groups (*74*, *92*, *93*, *96*–*98*). To overcome this issue, we followed Daru et al. (*99*) and McFadden et al. (*100*) and calculated phylogenetic turnover at deeper levels in the MCC metachronogram by collapsing all branches younger than a certain threshold, using the scripts provided by Daru et al. (*99*). We calculated ancient turnover at three thresholds: 5, 10, and 20 million years ago, and then performed variation partitioning to assess the influence of phylogenetic niche conservatism and dispersal limitation (see above).

**Phylogenetic regionalization**

Phyloregionalization or bioregionalization analysis provides a way to visualize and map patterns of spatial phylogenetic turnover purely based on the phylogeny and occurrence dataset, and independent from climatic or other types of data (*66*, *76*, *101*). A phylogenetic regionalization analysis (*76*, *77*, *101*) was performed using the Ward clustering algorithm implemented in the ‘hclust’ function of the stats package (*3*), which divided grid cells into between two and eight clusters based on their phylogenetic beta diversity. Clustering was performed on pantropical and continent levels, using full phylogenetic turnover, ancient turnover, and geographic residuals.

To investigate whether the resulting clusters are climatically different from each other, we tested whether the mean annual precipitation (Bio12), precipitation seasonality (Bio15), and dry season length values of all cells making up a cluster are significantly different from the climatic values of other clusters. This was done using the Wilcoxon rank sum test (‘wilcox.test’ function of the stats package (*3*)) for comparisons of two clusters, and the Kruskal-Wallis rank sum test (‘kruskal.test’ of the stats package) for comparisons of more than two clusters. In case of a significant outcome of the Kruskal-Wallis test, Dunn’s multiple comparison test with Bonferroni adjustment for multiple comparisons (using the ‘dunnTest’ function of the FSA package (*102*), which relies on the dunn.test package (*103*)) was used to identify the number of climatically distinct clusters.

#### 1.4 Optimisations of climate and geography

##### Optimisation of climate

Precipitation was optimised across the MCC tree of the metachronogram using the ‘contMap’ function of the phytools package (*43*). Two independent optimisations were performed, using the median values per species of mean annual precipitation (Bio12) and dry season length. To increase the accuracy of the optimisation especially at deeper levels in the Mimosoid tree, data of the 77 non-Mimosoid Caesalpinioideae outgroup taxa in the phylogeny (see section 1.2) were included in the analyses. While the outgroup sampling is less dense than sampling within the Mimosoid clade, the 77 outgroup taxa provide a reasonable representation of the climatic and geographic distributions of the non-Mimosoid Caesalpinioideae, and include taxa belonging to 58 of all 62 non-Mimosoid Caesalpinioideae genera.

Once the optimisation was finished, all nodes and tips outside the Mimosoid clade were removed from the tree. Nodes were then divided into three rainfall regimes following Neves et al. (*104*): dry (< 1200 mm/year), wet (> 1800 mm/year), and intermediate (between 1200 and 1800 mm/year). These categories were used to calculate the number, location, and age of niche shifts, with niche shifts defined as changes in rainfall regime (i.e., between dry, wet, and intermediate) that also incorporate a change in mean annual precipitation of at least 250 mm, optimised on a single branch. Niche shifts are assumed to have taken place on the middle of a branch, i.e. their age is the midpoint of the ages of the parent and child nodes.

For the optimisation of the dry season length, nodes were also divided into three regimes: dry (> 8 months), wet (< 4 months), and intermediate (between 4 and 8 months). Niche shifts were defined as changes in rainfall regime that also incorporate a change in dry season length of at least one full month.

##### Phylogenetic signal

Phylogenetic signal, expressed as Pagel’s lambda (*105*), was calculated using the ‘phylosig’ function in the phytools package (*43*), the MCC tree of the metachronogram, and median values of all Mimosoid species of mean annual precipitation (Bio12) and dry season length. Significance was assessed using a hypothesis test (option ‘test’ of ‘phylosig’).

##### Optimisation of geography

Geography was optimised across the MCC tree of the metachronogram, including non-Mimosoid Caesalpinioideae outgroups (see above), using BioGeoBEARS (*106*). To do so all taxa in the metachronogram were assigned to eight possible areas using their distribution data: North America, South America, Africa, Madagascar, Asia, Australia, Oceania, and the European Mediterranean. Taxa were allowed to occupy more than one area.

The BioGeoBEARS R package (*106*–*108*) was used to fit six different models: DEC, DEC+J, DIVALIKE, DIVALIKE+J, BAYAREALIKE, and BAYAREALIKE+J, and Akaike's corrected information criterion was used to select the best-fitting model.

After removing nodes representing non-Mimosoid Caesalpinioideae outgroup taxa from the BioGeoBEARS output, the number, location, and age of trans-oceanic dispersal events was assessed. To do so, reconstructed combined states at all internal nodes were simplified to the eight primary regions. Next, to exclude dispersal events between North and South America, these two areas were merged across all nodes and tips. Then the most probable range of each internal node was determined as the combination of all the most probable areas that together have a probability > 50%. Trans-oceanic dispersal was assumed to have taken place if the most probable areas of the child node are not all included among the most probable areas of the parent node, or if not all areas of a taxon (i.e., a tip) are included among the most probable areas of the parent node. As with niche shifts, dispersal events are assumed to have taken place on the middle of a branch, i.e. their age is the midpoint of the ages of the parent and child nodes.

As a robustness analysis, dispersal events were recalculated using a stricter definition of trans-oceanic dispersal, by combining the eight areas into three: North and South America, Africa-Madagascar-Asia-Australia-Mediterranean, and Oceania.

#### 1.5 Phylogenetic turnover across time

To explore scenarios of lineage turnover through time in relation to Cenozoic climate cooling that led to an increase in dry habitats across the planet (*109*), we ran a set of analyses with BAMM (*110*), using the MCC tree of the metachronogram set with the outgroup removed except for *Erythrophleum* and *Pachyelasma* (the sister group of the Mimosoid clade) in order to include the stem lineage of the Mimosoid clade. This method has been criticized (*111*, *112*) but the author of the program has responded to these criticisms (*113*, *114*) and, more generally, any diversification rate estimation method suffers from a lack of power to estimate extinction rates (*115*), meaning also that speciation rates are not identifiable from phylogenies (*116*). Therefore, we ran analyses across a wide range of fixed extinction rates to assess how speciation rates would vary through time under various levels of turnover, while making use of the powerful way in which BAMM can take unsampled diversity into account by assigning sampling fractions to genera or clades, which we estimated based on our Mimosoid checklist and taxonomic expertise. Priors were set using the setBAMMpriors option of the BAMMtools R package (*117*), extinction rates were fixed across different analyses at 0.05 lineages/Mya and ranging from 0.5 to 3.5 with intervals of 0.5. Speciation rates and rate shifts (or more accurately, shifts to different speciation rate regimes as the model also includes time-variable speciation rates) were left as free parameters to be estimated during the analysis. BAMM was then run for 10 million generations while saving parameters every 1,000 generations. Phylorate plots and rate-through-time plots were drawn using the BAMMtools R package (*117*).

The table with sampling fractions has been made available as online supplementary data to this article.

### 2. Results

Only results not included in the main text or in main or supplementary figures or tables are presented here.

#### 2.1 Phylogenetic inference

##### 2.1.1 Backbone topology

###### Excluded taxa

Four taxa were excluded from the final phylogenetic analyses: *Calliandra umbrosa*, *Vouacapoua americana*, *Pterogyne nitens*, and *Albizia subdimidiata var. subdimidiata*. The sequencing of *C. umbrosa* was unsuccessful, generating no reads that could be used in the analyses, probably because of low DNA quality and yield (circa 0.01 µg, the lowest yield of all samples) from poor quality herbarium material. While the sequencing of *V. americana* was more successful, only a few genes could be recovered for this taxon (Table S1), and the vast majority of these were removed as outliers during data cleaning, leaving too few genes to be confident about the phylogenetic placement of this taxon. *Pterogyne nitens* and *Albizia subdimidiata var. subdimidiata* were placed in the species trees, but their placements were highly unexpected, and inspection of gene trees and alignments heavily implies sequences of these two taxa were affected by contamination. They were therefore removed from all analyses.

Additionally, in every phylogeny *Hultholia mimosoides* was found to be nested within *Entada*, a placement that is clearly incorrect, as previous studies have shown *Hultholia* to be a member of the distantly-related Caesalpinia group (*10*, *68*). Although *H. mimosoides* should clearly not be placed in the Mimosoid clade, its highly divided bipinnate leaves are nevertheless Mimosoid-like (hence its specific epithet), and its geographic distribution overlaps with that of several *Entada* species (i.e., *E. glandulosa*, *E. phaseoloides*, *E. reticulata*, and *E. rheedii*). We therefore suggest that a sample or identification mix-up has resulted in sequencing DNA from an unidentified *Entada* species, instead of *H. mimosoides*, and have renamed this sample '*Entada sp.* van Beusekom et al. 4706'.

With the exclusion or renaming of these five taxa, samples of 420 ingroup taxa belonging to 147 of 152 Caesalpinioideae genera are included in the final phylogenies, as well as five outgroup taxa. This means that only five Caesalpinioideae genera are not included in the phylogenies: *Stenodrepanum* and *Microlobius* (not sequenced), and *Hultholia*, *Vouacapoua*, and *Pterogyne* (excluded from phylogenetic inference).

###### Sequencing results and data quality

Sequencing, data cleaning, and target assembly results per accession are presented in Table S1.

Fractions of duplicated reads, as determined by FastUniq (*15*), were moderately high, probably reflecting the large number of samples derived from old herbarium specimens with partially degraded DNA (Figure S6): TapeStation analyses showed that many DNA extractions yielded highly-fragmented DNA (average fragment sizes of circa 1000 bp) in low quantities (often < 1 µg) (results not shown). Fractions of reads filtered by Trimmomatic (*16*), in contrast, were low (Figure S7). The target capture was successful, as shown by the high percentages of reads on target (Figure S8) and the high numbers of genes that were recovered with at least 75% of the target length (Figure S9) by

HybPiper (*17*). Our results therefore confirm previous findings that herbarium specimens can be rich sources of DNA for hybrid capture studies (*1*, *118*, *119*). The hybrid capture was less successful with lower percentage of reads on target in most samples from Koenen et al. (*4*) in part due to the significantly larger bait set used for the hybrid capture reaction in that study. For most samples, additional, potentially paralogous sequences were only recovered for a small number of genes, but in a small minority of samples significantly higher numbers of genes appear to be duplicated, suggesting that these taxa are possible polyploids (Figure S10).

Of the 997 target genes, 167 were treated as multi-copy (i.e., potentially paralogous sequences were present in > 5% of taxa), and the remaining 830 genes were deemed to be single-copy (Figure S2). Outgroup sequences were obtained for 918 genes (Table S2, Figure S2). Alignment lengths varied between 117 and 7,752 nucleotide sites, with a median length of 843 (Figure S11). Numbers of taxa (including outgroups) per alignment varied between 18 and 426 (Figure S12). The much lower number of taxa in the alignments of the multi-copy genes with paralogs (median number of taxa: 42) compared to alignments of the single-copy genes (median: 409) and multi-copy genes without paralogs (median: 412), is caused by the orthology inferences for the former, which cut up larger gene trees into smaller, orthologous clusters (*21*).

**Phylogenetics**

Only a handful of gene trees had relatively high root-to-tip variances (i.e., > 0.009; Figure S13), namely nine out of 830 single-copy gene trees, three out of 169 multi-copy gene trees without paralogs, and two out of 667 multi-copy gene trees with paralogs, and these were excluded from further analyses. Concatenated alignments of the remaining loci were large, with many informative sites and relatively few gaps (Table S11, Figure S4).

A total of ten nuclear species trees were produced: three made with ASTRAL (Figure S3: Figures S14 – S16), six with RAxML (Figure S4: Figures S17 – S22), and one with PhyloBayes (Figure S5: Figure S23). All nuclear trees are robustly supported and largely congruent with each other (Figures S24 – S27), as evidenced by high topological similarities between trees (Table S12, Figure S28). The chloroplast phylogeny (Figure S29) has a significant number of missing taxa (i.e., it contains 383 taxa, versus 427 in the nuclear phylogenies) and shows more conflict with the nuclear trees than there is among the different nuclear estimates.

Gene tree conflict is ubiquitous in Caesalpinioideae: not a single bipartition in the single-copy ASTRAL phylogeny is supported by all 821 single-copy gene trees (Figures S30 and S31). Approximately one third of the nodes in the species tree are supported by at least half of all gene trees, and over half of species tree nodes are supported by at least a quarter of the gene trees. In contrast, for half of the species tree bipartitions there are more conflicting than supporting gene trees. In general there are many different conflicting topologies among the gene trees for each node, rather than a few abundant ones, indicating that gene tree conflict is mainly caused by lack of signal rather than common alternative topologies (*4*). Lack of signal in the gene trees is also directly evident given that for circa 20% of species tree nodes the majority of gene trees do not provide any information, be it in support or conflict. Some parts of the tree are especially prone to gene tree conflict, including along the ingoid backbone, the Madagascan clades in *Albizia* and *Dichrostachys* – *Alantsilodendron*, the Australasian clade comprising *Acacia*, *Archidendron*, and several smaller genera, and across the recently diversified largely Amazonian rain forest genera *Macrosamanea*, *Zygia*, and *Inga*. Gene tree conflict is also indicated by the two types of internode certainty values, ICA and EQP-IC, which are strongly correlated with each other (Pearson’s correlation coefficient 0.85), although on average the quartet-based EQP-IC scores are lower than the bipartition-based ICA values (Figure S32), as found by Zhou et al. (*40*).

ASTRAL's polytomy test shows the locations of several potential hard polytomies in the ASTRAL single-copy phylogeny (Figure S33), which closely coincide with nodes and clades associated with considerable gene tree conflict (Figures S30 – S32). The polytomies fall into two categories. The first category relates to the position of several difficult-to-place individual taxa, such as *Cylicodiscus gabunensis*, *Albizia leonardii*, and *Cedrelinga cateniformis*, while the second category comprises polytomies at the base of or within clades that most likely represent rapid radiations, such as along the ingoid backbone, in the Madagascan clades of *Albizia* and *Dichrostachys* – *Alantsilodendron*, within the Australasian clade, and within and among the recently diversified rain forest genera *Macrosamanea*, *Zygia*, and *Inga*. Several taxa in these clades are often identified by RogueNaRok as rogue taxa (Table S13), based on their variable placements in the RAxML bootstrap trees, indicating the difficulty, or perhaps impossibility, of resolving and representing the relationships resulting from rapid radiations as fully bifurcating topologies.

In the rest of this study, the single-copy ASTRAL phylogeny (Figure S14) is used as the 'reference' Caesalpinioideae phylogeny. There are several reasons to pick this particular tree. First, due to the much larger size of the nuclear dataset based on large numbers of independent loci, the high fraction of missing data and taxa in the chloroplast alignment, and the low bootstrap support values and short branches across parts of the plastid phylogeny, we consider the nuclear trees to provide a more reliable estimate of the species tree than the chloroplast phylogeny. Second, by selecting a phylogeny based on only the single-copy genes, the effects of the (absence of) orthology assessment, although minor, do not have to be considered. Furthermore, whether expressed in absolute gene numbers (Figure S30) or bipartition- or quartet-based internode certainty values (Figure S32), there is extensive conflict among individual gene trees, which violates the central assumption of the concatenation model (*120*). Finally, the multi-species coalescent model has been shown to consistently outperform the concatenation model on a range of phylogenomic datasets (*120*), which strongly suggests that the ASTRAL trees provide the most accurate approximation of the Caesalpinioideae species tree.

The eleven nuclear and plastid Caesalpinioideae phylogenies have been made available as online supplementary data for this article.

##### 2.1.2 Time calibration

The time-calibrated Caesalpinioideae phylogeny is shown in Figure S31.

A time-calibrated tree file and the BEAST xml file, which includes the alignment of the 100 clock-like genes used for dating, have been made available as online supplementary data for this article.

##### 2.1.3 Metachronogram

The Mimosoid clade in the maximum clade credibility (MCC) tree of the metachronogram is shown in Figure 1. Figure S34 depicts the same tree, with all added subtrees labelled and highlighted.

The 1000 metachronograms and the MCC tree with age intervals and posterior probabilities have been made available as online supplementary data for this article. Scripts for metachronogram analyses and the alignments of the subtrees are available on https://github.com/erikkoenen/metachronogram.

#### 2.2 Taxonomic checklist and occurrence dataset

##### Mimosoid clade

Based on the taxonomic checklist assembled here (Appendix 2), the Mimosoid clade contains at least 3,469 accepted species and 680 infraspecific taxa (Table S14). A further 6,155 synonyms are associated with these taxa focusing on recent synonyms that are likely to have been used on herbarium specimen determinations.

Using the taxonomic checklist, we assembled a dataset with quality-controlled occurrence records for 3,233 Mimosoid species and 522 infraspecific taxa, i.e. 93% of species and 91% of all taxa in the Mimosoid clade (Table S14). All 90 Mimosoid genera are represented in this dataset. This cleaned occurrence dataset contains 424,333 records, with a mean of 113 and a median of 33 records per taxon. Taxon richness quantified as the number of taxa per half degree longitude/latitude grid cell (Figure S35) implies that the Mimosoid clade is the most diverse in the Neotropics and Australia. However, generic richness (Figure S35) shows that this conclusion is strongly affected by the hyperdiverse genus *Acacia*, as large parts of Australia are characterised by high numbers of *Acacia* taxa but very low numbers of other Mimosoid genera. Furthermore, under-representation of many African regions in online repositories of occurrence data (*121*) also introduces sampling bias, revealed by the fact that many African cells have low taxon richness but relatively high genus richness (Figure S35). In Asia, the low taxon and genus richness is attributable to the relatively low number of Asian Mimosoid lineages and the limited availability of data, especially from the Indian subcontinent.

Figure 1e shows Mimosoid growth form variation across the tropical precipitation gradient. From left to right, the following species are shown:

- *Prosopis tamarugo*, in the hyper arid Pampa del Tamarugal (northern Chile). Photographer: Oliver Whaley;
- *Vachellia tortilis*, a typical 'umbrella thorn Acacia' of African savannas, in Serengeti National Park (Tanzania). Photographer: Robur.q (https://commons.wikimedia.org/wiki/File:Vachellia_(ex_Acacia)_tortilis.jpg, last visited on 31/05/2022);
- *Mimosa pumilio*, a geoxyle in the Brazilian Cerrado. Photographer: Marcelo F. Simon;
- *Entada rheedei*, a liana in humid tropical forest, Mozambique. Photographer: Bart T. Wursten (www.mozambiqueflora.com) (*122*);
- *Parkia pendula*, in tropical rainforest in Brazil. Photographer: Marcelo F. Simon.

##### Non-Mimosoid Caesalpinioideae

The occurrence dataset of the non-Mimosoid Caesalpinioideae contains 18,346 records of all 77 Caesalpinioideae outgroups in the phylogeny, with a mean of 238 and a median of 109 records per taxon.

The occurrence datasets of the Mimosoids and the non-Mimosoid Caesalpinioideae present in the phylogenomic backbone have been made available as online supplementary data for this article.

##### Abundance of Mimosoids across biomes

Mimosoids represent 5.4% of all tree species in the Amazon according to ter Steege et al. (*69*) and 4.5% following Cardoso et al. (*70*), 7.7% of all tree species in Neotropical dry forests (*71*), 17.3% of all African savanna tree species (*72*), and 5.4% of all native Australian Angiosperm species.

#### 2.3 Phylogenetic turnover across space

##### Robustness tests

The GDMs, variation partitioning and phyloregionalization results obtained using the metachronogram and the genus-level tree used as a robustness test are quantitatively and qualitatively highly similar. The main metachronogram results are presented Figures 2 and 3 and Tables S9 and S15. Figures S36 – S41 show metachronogram phyloregionalization results per region with different numbers of clusters, as well as those based on the geographic residuals of phylogenetic turnover and ancient turnover. Figure S42 presents the metachronogram phyloregionalization results with a global distribution of isohyets. The climatic distinctiveness of the metachronogram phyloregions is shown in Figure S43.

Phyloregionalization and variation partitioning results obtained using the genus-level Mimosoid phylogeny are presented in Figures S44 and S45 and Tables S10 and S16.

For all phyloregionalization figures, the underlying map is the ‘countriesHigh’ world map of the rworldextra package (*123*).

Analyses performed at the coarser spatial scale of one-by-one degree grid cells yielded highly similar results to analyses performed at a half-by-half degree scale, and are therefore not shown there.

#### 2.4 Optimisations of climate and geography

##### Optimisation of climate

Estimated niche shifts through time are highly similar regardless of whether mean annual precipitation (Figure 1) or dry season length (Figure S46) are used to calculate ancestral niches. Figure 1b shows the fraction of niche shifts per speciation event per five million years averaged across both optimisations, whereas the inset in Figure S46 shows the same just based on shifts in dry season length niches.

##### Optimisation of geography

The DEC+J model was recovered as the best-fitting model for the data, with the DEC model recovered as second-best. However, the DEC+J model, the DEC model, and the jump (+J) parameter have been criticised on theoretical grounds (*124*). To prevent drawing conclusions based on a single, possibly flawed model, we therefore inferenced ancestral distributions using four different models (DEC+J, DEC, DIVALIKE, and BAYAREALIKE), and estimated numbers of dispersal events with two different definitions of trans-oceanic dispersal (with seven regions (North and South America combined, Africa, Madagascar, Asia, Australia, Oceania, and the Mediterranean) and three regions (North and South America combined, Africa, Madagascar, Asia, Australia, and the Mediterranean combined, and Oceania)), resulting in a total of eight estimates.

Fractions of dispersal events through time (Figure 1b, Table S17) were averaged across all eight estimates. Dispersal events mapped onto the metachronogram (Figure 1a) are those inferred by the DEC+J model with seven regions. Ancestral states across Caesalpinioideae, based on the DEC+J model, and inferred trans-oceanic dispersal events in the Mimosoid clade, based on the model with seven regions, are shown in Figure S47.

#### 2.5 Phylogenetic turnover across time

The BAMM results are presented in Figures S48 and S49. Figure S48 shows the distribution of speciation rates across the metachronogram under different extinction scenarios. The top subfigure in Figure S49 shows the median speciation rate across the phylogeny in 100 time bins under the same extinction scenarios. Paleotemperatures in the middle figure are based on Zachos et al. (*125*), and were obtained using RPANDA (*126*). The phenogram was plotted using the 'phenogram' function in phytools (*43*).

An important limitation of the BAMM approach used here is that extinction rates are held constant through time. Under a model of episodic turnover, which we hypothesize (see main text), extinction rates are also likely to vary substantially through time, with relatively high levels of extinction likely during times of rapid historical climate change, and in particular, at the Eocene-Oligocene transition.

### 3. Discussion

Only aspects of the results not discussed in the main paper are included here.

##### Phylogenetic inference

In order to encompass the ever-growing complexities of phylogenomic methods and approaches we employed various approaches to infer 11 phylogenies, rather than relying on a single method, generating three nuclear ASTRAL trees (Figures S14 – S16), six nuclear RAxML phylogenies (Figures S17 – S22), one nuclear PhyloBayes tree (Figure S23), and one plastid RAxML tree (Figure S29). The workflows for generating these phylogenies differ in five important aspects: the genomic origin of the sequence data (i.e., nuclear or chloroplast genome), the model for handling the large number of genes in the nuclear dataset (i.e., using a gene tree-based multi-species coalescent model or concatenation), how paralogy was dealt with (i.e., with or without a full orthology assessment), the type of tree building algorithm (i.e., maximum likelihood or Bayesian), and the type of data used in the analyses (i.e., nucleotide or amino acid). While the resulting phylogenies are overall highly similar in terms of topology (Figure S28), support values, and taxonomic implications, there are some minor, yet important, differences between them (Figures S24 – S27). These differences reflect and result from the diverse ways the phylogenies were produced. Although the 'true' Caesalpinioideae species tree clearly remains unknown, and we therefore have no way of knowing which approaches yielded the most accurate phylogeny, comparing the 11 trees is informative about the impacts of the various workflows on the resulting phylogenies, as they are all largely based on the same underlying data.

Comparison of the 11 phylogenies, in which the (dis)similarities between the topologies of the trees are expressed as Robinson-Foulds (RF) distances ((*41*); Figure S28; see Table S12 for the actual values), shows that the factor that has the single greatest effect on the topology is the type of data the tree is based on, i.e. nuclear or plastid. All nuclear phylogenies, regardless of how they were inferred, are more similar to each other than they are to the chloroplast tree. The chloroplast and nuclear genomes have distinct evolutionary histories, which can conflict with each other (e.g., (*127–129*)), and there are several cases of such cytonuclear discordance in Caesalpinioideae, most notably regarding the monophyly, or not, of the genus *Senegalia* (Terra et al. in press). The fact that the chloroplast alignment is much smaller (circa 65,000 nucleotide sites, compared to circa 1,000,000 in the nuclear datasets), less informative (circa 36,600 alignment patterns, versus over 800,000 in the nuclear datasets), and considerably more gappy (almost 50% gaps, versus circa 12% in the nuclear alignments) than the various nuclear alignments (Table S11), likely also contributes to the differences between the plastid and nuclear phylogenies and the on average lower bootstrap support and shorter branch lengths across the plastid phylogeny.

When only the nuclear trees are considered, the largest influence on the resulting topologies is whether the multi-species coalescent model or a concatenation approach is used to infer a species tree, which remains one of the most contentious issues in phylogenomics (*120*, *130*, *131*). Proponents of the concatenation (or supermatrix) approach argue that concatenating several independent loci can result in finding 'hidden support' for relationships not inferred by individual gene trees (*130*). They also suggest that one of the main reasons for using the multi-species coalescent model, i.e. accounting for variation among gene trees due to incomplete lineage sorting (ILS), is often not applicable, as ILS is not a major cause of conflict among gene trees (*132*), and gene tree conflict is more often caused by analytical rather than biological issues (*133*). Advocates of the multi-species coalescent model contend that a major assumption of the concatenation model, that loci have topologically congruent genealogies, is often violated (as found here), and that concatenation does not account for many important biological phenomena, such as deep coalescence (*120*). Our results indicate that several shallow and deeper relationships differ between the concatenation and coalescence phylogenies (Figures S24 – S26; also see below) highlighting the impacts of the choice between these two approaches (*120*, *130*). However, it should be noted that the coalescence and concatenation phylogenies still yield largely congruent species tree estimates, and that any conflicts deeper in the phylogeny largely involve unsupported positions of clades on nodes that might better be regarded as polytomies (*4*) (see the PhyloBayes tree (Figure S23), ASTRAL's polytomy test (Figure S33), and below).

Interestingly, while the gene jackknifing approach applied to infer the PhyloBayes tree is based on a concatenated dataset, judging by the RF distances the topology of the resulting phylogeny appears almost intermediate between the (concatenated) RAxML trees and the (multi-species coalescent) ASTRAL phylogenies, although the PhyloBayes topology is slightly more similar to the RAxML phylogenies (mean RF distance 64) than to the ASTRAL trees (mean RF distance 83). The fact that the PhyloBayes phylogeny is a consensus tree in which poorly-supported relationships are collapsed into polytomies may help to explain why its topology is almost equally similar to the ASTRAL and RAxML trees, all of which are fully bifurcating. It is also possible that the PhyloBayes' advanced amino acid substitution model, CAT, which takes into account among-site rate heterogeneity and is able to suppress long-branch attraction to some degree (*33*–*35*, *134*), moves the resulting phylogeny away from the RAxML trees towards the ASTRAL topologies in the tree space. However, the substitution model alone cannot explain the surprising result that the amino acid-based RAxML trees are significantly more similar to the nucleotide-based RAxML phylogenies than to the amino acid-based PhyloBayes tree, especially since the amino acid substitution model employed in the RAxML analyses, LG4X, is, like PhyloBayes' CAT model, also adept at accounting for among-site rate heterogeneity (*32*). The similarity of the RAxML trees, and the relative dissimilarity between the RAxML and PhyloBayes phylogenies, might therefore better be explained by the differences resulting from the maximum likelihood-based tree search algorithm of RAxML and the Bayesian method of PhyloBayes. Our results suggest that the choice of tree building approach largely overrides most effects of expressing the dataset as nucleotides or amino acids, notwithstanding that the optimal type of character data has been a contentious issue in phylogenetics (e.g., (Simmons 2017)). Nevertheless, it should be noted that the difference between the amino acid-based RAxML and PhyloBayes phylogenies is small, such that, other than several shallow and difficult to resolve relationships, the phylogenies have largely congruent topologies (Figure S27).

Finally, and perhaps surprisingly, the factor that appears to have the least impact on the topology of the phylogenies is the orthology assessment. Trees based on just the single-copy genes, or on the single-copy genes plus the multi-copy genes which are treated as single-copy, or on the single-copy genes plus the multi-copy genes which are subjected to full orthology assessment (see Methods and Figures S2 – S4), are all highly congruent, regardless of whether these phylogenies were inferred by RAxML or ASTRAL and were based on nucleotide or amino acid data. This result is somewhat unexpected, given that rigorous orthology inference is widely suggested to be critical for phylogenomic analysis (*21*, *136*, *137*). One explanation could be that phylogenetic signal in the much larger number (821) of single-copy genes swamps the signal present in the relatively few (165) multi-copy genes. While this does not make orthology assessment irrelevant, it suggests that most of the topology of the resulting phylogeny is shaped by the single-copy genes. The marginally lower RF distances between trees based on just the single-copy genes and trees based on all genes with orthology inference than to trees without orthology assessment seem to support this possibility.

However, this hypothesis appears to be rejected by the finding that the single-copy ASTRAL tree is more similar to an ASTRAL tree based on just the 165 multi-copy genes without orthology assessment (RF distance 74; results not shown), than to a tree based on the same genes subjected to orthology inference (i.e., the 665 ortholog trees; RF distance 86; results not shown), implying that orthology assessment of the multi-copy genes actually shifts the resulting phylogeny further away from the single-copy genes phylogeny in tree space. A likely alternative explanation is that orthology assessment is of only marginal relevance for phylogenetic inference of Caesalpinioideae because this subfamily is unlikely to be directly subtended by a whole genome duplication (WGD) event (Koenen et al. 2020). While polyploidisation events have occurred several times within Caesalpinioideae, for example in *Leucaena* (*28*), *Vachellia*, and *Mimosa* (*58*, *138*), these WGDs appear to be placed relatively close to the tips of the tree, affecting just particular genera, subclades within genera, or individual species, and orthology assessment of the resulting paralogs will therefore only have a minimal effect on the overall topology of the phylogeny. In this context it is likely that detailed orthology assessment would have a much more significant and beneficial effect on the phylogenetic inference for other legume subfamilies, such as Papilionoideae which is thought to be subtended by two successive WGDs, or Detarioideae which is also thought to be subtended by a WGD (Koenen et al. 2020). A second and related explanation for the lack of clear benefits of orthology assessment in the Caesalpinioideae data may lie in the way multi-copy genes were identified and processed (see Methods and Figure S2). All genes with potential paralogs, as recovered by HybPiper (*17*), in over five percent of taxa were deemed multi-copy and subjected to orthology inference. During this process, gene trees were built from which orthologous clades containing at least 22 taxa were extracted, whereas clades with fewer taxa than this threshold were discarded. This process affects genes with different fractions of paralogs in contrasting ways: a hypothetical gene that has two orthologous copies in all taxa in the phylogeny will simply be divided into two orthologs, both of which are informative for all relationships in the phylogeny. In contrast, a gene that only has paralogous copies in a few taxa scattered across the phylogeny will be divided into many small clades, a large fraction of which are likely to contain fewer than 22 taxa and would therefore be discarded. As a result, orthology inference of low-copy genes, although theoretically appropriate, can result in less taxon-rich, more gappy, and less informative alignments than if the paralogs of these same genes were simply never considered (Figure S12). Given that only 31 of the 167 genes treated here as multi-copy have potential paralogs in more than a quarter of all taxa, the overall effect of the orthology assessment of these genes may have been to remove more phylogenetically informative sites than were gained. This suggests that the beneficial impacts of orthology assessment on phylogeny reconstruction are likely to be maximised if only those few genes that are duplicated across large parts of the phylogeny are considered.

**Taxonomic consequences**

The phylogenies inferred here reveal extensive generic non-monophyly across Caesalpinioideae, especially in the Mimosoid clade, which is discussed elsewhere (Ringelberg et al. submitted) and the taxonomic consequences of which are being dealt with in a forthcoming Special Issue of the journal *PhytoKeys*.

### 4. References

1. F. T. Bakker, A. Antonelli, J. A. Clarke, J. A. Cook, S. V. Edwards, P. G. P. Ericson, S. Faurby, N. Ferrand, M. Gelang, R. G. Gillespie, M. Irestedt, K. Lundin, E. Larsson, P. Matos-Maraví, J. Müller, T. von Proschwitz, G. K. Roderick, A. Schliep, N. Wahlberg, J. Wiedenhoeft, M. Källersjö, The global museum: Natural history collections and the future of evolutionary science and public education. *PeerJ*. **8**, 1–40 (2020).

2. E. K. Meineke, T. J. Davies, B. H. Daru, C. C. Davis, Biological collections for understanding biodiversity in the Anthropocene. *Philos. Trans. R. Soc. B Biol. Sci.* **374** (2019), doi:10.1098/rstb.2017.0386.

3. R Core Team, R: A language and environment for statistical computing. R Foundation for Statistical Computing, Vienna, Austria. URL https://www.R-project.org/ (2022).

4. E. J. M. Koenen, C. A. Kidner, É. R. de Souza, M. F. Simon, J. R. V. Iganci, J. A. Nicholls, G. K. Brown, L. P. de Queiroz, M. A. Luckow, G. P. Lewis, R. T. Pennington, C. E. Hughes, Hybrid capture of 964 nuclear genes resolves evolutionary relationships in the mimosoid legumes and reveals the polytomous origins of a large pantropical radiation. *Am. J. Bot.* **107**, 1710–1735 (2020).

5. V. A. Funk, R. J. Bayer, S. Keeley, R. Chan, L. Watson, B. Gemeinholzer, E. E. Schilling, J. L. Panero, B. G. Baldwin, N. Garcia-Jacas, A. Susanna, R. K. Jansen, Everywhere but Antarctica: Using a supertree to understand the diversity and distribution of the Compositae. *Biol. Skr.* **55**, 343–373 (2005).

6. V. A. Funk, C. D. Specht, Meta-trees: grafting for a global perspective. *Proc. Biol. Soc. Washingt.* **120**, 232–240 (2007).

7. E. L. Spriggs, P. A. Christin, E. J. Edwards, C4 photosynthesis promoted species diversification during the miocene grassland expansion. *PLoS One*. **9**, e97722 (2014).

8. LPWG, A new subfamily classification of the Leguminosae based on a taxonomically comprehensive phylogeny – The Legume Phylogeny Working Group (LPWG). *Taxon*. **66**, 44–77 (2017).

9. J. A. Nicholls, R. T. Pennington, E. J. M. Koenen, C. E. Hughes, J. Hearn, L. Bunnefeld, K. G. Dexter, G. N. Stone, C. A. Kidner, Using targeted enrichment of nuclear genes to increase phylogenetic resolution in the neotropical rain forest genus Inga (Leguminosae: Mimosoideae). *Front. Plant Sci.* **6** (2015), doi:10.3389/fpls.2015.00710.

10. E. Gagnon, A. Bruneau, C. E. Hughes, L. de Queiroz, G. P. Lewis, A new generic system for the pantropical Caesalpinia group (Leguminosae). *PhytoKeys*. **71**, 1–160 (2016).

11. P. G. Ribeiro, M. Luckow, G. P. Lewis, M. F. Simon, D. Cardoso, É. R. de Souza, A. P. C. Silva, M. C. Jesus, F. A. R. dos Santos, V. Azevedo, L. P. de Queiroz, Lachesiodendron, a new monospecific genus segregated from Piptadenia (Leguminosae: Caesalpinioideae: Mimosoid clade): Evidence from morphology and molecules. *Taxon*. **67**, 37–54 (2018).

12. M. F. Simon, J. F. B. Pastore, A. F. Souza, L. M. Borges, V. R. Scalon, P. G. Ribeiro, J. Santos-Silva, V. C. Souza, L. P. Queiroz, Molecular Phylogeny of Stryphnodendron (Mimosoideae, Leguminosae) and Generic Delimitations in the Piptadenia Group. *Int. J. Plant Sci.* **177**, 44–59 (2016).

13. S. F. Altschul, W. Gish, W. Miller, E. W. Myers, D. J. Lipman, Basic Local Alignment Search Tool. *J. Mol. Biol.* **215**, 403–410 (1990).

14. W. J. Kent, BLAT---The BLAST-Like Alignment Tool. *Genome Res.* **12**, 656–664 (2002).

15. H. Xu, X. Luo, J. Qian, X. Pang, J. Song, G. Qian, J. Chen, S. Chen, FastUniq: A Fast De Novo Duplicates Removal Tool for Paired Short Reads. *PLoS One*. **7**, 1–6 (2012).

16. A. M. Bolger, M. Lohse, B. Usadel, Trimmomatic: A flexible trimmer for Illumina sequence data. *Bioinformatics*. **30**, 2114–2120 (2014).

17. M. G. Johnson, E. M. Gardner, Y. Liu, R. Medina, B. Goffinet, A. J. Shaw, N. J. C. Zerega, N. J. Wickett, HybPiper: Extracting Coding Sequence and Introns for Phylogenetics from High-Throughput Sequencing Reads Using Target Enrichment. *Appl. Plant Sci.* **4**, 1600016 (2016).

18. H. Li, R. Durbin, Fast and accurate short read alignment with Burrows-Wheeler transform. *Bioinformatics*. **25**, 1754–1760 (2009).

19. A. Bankevich, S. Nurk, D. Antipov, A. A. Gurevich, M. Dvorkin, A. S. Kulikov, V. M. Lesin, S. I. Nikolenko, S. Pham, A. D. Prjibelski, A. V. Pyshkin, A. V. Sirotkin, N. Vyahhi, G. Tesler, M. A. Alekseyev, P. A. Pevzner, SPAdes: A new genome assembly algorithm and its applications to single-cell sequencing. *J. Comput. Biol.* **19**, 455–477 (2012).

20. G. S. C. Slater, E. Birney, Automated generation of heuristics for biological sequence comparison. *BMC Bioinformatics*. **6**, 1–11 (2005).

21. Y. Yang, S. A. Smith, Orthology Inference in Nonmodel Organisms Using Transcriptomes and Low-Coverage Genomes: Improving Accuracy and Matrix Occupancy for Phylogenomics. *Mol. Biol. Evol.* **31**, 3081–3092 (2014).

22. V. Ranwez, S. Harispe, F. Delsuc, E. J. P. Douzery, MACSE: Multiple alignment of coding SEquences accounting for frameshifts and stop codons. *PLoS One*. **6**, e22594 (2011).

23. J. W. Brown, J. F. Walker, S. A. Smith, Phyx: Phylogenetic tools for unix. *Bioinformatics*. **33**, 1886–1888 (2017).

24. A. Stamatakis, RAxML version 8: A tool for phylogenetic analysis and post-analysis of large phylogenies. *Bioinformatics*. **30**, 1312–1313 (2014).

25. U. Mai, S. Mirarab, TreeShrink: Fast and accurate detection of outlier long branches in collections of phylogenetic trees. *BMC Genomics*. **19**, 23–40 (2018).

26. A. Criscuolo, S. Gribaldo, BMGE (Block Mapping and Gathering with Entropy): A new software for selection of phylogenetic informative regions from multiple sequence alignments. *BMC Evol. Biol.* **10**, 210 (2010).

27. J. J. Doyle, in *Polyploidy and genome evolution*, P. S. Soltis, D. E. Soltis, Eds. (Springer-Verlag, Berlin - Heidelberg, Germany, 2012), pp. 147–180.

28. R. Govindarajulu, C. E. Hughes, P. J. Alexander, C. Donovan Bailey, The complex evolutionary dynamics of ancient and recent polyploidy in Leucaena (Leguminosae; Mimosoideae). *Am. J. Bot.* **98**, 2064–2076 (2011).

29. S. Mirarab, Species Tree Estimation Using ASTRAL: Practical Considerations, 1–21 (2019).

30. C. Zhang, M. Rabiee, E. Sayyari, S. Mirarab, ASTRAL-III: Polynomial time species tree reconstruction from partially resolved gene trees. *BMC Bioinformatics*. **19**, 15–30 (2018).

31. A. Stamatakis, RAxML-VI-HPC: Maximum likelihood-based phylogenetic analyses with thousands of taxa and mixed models. *Bioinformatics*. **22**, 2688–2690 (2006).

32. S. Q. Le, C. C. Dang, O. Gascuel, Modeling protein evolution with several amino acid replacement matrices depending on site rates. *Mol. Biol. Evol.* **29**, 2921–2936 (2012).

33. N. Lartillot, N. Rodrigue, D. Stubbs, J. Richer, PhyloBayes MPI: Phylogenetic Reconstruction with Infinite Mixtures of Profiles in a Parallel Environment. *Syst. Biol.* **62**, 11–615 (2013).

34. N. Lartillot, H. Brinkmann, H. Philippe, Suppression of long-branch attraction artefacts in the animal phylogeny using a site-heterogeneous model. *BMC Evol. Biol.* **7** (2007), doi:10.1186/1471-2148-7-S1-S4.

35. N. Lartillot, H. Philippe, A Bayesian mixture model for across-site heterogeneities in the amino-acid replacement process. *Mol. Biol. Evol.* **21**, 1095–1109 (2004).

36. M. Plummer, N. Best, K. Cowles, K. Vines, CODA: Convergence Diagnosis and Output Analysis for MCMC. *R News*. **6**, 7–11 (2006).

37. D. V Dugas, D. Hernandez, E. J. M. Koenen, E. Schwarz, S. Straub, C. E. Hughes, R. K. Jansen, M. Nageswara-Rao, M. Staats, J. T. Trujillo, N. H. Hajrah, N. S. Alharbi, A. L. Al-Malki, J. S. M. Sabir, C. D. Bailey, Mimosoid legume plastome evolution: IR expansion, tandem repeat expansions, and accelerated rate of evolution in clpP. *Sci. Rep.* **5** (2015), doi:10.1038/srep16958.

38. S. Huang, W. Wu, Z. Chen, Q. Zhu, W. L. Ng, Q. Zhou, Characterization of the chloroplast genome of Erythrophleum fordii (Fabaceae). *Conserv. Genet. Resour.* **11**, 165–167 (2019).

39. S. A. Smith, M. J. Moore, J. W. Brown, Y. Yang, Analysis of phylogenomic datasets reveals conflict, concordance, and gene duplications with examples from animals and plants. *BMC Evol. Biol.* **15**, 150 (2015).

40. X. Zhou, S. Lutteropp, L. Czech, A. Stamatakis, M. Von Looz, A. Rokas, Quartet-Based Computations of Internode Certainty Provide Robust Measures of Phylogenetic Incongruence. *Syst. Biol.* **69**, 308–324 (2020).

41. D. F. Robinson, L. R. Foulds, Comparison of phylogenetic trees. *Math. Biosci.* **53**, 131–147 (1981).

42. K. P. Schliep, phangorn: phylogenetic analysis in R. *Bioinformatics*. **27**, 592–593 (2011).

43. L. J. Revell, phytools: An R package for phylogenetic comparative biology (and other things). *Methods Ecol. Evol.* **3**, 217–223 (2012).

44. A. J. Aberer, D. Krompass, A. Stamatakis, Pruning rogue taxa improves phylogenetic accuracy: An efficient algorithm and webservice. *Syst. Biol.* **62**, 162–166 (2013).

45. M. A. Suchard, P. Lemey, G. Baele, D. L. Ayres, A. J. Drummond, A. Rambaut, Bayesian phylogenetic and phylodynamic data integration using BEAST 1.10. *Virus Evol.* **4**, 1–5 (2018).

46. A. J. Drummond, A. Rambaut, BEAST: Bayesian evolutionary analysis by sampling trees. *BMC Evol. Biol.* **7**, 1–8 (2007).

47. S. A. Smith, J. W. Brown, J. F. Walker, So many genes, so little time: A practical approach to divergence-time estimation in the genomic era. *PLoS One*. **13** (2018), doi:10.1371/journal.pone.0197433.

48. E. J. M. Koenen, D. I. Ojeda, R. Steeves, J. Migliore, F. T. Bakker, J. J. Wieringa, C. Kidner, O. J. Hardy, R. T. Pennington, A. Bruneau, C. E. Hughes, Large-scale genomic sequence data resolve the deepest divergences in the legume phylogeny and support a near-simultaneous

evolutionary origin of all six subfamilies. *New Phytol.* **225**, 1355–1369 (2020).

49. M. Lavin, B. P. Schrire, G. P. Lewis, R. T. Pennington, A. Delgado-Salinas, M. Thulin, C. E. Hughes, A. B. Matos, M. F. Wojciechowski, Metacommunity process rather than continental tectonic history better explains geographically structured phylogenies in legumes. *Philos. Trans. R. Soc. Lond. B. Biol. Sci.* **359**, 1509–1522 (2004).

50. A. Bruneau, M. Mercure, G. P. Lewis, P. S. Herendeen, Phylogenetic patterns and diversification in the caesalpinioid legumes. *Botany*. **86**, 697–718 (2008).

51. M. F. Simon, R. Grether, L. P. de Queiroz, C. Skema, R. T. Pennington, C. E. Hughes, Recent assembly of the Cerrado, a neotropical plant diversity hotspot, by in situ evolution of adaptations to fire. *Proc. Natl. Acad. Sci. U.S.A.* **106**, 20359–20364 (2009).

52. J. P. Huelsenbeck, F. Ronquist, MRBAYES: Bayesian inference of phylogeny. *Bioinformatics*. **17**, 754–755 (2001).

53. A. Rambaut, A. J. Drummond, D. Xie, G. Baele, M. A. Suchard, Posterior summarisation in Bayesian phylogenetics using Tracer 1.7. *Syst. Biol.* **67**, 901–904 (2018).

54. B. D. Mishler, N. Knerr, C. E. González-Orozco, A. D. Thornhill, S. W. Laffan, J. T. Miller, Phylogenetic measures of biodiversity and neo- and paleo-endemism in Australian Acacia. *Nat. Commun.* **5**, 4473 (2014).

55. É. R. de Souza, G. P. Lewis, F. Forest, A. S. Schnadelbach, C. van den Berg, L. P. de Queiroz, Phylogeny of Calliandra (Leguminosae: Mimosoideae) based on nuclear and plastid molecular markers. *Taxon*. **62**, 1200–1219 (2013).

56. C. E. Hughes, C. D. Bailey, S. Krosnick, M. A. Luckow, Relationships among genera of the informal Dichrostachys and Leucaena groups (Mimosoideae) inferred from nuclear ribosomal ITS sequences. *Adv. Legum. Syst.* **10**, 221–238 (2003).

57. S. B. Cannon, M. R. McKain, A. Harkess, M. N. Nelson, S. Dash, M. K. Deyholos, Y. Peng, B. Joyce, C. N. Stewart, M. Rolf, T. Kutchan, X. Tan, C. Chen, Y. Zhang, E. Carpenter, G. K. S. Wong, J. J. Doyle, J. Leebens-Mack, Multiple polyploidy events in the early radiation of nodulating and nonnodulating legumes. *Mol. Biol. Evol.* **32**, 193–210 (2015).

58. M. F. Simon, R. Grether, L. P. de Queiroz, T. E. Särkinen, V. F. Dutra, C. E. Hughes, The evolutionary history of Mimosa (Leguminosae): Toward a phylogeny of the sensitive plants. *Am. J. Bot.* **98**, 1201–1221 (2011).

59. T. N. C. Vasconcelos, S. Alcantara, C. O. Andrino, F. Forest, M. Reginato, M. F. Simon, J. R. Pirani, Fast diversification through a mosaic of evolutionary histories characterizes the endemic flora of ancient Neotropical mountains. *Proc. R. Soc. B Biol. Sci.* **287**, 20192933 (2020).

60. J. S. Boatwright, O. Maurin, M. van der Bank, Phylogenetic position of Madagascan species of Acacia s.l. and new combinations in Senegalia and Vachellia (Fabaceae, Mimosoideae, Acacieae). *Bot. J. Linn. Soc.* **179**, 288–294 (2015).

61. D. F. Comben, G. A. McCulloch, G. K. Brown, G. H. Walter, Phylogenetic placement and the timing of diversification in Australia’s endemic Vachellia (Caesalpinioideae, Mimosoid Clade, Fabaceae) species. *Aust. Syst. Bot.* **33**, 103–109 (2020).

62. V. Terra, F. C. P. Garcia, L. P. de Queiroz, M. van der Bank, J. T. Miller, Phylogenetic Relationships in Senegalia (Leguminosae-Mimosoideae) Emphasizing the South American Lineages. *Syst. Bot.* **42**, 458–464 (2017).

63. J. Ferm, A preliminary phylogeny of Zapoteca (Fabaceae: Caesalpinioideae: Mimosoid clade). *Plant Syst. Evol.* **305**, 341–352 (2019).

64. É. R. de Souza, M. V. Krishnaraj, L. P. de Queiroz, Sanjappa, a new genus in the tribe Ingeae (Leguminosae: Mimosoideae) from India. *Rheedea*. **26**, 1–12 (2016).

65. J. Ferm, P. Korall, G. P. Lewis, B. Ståhl, Phylogeny of the Neotropical legume genera Zygia and Marmaroxylon and close relatives. *Taxon*. **68**, 661–672 (2019).

66. C. E. González-Orozco, S. W. Laffan, N. Knerr, J. T. Miller, A biogeographical regionalization of Australian Acacia species. *J. Biogeogr.* **40**, 2156–2166 (2013).

67. J. J. Ringelberg, N. E. Zimmermann, A. Weeks, M. Lavin, C. E. Hughes, Biomes as evolutionary arenas: Convergence and conservatism in the trans-continental succulent biome. *Glob. Ecol. Biogeogr.* **29**, 1100–1113 (2020).

68. E. Gagnon, J. J. Ringelberg, A. Bruneau, G. P. Lewis, C. E. Hughes, Global Succulent Biome phylogenetic conservatism across the pantropical Caesalpinia Group (Leguminosae). *New Phytol.* **222**, 1994–2008 (2019).

69. H. ter Steege, N. C. A. Pitman, D. Sabatier, C. Baraloto, R. P. Salomão, J. E. Guevara, O. L. Phillips, C. V. Castilho, W. E. Magnusson, J. F. Molino, A. Monteagudo, P. Núñez Vargas, J. C. Montero, T. R. Feldpausch, E. N. H. Coronado, T. J. Killeen, B. Mostacedo, R. Vasquez, R. L. Assis, J. Terborgh, F. Wittmann, A. Andrade, W. F. Laurance, S. G. W. Laurance, B. S. Marimon, B. H. Marimon, I. C. Guimarães Vieira, I. L. Amaral, R. Brienen, H. Castellanos, D. Cárdenas López, J. F. Duivenvoorden, H. F. Mogollón, F. D. A. Matos, N. Dávila, R. García-Villacorta, P. R. Stevenson Diaz, F. Costa, T. Emilio, C. Levis, J. Schietti, P. Souza, A. Alonso, F. Dallmeier, A. J. D. Montoya, M. T. Fernandez Piedade, A. Araujo-Murakami, L. Arroyo, R. Gribel, P. V. A. Fine, C. A. Peres, M. Toledo, G. A. Aymard C, T. R. Baker, C. Cerón, J. Engel, T. W. Henkel, P. Maas, P. Petronelli, J. Stropp, C. E. Zartman, D. Daly, D. Neill, M. Silveira, M. R. Paredes, J. Chave, D. A. Lima Filho, P. M. Jørgensen, A. Fuentes, J. Schöngart, F. Cornejo Valverde, A. Di Fiore, E. M. Jimenez, M. C. Peñuela Mora, J. F. Phillips, G. Rivas, T. R. van Andel, P. von Hildebrand, B. Hoffman, E. L. Zent, Y. Malhi, A. Prieto, A. Rudas, A. R. Ruschel, N. Silva, V. A. Vos, S. Zent, A. A. Oliveira, A. C. Schutz, T. Gonzales, M. Trindade Nascimento, H. Ramirez-Angulo, R. Sierra, M. Tirado, M. N. Umaña Medina, G. van der Heijden, C. I. A. Vela, E. Vilanova Torre, C. Vriesendorp, O. Wang, K. R. Young, C. Baider, H. Balslev, C. Ferreira, I. Mesones, A. Torres-Lezama, L. E. Urrego Giraldo, R. Zagt, M. N. Alexiades, L. Hernandez, I. Huamantupa-Chuquimaco, W. Milliken, W. Palacios Cuenca, D. Pauletto, E. Valderrama Sandoval, L. Valenzuela Gamarra, K. G. Dexter, K. Feeley, G. Lopez-Gonzalez, M. R. Silman, Hyperdominance in the Amazonian tree flora. *Science*. **342**, 1243092 (2013).

70. D. Cardoso, T. Särkinen, S. Alexander, A. M. Amorim, V. Bittrich, M. Celis, D. C. Daly, P. Fiaschi, V. A. Funk, L. L. Giacomin, R. Goldenberg, G. Heiden, J. Iganci, C. L. Kelloff, S. Knapp, H. Cavalcante de Lima, A. F. P. Machado, R. M. dos Santos, R. Mello-Silva, F. A. Michelangeli, J. Mitchell, P. Moonlight, P. L. R. de Moraes, S. A. Mori, T. S. Nunes, T. D. Pennington, J. R. Pirani, G. T. Prance, L. P. de Queiroz, A. Rapini, R. Riina, C. A. V. Rincon, N. Roque, G. Shimizu, M. Sobral, J. R. Stehmann, W. D. Stevens, C. M. Taylor, M. Trovó, C. van den Berg, H. van der Werff, P. L. Viana, C. E. Zartman, R. C. Forzza, Amazon plant diversity revealed by a taxonomically verified species list. *Proc. Natl. Acad. Sci. U.S.A.* **114**, 201706756 (2017).

71. DRYFLOR, K. Banda-R, A. Delgado-Salinas, K. G. Dexter, R. Linares-Palomino, A. Oliveira-Filho, D. Prado, M. Pullan, C. Quintana, R. Riina, G. M. Rodríguez, J. Weintritt, P. Acevedo-Rodríguez, J. Adarve, E. Álvarez, Ba. Aranguren, G. A. Julián Camilo Arteaga, A. Castaño, Á. C. Natalia Ceballos-Mago, H. Cuadros, D. Freddy, W. Devia, H. Dueñas, L. Fajaro, Á. Fernández,

M. Á. Fernández, J. Franklin, E. H. Freid, L. A. Galetti, R. Gonto, R. González-M., R. Graveson, E. H. Helmer, Á. Idárrago, R. Lopéz, H. Marcano-Vega, O. G. Martínez, H. M. Maturo, M. McDonald, K. McLaren, O. Melo, F. Mijares, V. Mogni, D. Molina, N. del P. Moreno, J. M. Nassar, D. M. Neves, L. J. Oakley, M. Oatham, A. R. Olvera-Luna, F. F. Pezzini, O. J. R. Dominguez, M. E. Ríos, O. Rivera, N. Rodríguez, A. Rojas, T. E. Särkinen, R. Sánchez, M. Smith, C. Vargas, B. Villanueva, R. T. Pennington, Plant diversity patterns in neotropical dry forests and their conservation implications. *Science*. **353**, 1383–1387 (2016).

72. A. Fayolle, M. D. Swaine, J. Aleman, A. F. Azihou, D. Bauman, M. te Beest, E. N. Chidumayo, J. P. G. M. Cromsigt, H. Dessard, M. Finckh, F. M. P. Gonçalves, J. F. Gillet, A. Gorel, A. Hick, R. Holdo, B. Kirunda, G. Mahy, I. McNicol, C. M. Ryan, R. Revermann, A. Plumptre, R. Pritchard, P. Nieto-Quintano, C. B. Schmitt, J. Seghieri, A. Swemmer, H. Talila, E. Woollen, A sharp floristic discontinuity revealed by the biogeographic regionalization of African savannas. *J. Biogeogr.* **46**, 454–465 (2019).

73. A. Baselga, Partitioning the turnover and nestedness components of beta diversity. *Glob. Ecol. Biogeogr.* **19**, 134–143 (2010).

74. F. Leprieur, C. Albouy, J. de Bortoli, P. F. Cowman, D. R. Bellwood, D. Mouillot, Quantifying phylogenetic beta diversity: Distinguishing between “true” turnover of lineages and phylogenetic diversity gradients. *PLoS One*. **7** (2012), doi:10.1371/journal.pone.0042760.

75. G. G. Simpson, Mammals and the nature of continents. *Am. J. Sci.* **241** (1943), pp. 1–31.

76. H. Kreft, W. Jetz, A framework for delineating biogeographical regions based on species distributions. *J. Biogeogr.* **37**, 2029–2053 (2010).

77. A. Castro-Insua, C. Gómez-Rodríguez, A. Baselga, Dissimilarity measures affected by richness differences yield biased delimitations of biogeographic realms. *Nat. Commun.* **9**, 9–11 (2018).

78. A. Baselga, D. Orme, S. Villeger, J. de Bortoli, F. Leprieur, betapart: Partitioning Beta Diversity into Turnover and Nestedness Components. R package version 1.5.1. (2018).

79. G. P. Lewis, B. Schrire, B. Mackinder, M. Lock, *Legumes of the world* (Royal Botanic Gardens, Kew, United Kingdom, 2005).

80. D. Nychka, R. Furrer, J. Paige, S. Sain, “fields: Tools for spatial data.” doi: 10.5065/D6W957CT (URL: https://doi.org/10.5065/D6W957CT), R package version 10.3, <URL: https://github.com/NCAR/Fields> (2017).

81. M. C. Fitzpatrick, K. Mokany, G. Manion, M. Lisk, S. Ferrier, D. Nieto-Lugilde, gdm: Generalized Dissimilarity Modeling. R package version 1.4. (2020).

82. D. N. Karger, O. Conrad, J. Böhner, T. Kawohl, H. Kreft, R. W. Soria-Auza, N. E. Zimmermann, H. P. Linder, M. Kessler, Climatologies at high resolution for the earth’s land surface areas. *Sci. Data*. **4**, 1–20 (2017).

83. A. M. Wilson, W. Jetz, Remotely Sensed High-Resolution Global Cloud Dynamics for Predicting Ecosystem and Biodiversity Distributions. *PLoS Biol.* **14**, e1002415 (2016).

84. R. J. Hijmans, raster: Geographic Data Analysis and Modeling. R package version 2.8-4. https://CRAN.R-project.org/package=raster (2018).

85. D. F. Rosauer, S. Ferrier, K. J. Williams, G. Manion, J. S. Keogh, S. W. Laffan, Phylogenetic generalised dissimilarity modelling: A new approach to analysing and predicting spatial turnover in the phylogenetic composition of communities. *Ecography (Cop.).* **37**, 21–32 (2014).

86. S. Ferrier, G. Manion, J. Elith, K. Richardson, Using generalized dissimilarity modelling to analyse and predict patterns of beta diversity in regional biodiversity assessment. *Divers. Distrib.* **13**, 252–264 (2007).

87. M. C. Fitzpatrick, N. J. Sanders, S. Normand, J. C. Svenning, S. Ferrier, A. D. Gove, R. R. Dunn, Environmental and historical imprints on beta diversity: Insights from variation in rates of species turnover along gradients. *Proc. R. Soc. B Biol. Sci.* **280**, 20131201 (2013).

88. C. H. Graham, P. V. A. Fine, Phylogenetic beta diversity: Linking ecological and evolutionary processes across space in time. *Ecol. Lett.* **11**, 1265–1277 (2008).

89. C. König, P. Weigelt, H. Kreft, Dissecting global turnover in vascular plants. *Glob. Ecol. Biogeogr.* **26**, 228–242 (2017).

90. W. L. Eiserhardt, J.-C. Svenning, W. J. Baker, T. L. P. Couvreur, H. Balslev, Dispersal and niche evolution jointly shape the geographic turnover of phylogenetic clades across continents. *Sci. Rep.* **3**, 1164 (2013).

91. B. G. Holt, G. C. Costa, C. Penone, J. P. Lessard, T. M. Brooks, A. D. Davidson, S. Blair Hedges, V. C. Radeloff, C. Rahbek, C. Rondinini, C. H. Graham, Environmental variation is a major predictor of global trait turnover in mammals. *J. Biogeogr.* **45**, 225–237 (2018).

92. H. Qian, N. G. Swenson, J. Zhang, Phylogenetic beta diversity of angiosperms in North America. *Glob. Ecol. Biogeogr.* **22**, 1152–1161 (2013).

93. C. Penone, G. C. Costa, B. G. Weinstein, C. H. Graham, A. D. Davidson, T. M. Brooks, C. Rondinini, S. B. Hedges, Global mammal beta diversity shows parallel assemblage structure in similar but isolated environments. *Proc. R. Soc. B Biol. Sci.* **283**, 1–9 (2016).

94. B. Saladin, W. Thuiller, C. H. Graham, S. Lavergne, L. Maiorano, N. Salamin, N. E. Zimmermann, Environment and evolutionary history shape phylogenetic turnover in European tetrapods. *Nat. Commun.* **10**, 1–9 (2019).

95. D. L. Warren, M. Cardillo, D. F. Rosauer, D. I. Bolnick, Mistaking geography for biology: Inferring processes from species distributions. *Trends Ecol. Evol.* **29**, 572–580 (2014).

96. H. Qian, Y. Jin, F. Leprieur, X. Wang, T. Deng, Geographic patterns and environmental correlates of taxonomic and phylogenetic beta diversity for large-scale angiosperm assemblages in China. *Ecography (Cop.).*, 1–11 (2020).

97. J. N. Pinto-Ledezma, D. J. Larkin, J. Cavender-Bares, Patterns of beta diversity of vascular plants and their correspondence with biome boundaries across North America. *Front. Ecol. Evol.* **6**, 194 (2018).

98. C. H. Graham, J. L. Parra, C. Rahbek, J. A. McGuire, Phylogenetic structure in tropical hummingbird communities. *Proc. Natl. Acad. Sci. U.S.A.* **106**, 19673–19678 (2009).

99. B. H. Daru, M. van der Bank, T. J. Davies, Unravelling the evolutionary origins of biogeographic assemblages. *Divers. Distrib.* **24**, 313–324 (2018).

100. I. R. McFadden, M. T. P. Coelho, R. O. Wüest, F. A. S. Cassemiro, N. E. Zimmermann, L. Pellissier, T. F. Rangel, C. H. Graham, Global hotspots of recent and ancestral turnover in birds. *Res. Sq.* (2020), doi:https://doi.org/10.21203/rs.3.rs-131370/v2.

101. B. H. Daru, T. L. Elliott, D. S. Park, T. J. Davies, Understanding the Processes Underpinning Patterns of Phylogenetic Regionalization. *Trends Ecol. Evol.* **32**, 845–860 (2017).

102. D. H. Ogle, J. C. Doll, P. Wheeler, A. Dinno, FSA: Fisheries Stock Analysis. R package version 0.9.1, https://github.com/droglenc/FSA (2021).

103. A. Dinno, dunn.test: Dunn's Test of Multiple Comparisons Using Rank Sums. R package version 1.3.5. https://CRAN.R-project.org/package=dunn.test (2017).

104. D. M. Neves, K. G. Dexter, T. R. Baker, F. Coelho de Souza, A. T. Oliveira-Filho, L. P. Queiroz, H. C. Lima, M. F. Simon, G. P. Lewis, R. A. Segovia, L. Arroyo, C. Reynel, J. L. Marcelo-Peña, I. Huamantupa-Chuquimaco, D. Villarroel, G. A. Parada, A. Daza, R. Linares-Palomino, L. V. Ferreira, R. P. Salomão, G. S. Siqueira, M. T. Nascimento, C. N. Fraga, R. T. Pennington, Evolutionary diversity in tropical tree communities peaks at intermediate precipitation. *Sci. Rep.* **10**, 1–7 (2020).

105. M. Pagel, Inferring the historical patterns of biological evolution. *Nature*. **401**, 877–884 (1999).

106. N. J. Matzke, Probabilistic historical biogeography: new models for founder-event speciation, imperfect detection, and fossils allow improved accuracy and model-testing. *Front. Biogeogr.* **5**, 242–248 (2013).

107. N. J. Matzke, _BioGeoBEARS: BioGeography with Bayesian (and Likelihood) Evolutionary Analysis in R Scripts_. University of California, Berkeley, Berkeley, CA (2013).

108. N. J. Matzke, Model selection in historical biogeography reveals that founder-event speciation is a crucial process in island clades. *Syst. Biol.* **63**, 951–970 (2014).

109. M. Arakaki, P.-A. Christin, R. Nyffeler, A. Lendel, U. Eggli, R. M. Ogburn, E. Spriggs, M. J. Moore, E. J. Edwards, Contemporaneous and recent radiations of the world's major succulent plant lineages. *Proc. Natl. Acad. Sci. U.S.A.* **108**, 8379–8384 (2011).

110. D. L. Rabosky, Automatic detection of key innovations, rate shifts, and diversity-dependence on phylogenetic trees. *PLoS One*. **9** (2014), doi:10.1371/journal.pone.0089543.

111. B. R. Moore, S. Höhna, M. R. May, B. Rannala, J. P. Huelsenbeck, Critically evaluating the theory and performance of Bayesian analysis of macroevolutionary mixtures. *Proc. Natl. Acad. Sci. U.S.A.* **113**, 9569–9574 (2016).

112. A. L. S. Meyer, J. J. Wiens, Estimating diversification rates for higher taxa: BAMM can give problematic estimates of rates and rate shifts. *Evolution*. **72**, 39–53 (2018).

113. D. L. Rabosky, J. S. Mitchell, J. Chang, Is BAMM Flawed? Theoretical and Practical Concerns in the Analysis of Multi-Rate Diversification Models. *Syst. Biol.* **66**, 477–498 (2017).

114. D. L. Rabosky, BAMM at the court of false equivalency: A response to Meyer and Wiens. *Evolution*. **72**, 2246–2256 (2018).

115. D. L. Rabosky, Extinction rates should not be estimated from molecular phylogenies. *Evolution*. **64**, 1816–1824 (2010).

116. S. Louca, M. W. Pennell, Extant timetrees are consistent with a myriad of diversification histories. *Nature*. **580**, 502–505 (2020).

117. D. L. Rabosky, M. Grundler, C. Anderson, P. Title, J. J. Shi, J. W. Brown, H. Huang, J. G. Larson, BAMMtools: An R package for the analysis of evolutionary dynamics on phylogenetic trees. *Methods Ecol. Evol.* **5**, 701–707 (2014).

118. G. E. Brewer, J. J. Clarkson, O. Maurin, A. R. Zuntini, V. Barber, S. Bellot, N. Biggs, R. S. Cowan,

N. M. J. Davies, S. Dodsworth, S. L. Edwards, W. L. Eiserhardt, N. Epitawalage, S. Frisby, A. Grall, P. J. Kersey, L. Pokorny, I. J. Leitch, F. Forest, W. J. Baker, Factors Affecting Targeted Sequencing of 353 Nuclear Genes From Herbarium Specimens Spanning the Diversity of Angiosperms. *Front. Plant Sci.* **10**, 1–14 (2019).

119. M. L. Hart, L. L. Forrest, J. A. Nicholls, C. A. Kidner, Retrieval of hundreds of nuclear loci from herbarium specimens. *Taxon*. **65**, 1081–1092 (2016).

120. X. Jiang, S. V Edwards, L. Liu, The Multispecies Coalescent Model Outperforms Concatenation Across Diverse Phylogenomic Data Sets. *Syst. Biol.* **69**, 795–812 (2020).

121. C. Meyer, P. Weigelt, H. Kreft, Multidimensional biases, gaps and uncertainties in global plant occurrence information. *Ecol. Lett.* **19**, 992–1006 (2016).

122. M. A. Hyde, B. T. Wursten, P. Ballings, M. Coates Palgrave, Flora of Mozambique: Species information: individual images: Entada rheedii. https://www.mozambiqueflora.com/speciesdata/image-display.php?species_id=126520&image_id=13, retrieved 31 May 2022.

123. A. South, rworldxtra: Country boundaries at high resolution. R package version 1.01. https://CRAN.R-project.org/package=rworldxtra (2012).

124. R. H. Ree, I. Sanmartín, Conceptual and statistical problems with the DEC+J model of founder-event speciation and its comparison with DEC via model selection. *J. Biogeogr.* **45**, 741–749 (2018).

125. J. C. Zachos, G. R. Dickens, R. E. Zeebe, An early Cenozoic perspective on greenhouse warming and carbon-cycle dynamics. *Nature*. **451**, 279–283 (2008).

126. H. Morlon, E. Lewitus, F. L. Condamine, M. Manceau, J. Clavel, J. Drury, RPANDA: An R package for macroevolutionary analyses on phylogenetic trees. *Methods Ecol. Evol.* **7**, 589–597 (2016).

127. J. P. Rose, C. A. P. Toledo, E. M. Lemmon, A. R. Lemmon, K. J. Sytsma, Out of Sight, Out of Mind: Widespread Nuclear and Plastid-Nuclear Discordance in the Flowering Plant Genus Polemonium (Polemoniaceae) Suggests Widespread Historical Gene Flow Despite Limited Nuclear Signal. *Syst. Biol.* **70**, 162–180 (2021).

128. J. A. Lee-Yaw, C. J. Grassa, S. Joly, R. L. Andrew, L. H. Rieseberg, An evaluation of alternative explanations for widespread cytonuclear discordance in annual sunflowers (Helianthus). *New Phytol.* **221**, 515–526 (2019).

129. S. Bruun-Lund, W. L. Clement, F. Kjellberg, N. Rønsted, First plastid phylogenomic study reveals potential cyto-nuclear discordance in the evolutionary history of Ficus L. (Moraceae). *Mol. Phylogenet. Evol.* **109**, 93–104 (2017).

130. J. Gatesy, M. S. Springer, Phylogenetic analysis at deep timescales: Unreliable gene trees, bypassed hidden support, and the coalescence/concatalescence conundrum. *Mol. Phylogenet. Evol.* **80**, 231–266 (2014).

131. M. S. Springer, J. Gatesy, The gene tree delusion. *Mol. Phylogenet. Evol.* **94**, 1–33 (2016).

132. C. Scornavacca, N. Galtier, Incomplete lineage sorting in mammalian phylogenomics. *Syst. Biol.* **66**, 112–120 (2017).

133. E. J. Richards, J. M. Brown, A. J. Barley, R. A. Chong, R. C. Thomson, Variation across mitochondrial gene trees provides evidence for systematic error: How much gene tree

variation is biological? *Syst. Biol.* **67**, 847–860 (2018).

134. N. Lartillot, H. Philippe, Computing Bayes factors using thermodynamic integration. *Syst. Biol.* **55**, 195–207 (2006).

135. M. P. Simmons, Relative benefits of amino-acid, codon, degeneracy, DNA, and purine-pyrimidine character coding for phylogenetic analyses of exons. *J. Syst. Evol.* **55**, 85–109 (2017).

136. A. J. Moore, J. M. De Vos, L. P. Hancock, E. Goolsby, E. J. Edwards, Targeted Enrichment of Large Gene Families for Phylogenetic Inference: Phylogeny and Molecular Evolution of Photosynthesis Genes in the Portullugo Clade (Caryophyllales). *Syst. Biol.* **67**, 367–383 (2018).

137. N. Karimi, C. E. Grover, J. P. Gallagher, J. F. Wendel, C. Ané, D. A. Baum, Reticulate Evolution Helps Explain Apparent Homoplasy in Floral Biology and Pollination in Baobabs (Adansonia; Bombacoideae; Malvaceae). *Syst. Biol.* **69**, 462–478 (2019).

138. N. Dahmer, M. F. Simon, M. T. Schifino-Wittmann, C. E. Hughes, S. T. S. Miotto, J. C. Giuliani, Chromosome numbers in the genus Mimosa L.: Cytotaxonomic and evolutionary implications. *Plant Syst. Evol.* **291**, 211–220 (2011).

139. D. J. Bertioli, S. B. Cannon, L. Froenicke, G. Huang, A. D. Farmer, E. K. S. Cannon, X. Liu, D. Gao, J. Clevenger, S. Dash, L. Ren, M. C. Moretzsohn, K. Shirasawa, W. Huang, B. Vidigal, B. Abernathy, Y. Chu, C. E. Niederhuth, P. Umale, A. C. G. Arajo, A. Kozik, K. Do Kim, M. D. Burow, R. K. Varshney, X. Wang, X. Zhang, N. Barkley, P. M. Guimares, S. Isobe, B. Guo, B. Liao, H. T. Stalker, R. J. Schmitz, B. E. Scheffler, S. C. M. Leal-Bertioli, X. Xun, S. A. Jackson, R. Michelmore, P. Ozias-Akins, The genome sequences of Arachis duranensis and Arachis ipaensis, the diploid ancestors of cultivated peanut. *Nat. Genet.* **48**, 438–446 (2016).

140. J. S. Stai, A. Yadav, C. Sinou, A. Bruneau, J. J. Doyle, D. Fernández-Baca, S. B. Cannon, Cercis: A non-polyploid genomic relic within the generally polyploid legume family. *Front. Plant Sci.* **10**, 1–18 (2019).

141. J. Schmutz, S. B. Cannon, J. Schlueter, J. Ma, T. Mitros, W. Nelson, D. L. Hyten, Q. Song, J. J. Thelen, J. Cheng, D. Xu, U. Hellsten, G. D. May, Y. Yu, T. Sakurai, T. Umezawa, M. K. Bhattacharyya, D. Sandhu, B. Valliyodan, E. Lindquist, M. Peto, D. Grant, S. Shu, D. Goodstein, K. Barry, M. Futrell-Griggs, B. Abernathy, J. Du, Z. Tian, L. Zhu, N. Gill, T. Joshi, M. Libault, A. Sethuraman, X. C. Zhang, K. Shinozaki, H. T. Nguyen, R. A. Wing, P. Cregan, J. Specht, J. Grimwood, D. Rokhsar, G. Stacey, R. C. Shoemaker, S. A. Jackson, Genome sequence of the palaeopolyploid soybean. *Nature*. **463**, 178–183 (2010).

142. N. D. Young, F. Debellé, G. E. D. Oldroyd, R. Geurts, S. B. Cannon, M. K. Udvardi, V. A. Benedito, K. F. X. Mayer, J. Gouzy, H. Schoof, Y. Van De Peer, S. Proost, D. R. Cook, B. C. Meyers, M. Spannagl, F. Cheung, S. De Mita, V. Krishnakumar, H. Gundlach, S. Zhou, J. Mudge, A. K. Bharti, J. D. Murray, M. A. Naoumkina, B. Rosen, K. A. T. Silverstein, H. Tang, S. Rombauts, P. X. Zhao, P. Zhou, V. Barbe, P. Bardou, M. Bechner, A. Bellec, A. Berger, H. Bergès, S. Bidwell, T. Bisseling, N. Choisne, A. Couloux, R. Denny, S. Deshpande, X. Dai, J. J. Doyle, A. M. Dudez, A. D. Farmer, S. Fouteau, C. Franken, C. Gibelin, J. Gish, S. Goldstein, A. J. González, P. J. Green, A. Hallab, M. Hartog, A. Hua, S. J. Humphray, D. H. Jeong, Y. Jing, A. Jöcker, S. M. Kenton, D. J. Kim, K. Klee, H. Lai, C. Lang, S. Lin, S. L. MacMil, G. Magdelenat, L. Matthews, J. McCorrison, E. L. Monaghan, J. H. Mun, F. Z. Najar, C. Nicholson, C. Noirot, M. O'Bleness, C. R. Paule, J. Poulain, F. Prion, B. Qin, C. Qu, E. F. Retzel, C. Riddle, E. Sallet, S. Samain, N. Samson, I. Sanders, O. Saurat, C. Scarpelli, T. Schiex, B. Segurens, A. J. Severin, D. J. Sherrier, R. Shi, S. Sims, S. R. Singer, S. Sinharoy, L. Sterck, A. Viollet, B. B. Wang, K. Wang, M.

Wang, X. Wang, J. Warfsmann, J. Weissenbach, D. D. White, J. D. White, G. B. Wiley, P. Wincker, Y. Xing, L. Yang, Z. Yao, F. Ying, J. Zhai, L. Zhou, A. Zuber, J. Dénarié, R. A. Dixon, G. D. May, D. C. Schwartz, J. Rogers, F. Quétier, C. D. Town, B. A. Roe, The Medicago genome provides insight into the evolution of rhizobial symbioses. *Nature*. **480**, 520–524 (2011).

143. M. Griesmann, Y. Chang, X. Liu, Y. Song, G. Haberer, M. B. Crook, B. Billault-Penneteau, D. Lauressergues, J. Keller, L. Imanishi, Y. P. Roswanjaya, W. Kohlen, P. Pujic, K. Battenberg, N. Alloisio, Y. Liang, H. Hilhorst, M. G. Salgado, V. Hocher, H. Gherbi, S. Svistoonoff, J. J. Doyle, S. He, Y. Xu, S. Xu, J. Qu, Q. Gao, X. Fang, Y. Fu, P. Normand, A. M. Berry, L. G. Wall, J. M. Ané, K. Pawlowski, X. Xu, H. Yang, M. Spannagl, K. F. X. Mayer, G. K. S. Wong, M. Parniske, P. M. Delaux, S. Cheng, Phylogenomics reveals multiple losses of nitrogen-fixing root nodule symbiosis. *Science*. **361**, eaat1743 (2018).

144. S. L. Wing, F. Herrera, C. A. Jaramillo, C. Gómez-Navarro, P. Wilf, C. C. Labandeira, Late Paleocene fossils from the Cerrejón Formation, Colombia, are the earliest record of Neotropical rainforest. *Proc. Natl. Acad. Sci. U.S.A.* **106**, 18627–18632 (2009).

145. F. Herrera, M. R. Carvalho, S. L. Wing, C. Jaramillo, P. S. Herendeen, Middle to Late Paleocene Leguminosae fruits and leaves from Colombia. *Aust. Syst. Bot.* **32**, 385–408 (2019).

146. H. D. MacGinitie, in *Carnegie Institution of Washington publication 599. Contributions of Paleontology* (Washington D.C., USA, 1953).

147. P. S. Herendeen, F. Herrera, Eocene fossil legume leaves referable to the extant genus Arcoa (Caesalpinioideae, leguminosae). *Int. J. Plant Sci.* **180**, 220–231 (2019).

148. P. S. Herendeen, in *Advances in Legume Systematics part 4: The Fossil Record*, P. S. Herendeen, D. L. Dilcher, Eds. (Royal Botanic Gardens, Kew, United Kingdom, 1992), pp. 85–160.

149. W. L. Crepet, D. L. Dilcher, Investigations of Angiosperms from the Eocene of North America: A Mimosoid Inflorescence. *Am. J. Bot.* **64**, 714–725 (1977).

150. P. Guinet, E. Sabrouty, H. A. Soliman, A. M. Omrah, Études des caractères du pollen des Légumineuses - Mimosoideae des sédiments Tertiaires du Nord Ouest de l'Égypte. *Mém. Travel. Ec. Prat. des Hautes Etudes, Inst. Montpellier*. **17**, 159–171 (1987).

151. J. T. Miller, D. J. Murphy, S. Y. W. Ho, D. J. Cantrill, D. Seigler, Comparative dating of Acacia: Combining fossils and multiple phylogenies to infer ages of clades with poor fossil records. *Aust. J. Bot.* **61**, 436–445 (2013).

152. M. Caccavari, V. Barreda, A new calymmate mimosoid polyad from the Miocene of Argentina. *Rev. Palaeobot. Palynol.* **109**, 197–203 (2000).

### 5. Supplementary tables

**Table S1.** Names and voucher details for species and accessions sampled, European Nucleotide Archive (ENA) accession codes, and the sequencing and target assembly results, including numbers of sequence reads on target, genes recovered, and paralog warnings for each accession.

| Taxon | Voucher | ENA accession number | Source | Fraction of duplicated reads | Fraction of filtered reads | Reads left | Reads mapped | Genes with sequences | Genes recovered with 75% of target length | Potential paralogs |
|---|---|---|---|---|---|---|---|---|---|---|
| *Abarema cochliacarpos* | de Queiroz 15538 (HUEFS) | ERS4812838 | Koenen et al. 2020 | 0.552 | 0.07 | 8499987 | 3150000 (37.1%) | 971 | 853 | 15 |
| *Abarema diamantina* | Guerra 148 (ICN) | ERS11697078 | This study | 0.194 | 0.005 | 6944231 | 5312876 (76.5%) | 992 | 966 | 54 |
| *Acacia alata* var. *biglandulosa* | Murphy 464 (MELU) | ERS11697096 | This study | 0.142 | 0.005 | 3811009 | 3043513 (79.9%) | 992 | 961 | 35 |
| *Acacia ampliceps* | Murphy 323 (MELU) | ERS11697097 | This study | 0.146 | 0.006 | 6059169 | 4736028 (78.2%) | 994 | 970 | 39 |
| *Acacia auriculiformis* | Brown 154 (MEL) | ERS11697098 | This study | 0.151 | 0.006 | 5079017 | 4191953 (82.5%) | 994 | 964 | 38 |
| *Acacia ausfeldii* | Karunajeewa 1149 (MEL) | ERS11697099 | This study | 0.13 | 0.007 | 5487545 | 4044020 (73.7%) | 994 | 972 | 44 |
| *Acacia colei* var. *colei* | Murphy 326 (MELU) | ERS11697100 | This study | 0.158 | 0.006 | 6192508 | 4808345 (77.6%) | 993 | 939 | 46 |
| *Acacia deanei* subsp. *paucijuga* | Murphy 599 (MELU) | ERS11697101 | This study | 0.148 | 0.006 | 7159214 | 5527391 (77.2%) | 992 | 966 | 43 |
| *Acacia longifolia* | Koenen 182 (Z) | ERS4812840 | Koenen et al. 2020 | 0.012 | 0.018 | 3705539 | 115743 (3.1%) | 772 | 565 | 0 |
| *Acacia lycopodiifolia* | Murphy 339 (MELU) | ERS11697102 | This study | 0.149 | 0.006 | 2250348 | 1830789 (81.4%) | 993 | 966 | 19 |
| *Acacia montana* | Murphy 672 (MELU) | ERS11697103 | This study | 0.152 | 0.006 | 5357202 | 4096344 (76.5%) | 992 | 966 | 43 |
| *Acacia murrayana* | Ariati 110 (MELU) | ERS11697104 | This study | 0.148 | 0.005 | 5154241 | 3942099 (76.5%) | 992 | 960 | 48 |
| *Acacia oswaldii* | Murphy 573 (MELU) | ERS11697105 | This study | 0.153 | 0.005 | 5809958 | 4471048 (77%) | 992 | 965 | 47 |
| *Acacia platycarpa* | Murphy 327 (MELU) | ERS11697106 | This study | 0.152 | 0.005 | 4125748 | 3097650 (75.1%) | 992 | 969 | 38 |
| *Acacia pycnantha* | Murphy 670 (MELU) | ERS11697107 | This study | 0.17 | 0.006 | 7377943 | 5368248 (72.8%) | 991 | 963 | 46 |
| *Acacia pyrifolia* | Murphy 337 (MELU) | ERS11697108 | This study | 0.244 | 0.007 | 2763767 | 2293777 (83%) | 993 | 953 | 22 |
| *Acacia rostellifera* | Murphy 466 (MELU) | ERS11697109 | This study | 0.19 | 0.005 | 8302200 | 6384932 (76.9%) | 993 | 976 | 42 |
| *Acacia sibirica* | Murphy 486 (MEL) | ERS11697110 | This study | 0.159 | 0.005 | 6046537 | 4625699 (76.5%) | 988 | 954 | 55 |

| | | | | | | | | | | |
|---|---|---|---|---|---|---|---|---|---|---|
| *Acacia triptera* | Karunajeewa 1446 (MEL) | ERS11697111 | This study | 0.165 | 0.006 | 7506934 | 5657563 (75.4%) | 992 | 969 | 49 |
| *Acacia tumida* | Murphy 306 (MELU) | ERS11697112 | This study | 0.194 | 0.007 | 2415907 | 2004246 (83%) | 991 | 949 | 26 |
| *Acacia vernici-flua* | Karunajeewa 1012 (MEL) | ERS11697113 | This study | 0.137 | 0.006 | 3949459 | 3006398 (76.1%) | 992 | 972 | 33 |
| *Acacia victoriae* | Ariati 260 (MELU) | ERS11697114 | This study | 0.155 | 0.006 | 6876195 | 5437933 (79.1%) | 994 | 969 | 45 |
| *Acaciella villosa* | Hughes 2635 (FHO) | ERS4812841 | Koenen et al. 2020 | 0.037 | 0.02 | 2911372 | 74570 (2.6%) | 598 | 314 | 0 |
| *Acrocarpus fraxinifolius* | Manos 1416 (DUKE) | ERS11697115 | This study | 0.236 | 0.008 | 5998945 | 3865675 (64.4%) | 955 | 746 | 89 |
| *Adenanthera pavonina* | Ambriansyah and Arifin AA295 (K) | ERS4812842 | Koenen et al. 2020 | 0.016 | 0.018 | 4242045 | 164600 (3.9%) | 882 | 588 | 12 |
| *Adenopodia patens* | Sandoval MS343 (K) | ERS4812843 | Koenen et al. 2020 | 0.043 | 0.022 | 6434933 | 443279 (6.9%) | 968 | 778 | 8 |
| *Adenopodia scelerata* | Jongkind 10602 (WAG) | ERS4812844 | Koenen et al. 2020 | 0.047 | 0.008 | 9208280 | 707958 (7.7%) | 981 | 874 | 16 |
| *Afrocalliandra gilbertii* | Gillett and Hemming 24799 (PRE) | ERS11697116 | This study | 0.202 | 0.006 | 4355148 | 3628668 (83.3%) | 990 | 941 | 29 |
| *Afrocalliandra redacta* | Germishuizen 5585 (PRE) | ERS11697117 | This study | 0.248 | 0.006 | 7960105 | 6666175 (83.7%) | 992 | 947 | 27 |
| *Alantsiloden-dron glomera-tum* | Koenen 257 (G, K) | ERS11697118 | This study | 0.139 | 0.006 | 4313074 | 3157671 (73.2%) | 992 | 946 | 47 |
| *Alantsiloden-dron pilosum* | Koenen 203 (Z) | ERS4812845 | Koenen et al. 2020 | 0.023 | 0.015 | 3141103 | 202237 (6.4%) | 928 | 721 | 2 |
| *Alantsiloden-dron villosum* | Koenen 409 (G, K) | ERS11697119 | This study | 0.156 | 0.005 | 3698603 | 2946722 (79.7%) | 992 | 945 | 30 |
| *Albizia adian-thifolia* | Wieringa 6278 (WAG) | ERS4812846 | Koenen et al. 2020 | 0.029 | 0.008 | 9142035 | 461499 (5%) | 973 | 834 | 10 |
| *Albizia adino-cephala* | Hughes 1070 (FHO) | ERS11697120 | This study | 0.344 | 0.007 | 4675412 | 3431491 (73.4%) | 995 | 980 | 42 |
| *Albizia altissi-ma* | Jongkind 10709 (WAG) | ERS4812847 | Koenen et al. 2020 | 0.526 | 0.066 | 9157157 | 3852321 (42.1%) | 973 | 856 | 19 |
| *Albizia anthel-mintica* | Maurin 363 (JRAU) | ERS4812848 | Koenen et al. 2020 | 0.028 | 0.01 | 5165679 | 327512 (6.3%) | 970 | 831 | 8 |
| *Albizia ataka-taka* | Koenen 229 (Z) | ERS4812849 | Koenen et al. 2020 | 0.647 | 0.089 | 12500723 | 4350341 (34.8%) | 972 | 807 | 14 |
| *Albizia auris-parsa* | Koenen 230 (Z) | ERS4812850 | Koenen et al. 2020 | 0.031 | 0.01 | 14432059 | 1021898 (7.1%) | 986 | 898 | 18 |

| | | | | | | | | | | |
|---|---|---|---|---|---|---|---|---|---|---|
| *Albizia bernieri* | Koenen 354 (Z) | ERS4812851 | Koenen et al. 2020 | 0.016 | 0.016 | 3429090 | 142276 (4.1%) | 879 | 687 | 8 |
| *Albizia bertero-ana* | Jiménez Rodríguez 1107 (NY) | ERS11697121 | This study | 0.251 | 0.006 | 4737523 | 3786287 (79.9%) | 993 | 977 | 29 |
| *Albizia boivinii* | Koenen 270 (Z) | ERS4812852 | Koenen et al. 2020 | 0.028 | 0.013 | 3362669 | 234729 (7%) | 955 | 784 | 14 |
| *Albizia brevi-folia* | Maurin 826 (JRAU) | ERS4812853 | Koenen et al. 2020 | 0.023 | 0.02 | 2433285 | 156865 (6.4%) | 906 | 662 | 5 |
| *Albizia burkar-tiana* | Stival-Santos 678 (RB) | ERS4812854 | Koenen et al. 2020 | 0.031 | 0.014 | 5440692 | 242245 (4.5%) | 956 | 792 | 7 |
| *Albizia car-bonaria* | Daza 16353 (K) | ERS11697122 | This study | 0.233 | 0.006 | 10728752 | 8913272 (83.1%) | 994 | 977 | 35 |
| *Albizia coripat-ensis* | Hughes 2433 (FHO) | ERS11697123 | This study | 0.258 | 0.006 | 7841663 | 6094428 (77.7%) | 993 | 975 | 41 |
| *Albizia decan-dra* | Vilhena 231 (NY) | ERS11697124 | This study | 0.377 | 0.01 | 830632 | 644911 (77.6%) | 994 | 957 | 12 |
| *Albizia din-klagei* | Jongkind 7359 (WAG) | ERS4812855 | Koenen et al. 2020 | 0.01 | 0.017 | 1903228 | 75207 (4%) | 581 | 441 | 3 |
| *Albizia edwallii* | Dalmaso 272 (RB) | ERS4812856 | Koenen et al. 2020 | 0.02 | 0.008 | 3116764 | 180288 (5.8%) | 921 | 764 | 8 |
| *Albizia eri-orhachis* | Breteler 507 (WAG) | ERS11697167 | This study | 0.217 | 0.006 | 7065837 | 5992229 (84.8%) | 995 | 967 | 34 |
| *Albizia ferrug-inea* | Jongkind 10762 (WAG) | ERS4812857 | Koenen et al. 2020 | 0.033 | 0.008 | 6199048 | 343428 (5.5%) | 965 | 839 | 10 |
| *Albizia glabrip-etala* | Lewis 1652 (K) | ERS11697125 | This study | 0.562 | 0.011 | 1176536 | 881434 (74.9%) | 990 | 905 | 32 |
| *Albizia grandi-bracteata* | Koenen 159 (WAG) | ERS4812858 | Koenen et al. 2020 | 0.72 | 0.095 | 7971575 | 2894494 (36.3%) | 969 | 791 | 12 |
| *Albizia inun-data* | Wood 26530 (K) | ERS4812859 | Koenen et al. 2020 | 0.725 | 0.096 | 7376881 | 2521660 (34.2%) | 968 | 785 | 7 |
| *Albizia leonardii* | Zanoni 34986 (NY) | ERS11697126 | This study | 0.365 | 0.008 | 1201177 | 943730 (78.6%) | 993 | 896 | 30 |
| *Albizia lepto-phylla* | Brummitt 14035 (K) | ERS11697292 | This study | 0.168 | 0.006 | 5071059 | 4377041 (86.3%) | 994 | 968 | 28 |
| *Albizia mahalao* | Koenen 216 (Z) | ERS4812860 | Koenen et al. 2020 | 0.711 | 0.096 | 15008953 | 6126074 (40.8%) | 972 | 802 | 12 |
| *Albizia masiko-rorum* | Koenen 237 (Z) | ERS4812861 | Koenen et al. 2020 | 0.032 | 0.011 | 11494741 | 723554 (6.3%) | 988 | 895 | 15 |
| *Albizia multi-flora* | Hughes 3090 (Z) | ERS11697127 | This study | 0.225 | 0.006 | 4764898 | 3772512 (79.2%) | 994 | 968 | 28 |
| *Albizia nio-poides* | Simon 1601 (CEN) | ERS11697128 | This study | 0.218 | 0.006 | 5693507 | 4724671 (83%) | 995 | 972 | 22 |

| | | | | | | | | | | |
|---|---|---|---|---|---|---|---|---|---|---|
| *Albizia obbia-densis* | Thulin 4163 (UPS) | ERS4812862 | Koenen et al. 2020 | 0.031 | 0.012 | 4969768 | 331864 (6.7%) | 966 | 838 | 10 |
| *Albizia obliqui-foliolata* | Wieringa 6519 (WAG) | ERS4812863 | Koenen et al. 2020 | 0.559 | 0.063 | 4785416 | 2047233 (42.8%) | 972 | 840 | 14 |
| *Albizia poly-phylla* | Koenen 256 (Z) | ERS4812864 | Koenen et al. 2020 | 0.029 | 0.008 | 2976472 | 209635 (7%) | 945 | 769 | 15 |
| *Albizia retusa* | Hyland 2732 (L) | ERS4812865 | Koenen et al. 2020 | 0.032 | 0.011 | 10800530 | 702560 (6.5%) | 981 | 864 | 10 |
| *Albizia rhombi-folia* | Deighton 3618 (K) | ERS11697168 | This study | 0.216 | 0.007 | 3323568 | 2721152 (81.9%) | 995 | 969 | 33 |
| *Albizia saha-fariensis* | Koenen 405 (Z) | ERS4812866 | Koenen et al. 2020 | 0.031 | 0.018 | 11553872 | 734373 (6.4%) | 983 | 885 | 16 |
| *Albizia sapon-aria* | Jobson 1041 (BH) | ERS4812867 | Koenen et al. 2020 | 0.7 | 0.095 | 8862638 | 3243698 (36.6%) | 972 | 803 | 15 |
| *Albizia sinalo-ensis* | Hughes 1576 (K) | ERS11697130 | This study | 0.222 | 0.007 | 3073383 | 2471008 (80.4%) | 993 | 976 | 23 |
| *Albizia sub-dimidiata* var. *minor* | Gorts 341 (K) | ERS11697131 | This study | 0.527 | 0.012 | 649626 | 455201 (70.1%) | 993 | 966 | 19 |
| *Albizia sub-dimidiata* var. *subdimidiata* | Ferreira 210 (K) | ERS11697129 | This study | 0.259 | 0.005 | 4907457 | 4054305 (82.6%) | 993 | 966 | 24 |
| *Albizia tomen-tosa* | Hughes 648 (K) | ERS11697132 | This study | 0.237 | 0.006 | 3848755 | 3084913 (80.2%) | 994 | 972 | 23 |
| *Albizia umbel-lata* | Jobson 1037 (BH) | ERS4812882 | Koenen et al. 2020 | 0.536 | 0.067 | 9829794 | 4228923 (43%) | 971 | 854 | 20 |
| *Albizia um-bellata* subsp. *umbellata* | Larsen 33736 (K) | ERS11697169 | This study | 0.243 | 0.01 | 2937052 | 2603852 (88.7%) | 982 | 789 | 93 |
| *Albizia versi-color* | Maurin 560 (JRAU) | ERS4812868 | Koenen et al. 2020 | 0.719 | 0.086 | 13969835 | 5950552 (42.6%) | 969 | 806 | 9 |
| *Albizia viridis* | Du Puy M251 (K) | ERS4812869 | Koenen et al. 2020 | 0.029 | 0.013 | 6447985 | 426750 (6.6%) | 973 | 836 | 15 |
| *Albizia xero-phytica* | Hughes 1435 (K) | ERS11697133 | This study | 0.211 | 0.006 | 3486392 | 2827768 (81.1%) | 995 | 977 | 24 |
| *Albizia zygia* | Wieringa 5915 (WAG) | ERS4812870 | Koenen et al. 2020 | 0.03 | 0.008 | 7296600 | 381783 (5.2%) | 969 | 820 | 9 |
| *Amblygonocar-pus andongen-sis* | Sokpon 1451 (WAG) | ERS4812871 | Koenen et al. 2020 | 0.02 | 0.016 | 4673226 | 157163 (3.4%) | 864 | 527 | 12 |
| *Anadenanthera colubrina* | de Queiroz 15685 (HUEFS) | ERS4812872 | Koenen et al. 2020 | 0.024 | 0.012 | 3882704 | 231474 (6%) | 943 | 755 | 7 |

| | | | | | | | | | | |
|---|---|---|---|---|---|---|---|---|---|---|
| *Arapatiella psilophylla* | de Lima et al. 7906 (RB) | ERS11697134 | This study | 0.168 | 0.006 | 4701923 | 3236767 (68.8%) | 985 | 861 | 70 |
| *Archidendron bubalinum* | Latiff and Zainudin ALM3503 (L) | ERS11697135 | This study | 0.226 | 0.005 | 4606643 | 3935161 (85.4%) | 994 | 964 | 22 |
| *Archidendron clypearia* | Wieringa 1849 (WAG) | ERS11697136 | This study | 0.197 | 0.006 | 8533396 | 7182846 (84.2%) | 994 | 968 | 32 |
| *Archidendron ellipticum* sub-sp. *ellipticum* | Kalat ARK 42 (L) | ERS11697137 | This study | 0.191 | 0.006 | 6306349 | 5430400 (86.1%) | 994 | 967 | 29 |
| *Archidendron grandiflorum* | Clarkson 6233 (L) | ERS11697138 | This study | 0.178 | 0.008 | 7556417 | 5729504 (75.8%) | 995 | 968 | 43 |
| *Archidendron jiringa* | Annable 3321 (NY) | ERS11697139 | This study | 0.195 | 0.006 | 4588076 | 3900989 (85%) | 995 | 970 | 25 |
| *Archidendron kanisii* | Ford and Holmes AF3669 (L) | ERS11697140 | This study | 0.185 | 0.006 | 5970058 | 4833236 (81%) | 994 | 970 | 40 |
| *Archidendron lucidum* | Wang and Lin 2534 (L) | ERS4812873 | Koenen et al. 2020 | 0.014 | 0.01 | 5795037 | 316866 (5.5%) | 971 | 824 | 9 |
| *Archidendron ptenopum* | Takeuchi and Ama 15334 (L) | ERS11697141 | This study | 0.149 | 0.006 | 4613690 | 3511372 (76.1%) | 993 | 968 | 46 |
| *Archidendron quocense* | Newman 2094 (E) | ERS4812874 | Koenen et al. 2020 | 0.726 | 0.092 | 11417868 | 4456246 (39%) | 970 | 806 | 9 |
| *Archidendron triplinervium* | Church et al. 1171 (L) | ERS11697142 | This study | 0.218 | 0.006 | 5563658 | 4815056 (86.5%) | 994 | 973 | 18 |
| *Archidendropsis granulosa* | McKee 38353 (L) | ERS4812875 | Koenen et al. 2020 | 0.021 | 0.009 | 12082020 | 831488 (6.9%) | 987 | 910 | 12 |
| *Archidendropsis xanthoxylon* | Hyland 9229 (L) | ERS11697143 | This study | 0.141 | 0.005 | 4667843 | 3839349 (82.3%) | 993 | 966 | 30 |
| *Arcoa gonaven-sis* | Gardner and Knees 7026 (E) | ERS11697144 | This study | 0.182 | 0.007 | 9685695 | 6011856 (62.1%) | 939 | 662 | 212 |
| *Arquita tricho-carpa* | Gagnon 2018 (MT) | ERS11697145 | This study | 0.247 | 0.007 | 3176819 | 1627215 (51.2%) | 953 | 677 | 35 |
| *Aubrevillea kerstingii* | Nimba Botanic Team JR957 (WAG) | ERS4812876 | Koenen et al. 2020 | 0.015 | 0.016 | 5685788 | 231500 (4.1%) | 926 | 708 | 10 |
| *Balizia elegans* | Iganci 870 (RB) | ERS11697146 | This study | 0.215 | 0.006 | 7937229 | 5995772 (75.5%) | 995 | 955 | 46 |
| *Balizia leucoc-alyx* | Aguilar 1939 (NY) | ERS11697147 | This study | 0.22 | 0.006 | 5789748 | 4546951 (78.5%) | 994 | 970 | 47 |
| *Balizia pedicel-laris* | de Queiroz 15529 (HUEFS) | ERS4812877 | Koenen et al. 2020 | 0.503 | 0.07 | 11239481 | 3679071 (32.7%) | 974 | 859 | 20 |
| *Balizia* sp. nov. | Morim 577 (RB) | ERS4812878 | Koenen et al. 2020 | 0.541 | 0.064 | 7968724 | 3021322 (37.9%) | 971 | 849 | 19 |
| *Balsamocarpon brevifolium* | Eggli and Leuen-berger 2986 (Z) | ERS11697148 | This study | 0.254 | 0.009 | 4616413 | 1847991 (40%) | 955 | 732 | 54 |
| *Batesia flori-bunda* | Grenand 3032 (CAY) | ERS11697149 | This study | 0.233 | 0.006 | 492546 | 291743 (59.2%) | 934 | 649 | 27 |

| | | | | | | | | | | |
|---|---|---|---|---|---|---|---|---|---|---|
| *Biancaea decapetala* | Hughes 2227 (FHO) | ERS11697150 | This study | 0.276 | 0.007 | 6890313 | 4093095 (59.4%) | 964 | 767 | 42 |
| *Blanchetiodendron blanchetii* | de Queiroz 15616 (HUEFS) | ERS4812879 | Koenen et al. 2020 | 0.029 | 0.008 | 6107868 | 314085 (5.1%) | 970 | 828 | 10 |
| *Burkea africana* | Smith 6 (WAG) | ERS11697151 | This study | 0.233 | 0.006 | 2546859 | 1774273 (69.7%) | 983 | 853 | 47 |
| *Bussea perrieri* | Randrianasolo 527 (WAG) | ERS11697152 | This study | 0.173 | 0.005 | 3607830 | 2443747 (67.7%) | 983 | 834 | 52 |
| *Caesalpinia cassioides* | Pennington 789 (K) | ERS11697153 | This study | 0.248 | 0.006 | 5725554 | 3462449 (60.5%) | 971 | 749 | 41 |
| *Caesalpinia crista* | Herendeen 1 V 99 3 (US) | ERS11697155 | This study | 0.285 | 0.007 | 6059665 | 3807786 (62.8%) | 962 | 718 | 31 |
| *Calliandra bella* | de Queiroz 15696 (HUEFS) | ERS11697156 | This study | 0.214 | 0.007 | 8940737 | 7202271 (80.6%) | 992 | 944 | 37 |
| *Calliandra haematocephala* | Hughes 2604 (FHO) | ERS11697157 | This study | 0.199 | 0.007 | 5482960 | 4308847 (78.6%) | 986 | 945 | 38 |
| *Calliandra haematomma* | Kass 2008 1 (BH) | ERS11697158 | This study | 0.212 | 0.006 | 6493219 | 5542035 (85.4%) | 990 | 940 | 32 |
| *Calliandra hygrophila* | de Queiroz 15542 (HUEFS) | ERS4812880 | Koenen et al. 2020 | 0.039 | 0.014 | 3725309 | 211230 (5.7%) | 919 | 717 | 7 |
| *Calliandra parviflora* | Wood 26606 (K) | ERS11697159 | This study | 0.18 | 0.007 | 7464597 | 6167415 (82.6%) | 984 | 943 | 38 |
| *Calliandra sessilis* | de Queiroz 15608 (HUEFS) | ERS11697160 | This study | 0.204 | 0.006 | 5678535 | 4695359 (82.7%) | 988 | 931 | 39 |
| *Calliandra* sp. nov. | Poilane 9150 (P) | ERS11697161 | This study | 0.232 | 0.006 | 2706841 | 2297453 (84.9%) | 991 | 943 | 29 |
| *Calliandra viscidula* | de Queiroz 15541 (HUEFS) | ERS11697162 | This study | 0.209 | 0.006 | 6337439 | 5203387 (82.1%) | 988 | 950 | 27 |
| *Calliandropsis nervosa* | Hughes 1784 (K) | ERS11697163 | This study | 0.214 | 0.006 | 6190848 | 4841250 (78.2%) | 992 | 959 | 39 |
| *Calpocalyx dinklagei* | Wieringa 6094 (WAG) | ERS4812881 | Koenen et al. 2020 | 0.023 | 0.013 | 10150267 | 312067 (3.1%) | 936 | 692 | 10 |
| *Calpocalyx heitzii* | van Valkenburg 2877 (WAG) | ERS11697164 | This study | 0.162 | 0.006 | 2555176 | 1877054 (73.5%) | 989 | 889 | 33 |
| *Campsiandra comosa* | Iganci 856 (RB) | ERS11697165 | This study | 0.18 | 0.008 | 8560245 | 6097949 (71.2%) | 986 | 908 | 94 |
| *Cassia cowanii* var. *guianensis* | Redden 3277 (US) | ERS11697166 | This study | 0.182 | 0.007 | 7359511 | 4902385 (66.6%) | 973 | 795 | 46 |
| *Cedrelinga cateniformis* | Pennington 17761 (K) | ERS4812883 | Koenen et al. 2020 | 0.02 | 0.019 | 3588744 | 223591 (6.2%) | 950 | 768 | 6 |
| *Cenostigma pluviosum* var. *maraniona* | Hughes 3105 (MT) | ERS11697170 | This study | 0.257 | 0.007 | 8289046 | 4853867 (58.6%) | 970 | 782 | 62 |
| *Ceratonia siliqua* | Wieringa 3477 (WAG) | ERS11697171 | This study | 0.224 | 0.006 | 4232224 | 2964154 (70%) | 966 | 722 | 63 |
| *Chamaecrista adiantifolia* | Iganci 861 (RB) | ERS11697172 | This study | 0.146 | 0.007 | 2899104 | 1813687 (62.6%) | 969 | 777 | 55 |

| | | | | | | | | | | |
|---|---|---|---|---|---|---|---|---|---|---|
| *Chamaecrista lineata* | Bradley 32006 (US) | ERS11697173 | This study | 0.19 | 0.006 | 4065468 | 2504843 (61.6%) | 947 | 671 | 27 |
| *Chamaecrista ramosa* | Lewis 3845 (K) | ERS11697174 | This study | 0.198 | 0.005 | 4926849 | 3116425 (63.3%) | 945 | 696 | 36 |
| *Chamaecrista viscosa* | Wood 26658 (K) | ERS11697175 | This study | 0.195 | 0.006 | 4228490 | 2512358 (59.4%) | 945 | 702 | 37 |
| *Chidlowia sanguinea* | Wieringa 4338 (WAG) | ERS4812884 | Koenen et al. 2020 | 0.02 | 0.015 | 8052913 | 247743 (3.1%) | 923 | 664 | 11 |
| *Chloroleucon mangense* var. *mangense* | Lozano 1166 (K) | ERS11697176 | This study | 0.134 | 0.005 | 3472053 | 2708124 (78%) | 994 | 971 | 24 |
| *Chloroleucon tenuiflorum* | de Queiroz 15514 (HUEFS) | ERS4812885 | Koenen et al. 2020 | 0.018 | 0.01 | 6789783 | 419135 (6.2%) | 973 | 868 | 15 |
| *Cojoba arborea* | Simon 1545 (CEN) | ERS4812886 | Koenen et al. 2020 | 0.047 | 0.008 | 9087281 | 556186 (6.1%) | 982 | 871 | 15 |
| *Cojoba filipes* | Colella 1331 (NY) | ERS11697177 | This study | 0.278 | 0.006 | 5763912 | 4870940 (84.5%) | 994 | 960 | 26 |
| *Cojoba rufescens* | Castroviejo 14683 (K) | ERS11697178 | This study | 0.206 | 0.005 | 4589082 | 3566079 (77.7%) | 994 | 967 | 42 |
| *Cojoba zanonii* | Zanoni 36337 (NY) | ERS11697179 | This study | 0.186 | 0.007 | 3584825 | 3121762 (87.1%) | 993 | 962 | 21 |
| *Colvillea racemosa* | Bruneau 1403 (MT) | ERS11697180 | This study | 0.19 | 0.005 | 5746997 | 3834447 (66.7%) | 980 | 884 | 65 |
| *Conzattia multiflora* | Hughes 2071 (FHO) | ERS11697181 | This study | 0.186 | 0.005 | 4260881 | 2903786 (68.1%) | 977 | 869 | 48 |
| *Cordeauxia edulis* | Kuchar 17803 (K) | ERS11697182 | This study | 0.164 | 0.007 | 2922067 | 1681248 (57.5%) | 967 | 761 | 44 |
| *Coulteria platyloba* | MacQueen 178 (FHO) | ERS11697154 | This study | 0.26 | 0.008 | 7199957 | 4181262 (58.1%) | 965 | 738 | 152 |
| *Cylicodiscus gabunensis* | Sosef 645A (WAG) | ERS4812887 | Koenen et al. 2020 | 0.037 | 0.009 | 6104369 | 404128 (6.6%) | 964 | 770 | 15 |
| *Delonix decaryi* | Koenen 238 (G, K) | ERS11697183 | This study | 0.178 | 0.006 | 4250257 | 2742084 (64.5%) | 977 | 872 | 67 |
| *Delonix edule* | Willing s.n. (K) | ERS11697227 | This study | 0.19 | 0.006 | 4981753 | 3367148 (67.6%) | 979 | 864 | 50 |
| *Denisophytum madagascariense* | Bruneau et al. 1348 (MT) | ERS11697184 | This study | 0.217 | 0.01 | 2930272 | 1392506 (47.5%) | 959 | 715 | 33 |
| *Desmanthus acuminatus* | Hughes 2314 (FHO) | ERS11697185 | This study | 0.14 | 0.005 | 3176892 | 2421869 (76.2%) | 995 | 963 | 35 |
| *Desmanthus balsensis* | Hughes 1825 (FHO) | ERS11697186 | This study | 0.136 | 0.006 | 5883758 | 4369334 (74.3%) | 993 | 965 | 42 |
| *Desmanthus leptophyllus* | Hughes 2035 (FHO) | ERS4812888 | Koenen et al. 2020 | 0.019 | 0.018 | 4309291 | 193040 (4.5%) | 917 | 686 | 7 |
| *Desmanthus virgatus* | Wood 26551 (K) | ERS11697187 | This study | 0.148 | 0.005 | 5846308 | 4447399 (76.1%) | 992 | 965 | 51 |

| | | | | | | | | | | |
|---|---|---|---|---|---|---|---|---|---|---|
| *Dichrostachys cinerea* | Maurin 256 (JRAU) | ERS4812889 | Koenen et al. 2020 | 0.03 | 0.018 | 4143707 | 247851 (6%) | 942 | 720 | 3 |
| *Dichrostachys myriophylla* | Koenen 301 (G, K) | ERS11697188 | This study | 0.145 | 0.005 | 3010514 | 2241701 (74.5%) | 995 | 959 | 37 |
| *Dichrostachys paucifoliolata* | Luckow 4157 (BH) | ERS11697189 | This study | 0.157 | 0.006 | 7700124 | 5511552 (71.6%) | 993 | 958 | 51 |
| *Dichrostachys richardiana* | Koenen 282 (G, K) | ERS11697190 | This study | 0.127 | 0.006 | 3899707 | 3113125 (79.8%) | 992 | 955 | 32 |
| *Dichrostachys tenuifolia* | Labat 3579 (P) | ERS11697191 | This study | 0.141 | 0.006 | 3335610 | 2699005 (80.9%) | 993 | 943 | 25 |
| *Dichrostachys unijuga* | Koenen 242 (G, K) | ERS11697192 | This study | 0.145 | 0.006 | 3262061 | 2495849 (76.5%) | 993 | 940 | 36 |
| *Dimorphandra davisii* | Luckow 4593 (BH) | ERS11697193 | This study | 0.174 | 0.005 | 3658031 | 2579975 (70.5%) | 986 | 870 | 64 |
| *Dimorphandra gardneriana* | Hughes 2409 (FHO) | ERS11697194 | This study | 0.228 | 0.006 | 4727541 | 3206179 (67.8%) | 987 | 881 | 75 |
| *Dimorphandra macrostachya* | Iganci 877 (RB) | ERS4812890 | Koenen et al. 2020 | 0.014 | 0.017 | 6153872 | 144945 (2.4%) | 823 | 586 | 19 |
| *Dinizia jueira-na-facao* | Folli 4889 (HUEFS, K) | ERS11697195 | This study | 0.167 | 0.009 | 3254995 | 2291449 (70.4%) | 986 | 887 | 66 |
| *Diptychandra aurantiaca* | Wood 26513 (K) | ERS4812891 | Koenen et al. 2020 | 0.011 | 0.015 | 7891660 | 67216 (0.9%) | 463 | 262 | 2 |
| *Ebenopsis con-finis* | Hughes 1539 (FHO) | ERS4812892 | Koenen et al. 2020 | 0.042 | 0.019 | 4878310 | 253534 (5.2%) | 953 | 736 | 9 |
| *Elephantorrhiza burkei* | van der Bank 15 (JRAU) | ERS11697196 | This study | 0.236 | 0.006 | 4778607 | 3486450 (73%) | 994 | 928 | 41 |
| *Elephantorrhiza elephantina* | Komape, Mabe, and Siebert 198 (JRAU) | ERS4812893 | Koenen et al. 2020 | 0.02 | 0.008 | 6908210 | 277014 (4%) | 926 | 734 | 14 |
| *Entada africana* | van der Maesen 7144 (WAG) | ERS11697197 | This study | 0.185 | 0.006 | 6429832 | 4515988 (70.2%) | 988 | 925 | 38 |
| *Entada are-naria* | Bamps 8098 (WAG) | ERS11697198 | This study | 0.192 | 0.007 | 4085220 | 3196694 (78.3%) | 986 | 893 | 27 |
| *Entada pervillei* | Koenen 302 (G, K) | ERS11697199 | This study | 0.216 | 0.006 | 5885323 | 4655217 (79.1%) | 992 | 927 | 36 |
| *Entada polys-tachya* | Amaral Santos 3326 (CEN) | ERS11697200 | This study | 0.181 | 0.005 | 5920321 | 4327808 (73.1%) | 986 | 908 | 32 |
| *Entada rheedei* | Koenen 496 (Z) | ERS4812894 | Koenen et al. 2020 | 0.016 | 0.008 | 2451784 | 103654 (4.2%) | 808 | 615 | 5 |
| *Entada* sp. | van Beusekom et al. 4706 | ERS11697222 | This study | 0.406 | 0.037 | 209536 | 90355 (43.1%) | 794 | 472 | 7 |
| *Entada tuber-osa* | Koenen 417 (G, K) | ERS11697201 | This study | 0.183 | 0.005 | 3895325 | 2740535 (70.4%) | 987 | 917 | 40 |
| *Enterolobium barinense* | Blanco 157 (NY) | ERS11697202 | This study | 0.297 | 0.01 | 5647074 | 4652940 (82.4%) | 992 | 970 | 31 |
| *Enterolobium barnebianum* | Villa 1775 (K) | ERS11697203 | This study | 0.223 | 0.006 | 8653164 | 7178035 (83%) | 993 | 956 | 48 |

| | | | | | | | | | | |
|---|---|---|---|---|---|---|---|---|---|---|
| *Enterolobium contortisiliq-uum* | de Queiroz 15579 (HUEFS) | ERS4812895 | Koenen et al. 2020 | 0.013 | 0.017 | 8146204 | 642461 (7.9%) | 928 | 702 | 11 |
| *Enterolobium cyclocarpum* | MacQueen and Styles 75 (K) | ERS11697204 | This study | 0.225 | 0.007 | 9560067 | 8015575 (83.8%) | 994 | 975 | 34 |
| *Enterolobium gummiferum* | Harley 28284 (K) | ERS11697205 | This study | 0.247 | 0.006 | 9400120 | 7936398 (84.4%) | 994 | 974 | 28 |
| *Enterolobium maximum* | Nascimento 34 (K) | ERS11697206 | This study | 0.234 | 0.006 | 7827154 | 6486086 (82.9%) | 994 | 973 | 36 |
| *Erythrophleum ivorense* | Wieringa 5487 (WAG) | ERS4812896 | Koenen et al. 2020 | 0.023 | 0.007 | 10744241 | 337474 (3.1%) | 942 | 711 | 24 |
| *Erythrophleum teysmannii* | Smitinand 10468 (K) | ERS11697209 | This study | 0.175 | 0.007 | 2684620 | 1818272 (67.7%) | 987 | 890 | 50 |
| *Erythrostemon coluteifolius* | Gagnon 207 (MT) | ERS11697210 | This study | 0.268 | 0.007 | 2617653 | 1142755 (43.7%) | 949 | 689 | 33 |
| *Erythrostemon mexicanus* | Gagnon 2010 015 (MT) | ERS11697211 | This study | 0.251 | 0.009 | 3472543 | 1723736 (49.6%) | 956 | 672 | 22 |
| *Faidherbia albida* | Maurin 3495 (JRAU) | ERS4812897 | Koenen et al. 2020 | 0.044 | 0.011 | 5705125 | 315581 (5.5%) | 963 | 793 | 9 |
| *Falcataria moluccana* | Ambri and Arifin W826A (K) | ERS4812898 | Koenen et al. 2020 | 0.032 | 0.015 | 6552373 | 420103 (6.4%) | 976 | 829 | 10 |
| *Fillaeopsis dis-cophora* | Wieringa 5498 (WAG) | ERS4812899 | Koenen et al. 2020 | 0.014 | 0.025 | 2028878 | 61595 (3%) | 487 | 279 | 0 |
| *Gagnebina commersoniana* | Koenen 374 (G, K) | ERS11697212 | This study | 0.153 | 0.005 | 4924185 | 3728358 (75.7%) | 993 | 959 | 37 |
| *Gelrebia ros-trata* | 6th International Legume Conference 5 (JRAU) | ERS11697213 | This study | 0.266 | 0.01 | 3977542 | 1979853 (49.8%) | 942 | 638 | 24 |
| *Gleditsia chin-ensis* | Koenen 604 (Z) | ERS11697214 | This study | 0.103 | 0.009 | 1852772 | 1015332 (54.8%) | 956 | 723 | 74 |
| *Guilandina bonduc* | Herendeen and Mbago 9 XII 97 3 (US) | ERS11697215 | This study | 0.256 | 0.008 | 3088619 | 1411360 (45.7%) | 962 | 763 | 34 |
| *Gymnocladus dioicus* | Jongkind and Wie-ringa 4426 (WAG) | ERS11697216 | This study | 0.185 | 0.005 | 4530437 | 2849766 (62.9%) | 963 | 782 | 96 |
| *Haematoxylum brasiletto* | Gagnon 2010 013 (MT) | ERS11697217 | This study | 0.343 | 0.008 | 7115301 | 4248340 (59.7%) | 959 | 710 | 40 |
| *Havardia cam-pylacantha* | Hughes 1404 (K) | ERS11697218 | This study | 0.186 | 0.006 | 5858307 | 5056883 (86.3%) | 996 | 945 | 28 |
| *Havardia pal-lens* | Hughes 2138 (FHO) | ERS4812900 | Koenen et al. 2020 | 0.042 | 0.012 | 5849724 | 298968 (5.1%) | 967 | 809 | 13 |
| *Hererolandia pearsonii* | Kolberg and Loots HK 1399 (K) | ERS11697219 | This study | 0.258 | 0.006 | 9555894 | 5524893 (57.8%) | 957 | 791 | 55 |
| *Hesperalbizzia occidentalis* | Hughes 1296 (FHO) | ERS4812901 | Koenen et al. 2020 | 0.022 | 0.006 | 5078817 | 321941 (6.3%) | 964 | 824 | 7 |
| *Heteroflorum sclerocarpum* | Hughes 1849 (FHO) | ERS11697220 | This study | 0.228 | 0.007 | 4860825 | 3374699 (69.4%) | 978 | 834 | 42 |

| | | | | | | | | | | |
|---|---|---|---|---|---|---|---|---|---|---|
| *Hoffmannseg-gia arequipen-sis* | Hughes 2342 (FHO) | ERS11697221 | This study | 0.241 | 0.008 | 5218642 | 3131359 (60%) | 942 | 640 | 16 |
| *Hydrochorea corymbosa* (1) | Bonadeu 655 (RB) | ERS4812902 | Koenen et al. 2020 | 0.705 | 0.091 | 8772810 | 2969459 (33.8%) | 970 | 799 | 12 |
| *Hydrochorea corymbosa* (2) | Iganci 862 (RB) | ERS4812903 | Koenen et al. 2020 | 0.537 | 0.071 | 7443935 | 3123121 (42%) | 971 | 850 | 18 |
| *Hydrochorea gonggrijpii* | Tillett 45696 (K) | ERS11697223 | This study | 0.22 | 0.006 | 5304614 | 4552594 (85.8%) | 994 | 960 | 26 |
| *Hydrochorea marginata* | Morim 563 (RB) | ERS11697224 | This study | 0.222 | 0.006 | 8308280 | 6365865 (76.6%) | 994 | 965 | 50 |
| *Indopiptadenia oudhensis* | Adhikari, Poulsen, and Parmar BAG31 (E) | ERS11697225 | This study | 0.165 | 0.006 | 4741919 | 3591620 (75.7%) | 943 | 678 | 295 |
| *Inga alata* | Coley and Kursar TAKPDC1673 | SAMEA3283847 | Nicholls et al. 2015 | 0.07 | 0.008 | 1310260 | 586851 (44.8%) | 962 | 827 | 23 |
| *Inga alba* | Coley and Kursar TAKPDC1677 (UT) | ERR776844 | Koenen et al. 2020 | 0.062 | 0.009 | 1496500 | 613302 (41%) | 969 | 822 | 27 |
| *Inga auristellae* | Coley and Kursar TAKPDC1681 (UT) | SAMEA3283845 | Nicholls et al. 2015 | 0.06 | 0.01 | 1369861 | 593606 (43.3%) | 966 | 821 | 31 |
| *Inga bourgonii* | Coley and Kursar TAKPDC1688 (UT) | SAMEA3283822 | Nicholls et al. 2015 | 0.055 | 0.01 | 1214463 | 543496 (44.8%) | 970 | 823 | 27 |
| *Inga brevipes* | Coley and Kursar TAKPDC1694 (UT) | SAMEA3283838 | Nicholls et al. 2015 | 0.011 | 0.011 | 1275553 | 527411 (41.3%) | 966 | 832 | 32 |
| *Inga cin-namomea* | Dexter 465 (E, MOL) | SAMEA3283820 | Nicholls et al. 2015 | 0.069 | 0.009 | 1462414 | 626418 (42.8%) | 968 | 822 | 33 |
| *Inga cylindrica* | Coley and Kursar TAKPDC1713 (UT) | SAMEA3283801 | Nicholls et al. 2015 | 0.067 | 0.009 | 1398145 | 617850 (44.2%) | 966 | 815 | 25 |
| *Inga edulis* | Coley and Kursar TAKPDC1719 (UT) | ERR776838 | Koenen et al. 2020 | 0.058 | 0.011 | 1427222 | 617452 (43.3%) | 965 | 819 | 25 |
| *Inga hetero-phylla* | Dexter 345 (E, MOL) | SAMEA3283844 | Nicholls et al. 2015 | 0.075 | 0.011 | 1217628 | 536299 (44%) | 967 | 813 | 25 |
| *Inga huberi* | Coley and Kursar TAKPDC1755 (UT) | ERR776810 | Koenen et al. 2020 | 0.058 | 0.011 | 1350368 | 593530 (44%) | 968 | 812 | 24 |
| *Inga laurina* | Dexter 398 (E) | ERR776816 | Koenen et al. 2020 | 0.078 | 0.009 | 1381539 | 571714 (41.4%) | 970 | 813 | 28 |
| *Inga leiocaly-cina* | Dexter 355 (E, MOL) | SAMEA3283810 | Nicholls et al. 2015 | 0.073 | 0.008 | 1325708 | 566598 (42.7%) | 966 | 817 | 27 |

| | | | | | | | | | | |
|---|---|---|---|---|---|---|---|---|---|---|
| *Inga longiflora* | Coley and Kursar TAKPDC1788 (UT) | SAMEA3283836 | Nicholls et al. 2015 | 0.066 | 0.009 | 1253600 | 543896 (43.4%) | 967 | 812 | 27 |
| *Inga marginata* | BCI 8582 | SAMEA3283827 | Nicholls et al. 2015 | 0.017 | 0.008 | 968533 | 394059 (40.7%) | 964 | 844 | 30 |
| *Inga nouragensis* | Coley and Kursar TAKPDC1819 (UT) | SAMEA3283830 | Nicholls et al. 2015 | 0.062 | 0.011 | 1273976 | 555161 (43.6%) | 969 | 831 | 26 |
| *Inga pezizifera* | BCI 8577 | SAMEA3283806 | Nicholls et al. 2015 | 0.017 | 0.007 | 790785 | 325289 (41.1%) | 961 | 832 | 25 |
| *Inga punctata* | BCI 8580 | SAMEA3283828 | Nicholls et al. 2015 | 0.016 | 0.009 | 1008652 | 387572 (38.4%) | 969 | 838 | 28 |
| *Inga ruiziana* | BCI 8589 | SAMEA3283824 | Nicholls et al. 2015 | 0.008 | 0.008 | 749402 | 243746 (32.5%) | 961 | 826 | 24 |
| *Inga sapindoides* | BCI 97 | SAMEA3283803 | Nicholls et al. 2015 | 0.072 | 0.008 | 1369321 | 514025 (37.5%) | 965 | 808 | 29 |
| *Inga setosa* | Dexter 343 (E, MOL) | SAMEA3283850 | Nicholls et al. 2015 | 0.055 | 0.011 | 1109681 | 479709 (43.2%) | 969 | 825 | 18 |
| *Inga stipularis* | Coley and Kursar TAKPDC1856 (UT) | ERR776821 | Koenen et al. 2020 | 0.067 | 0.008 | 1497618 | 629753 (42.1%) | 965 | 834 | 28 |
| *Inga tenuistipula* | Dexter 110 (E) | ERR776831 | Koenen et al. 2020 | 0.071 | 0.008 | 1246734 | 543681 (43.6%) | 967 | 822 | 29 |
| *Inga thibaudiana* | Coley and Kursar TAKPDC1859 (UT) | SAMEA3283785 | Nicholls et al. 2015 | 0.064 | 0.01 | 1302945 | 552562 (42.4%) | 966 | 816 | 31 |
| *Inga umbellifera* | BCI 103 | SAMEA3283799 | Nicholls et al. 2015 | 0.011 | 0.01 | 1355642 | 546555 (40.3%) | 964 | 841 | 32 |
| *Jacqueshuberia brevipes* | Redden 1240 (US) | ERS11697226 | This study | 0.231 | 0.005 | 5380639 | 3913315 (72.7%) | 980 | 861 | 58 |
| *Jupunba abbottii* | Zanoni 21220 (NY) | ERS11697071 | This study | 0.196 | 0.008 | 3747723 | 3197746 (85.3%) | 995 | 968 | 31 |
| *Jupunba aspleniifolia* | Ekman 6383 (NY) | ERS11697072 | This study | 0.442 | 0.009 | 959289 | 796982 (83.1%) | 994 | 954 | 24 |
| *Jupunba barbouriana* | Iganci 847 (RB) | ERS11697073 | This study | 0.233 | 0.006 | 7284640 | 5844453 (80.2%) | 993 | 962 | 38 |
| *Jupunba brachystachya* | de Lima 7438 (RB) | ERS11697074 | This study | 0.19 | 0.006 | 5455545 | 4232261 (77.6%) | 994 | 973 | 46 |
| *Jupunba cochleata* | Bonadeu 673 (RB) | ERS11697076 | This study | 0.18 | 0.006 | 5895755 | 4445955 (75.4%) | 994 | 969 | 46 |
| *Jupunba commutata* | Maguire 46145 (NY) | ERS11697077 | This study | 0.237 | 0.006 | 5221538 | 4211275 (80.7%) | 994 | 968 | 41 |
| *Jupunba filamentosa* | de Lima 7487 (RB) | ERS11697079 | This study | 0.188 | 0.006 | 5318746 | 4121885 (77.5%) | 993 | 965 | 50 |

| | | | | | | | | | | |
|---|---|---|---|---|---|---|---|---|---|---|
| *Jupunba floribunda* | Iganci 883 (RB) | ERS11697080 | This study | 0.156 | 0.005 | 5803035 | 4578418 (78.9%) | 994 | 966 | 61 |
| *Jupunba idiopoda* | Quesada 1718 (NY) | ERS11697081 | This study | 0.195 | 0.005 | 3232056 | 2713116 (83.9%) | 993 | 961 | 28 |
| *Jupunba laeta* | Mori 25147 (NY) | ERS11697083 | This study | 0.268 | 0.007 | 1665213 | 1237759 (74.3%) | 994 | 965 | 24 |
| *Jupunba langsdorffii* | Ribeiro 728 (RB) | ERS11697084 | This study | 0.178 | 0.006 | 5605517 | 4246609 (75.8%) | 992 | 971 | 45 |
| *Jupunba leucophylla* | Iganci 839 (RB) | ERS11697086 | This study | 0.24 | 0.006 | 10089905 | 7755339 (76.9%) | 993 | 959 | 62 |
| *Jupunba longipedunculata* | Cardona 2682 (NY) | ERS11697088 | This study | 0.295 | 0.006 | 3933788 | 3191531 (81.1%) | 994 | 961 | 30 |
| *Jupunba macradenia* | Lourteig 3021 (NY) | ERS11697089 | This study | 0.337 | 0.007 | 3391013 | 2796383 (82.5%) | 994 | 966 | 32 |
| *Jupunba microcalyx* | Iganci 855 (RB) | ERS11697090 | This study | 0.187 | 0.006 | 7478422 | 5711833 (76.4%) | 994 | 962 | 51 |
| *Jupunba nipensis* | Mayo 19662 (NY) | ERS11697091 | This study | 0.424 | 0.008 | 810725 | 605585 (74.7%) | 993 | 958 | 21 |
| *Jupunba oppositifolia* | Liegier 16014 (NY) | ERS11697092 | This study | 0.302 | 0.007 | 3447350 | 2868985 (83.2%) | 994 | 965 | 35 |
| *Jupunba oxyphyllidia* | Yonker 6157 (NY) | ERS11697093 | This study | 0.236 | 0.006 | 4429470 | 3679176 (83.1%) | 993 | 967 | 30 |
| *Jupunba rhombea* | Iganci 261 (RB) | ERS11697087 | This study | 0.188 | 0.006 | 5403648 | 4148748 (76.8%) | 993 | 970 | 47 |
| *Jupunba trapezifolia* var. *micradenia* | Simon 1600 (CEN) | ERS4812839 | Koenen et al. 2020 | 0.54 | 0.066 | 6111172 | 2117308 (34.6%) | 968 | 832 | 18 |
| *Jupunba villosa* | Borges 423 (RB) | ERS11697095 | This study | 0.194 | 0.007 | 6341126 | 4848824 (76.5%) | 995 | 968 | 43 |
| *Kanaloa kahoolawensis* | Lorence 7380 (PTBG) | ERS4812904 | Koenen et al. 2020 | 0.036 | 0.008 | 11308250 | 1138422 (10.1%) | 983 | 926 | 20 |
| *Lachesiodendron viridiflorum* | de Queiroz 15614 (HUEFS) | ERS4812905 | Koenen et al. 2020 | 0.04 | 0.009 | 16885363 | 1108078 (6.6%) | 985 | 933 | 26 |
| *Lemurodendron capuronii* | Koenen 435 (Z) | ERS4812906 | Koenen et al. 2020 | 0.029 | 0.013 | 6336745 | 487016 (7.7%) | 955 | 631 | 103 |
| *Leucaena trichandra* | Hughes 1128 (FHO) | ERS11697228 | This study | 0.171 | 0.005 | 7893619 | 5938340 (75.2%) | 949 | 816 | 462 |
| *Leucochloron bolivianum* | Hughes 2608 (FHO) | ERS4812907 | Koenen et al. 2020 | 0.023 | 0.009 | 7312893 | 334766 (4.6%) | 972 | 841 | 13 |
| *Leucochloron limae* | Chase 8250 (K) | ERS4812908 | Koenen et al. 2020 | 0.019 | 0.01 | 7205109 | 443002 (6.1%) | 976 | 848 | 13 |
| *Libidibia glabrata* | Lewis and Lozano 3073 (K) | ERS11697229 | This study | 0.244 | 0.008 | 4178019 | 2495534 (59.7%) | 958 | 696 | 21 |
| *Lophocarpinia aculeatifolia* | Vogt 1321 (G) | ERS11697230 | This study | 0.246 | 0.008 | 7044959 | 3807972 (54.1%) | 951 | 721 | 43 |

| | | | | | | | | | | |
|---|---|---|---|---|---|---|---|---|---|---|
| *Lysiloma candidum* | Marazzi 300 (ASU) | ERS4812909 | Koenen et al. 2020 | 0.005 | 0.019 | 1872260 | 35152 (1.9%) | 154 | 93 | 0 |
| *Lysiloma latisiliquum* | Pennington 9197 (K) | ERS11697231 | This study | 0.104 | 0.008 | 4349268 | 3444588 (79.2%) | 995 | 972 | 35 |
| *Macrosamanea amplissima* | Bonadeu 663 (RB) | ERS4812910 | Koenen et al. 2020 | 0.017 | 0.007 | 2228110 | 101748 (4.6%) | 758 | 584 | 3 |
| *Macrosamanea discolor* | Wurdack 42732 (K) | ERS11697232 | This study | 0.15 | 0.008 | 2575768 | 2112820 (82%) | 992 | 943 | 32 |
| *Macrosamanea duckei* | Simon 1646 (CEN) | ERS11697233 | This study | 0.179 | 0.005 | 3977417 | 3260309 (82%) | 990 | 948 | 34 |
| *Macrosamanea kegelii* | Prevost 1721 (NY) | ERS11697234 | This study | 0.218 | 0.006 | 2335405 | 1871666 (80.1%) | 990 | 954 | 25 |
| *Macrosamanea prancei* | da Silva 183 (NY) | ERS11697235 | This study | 0.229 | 0.007 | 3118916 | 2600083 (83.4%) | 989 | 934 | 24 |
| *Macrosamanea simabifolia* | Iganci 881 (RB) | ERS11697236 | This study | 0.181 | 0.006 | 6849903 | 4840869 (70.7%) | 994 | 960 | 64 |
| *Macrosamanea spruceana* | Steyermark 87793 (NY) | ERS11697237 | This study | 0.435 | 0.01 | 2262225 | 1781741 (78.8%) | 991 | 940 | 31 |
| *Mariosousa sericea* | Chase 18949 (K) | ERS4812911 | Koenen et al. 2020 | 0.037 | 0.007 | 7488756 | 682808 (9.1%) | 982 | 853 | 14 |
| *Melanoxylon brauna* | Lopes and Andrade 113 (K) | ERS11697238 | This study | 0.188 | 0.006 | 4898735 | 3379585 (69%) | 978 | 801 | 56 |
| *Mezoneuron kauaiensis* | Lorence and Wagner 8904 (PTBG) | ERS11697239 | This study | 0.27 | 0.008 | 5356803 | 3183152 (59.4%) | 962 | 749 | 37 |
| *Mimosa ceratonia* | de Queiroz 15484 (HUEFS) | ERS11697240 | This study | 0.14 | 0.005 | 2841672 | 2277796 (80.2%) | 988 | 923 | 28 |
| *Mimosa dolens* | Simon 879 (FHO) | ERS11697241 | This study | 0.143 | 0.006 | 4467084 | 3367839 (75.4%) | 956 | 683 | 148 |
| *Mimosa grandidieri* | Koenen 207 (Z) | ERS4812912 | Koenen et al. 2020 | 0.051 | 0.008 | 7052923 | 400295 (5.7%) | 963 | 792 | 12 |
| *Mimosa hexandra* | Wood 26499 (K) | ERS11697242 | This study | 0.138 | 0.007 | 3213510 | 2288855 (71.2%) | 989 | 911 | 42 |
| *Mimosa hondurana* | Simon 858 (MEXU) | ERS11697243 | This study | 0.139 | 0.007 | 3583743 | 2590167 (72.3%) | 983 | 929 | 36 |
| *Mimosa invisa* | Simon 715 (FHO) | ERS11697244 | This study | 0.144 | 0.006 | 4871784 | 3636864 (74.7%) | 985 | 919 | 25 |
| *Mimosa myriadenia* | Iganci 835 (RB) | ERS11697245 | This study | 0.148 | 0.006 | 4099090 | 3053023 (74.5%) | 991 | 958 | 27 |
| *Mimosa pigra* | Simon 820 (MEXU) | ERS11697246 | This study | 0.159 | 0.007 | 7042106 | 4955742 (70.4%) | 985 | 840 | 49 |
| *Mimosa pudica* | Simon 1480 (CEN) | ERS11697247 | This study | 0.182 | 0.005 | 7218521 | 5309829 (73.6%) | 971 | 813 | 30 |
| *Mimosa revoluta* | Hughes 3051 (Z) | ERS11697248 | This study | 0.142 | 0.005 | 3058656 | 2380707 (77.8%) | 991 | 957 | 29 |
| *Mimosa rubicaulis* subsp. *himalayana* | Thomas SM 24 1 (K) | ERS11697249 | This study | 0.165 | 0.006 | 5237948 | 3929708 (75%) | 988 | 932 | 28 |

| | | | | | | | | | | |
|---|---|---|---|---|---|---|---|---|---|---|
| *Mimosa speciosissima* | Simon 753 (FHO) | ERS11697250 | This study | 0.148 | 0.006 | 3608125 | 2665822 (73.9%) | 978 | 889 | 23 |
| *Mimosa tenuiflora* | de Queiroz 15498 (HUEFS) | ERS4812913 | Koenen et al. 2020 | 0.051 | 0.008 | 5629617 | 250289 (4.4%) | 927 | 733 | 10 |
| *Mimosa tequilana* | Simon 813 (MEXU) | ERS11697251 | This study | 0.137 | 0.005 | 5364099 | 3924410 (73.2%) | 979 | 882 | 33 |
| *Mimosa tricephala* | Simon 849 (MEXU) | ERS11697252 | This study | 0.139 | 0.006 | 5195320 | 3729774 (71.8%) | 946 | 719 | 223 |
| *Mimozyganthus carinatus* | Hughes 2476 (FHO) | ERS4812914 | Koenen et al. 2020 | 0.037 | 0.01 | 7315427 | 404905 (5.5%) | 962 | 805 | 14 |
| *Moldenhawera floribunda* | Klitgaard 30 (K) | ERS11697253 | This study | 0.171 | 0.007 | 2169956 | 1202944 (55.4%) | 979 | 818 | 40 |
| *Mora gonggrijpii* | Breteler 13792 (WAG) | ERS11697254 | This study | 0.205 | 0.008 | 7112635 | 4980946 (70%) | 984 | 874 | 85 |
| *Moullava spicata* | Gillis 9504 (MO) | ERS11697255 | This study | 0.412 | 0.018 | 198073 | 95544 (48.2%) | 813 | 324 | 6 |
| *Neptunia oleracea* | Koenen 283 (Z) | ERS4812915 | Koenen et al. 2020 | 0.029 | 0.032 | 8812926 | 529920 (6%) | 977 | 795 | 13 |
| *Newtonia hildebrandtii* | Maurin 2457 (JRAU) | ERS4812916 | Koenen et al. 2020 | 0.039 | 0.008 | 7884530 | 415593 (5.3%) | 966 | 761 | 19 |
| *Pachyelasma tessmannii* | Wieringa 5229 (WAG) | ERS4812917 | Koenen et al. 2020 | 0.021 | 0.009 | 10879659 | 415320 (3.8%) | 959 | 757 | 25 |
| *Painteria elachistophylla* | Guzman Cruz UG2169 (K) | ERS11697256 | This study | 0.2 | 0.006 | 4999242 | 3952300 (79.1%) | 993 | 957 | 36 |
| *Painteria leptophylla* | Tenorio and Manriquez 4067 (MEXU) | ERS11697257 | This study | 0.217 | 0.006 | 7125144 | 5829138 (81.8%) | 995 | 971 | 28 |
| *Parapiptadenia excelsa* | Hughes 2451 (FHO) | ERS11697258 | This study | 0.135 | 0.005 | 2284574 | 1771891 (77.6%) | 993 | 953 | 32 |
| *Parapiptadenia zehntneri* | de Queiroz 15692 (HUEFS) | ERS4812918 | Koenen et al. 2020 | 0.032 | 0.017 | 3906807 | 170055 (4.4%) | 898 | 701 | 7 |
| *Pararchidendron pruinosum* | Jobson 1039 (BH) | ERS4812919 | Koenen et al. 2020 | 0.018 | 0.007 | 7208030 | 382671 (5.3%) | 969 | 840 | 11 |
| *Parasenegalia visco* | Fortunato 7645 (SI) | ERS11697302 | This study | 0.14 | 0.006 | 7345791 | 5884473 (80.1%) | 993 | 971 | 38 |
| *Paraserianthes lophantha* subsp. *lophantha* | van Slageren and Newton MSRN648 (K) | ERS4812920 | Koenen et al. 2020 | 0.017 | 0.008 | 5986133 | 302423 (5.1%) | 964 | 827 | 9 |
| *Parkia bahiae* | de Queiroz 15699 (HUEFS) | ERS11697259 | This study | 0.153 | 0.005 | 7977371 | 6329950 (79.3%) | 993 | 970 | 48 |
| *Parkia bicolor* | Wieringa 6265 (WAG) | ERS11697260 | This study | 0.132 | 0.006 | 4451222 | 3569411 (80.2%) | 992 | 969 | 39 |
| *Parkia igneiflora* | Iganci 885 (RB) | ERS11697261 | This study | 0.148 | 0.005 | 4751363 | 3705148 (78%) | 991 | 966 | 47 |
| *Parkia panurensis* | Iganci 842 (RB) | ERS4812921 | Koenen et al. 2020 | 0.024 | 0.019 | 2295962 | 125886 (5.5%) | 837 | 582 | 4 |

| | | | | | | | | | | |
|---|---|---|---|---|---|---|---|---|---|---|
| *Parkia pendula* | Simon 1194 (CEN) | ERS11697262 | This study | 0.179 | 0.006 | 5659537 | 4390458 (77.6%) | 993 | 964 | 44 |
| *Parkia timori-ana* | Murphy 351 (NY) | ERS11697263 | This study | 0.144 | 0.007 | 4739305 | 3697930 (78%) | 989 | 967 | 41 |
| *Parkia ulei* | Poncy s.n. (K) | ERS11697264 | This study | 0.156 | 0.006 | 6025015 | 4186376 (69.5%) | 994 | 974 | 59 |
| *Parkinsonia andicola* | Hughes 2619 (FHO) | ERS11697265 | This study | 0.176 | 0.006 | 4517120 | 2911215 (64.4%) | 984 | 885 | 70 |
| *Paubrasilia echinata* | Filgueiras 3391 (NY) | ERS11697266 | This study | 0.242 | 0.007 | 3580881 | 1992192 (55.6%) | 973 | 768 | 32 |
| *Peltophorum africanum* | Koenen 601 (Z) | ERS4812922 | Koenen et al. 2020 | 0.009 | 0.026 | 2589158 | 45554 (1.8%) | 368 | 142 | 2 |
| *Peltophorum dubium* | Hughes 2436 (FHO) | ERS11697267 | This study | 0.182 | 0.006 | 7352587 | 5207658 (70.8%) | 984 | 861 | 57 |
| *Pentaclethra macroloba* | Boyle 6720 (K) | ERS11697268 | This study | 0.153 | 0.006 | 3213908 | 2491507 (77.5%) | 989 | 903 | 29 |
| *Pentaclethra macrophylla* | Galeuchet & Balth-azar 10 (Z) | ERS4812923 | Koenen et al. 2020 | 0.024 | 0.008 | 16663175 | 524022 (3.1%) | 968 | 780 | 16 |
| *Piptadenia buchtienii* | Hughes 2427 (FHO) | ERS11697269 | This study | 0.135 | 0.006 | 4154987 | 3328448 (80.1%) | 993 | 967 | 38 |
| *Piptadenia robusta* | Luckow 4633 (BH) | ERS4812924 | Koenen et al. 2020 | 0.038 | 0.011 | 3153147 | 216876 (6.9%) | 926 | 687 | 22 |
| *Piptadenia uaupensis* | Mori & Ishikawa 20836 (K) | ERS11697270 | This study | 0.133 | 0.006 | 4852695 | 4048317 (83.4%) | 990 | 966 | 39 |
| *Piptadenias-trum africanum* | Koenen 152 (WAG) | ERS4812925 | Koenen et al. 2020 | 0.018 | 0.008 | 8360522 | 385173 (4.6%) | 953 | 757 | 15 |
| *Piptadeniopsis lomentifera* | Luckow 4505 (BH) | ERS4812926 | Koenen et al. 2020 | 0.026 | 0.007 | 5891872 | 427637 (7.3%) | 973 | 812 | 9 |
| *Pithecellobium dulce* | Marazzi 309 (ASU) | ERS4812927 | Koenen et al. 2020 | 0.036 | 0.007 | 6013147 | 372328 (6.2%) | 974 | 835 | 14 |
| *Pithecellobium excelsum* | Hughes 3101 (Z) | ERS11697271 | This study | 0.187 | 0.006 | 4722927 | 3840361 (81.3%) | 994 | 969 | 36 |
| *Pithecellobium hymenaeifoli-um* | Arvigo 216 (NY) | ERS11697272 | This study | 0.216 | 0.005 | 5649519 | 4686247 (82.9%) | 993 | 968 | 33 |
| *Pithecellobium keyense* | Chase 8958 (K) | ERS11697273 | This study | 0.217 | 0.005 | 6251735 | 4912041 (78.6%) | 994 | 968 | 39 |
| *Pithecellobium macrandrium* | Aguilar 1894 (NY) | ERS11697274 | This study | 0.214 | 0.006 | 4881101 | 3956361 (81.1%) | 993 | 962 | 40 |
| *Pityrocarpa moniliformis* | Wood 26516 (K) | ERS4812928 | Koenen et al. 2020 | 0.041 | 0.014 | 5395524 | 221123 (4.1%) | 929 | 753 | 16 |
| *Plathymenia reticulata* | de Queiroz 15688 (HUEFS) | ERS4812929 | Koenen et al. 2020 | 0.009 | 0.019 | 2196912 | 116395 (5.3%) | 822 | 563 | 5 |
| *Pomaria jamesii* | Gagnon 2010 020 (MT) | ERS11697275 | This study | 0.228 | 0.007 | 2454719 | 1057749 (43.1%) | 944 | 622 | 21 |

| | | | | | | | | | | |
|---|---|---|---|---|---|---|---|---|---|---|
| *Prosopidastrum globosum* | Luckow s.n. (BH) | ERS4812930 | Koenen et al. 2020 | 0.033 | 0.018 | 5321583 | 337387 (6.3%) | 962 | 775 | 7 |
| *Prosopis africana* | Essou 2110 (WAG) | ERS4812931 | Koenen et al. 2020 | 0.033 | 0.009 | 10475738 | 879557 (8.4%) | 985 | 854 | 15 |
| *Prosopis argentina* | Guaglianone 1338 (NY) | ERS11697276 | This study | 0.174 | 0.007 | 4752670 | 4103193 (86.3%) | 995 | 958 | 21 |
| *Prosopis cineraria* | Hafisullah and Dilawar 266 (Z) | ERS11697277 | This study | 0.314 | 0.04 | 101697 | 65927 (64.8%) | 697 | 467 | 4 |
| *Prosopis farcta* | Kerimov 31 (NY) | ERS11697278 | This study | 0.243 | 0.006 | 3041167 | 2258136 (74.3%) | 992 | 955 | 33 |
| *Prosopis ferox* | Hughes 2618 (FHO) | ERS11697279 | This study | 0.163 | 0.005 | 5970858 | 4786921 (80.2%) | 993 | 962 | 27 |
| *Prosopis juliflora* | Hughes 1703 (FHO) | ERS11697280 | This study | 0.164 | 0.009 | 4009818 | 3161949 (78.9%) | 992 | 957 | 21 |
| *Prosopis kuntzei* | Hughes 2458 (FHO) | ERS11697281 | This study | 0.165 | 0.006 | 8818267 | 6735181 (76.4%) | 992 | 970 | 48 |
| *Prosopis laevigata* | Hughes 2058 (FHO) | ERS4812932 | Koenen et al. 2020 | 0.02 | 0.019 | 3056948 | 113584 (3.7%) | 772 | 590 | 7 |
| *Prosopis ruscifolia* | Fortunato 6773 (NY) | ERS11697282 | This study | 0.175 | 0.015 | 930099 | 732360 (78.7%) | 992 | 906 | 15 |
| *Prosopis strombulifera* | Kiesling 4828 (NY) | ERS11697283 | This study | 0.147 | 0.006 | 3704512 | 3164148 (85.4%) | 993 | 955 | 18 |
| *Pseudopiptadenia contorta* | de Queiroz 15582 (HUEFS) | ERS4812933 | Koenen et al. 2020 | 0.048 | 0.008 | 6924713 | 330101 (4.8%) | 962 | 821 | 22 |
| *Pseudopiptadenia psilostachya* | Simon 1245 (CEN) | ERS11697284 | This study | 0.147 | 0.006 | 2749436 | 1962195 (71.4%) | 993 | 963 | 38 |
| *Pseudopiptadenia schumanniana* | de Lima 7903 (RB) | ERS11697285 | This study | 0.134 | 0.006 | 2846698 | 2021242 (71%) | 994 | 960 | 45 |
| *Pseudoprosopis euryphylla* | Lock & Frison 88/83 (K) | ERS11697286 | This study | 0.146 | 0.007 | 2291993 | 1765731 (77%) | 987 | 858 | 28 |
| *Pseudoprosopis gilletii* | Wieringa 6021 (WAG) | ERS4812934 | Koenen et al. 2020 | 0.018 | 0.007 | 5604090 | 159884 (2.9%) | 857 | 573 | 11 |
| *Pseudoprosopis sericea* | Jongkind 9513 (WAG) | ERS11697287 | This study | 0.153 | 0.006 | 3850946 | 2800400 (72.7%) | 988 | 904 | 48 |
| *Pseudosamanea cubana* | Leon 12095 (NY) | ERS11697288 | This study | 0.161 | 0.007 | 3885869 | 3042116 (78.3%) | 995 | 977 | 33 |
| *Pseudosamanea guachapele* | Hughes 1198 (FHO) | ERS4812935 | Koenen et al. 2020 | 0.021 | 0.014 | 6673993 | 473494 (7.1%) | 982 | 858 | 12 |
| *Pseudosenegalia feddeana* | Atahuachi 1146 (FHO, LPB) | ERS11697289 | This study | 0.171 | 0.005 | 6105338 | 5021078 (82.2%) | 991 | 944 | 69 |
| *Pterolobium stellatum* | Herendeen and Mbago 17 XII 97 9 (F) | ERS11697290 | This study | 0.242 | 0.007 | 3887117 | 2366370 (60.9%) | 959 | 737 | 28 |
| *Punjuba callejasii* | Daly 5935 (NY) | ERS11697075 | This study | 0.235 | 0.008 | 1185683 | 780532 (65.8%) | 993 | 961 | 20 |

| | | | | | | | | | | |
|---|---|---|---|---|---|---|---|---|---|---|
| *Punjuba killipii* | Palodorios 6252 (NY) | ERS11697082 | This study | 0.227 | 0.006 | 2140489 | 1842980 (86.1%) | 991 | 957 | 23 |
| *Punjuba leh-mannii* | Escobar 7465 (NY) | ERS11697085 | This study | 0.169 | 0.007 | 3964907 | 3155020 (79.6%) | 993 | 972 | 47 |
| *Punjuba racem-iflora* | Jimenez and Soares 3626 (USJ) | ERS11697094 | This study | 0.177 | 0.005 | 5467696 | 4445341 (81.3%) | 994 | 976 | 40 |
| *Recordoxylon speciosum* | Redden 5983 (US) | ERS11697291 | This study | 0.17 | 0.007 | 3672456 | 2416333 (65.8%) | 970 | 806 | 54 |
| *Robrichia olde-manii* | Bonadeu 706 (NY) | ERS11697207 | This study | 0.159 | 0.005 | 6805644 | 5463859 (80.3%) | 993 | 963 | 46 |
| *Robrichia schomburgkii* | Maguire 56534 (K) | ERS11697208 | This study | 0.133 | 0.008 | 2398649 | 2094588 (87.3%) | 992 | 951 | 19 |
| *Samanea sa-man* | Hughes 421 (FHO) | ERS4812936 | Koenen et al. 2020 | 0.015 | 0.008 | 3134366 | 217362 (6.9%) | 942 | 800 | 9 |
| *Sanjappa cy-nometroides* | Krishnaraj 71501 (TBGT) | ERS11697293 | This study | 0.187 | 0.007 | 1909678 | 1523421 (79.8%) | 994 | 930 | 30 |
| *Schizolobium parahyba* | Klitgaard 694 (K) | ERS11697294 | This study | 0.165 | 0.005 | 2957135 | 1938266 (65.5%) | 961 | 710 | 241 |
| *Schleinitzia insularum* | Rinehart 17441 (K) | ERS11697295 | This study | 0.148 | 0.006 | 7684552 | 5971235 (77.7%) | 967 | 779 | 370 |
| *Schleinitzia megaladenia* | Ramos & Edaño 46708 (P) | ERS11697296 | This study | 0.228 | 0.008 | 1463032 | 1247392 (85.3%) | 986 | 746 | 104 |
| *Schleinitzia no-voguineensis* | Chaplin 57 84 (FHO) | ERS4812937 | Koenen et al. 2020 | 0.037 | 0.008 | 15322018 | 1503504 (9.8%) | 979 | 829 | 41 |
| *Senegalia atax-acantha* | Jongkind 10603 (WAG) | ERS4812938 | Koenen et al. 2020 | 0.054 | 0.007 | 10724616 | 680898 (6.3%) | 982 | 857 | 10 |
| *Senegalia bahiensis* | de Queiroz 15499 (HUEFS) | ERS11697298 | This study | 0.158 | 0.006 | 5216922 | 4016660 (77%) | 989 | 960 | 41 |
| *Senegalia born-eensis* | Ambriansyah AA1679 (L) | ERS11697299 | This study | 0.145 | 0.006 | 3048927 | 2292401 (75.2%) | 991 | 961 | 48 |
| *Senegalia ni-grescens* | Barnes 536 (FHO) | ERS11697300 | This study | 0.146 | 0.005 | 8049990 | 6425427 (79.8%) | 994 | 952 | 42 |
| *Senegalia pen-tagona* | Jongkind 10670 (WAG) | ERS11697301 | This study | 0.146 | 0.006 | 3503774 | 2761310 (78.8%) | 990 | 956 | 40 |
| *Senegalia saka-lava* | Koenen 215 (Z) | ERS4812939 | Koenen et al. 2020 | 0.042 | 0.022 | 10377918 | 492909 (4.7%) | 972 | 789 | 9 |
| *Senna cushina* | Hughes 3121 (Z) | ERS11697303 | This study | 0.188 | 0.006 | 4109680 | 2554233 (62.2%) | 969 | 795 | 52 |
| *Senna lasseig-niana* | Hughes 3086 (Z) | ERS11697304 | This study | 0.184 | 0.007 | 3435905 | 2083375 (60.6%) | 962 | 771 | 33 |
| *Senna leandrii* | Koenen 245 (G, K) | ERS11697305 | This study | 0.174 | 0.006 | 7120815 | 4396488 (61.7%) | 967 | 823 | 67 |
| *Senna mollis-sima* | Hughes 3150 (Z) | ERS11697306 | This study | 0.169 | 0.006 | 4492751 | 2659553 (59.2%) | 965 | 786 | 48 |
| *Senna rugosa* | de Queiroz 15592 (HUEFS) | ERS11697307 | This study | 0.186 | 0.005 | 4630639 | 2893029 (62.5%) | 963 | 746 | 29 |
| *Senna velutina* | Wood 26598 (K) | ERS11697308 | This study | 0.21 | 0.006 | 4123567 | 2664458 (64.6%) | 965 | 767 | 43 |

| | | | | | | | | | | |
|---|---|---|---|---|---|---|---|---|---|---|
| *Serianthes calycina* | Barrabé 1158 (NOU) | ERS11697309 | This study | 0.14 | 0.006 | 3888534 | 2912175 (74.9%) | 994 | 973 | 41 |
| *Serianthes nelsonii* | Moore 1241 (L) | ERS4812940 | Koenen et al. 2020 | 0.016 | 0.01 | 6673348 | 332881 (5%) | 968 | 826 | 8 |
| *Sphinga acatlensis* | Hughes 2112 (FHO) | ERS4812941 | Koenen et al. 2020 | 0.02 | 0.011 | 8449227 | 450528 (5.3%) | 976 | 849 | 11 |
| *Stachyothyrsus staudtii* | van Andel 4054 (WAG) | ERS11697310 | This study | 0.2 | 0.006 | 7505359 | 5218629 (69.5%) | 988 | 884 | 82 |
| *Stryphnodendron adstringens* | de Queiroz 15580 (HUEFS) | ERS11697311 | This study | 0.157 | 0.005 | 4934723 | 3893905 (78.9%) | 995 | 971 | 36 |
| *Stryphnodendron duckeanum* | Simon 1606 (CEN) | ERS11697312 | This study | 0.146 | 0.006 | 3948999 | 3097143 (78.4%) | 991 | 966 | 38 |
| *Stryphnodendron paniculatum* | Simon 1058 (CEN) | ERS11697313 | This study | 0.154 | 0.006 | 5781510 | 4469062 (77.3%) | 993 | 972 | 42 |
| *Stryphnodendron pulcherrimum* | de Queiroz 15482 (HUEFS) | ERS4812942 | Koenen et al. 2020 | 0.038 | 0.007 | 10894656 | 789387 (7.2%) | 983 | 887 | 22 |
| *Stuhlmannia moavi* | Keraudren and Aymonin 25628 (MO, P) | ERS11697314 | This study | 0.195 | 0.006 | 2677061 | 1587577 (59.3%) | 971 | 774 | 44 |
| *Sympetalandra schmutzii* | Schmutz 3764 (MO) | ERS11697315 | This study | 0.265 | 0.008 | 4370229 | 3347867 (76.6%) | 948 | 614 | 224 |
| *Sympetalandra unijuga* | Sidiyasa 1320 (K) | ERS11697316 | This study | 0.184 | 0.006 | 4942770 | 3832570 (77.5%) | 935 | 676 | 286 |
| *Tachigali bracteolata* | Mori 24793 (NY) | ERS11697317 | This study | 0.171 | 0.006 | 3385924 | 2220995 (65.6%) | 984 | 851 | 77 |
| *Tachigali guianensis* | Mori 22791 (NY) | ERS11697297 | This study | 0.193 | 0.006 | 4060573 | 2758090 (67.9%) | 985 | 857 | 64 |
| *Tachigali odoratissima* | Morim 562 (RB) | ERS4812943 | Koenen et al. 2020 | 0.012 | 0.015 | 7331095 | 127595 (1.7%) | 777 | 513 | 11 |
| *Tachigali paniculata* | Henkel 657 (NY) | ERS11697318 | This study | 0.166 | 0.006 | 3664973 | 2347363 (64%) | 984 | 845 | 68 |
| *Tachigali vasquezii* | Neill 13998 (MO) | ERS11697319 | This study | 0.162 | 0.006 | 2660201 | 1794909 (67.5%) | 984 | 840 | 47 |
| *Tara spinosa* | Eastwood 36 (FHO) | ERS11697320 | This study | 0.26 | 0.009 | 4787915 | 2560721 (53.5%) | 967 | 761 | 40 |
| *Tetrapleura tetraptera* | Koenen 155 (WAG) | ERS4812944 | Koenen et al. 2020 | 0.017 | 0.01 | 3936371 | 156722 (4%) | 873 | 612 | 13 |
| *Tetrapterocarpon geayi* | Koenen 231 (G, K) | ERS11697321 | This study | 0.227 | 0.006 | 6655137 | 4025160 (60.5%) | 942 | 672 | 63 |
| *Thailentadopsis nitida* | Kostermans 28234 (K) | ERS11697322 | This study | 0.23 | 0.005 | 3128413 | 2452697 (78.4%) | 992 | 964 | 42 |
| *Thailentadopsis tenuis* | Larsen & Larsen 33960 (K) | ERS11697323 | This study | 0.175 | 0.006 | 2236040 | 1901846 (85.1%) | 992 | 955 | 20 |

| | | | | | | | | | | |
|---|---|---|---|---|---|---|---|---|---|---|
| *Umtiza listeriana* | 6th International Legume Conference 10 (JRAU) | ERS11697324 | This study | 0.208 | 0.006 | 5975233 | 3930911 (65.8%) | 967 | 776 | 98 |
| *Vachellia erioloba* | living collection Botanical Garden Zurich | ERS11697325 | This study | 0.153 | 0.006 | 10546334 | 7761717 (73.6%) | 958 | 789 | 368 |
| *Vachellia farnesiana* | living collection Botanical Garden Zurich | ERS11697326 | This study | 0.137 | 0.006 | 6609686 | 4552453 (68.9%) | 962 | 836 | 432 |
| *Vachellia nilotica* | Eastwood 117 (FHO) | ERS11697327 | This study | 0.136 | 0.007 | 7162697 | 5462646 (76.3%) | 932 | 774 | 362 |
| *Vachellia tortilis* | Koenen 603 (Z) | ERS4812945 | Koenen et al. 2020 | 0.025 | 0.02 | 4704801 | 268630 (5.7%) | 944 | 689 | 7 |
| *Vachellia viguieri* | Koenen 199 (Z) | ERS4812946 | Koenen et al. 2020 | 0.037 | 0.013 | 4330329 | 321296 (7.4%) | 938 | 727 | 25 |
| *Viguieranthus glaber* | Koenen 325 (Z) | ERS4812947 | Koenen et al. 2020 | 0.047 | 0.013 | 9401416 | 671920 (7.1%) | 986 | 837 | 13 |
| *Wallaceodendron celebicum* | Flynn 7173 (NY) | ERS11697328 | This study | 0.158 | 0.005 | 6008113 | 4567544 (76%) | 994 | 973 | 33 |
| *Xerocladia viridiramis* | Kolberg and Tholkes HK2493 (WIND) | ERS11697329 | This study | 0.2 | 0.007 | 10389113 | 8119361 (78.2%) | 994 | 959 | 40 |
| *Xylia evansii* | Jongkind 9064 (WAG) | ERS11697330 | This study | 0.176 | 0.005 | 6704436 | 4864344 (72.6%) | 989 | 914 | 56 |
| *Xylia hoffmannii* | Koenen 402 (Z) | ERS4812948 | Koenen et al. 2020 | 0.022 | 0.022 | 6632000 | 276998 (4.2%) | 922 | 678 | 13 |
| *Xylia torreana* | Maurin et al. RBN171 (JRAU) | ERS11697331 | This study | 0.172 | 0.006 | 2829838 | 2042888 (72.2%) | 989 | 899 | 37 |
| *Zapoteca aculeata* | Delinks 332 (NY) | ERS11697332 | This study | 0.228 | 0.007 | 2357164 | 1747778 (74.1%) | 989 | 944 | 35 |
| *Zapoteca amazonica* | Graham 270 (K) | ERS11697333 | This study | 0.291 | 0.009 | 292718 | 59749 (20.4%) | 572 | 283 | 1 |
| *Zapoteca caracasana* | Hughes 3071 (Z) | ERS4812949 | Koenen et al. 2020 | 0.046 | 0.013 | 4703875 | 149537 (3.2%) | 861 | 622 | 6 |
| *Zapoteca nervosa* | Garcia 665 (NY) | ERS11697334 | This study | 0.317 | 0.009 | 3264117 | 2740207 (83.9%) | 992 | 950 | 19 |
| *Zuccagnia punctata* | Fortunato 5545 (MO, BAB) | ERS11697335 | This study | 0.301 | 0.009 | 3141469 | 1138914 (36.3%) | 948 | 696 | 33 |
| *Zygia ampla* | Prance 26527 (NY) | ERS11697336 | This study | 0.159 | 0.007 | 5076411 | 4296146 (84.6%) | 993 | 948 | 34 |
| *Zygia basijuga* | Wurdack 1937 (NY) | ERS11697337 | This study | 0.15 | 0.006 | 3283801 | 2665921 (81.2%) | 992 | 958 | 33 |
| *Zygia bisingula* | Ortega Mendoza 2561 (NY) | ERS11697338 | This study | 0.163 | 0.006 | 7127321 | 5671760 (79.6%) | 994 | 956 | 43 |
| *Zygia cataractae* | Bonadeu 647 (RB) | ERS11697339 | This study | 0.183 | 0.006 | 7213847 | 5544614 (76.9%) | 994 | 950 | 53 |
| *Zygia claviflora* | Iganci 841 (RB) | ERS4812950 | Koenen et al. 2020 | 0.014 | 0.012 | 2246421 | 94113 (4.2%) | 719 | 516 | 2 |

| | | | | | | | | | | |
|---|---|---|---|---|---|---|---|---|---|---|
| *Zygia conzattii* | Ortega 596 (K) | ERS11697340 | This study | 0.136 | 0.008 | 2174933 | 1843911 (84.8%) | 991 | 944 | 26 |
| *Zygia inaequalis* | Iganci 832 (RB) | ERS4812951 | Koenen et al. 2020 | 0.012 | 0.015 | 5158351 | 250135 (4.8%) | 956 | 771 | 7 |
| *Zygia inundata* | Poncy 361 (NY) | ERS11697341 | This study | 0.168 | 0.006 | 6246417 | 5038331 (80.7%) | 994 | 967 | 33 |
| *Zygia latifolia* var. *communis* | Simon 1649 (CEN) | ERS11697342 | This study | 0.197 | 0.005 | 11242303 | 8987934 (79.9%) | 993 | 945 | 50 |
| *Zygia longifolia* | MacQueen 609 (K) | ERS11697343 | This study | 0.172 | 0.007 | 4463055 | 3773313 (84.5%) | 990 | 943 | 29 |
| *Zygia morongii* | Krapovickas 23604 (NY) | ERS11697344 | This study | 0.152 | 0.008 | 3929083 | 3227141 (82.1%) | 993 | 922 | 37 |
| *Zygia obolin-goides* | Krukoff 10969 (NY) | ERS11697345 | This study | 0.191 | 0.007 | 4781761 | 3877913 (81.1%) | 992 | 950 | 30 |
| *Zygia ocuma-rensis* | Stergios 14780 (NY) | ERS11697346 | This study | 0.158 | 0.006 | 4060642 | 3247793 (80%) | 991 | 961 | 31 |
| *Zygia racemosa* | Simon 1658 (CEN) | ERS4812952 | Koenen et al. 2020 | 0.02 | 0.007 | 10085857 | 397461 (3.9%) | 976 | 830 | 10 |
| *Zygia ramiflora* | de Lima 2751 (NY) | ERS11697347 | This study | 0.388 | 0.011 | 257984 | 173996 (67.4%) | 964 | 847 | 8 |
| *Zygia rhytido-carpa* | Yuncker 8553 (NY) | ERS11697348 | This study | 0.205 | 0.005 | 7756608 | 6172803 (79.6%) | 993 | 937 | 39 |
| *Zygia sabatieri* | Sabatier 4838 (K) | ERS11697349 | This study | 0.204 | 0.007 | 5239416 | 4461118 (85.1%) | 992 | 959 | 31 |
| *Zygia sp.* | Coley and Kursar Tip917 (UT) | ERR776824 | Koenen et al. 2020 | 0.07 | 0.008 | 1229023 | 508986 (41.4%) | 960 | 802 | 36 |
| *Zygia unifolio-lata* | Wendt 4388 (K) | ERS11697350 | This study | 0.163 | 0.005 | 5591357 | 4408016 (78.8%) | 993 | 950 | 50 |

**Table S2.** Outgroup species with numbers of genes matching the target genes used here which were recovered by BLAST and BLAT from genome sequences.

| Taxon | Reference | BLAST | BLAT | Total |
|---|---|---|---|---|
| *Arachis ipaensis* | (*139*) | 451 | 195 | 646 |
| *Cercis canadensis* | (*140*) | 694 | 167 | 861 |
| *Glycine max* | (*141*) | 623 | 135 | 758 |
| *Medicaga trunculata* | (*142*) | 488 | 158 | 646 |
| *Nissolia schotii* | (*143*) | 527 | 191 | 718 |

| | |
|---|---|
| Total unique genes | 918 |

**Table S3.** Identities, ages, and placements of fossil constraints used for time-calibrating the Hybseq backbone phylogeny. SG = Stem Group; MRCA = Most Recent Common Ancestor

| Node | Fossil taxon | Min age (Mya) | Reference |
|---|---|---|---|
| Caesalpinioideae SG | Unnamed bipinnate leaves | 58 | (*144, 145*) |
| *Arcoa* SG | *Arcoa* | 48.5 | (*146, 147*) |
| *Senna*/*Cassia* split | *Senna* fruits | 37.8 | (*148*) |
| *Dinizia*/Mimosoid clade MRCA | *Eumimosoidea plumosa* flowers, leaves and fruits | 37.8 | (*149*) |
| *Vachellia*/ingoid clade split | Flattened 16-celled polyads similar to *Vachellia, Albizia* and other ingoids | 33.9 | (*150*) |
| *Acacia* SG | *Acacia* polyads | 23 | (*151*) |
| *Calliandra* SG | *Calliandra* polyads | 16 | (*152*) |

**Table S4.** GenBank accession numbers of new *Acacia* samples included in the metachronogram.

| Taxon name | GenBank accession code |
|---|---|
| Acacia_acellerata_2617 | XXX |
| Acacia_aculeiformis_2620 | XXX |
| Acacia_alleniana_1786 | XXX |
| Acacia_ammophila_2934 | XXX |
| Acacia_amoena_3553 | XXX |
| Acacia_anarthros_2658 | XXX |
| Acacia_aptaneura_9275 | XXX |
| Acacia_awestoniana_2797 | XXX |
| Acacia_barbinervis_2839 | XXX |
| Acacia_barrancana_3530 | XXX |
| Acacia_barrettiorum_3518 | XXX |
| Acacia_bartleana_2837 | XXX |
| Acacia_blaxellii_2848 | XXX |
| Acacia_botrydion_3154 | XXX |
| Acacia_brachycarpa_2843 | XXX |
| Acacia_brumalis_3557 | XXX |
| Acacia_brunioides_2868 | XXX |
| Acacia_burbidgeae_2888 | XXX |
| Acacia_bynoeana_2887 | XXX |
| Acacia_caerulescens_2886 | XXX |
| Acacia_caesaneura_9367 | XXX |
| Acacia_caesariata_2885 | XXX |
| Acacia_calcarata_2883 | XXX |
| Acacia_caleyi_2882 | XXX |
| Acacia_calyculata_3866 | XXX |
| Acacia_cambagei_3468 | XXX |
| Acacia_carens_2876 | XXX |
| Acacia_caroliron42ca | XXX |
| Acacia_cataractae_2898 | XXX |
| Acacia_centrinervia_2893 | XXX |
| Acacia_cerastes_2892 | XXX |
| Acacia_chalkeri_2891 | XXX |
| Acacia_chamaeleon_2890 | XXX |
| Acacia_chapmanii_3489 | XXX |
| Acacia_chippendalei_3488 | XXX |
| Acacia_chrysocephala_827 | XXX |
| Acacia_chrysotrina_2188 | XXX |
| Acacia_cincinnata_1778 | XXX |
| Acacia_cochlocarpa_3297 | XXX |
| Acacia_convallium_3302 | XXX |
| Acacia_convenyi_2811 | XXX |
| Acacia_costiniana_3609 | XXX |
| Acacia_crassistipula_3304 | XXX |
| Acacia_crassuloides_3308 | XXX |
| Acacia_cretata_1770 | XXX |
| Acacia_crispula_3311 | XXX |
| Acacia_cummingiana_3128 | XXX |
| Acacia_dacrydioides_3538 | XXX |
| Acacia_daviesioides_3650 | XXX |
| Acacia_delicatula_3315 | XXX |
| Acacia_deltoidea_3519 | XXX |
| Acacia_dermatophylla_3255 | XXX |
| Acacia_diaphyllodinea_3252 | XXX |
| Acacia_dilatata_3248 | XXX |
| Acacia_diminuta_3247 | XXX |
| Acacia_disticha_3246 | XXX |
| Acacia_drewiana_3243 | XXX |
| Acacia_durabilis_3240 | XXX |
| Acacia_echinuliflora_3232 | XXX |
| Acacia_effusifolia_3416 | XXX |
| Acacia_ericksoniae_3230 | XXX |
| Acacia_erioclada_3220 | XXX |
| Acacia_fasciculifera_305 | XXX |
| Acacia_filamentosa_3419 | XXX |
| Acacia_flavescens_3612 | XXX |
| Acacia_flavipila_3832 | XXX |
| Acacia_floydii_3773 | XXX |
| Acacia_frigescens_3640 | XXX |
| Acacia_froggattii_3515 | XXX |
| Acacia_fubescens_2251 | XXX |
| Acacia_fuscaneura_9778 | XXX |
| Acacia_gemina_3560 | XXX |
| Acacia_glaucocaesia_2859 | XXX |
| Acacia_gordonii_2138 | XXX |
| Acacia_gracilenta_3420 | XXX |
| Acacia_helicophylla_3601 | XXX |
| Acacia_hemignosta_3520 | XXX |
| Acacia_humifusa_3867 | XXX |
| Acacia_hyaloneura_3744 | XXX |
| Acacia_imitans_3168 | XXX |
| Acacia_incognita_3425 | XXX |
| Acacia_incurva_873 | XXX |
| Acacia_incurvaneura_9646 | XXX |
| Acacia_ingrata_3165 | XXX |
| Acacia_inophloia_3588 | XXX |
| Acacia_intricata_3571 | XXX |
| Acacia_jancada_2304 | XXX |
| Acacia_jasperensis_3428 | XXX |
| Acacia_kelleri_3536 | XXX |
| Acacia_kenneallyi_3517 | XXX |
| Acacia_kettlewelliae_3434 | XXX |

| | |
|---|---|
| Acacia_kimberleyensis_3537 | XXX |
| Acacia_kochii_3293 | XXX |
| Acacia_lacertensis_3291 | XXX |
| Acacia_lachnophylla_3286 | XXX |
| Acacia_larigerava_2298 | XXX |
| Acacia_lasiocarpa_3655 | XXX |
| Acacia_lateriticola_157 | XXX |
| Acacia_latifolia_3437 | XXX |
| Acacia_latior_3283 | XXX |
| Acacia_leeuweniana_3281 | XXX |
| Acacia_leichhardtii_1986 | XXX |
| Acacia_leiophylla_3652 | XXX |
| Acacia_lentiginea_3525 | XXX |
| Acacia_leprosa_3822 | XXX |
| Acacia_leptoclada_DM1 | XXX |
| Acacia_leptostachya_3586 | XXX |
| Acacia_limbata_3507 | XXX |
| Acacia_linarioides_3439 | XXX |
| Acacia_linophylla_1866 | XXX |
| Acacia_littorea_3564 | XXX |
| Acacia_lucasii_3277 | XXX |
| Acacia_lullfitziorum_3441 | XXX |
| Acacia_luteola_2100 | XXX |
| Acacia_macraneura_9178 | XXX |
| Acacia_maidenii_3004 | XXX |
| Acacia_marangensis_2152 | XXX |
| Acacia_matthewii_2121 | XXX |
| Acacia_megalantha_3585 | XXX |
| Acacia_merrallii_3607 | XXX |
| Acacia_merrickiae_448 | XXX |
| Acacia_microcalyx_3325 | XXX |
| Acacia_microneura_3275 | XXX |
| Acacia_mimica_3273 | XXX |
| Acacia_moirii_2101 | XXX |
| Acacia_mooreana_3270 | XXX |
| Acacia_mulganeura_9276 | XXX |
| Acacia_multistipulosa_3267 | XXX |
| Acacia_neriifolia_2033 | XXX |
| Acacia_nitidula_3265 | XXX |
| Acacia_nodiflora_3583 | XXX |
| Acacia_octonervia_3264 | XXX |
| Acacia_ophiolithica_3262 | XXX |
| Acacia_oraria_3722 | XXX |
| Acacia_orbifolia_3261 | XXX |
| Acacia_papulosa_3323 | XXX |
| Acacia_peregrinalis_2366 | XXX |
| Acacia_phaeocalyx_3322 | XXX |
| Acacia_pharangites_3320 | XXX |
| Acacia_piligera_2120 | XXX |
| Acacia_pinguiculosa_3318 | XXX |
| Acacia_plicata_3219 | XXX |
| Acacia_polystachya_3327 | XXX |
| Acacia_pteraneura_3066 | XXX |
| Acacia_ptychoclada_3645 | XXX |
| Acacia_pubescens_2275 | XXX |
| Acacia_pubirhachis_3331 | XXX |
| Acacia_quadrimarginea_3570 | XXX |
| Acacia_quadrisulcata_3562 | XXX |
| Acacia_quinquinervia_3334 | XXX |
| Acacia_resinicostata_2147 | XXX |
| Acacia_resinimarginea_2924 | XXX |
| Acacia_resinosa_3335 | XXX |
| Acacia_restiacea_3582 | XXX |
| Acacia_riceana_3794 | XXX |
| Acacia_rigescens_3209 | XXX |
| Acacia_rossei_209 | XXX |
| Acacia_rothii_3604 | XXX |
| Acacia_rotundifolia_3600 | XXX |
| Acacia_sclerosperma_3697 | XXX |
| Acacia_semirigida_3608 | XXX |
| Acacia_sericata_3339 | XXX |
| Acacia_sericocarpa_3344 | XXX |
| Acacia_sessilis_3599 | XXX |
| Acacia_setulifera_3804 | XXX |
| Acacia_shirleyi_3748 | XXX |
| Acacia_signata_3556 | XXX |
| Acacia_sp_3118 | XXX |
| Acacia_sp_3119 | XXX |
| Acacia_sp_3636 | XXX |
| Acacia_sp_3721 | XXX |
| Acacia_sp_3730 | XXX |
| Acacia_sp_3757 | XXX |
| Acacia_sp_3768 | XXX |
| Acacia_sp_3775 | XXX |
| Acacia_sp_3864 | XXX |
| Acacia_speckii_3346 | XXX |
| Acacia_spinosissima_3349 | XXX |
| Acacia_spondylophylla_934 | XXX |
| Acacia_startii_3702 | XXX |
| Acacia_steedmanii_3563 | XXX |
| Acacia_stellaticeps_3784 | XXX |
| Acacia_stereophylla_3704 | XXX |
| Acacia_subflexuosa_3597 | XXX |
| Acacia_sublanata_3353 | XXX |

| | |
|---|---|
| Acacia_subporosa_3783 | XXX |
| Acacia_subracemosa_3567 | XXX |
| Acacia_subternata_3806 | XXX |
| Acacia_sulcata_3598 | XXX |
| Acacia_teretifolia_3594 | XXX |
| Acacia_trigonophylla_3593 | XXX |
| Acacia_trinalis_3226 | XXX |
| Acacia_trinervata_3568 | XXX |
| Acacia_trineura_3590 | XXX |
| Acacia_triptycha_3581 | XXX |
| Acacia_ulicifolia_3579 | XXX |
| Acacia_ulicina_3580 | XXX |
| Acacia_uliginosa_3224 | XXX |
| Acacia_uncifera_3732 | XXX |
| Acacia_uncinella_3591 | XXX |
| Acacia_unguicula_3223 | XXX |
| Acacia_urandangi_3587 | XXX |
| Acacia_urophylla_3576 | XXX |
| Acacia_veronica_3780 | XXX |
| Acacia_verticillata_2950 | XXX |
| Acacia_vincentii_3522 | XXX |
| Acacia_vittata_3221 | XXX |
| Acacia_wardellii_3355 | XXX |
| Acacia_wickhamii_3578 | XXX |
| Acacia_williamsonii_3577 | XXX |

**Table S5.** New *Albizia* samples in the metachronogram, including European Nucleotide Archive accession numbers.

| Species name | Voucher | Herbarium | ENA accession number |
|---|---|---|---|
| *Albizia petersiana* | Greenway & Kanuri 15362 | PRE | XXX |
| *Albizia evansii* | Correia & Marques 843 | PRE | XXX |
| *Albizia gummifera* | Tawakali & Kaunda 436 | PRE | XXX |
| *Albizia mainaea* | Du Puy M682 | K | XXX |
| *Albizia laurentii* | Breteler 15764 | WAG | XXX |
| *Albizia gillardinii* | Deschamps 256 | K | XXX |
| *Albizia julibrissin* | Fumihiro Konta 18313 | P | XXX |
| *Albizia kalkora* | Xiao Bai-Zhong | K | XXX |
| *Albizia chinensis* | But 79-100A | K | XXX |
| *Albizia philippinensis* | Ahern's collector 215 | P | XXX |
| *Albizia sherriffii* | Grierson 4314 | K | XXX |
| *Albizia salomonensis* | Whitmore BSIP 4424 | K | XXX |
| *Albizia guillainii* | Veillon 8151 | NOU | XXX |
| *Albizia lebbekoides* | Larsen 31706 | K | XXX |
| *Albizia vialeana* | Larsen 32204 | P | XXX |
| *Albizia odoratissima* | Chow 80078 | K | XXX |
| *Albizia corniculata* | Shiu Ying Hu 10383 | K | XXX |
| *Albizia rufa* | Zollinger 80 | P | XXX |
| *Albizia myriophylla* | Larsen 33455 | P | XXX |
| *Albizia tomentella* | Leach & Dunlop 3801 | L | XXX |
| *Albizia crassiramea* | Parnell 95-401 | K | XXX |
| *Albizia duclouxii* | Ducloux 6112 | P | XXX |
| *Albizia lebbeck* | Nee 28682 | NY | XXX |
| *Albizia procera* | Matthew RHT 63670 | K | XXX |
| *Albizia attopeuensis* | Maxwell 92-747 | P | XXX |
| *Albizia canescens* | Forster PIF9580 | L | XXX |
| *Albizia pedicellata* | Ambriansyah AA702 | K | XXX |
| *Albizia rosulata* | Ariffin Kalat BRUN 17375 | K | XXX |
| *Albizia acle* | Villavicencio 27204 | P | XXX |
| *Albizia chevalieri* | Okafov & Daramota 54622 | FHO | XXX |
| *Albizia harveyi* | Maurin 0773 | JRAU | XXX |
| *Albizia suluensis* | Maurin 1810 | JRAU | XXX |
| *Albizia malacophylla* | de Wilde & de Wilde-Duyfjes 8944 | PRE | XXX |
| *Albizia antunesiana* | Tinley 1454 | PRE | XXX |
| *Albizia tanganyicensis* | Maurin 1972 | JRAU | XXX |
| *Albizia coriaria* | Jongkind 2482 | WAG | XXX |
| *Albizia odorata* | Du Puy M335 | K | XXX |
| *Albizia tulearensis* | Service Forestier Madagascar & Capuron 29107 sf | P | XXX |
| *Albizia vaughanii* | Vaughan 10763 | K | XXX |
| *Albizia divaricata* | Randrianaivo 485 | P | XXX |
| *Albizia forbesii* | Maurin 0228 | JRAU | XXX |
| *Albizia morombensis* | Phillipson & Manjakahery 5985 | P | XXX |
| *Albizia numidarum* | Labat 3552 | K | XXX |
| *Albizia arenicola* | Du Puy M296 | K | XXX |

| | | | |
|---|---|---|---|
| *Albizia boinensis* | Du Puy M523 | WAG | XXX |
| *Albizia verrucosa* | Service Forestier Madagascar & Capuron 24093 sf | P | XXX |
| *Albizia perrieri* | Nusbaumer LN 1876 | K | XXX |
| *Albizia androyensis* | Randrianaivo et al. 1742 | P | XXX |
| *Albizia commiphoroides* | Phillipson 2781 | P | XXX |
| *Albizia balabaka* | Du Puy M123 | K | XXX |
| *Albizia jaubertiana* | Service Forestier Madagascar & Capuron 24619 sf | P | XXX |
| *Albizia* sp. | Koenen 387 | G | XXX |
| *Albizia isenbergiana* | Verdcourt 3237B | K | XXX |
| *Albizia schimperiana* | Van Wyk BSA 2748 | PRE | XXX |
| *Albizia glaberrima* | Maurin 2605 | JRAU | XXX |
| *Albizia amara* subsp. *sericoceph-ala* | Maurin 3368 | JRAU | XXX |
| *Albizia amara* subsp. *amara* | Semsei 3274 | PRE | XXX |

**Table S6.** GenBank accession numbers of *Leucaena* samples included in the metachronogram.

| **Sample** | **GenBank accession number** |
|---|---|
| Leucaena_alvadorensis | XXX |
| Leucaena_lempirana | XXX |
| Leucaena_multicapitula | XXX |
| Leucaena_zacapana | XXX |
| Leucaena_collinsii | XXX |
| Leucaena_shannonii | XXX |
| Leucaena_trichandra | XXX |
| Leucaena_magnifica | XXX |
| Leucaena_confertiflora_adenotheloidia | XXX |
| Leucaena_lanceolata_lanceolata | XXX |
| Leucaena_cruziana | XXX |
| Leucaena_trichodes | XXX |
| Leucaena_macrophylla_macrophylla | XXX |
| Leucaena_pueblana | XXX |
| Leucaena_esculenta | XXX |
| Leucaena_involucrata | XXX |
| Leucaena_pallida | XXX |
| Leucaena_matudae | XXX |
| Leucaena_leucocephala_leucocephala | XXX |
| Leucaena_pulverulenta | XXX |
| Leucaena_diversifolia | XXX |
| Leucaena_greggii | XXX |
| Leucaena_retusa | XXX |
| Leucaena_cuspidata | XXX |

**Table S7.** GenBank accession numbers of new *Mimosa* samples included in the metachronogram.

| **Name in metachronogram** | **Species name** | **Voucher** | **GenBank accession number** |
|---|---|---|---|
| Mimosa_aff_custodis_QUIR6 | *Mimosa* sp. | QUIR 6 | ON530940 |
| Mimosa_aff_pigra_MFS3066 | *Mimosa pigra* | M.F. Simon 3066 | ON530935 |
| Mimosa_afranioi_ACS6669 | *Mimosa afranioi* | A.C. Sevilha 6669 (MG) | OM891221 |
| Mimosa_burchellii_MFS2826 | *Mimosa burchellii* | M.F. Simon 2826 (MA) | OM891222 |
| Mimosa_burkartii_BK1 | *Mimosa burkartii* | BK1 | ON530912 |
| Mimosa_callydryas_MFS3568 | *Mimosa callidryas* | M.F. Simon 3568 | ON530936 |
| Mimosa_campicola_LSBJ354 | *Mimosa campicola* | L.S.B. Jordão 354 (BA) | OM891223 |
| Mimosa_crumenarioides_LSBJ215 | *Mimosa crumenarioides* | L.S.B. Jordão 215 (BA) | OM891224 |
| Mimosa_demissa_MM5128 | *Mimosa demissa* | MM 5128 | ON530932 |
| Mimosa_distans_MFS2462 | *Mimosa distans* | M.F. Simon 2462 | ON530924 |
| Mimosa_diversipila_MM2026_DE | *Mimosa diversipila* | MM2026 DE | ON530911 |
| Mimosa_dolens_foliolosa_DE13 | *Mimosa dolens* var. *foliolosa* | DE13 | ON530920 |
| Mimosa_dolens_rudis_BOTU3 | *Mimosa dolens* var. *rudis* | BOTU3 | ON530919 |
| Mimosa_dryandroides_LJ427 | *Mimosa dryandroides* | LJ427 | ON530922 |
| Mimosa_emaensis_LSBJ424 | *Mimosa emaensis* | L.S.B. Jordão 424 (GO) | OM891226 |
| Mimosa_falcipinna_MM4980 | *Mimosa falcipinna* | MM4980 | ON530925 |
| Mimosa_filiformis_LSBJ303 | *Mimosa gracilis* subsp. *filiformis* | L.S.B. Jordão 303 (MG) | OM891227 |
| Mimosa_flabellifolia_LSBJ439 | *Mimosa flabellifolia* | L.S.B. Jordão 439 (TO) | OM891228 |
| Mimosa_flavocaesia_MFS3193 | *Mimosa flavocaesia* | M.F. Simon 3193 | ON530933 |
| Mimosa_gracilis_brevissima_MFS2606 | *Mimosa gracilis* subsp. *brevissi-ma* | M.F. Simon 2606 (GO) | OM891230 |
| Mimosa_gracilis_gracilis_LSBJ300 | *Mimosa gracilis* subsp. *gracilis* | L.S.B. Jordão 300 (MG) | OM891231 |
| Mimosa_hatschbachii_EL4109 | *Mimosa hatschbachii* | EL4190 | ON530915 |
| Mimosa_intricata_INT8 | *Mimosa intricata* | INT8 | ON530914 |
| Mimosa_longipes_MFS2467 | *Mimosa longipes* | M.F. Simon 2467 | ON530930 |
| Mimosa_macrocephala_MFS2475 | *Mimosa macrocephala* | M.F. Simon 2475 | ON530926 |
| Mimosa_maracayuensis_LSBJ410 | *Mimosa maracayuensis* | L.S.B. Jordão 410 (MS) | OM891232 |
| Mimosa_micropteris_MM2124 | *Mimosa micropteris* | MM 2124 | ON530938 |
| Mimosa_multiceps_LSBJ419 | *Mimosa multiceps* | L.S.B. Jordão 419 (MS) | OM891233 |
| Mimosa_osmarii_LSBJ279 | *Mimosa osmarii* | L.S.B. Jordão 279 (MG) | OM891234 |
| Mimosa_pabstiana_ACS6809 | *Mimosa pabstiana* | A.C. Sevilha 6809 (MG) | MT044447 |
| Mimosa_palmetorum_DMN1581 | *Mimosa palmetorum* | D.M. Neves 1581 (BA) | OM891235 |
| Mimosa_paraibana_MFS2813 | *Mimosa paraibana* | M.F. Simon 2813 (MA) | MT044446 |
| Mimosa_phyllodinea_LSBJ394 | *Mimosa phyllodinea* | L.S.B. Jordão 394 (GO) | OM891237 |
| Mimosa_pinetorum_LSBJ243 | *Mimosa pinetorum* | L.S.B. Jordão 243 (BA) | OM891238 |
| Mimosa_rava_MFS2658 | *Mimosa rava* | M.F. Simon 2658 | ON530928 |
| Mimosa_reduviosa_TBC3846 | *Mimosa reduviosa* | TBC 3846 | ON530941 |
| Mimosa_regnellii_MM2118 | *Mimosa regnellii* | MM 2118 | ON530937 |
| Mimosa_regnellii_supersetosa_ MM2130 | *Mimosa regnellii* var. *supersetosa* | MM 2130 | ON530939 |
| Mimosa_selloi_LSBJ430 | *Mimosa selloi* | L.S.B. Jordão 430 (SP) | OM891239 |
| Mimosa_serpens_EDL2678 | *Mimosa serpens* | E.D. Lozano 2678 (PR) | OM891240 |
| Mimosa_sobralii_A17 | *Mimosa sobralii* | A17 | ON530913 |

| | | | |
|---|---|---|---|
| Mimosa_sp_nov_pachyc_MFS3060 | *Mimosa* sp. | M.F. Simon 3060 | ON530927 |
| Mimosa_sp_nov_polycepha_MFS3061 | *Mimosa pseudoracemosa* | M.F. Simon 3061 | ON530934 |
| Mimosa_sp_nov_TBC3839 | *Mimosa* sp. | TBC 3839 | ON530918 |
| Mimosa_sp_stipel_PSE4 | *Mimosa* sp. | PSE 4 | ON530917 |
| Mimosa_subenervis_LSBJ336 | *Mimosa subenervis* | L.S.B. Jordão 336 (BA) | OM891241 |
| Mimosa_suburbana_LSBJ393 | *Mimosa suburbana* | L.S.B. Jordão 393 (DF) | OM891242 |
| Mimosa_supravisa_LSBJ442 | *Mimosa supravisa* | L.S.B. Jordão 442 (GO) | OM891243 |
| Mimosa_thomista_MB2768 | *Mimosa thomista* | MB 2768 | ON530923 |

**Table S8.** GenBank accession numbers of Parkia clade samples included in the metachronogram.

| **Species** | **GenBank accession code** |
|---|---|
| *Anadenanthera peregrina* | AF521814 |
| *Parkia bahiae* | KY046204 |
| *Parkia biglobosa* | KY046215 |
| *Parkia decussata* | KY118189 |
| *Parkia discolor* | KY045957 |
| *Parkia leiophylla* | KY046207 |
| *Parkia madagascariensis* | KY046208 |
| *Parkia multijuga* | KY046209 |
| *Parkia nitida* | KY046210 |
| *Parkia panurensis* | KY046211 |
| *Parkia pendula* | KY046083 |
| *Parkia platycephala* | KY045958 |
| *Parkia sumatrana* | KY046214 |
| *Parkia ulei* | KY046045 |
| *Parkia velutina* | KY046084 |
| *Vachellia bidwillii* | MK923636 |
| *Vachellia clarksoniana* | MK923634 |
| *Vachellia ditricha* | MK923632 |
| *Vachellia douglasica* | MK923631 |
| *Vachellia pachyphloia* | MK923630 |
| *Vachellia suberosa* | MK923629 |
| *Vachellia sutherlandii* | MK923627 |
| *Vachellia valida* | MK923623 |

**Table S9.** Ancient phylogenetic turnover variation partitioning results obtained with the metachronogram per region. The relative contribution of each predictor variable is shown, as well as the total deviance explained by the full model.

**pantropics**

| **Cut-off** | **Full** | **5 Ma** | **10 Ma** | **20 Ma** |
|---|---|---|---|---|
| **Correlation with taxonomic turnover** | 0.70 | 0.62 | 0.56 | 0.35 |
| **Correlation with full phylogenetic turnover** | 1.00 | 0.99 | 0.95 | 0.64 |
| **Variation partitioning** | | | | |
| **Climatic distance** | 0.04 | 0.03 | 0.03 | 0.00 |
| **Interaction** | 0.31 | 0.29 | 0.26 | 0.00 |
| **Spatial distance** | 0.64 | 0.67 | 0.71 | 0.00 |
| **Percentage deviance explained** | 54.47 | 51.68 | 47.01 | 0.00 |

**North America**

| **Cut-off** | **Full** | **5 Ma** | **10 Ma** | **20 Ma** |
|---|---|---|---|---|
| **Correlation with taxonomic turnover** | 0.70 | 0.61 | 0.52 | 0.41 |
| **Correlation with full phylogenetic turnover** | 1.00 | 0.99 | 0.96 | 0.87 |
| **Variation partitioning** | | | | |
| **Climatic distance** | 0.48 | 0.50 | 0.56 | 0.61 |
| **Interaction** | 0.51 | 0.49 | 0.44 | 0.39 |
| **Spatial distance** | 0.01 | 0.00 | 0.00 | 0.00 |
| **Percentage deviance explained** | 29.92 | 25.78 | 22.18 | 16.74 |

**South America**

| **Cut-off** | **Full** | **5 Ma** | **10 Ma** | **20 Ma** |
|---|---|---|---|---|
| **Correlation with taxonomic turnover** | 0.64 | 0.53 | 0.48 | 0.36 |
| **Correlation with full phylogenetic turnover** | 1.00 | 0.98 | 0.96 | 0.82 |
| **Variation partitioning** | | | | |
| **Climatic distance** | 0.55 | 0.62 | 0.63 | 0.64 |
| **Interaction** | 0.40 | 0.34 | 0.33 | 0.30 |
| **Spatial distance** | 0.05 | 0.04 | 0.04 | 0.06 |
| **Percentage deviance explained** | 21.84 | 18.79 | 17.11 | 12.05 |

**Africa**

| **Cut-off** | **Full** | **5 Ma** | **10 Ma** | **20 Ma** |
|---|---|---|---|---|
| **Correlation with taxonomic turnover** | 0.78 | 0.70 | 0.63 | 0.53 |
| **Correlation with full phylogenetic turnover** | 1.00 | 0.99 | 0.97 | 0.90 |
| **Variation partitioning** | | | | |
| **Climatic distance** | 0.63 | 0.66 | 0.69 | 0.71 |
| **Interaction** | 0.30 | 0.28 | 0.26 | 0.25 |
| **Spatial distance** | 0.06 | 0.05 | 0.04 | 0.03 |
| **Percentage deviance explained** | 28.10 | 27.41 | 26.62 | 23.61 |

**Asia**

| **Cut-off** | **Full** | **5 Ma** | **10 Ma** | **20 Ma** |
|---|---|---|---|---|
| **Correlation with taxonomic turnover** | 0.86 | 0.77 | 0.65 | 0.29 |
| **Correlation with full phylogenetic turnover** | 1.00 | 0.99 | 0.94 | 0.59 |
| **Variation partitioning** | | | | |
| **Climatic distance** | 0.20 | 0.26 | 0.34 | 1.00 |
| **Interaction** | 0.39 | 0.35 | 0.29 | 0.00 |
| **Spatial distance** | 0.41 | 0.39 | 0.37 | 0.00 |
| **Percentage deviance explained** | 34.58 | 31.87 | 27.19 | 15.79 |

| Australia | Cut-off | Full | 5 Ma | 10 Ma | 20 Ma |
|---|---|---|---|---|---|
| | Correlation with taxonomic turnover | 0.77 | 0.64 | 0.47 | 0.12 |
| | Correlation with full phylogenetic turnover | 1.00 | 0.96 | 0.82 | 0.26 |
| | Variation partitioning | | | | |
| | Climatic distance | 0.47 | 0.51 | 0.59 | 0.00 |
| | Interaction | 0.47 | 0.44 | 0.38 | 0.00 |
| | Spatial distance | 0.07 | 0.05 | 0.03 | 0.00 |
| | Percentage deviance explained | 44.67 | 37.96 | 25.13 | 0.00 |

**Table S10.** Ancient phylogenetic turnover variation partitioning results obtained with the genus-level Mimosoid tree per region. The relative contribution of each predictor variable is shown, as well as the total deviance explained by the full model.

| pantropics | Cut-off | Full | 5 Ma | 10 Ma | 20 Ma |
|---|---|---|---|---|---|
| | Correlation with taxonomic turnover | 0.97 | 0.96 | 0.92 | 0.54 |
| | Correlation with full phylogenetic turnover | 1.00 | 1.00 | 0.99 | 0.69 |
| | Variation partitioning | | | | |
| | Climatic distance | 0.41 | 0.41 | 0.41 | 0.54 |
| | Interaction | 0.50 | 0.50 | 0.50 | 0.39 |
| | Spatial distance | 0.08 | 0.08 | 0.09 | 0.07 |
| | Percentage deviance explained | 51.98 | 49.87 | 46.22 | 29.58 |

| North America | Cut-off | Full | 5 Ma | 10 Ma | 20 Ma |
|---|---|---|---|---|---|
| | Correlation with taxonomic turnover | 0.88 | 0.82 | 0.73 | 0.54 |
| | Correlation with full phylogenetic turnover | 1.00 | 0.99 | 0.96 | 0.82 |
| | Variation partitioning | | | | |
| | Climatic distance | 0.43 | 0.46 | 0.50 | 0.55 |
| | Interaction | 0.53 | 0.51 | 0.49 | 0.45 |
| | Spatial distance | 0.04 | 0.03 | 0.02 | 0.00 |
| | Percentage deviance explained | 36.74 | 33.57 | 28.74 | 22.62 |

| South America | Cut-off | Full | 5 Ma | 10 Ma | 20 Ma |
|---|---|---|---|---|---|
| | Correlation with taxonomic turnover | 0.86 | 0.78 | 0.60 | 0.44 |
| | Correlation with full phylogenetic turnover | 1.00 | 0.99 | 0.88 | 0.75 |
| | Variation partitioning | | | | |
| | Climatic distance | 0.41 | 0.48 | 0.65 | 0.71 |
| | Interaction | 0.49 | 0.44 | 0.30 | 0.24 |
| | Spatial distance | 0.09 | 0.08 | 0.04 | 0.05 |
| | Percentage deviance explained | 31.86 | 29.72 | 25.18 | 18.14 |

| Africa | Cut-off | Full | 5 Ma | 10 Ma | 20 Ma |
|---|---|---|---|---|---|
| | Correlation with taxonomic turnover | 0.91 | 0.87 | 0.81 | 0.57 |
| | Correlation with full phylogenetic turnover | 1.00 | 1.00 | 0.98 | 0.80 |
| | Variation partitioning | | | | |
| | Climatic distance | 0.42 | 0.45 | 0.50 | 0.67 |
| | Interaction | 0.46 | 0.44 | 0.41 | 0.29 |
| | Spatial distance | 0.12 | 0.11 | 0.09 | 0.04 |
| | Percentage deviance explained | 41.65 | 41.54 | 41.34 | 36.03 |

| Asia | | | | |
|---|---|---|---|---|
| **Cut-off** | **Full** | **5 Ma** | **10 Ma** | **20 Ma** |
| **Correlation with taxonomic turnover** | 0.88 | 0.83 | 0.78 | 0.39 |
| **Correlation with full phylogenetic turnover** | 1.00 | 1.00 | 0.98 | 0.62 |
| **Variation partitioning** | | | | |
| **Climatic distance** | 0.09 | 0.09 | 0.10 | 0.46 |
| **Interaction** | 0.45 | 0.45 | 0.45 | 0.33 |
| **Spatial distance** | 0.47 | 0.46 | 0.45 | 0.21 |
| **Percentage deviance explained** | 38.63 | 35.64 | 32.16 | 10.93 |

| Australia | | | | |
|---|---|---|---|---|
| **Cut-off** | **Full** | **5 Ma** | **10 Ma** | **20 Ma** |
| **Correlation with taxonomic turnover** | 0.97 | 0.95 | 0.92 | 0.52 |
| **Correlation with full phylogenetic turnover** | 1.00 | 0.95 | 0.99 | 0.69 |
| **Variation partitioning** | | | | |
| **Climatic distance** | 0.34 | 0.33 | 0.33 | 0.41 |
| **Interaction** | 0.54 | 0.54 | 0.54 | 0.45 |
| **Spatial distance** | 0.12 | 0.12 | 0.13 | 0.13 |
| **Percentage deviance explained** | 53.17 | 50.85 | 46.89 | 21.73 |

**Table S11.** Details of the concatenated alignments showing alignment lengths, phylogenetic informativeness, and fractions of missing data.

| **Alignment** | **Type** | **Sites** | **Alignment patterns** | **Gaps** |
|---|---|---|---|---|
| All genes with paralogs | NT | 1544271 | 1099898 | 0.38 |
| All genes with paralogs | AA | 514757 | 429244 | 0.38 |
| All genes without paralogs | NT | 1091682 | 945847 | 0.116 |
| All genes without paralogs | AA | 363894 | 348473 | 0.116 |
| Single-copy genes | NT | 944871 | 824713 | 0.119 |
| Single-copy genes | AA | 314957 | 302861 | 0.119 |
| Chloroplast | NT | 65409 | 36662 | 0.488 |

**Table S12.** Robinson-Foulds distances between various species trees. Abbreviations are as follows 'AS' = ASTRAL-3; 'RA' = RAxML; '997' = all genes without paralogs, '997p'= all genes with paralogs, 'SC' = single-copy genes; 'NT' = nucleotide alignment; 'AA' = amino acid alignment.

| **Robinson-Foulds distances** | AS-SC | AS-997 | AS-997p | RA-SC-NT | RA-997-NT | RA-997p-NT | RA-SC-AA | RA-997-AA | RA-997p-AA | Phy-loBayes |
|---|---|---|---|---|---|---|---|---|---|---|
| AS-997 | 14 | | | | | | | | | |
| AS-997p | 10 | 12 | | | | | | | | |
| RA-SC-NT | 98 | 98 | 100 | | | | | | | |
| RA-997-NT | 98 | 102 | 100 | 28 | | | | | | |
| RA-997p-NT | 86 | 86 | 88 | 20 | 34 | | | | | |
| RA-SC-AA | 94 | 92 | 94 | 30 | 50 | 34 | | | | |
| RA-997-AA | 90 | 90 | 92 | 38 | 44 | 34 | 16 | | | |
| RA-997p-AA | 90 | 90 | 92 | 30 | 46 | 26 | 8 | 12 | | |
| PhyloBayes | 83 | 83 | 83 | 65 | 63 | 63 | 63 | 65 | 63 | |
| Chloroplast | 288 | 290 | 288 | 306 | 306 | 300 | 300 | 302 | 300 | 265 |

**Table S13.** Number of times taxa were identified as rogue taxa across 54 RogueNaRok analyses.

| Taxon | Times identified as rogue |
|---|---|
| *Zygia inaequalis* | 12 |
| *Zygia longifolia* | 8 |
| *Archidendron quocense* | 7 |
| *Archidendron lucidum* | 7 |
| *Inga cinnamomea* | 7 |
| *Leucochloron limae* | 6 |
| *Zygia ramiflora* | 5 |
| *Paraserianthes lophantha* | 3 |
| *Abarema langsdorfii* | 2 |
| *Hydrochorea corymbosa* 1 | 2 |
| *Hydrochorea corymbosa* 2 | 2 |
| *Albizia obbiadensis* | 1 |
| *Albizia viridis* | 1 |
| *Archidendron ellipticum* subsp. *ellipticum* | 1 |
| *Archidendron triplinervium* | 1 |
| *Caesalpinia cassioides* | 1 |
| *Albizia obliquifoliolata* | 1 |
| *Albizia rhombifolia* | 1 |
| *Denisophytum madagascariense* | 1 |
| *Hydrochorea gonggrijpii* | 1 |
| *Hydrochorea marginata* | 1 |
| *Inga heterophylla* | 1 |
| *Inga setosa* | 1 |
| *Inga stipularis* | 1 |
| *Samanea saman* | 1 |
| *Wallaceodendron celebicum* | 1 |

**Table S14.** Numbers of accepted taxa, accepted species, and synonyms in the taxonomic checklist, and numbers of records, taxa, and species in the occurrence dataset. See Appendix 1 for details about the assembly and contents of the checklist and occurrence dataset, and Appendix 2 for the full taxonomic checklist. Note that the number of species listed for *Abarema* includes five poorly-known taxa unlikely to belong to *Abarema*; see Appendix 1 for details.

| | Taxonomic checklist | | | Occurrence dataset | | |
|---|---|---|---|---|---|---|
| **Genus** | **Accepted taxa** | **Accepted species** | **Synonyms** | **Occurrences** | **Taxa** | **Species** |
| ***Abarema*** | 7 | 7 | 13 | 407 | 5 | 5 |
| ***Acacia*** | 1098 | 1093 | 64 | 178367 | 1074 | 1068 |
| ***Acaciella*** | 25 | 15 | 107 | 4521 | 23 | 14 |
| ***Adenanthera*** | 15 | 13 | 13 | 628 | 11 | 10 |
| ***Adenopodia*** | 7 | 7 | 18 | 364 | 7 | 7 |
| ***Afrocalliandra*** | 2 | 2 | 3 | 51 | 2 | 2 |
| ***Alantsilodendron*** | 9 | 9 | 8 | 145 | 9 | 9 |
| ***Albizia* (New World)** | 30 | 24 | 83 | 3483 | 26 | 20 |
| ***Albizia* (Old World)** | 127 | 93 | 234 | 9078 | 111 | 90 |
| ***Amblygonocarpus*** | 1 | 1 | 6 | 82 | 1 | 1 |
| ***Anadenanthera*** | 6 | 2 | 38 | 4360 | 6 | 2 |
| ***Archidendron*** | 120 | 99 | 388 | 4170 | 97 | 87 |
| ***Archidendropsis*** | 19 | 14 | 20 | 489 | 19 | 14 |
| ***Aubrevillea*** | 2 | 2 | 2 | 112 | 2 | 2 |
| ***Balizia*** | 3 | 3 | 12 | 528 | 3 | 3 |
| ***Blanchetiodendron*** | 1 | 1 | 4 | 154 | 1 | 1 |
| ***Calliandra*** | 203 | 148 | 464 | 16053 | 188 | 137 |
| ***Calliandra umbrosa*** | 3 | 1 | 6 | 0 | 0 | 0 |
| ***Calliandropsis*** | 1 | 1 | 2 | 68 | 1 | 1 |
| ***Calpocalyx*** | 11 | 11 | 10 | 755 | 11 | 11 |
| ***Cedrelinga*** | 1 | 1 | 3 | 249 | 1 | 1 |
| ***Chidlowia*** | 1 | 1 | 0 | 137 | 1 | 1 |
| ***Chloroleucon*** | 17 | 11 | 68 | 1802 | 17 | 11 |
| ***Cojoba*** | 19 | 14 | 191 | 2527 | 17 | 12 |
| ***Cylicodiscus*** | 1 | 1 | 1 | 90 | 1 | 1 |
| ***Desmanthus*** | 28 | 24 | 107 | 3391 | 26 | 24 |
| ***Dichrostachys*** | 33 | 13 | 59 | 1050 | 14 | 13 |
| ***Ebenopsis*** | 3 | 3 | 15 | 258 | 3 | 3 |
| ***Elephantorrhiza*** | 15 | 9 | 16 | 925 | 12 | 9 |
| ***Entada*** | 41 | 31 | 82 | 3490 | 35 | 28 |
| ***Enterolobium*** | 8 | 8 | 3 | 2768 | 8 | 8 |
| ***Faidherbia*** | 1 | 1 | 8 | 624 | 1 | 1 |
| ***Falcataria*** | 3 | 3 | 11 | 433 | 3 | 3 |
| ***Fillaeopsis*** | 1 | 1 | 0 | 84 | 1 | 1 |
| ***Gagnebina*** | 10 | 8 | 19 | 168 | 10 | 8 |
| ***Havardia*** | 5 | 5 | 22 | 1283 | 5 | 5 |
| ***Hesperalbizia*** | 1 | 1 | 7 | 465 | 1 | 1 |
| ***Hydrochorea*** | 6 | 3 | 21 | 862 | 6 | 3 |
| ***Indopiptadenia*** | 1 | 1 | 1 | 25 | 1 | 1 |
| ***Inga*** | 288 | 276 | 609 | 39332 | 279 | 266 |

| | | | | | | |
|---|---|---|---|---|---|---|
| ***Jupunba*** | 52 | 37 | 164 | 3246 | 51 | 37 |
| ***Kanaloa*** | 1 | 1 | 0 | 1 | 1 | 1 |
| ***Lachesiodendron*** | 1 | 1 | 10 | 662 | 1 | 1 |
| ***Lemurodendron*** | 1 | 1 | 0 | 5 | 1 | 1 |
| ***Leucaena*** | 33 | 26 | 67 | 2172 | 25 | 23 |
| ***Leucochloron*** | 5 | 5 | 7 | 255 | 5 | 5 |
| ***Lysiloma*** | 8 | 8 | 45 | 5577 | 8 | 8 |
| ***Macrosamanea*** | 14 | 12 | 33 | 642 | 13 | 11 |
| ***Mariosousa*** | 13 | 13 | 30 | 1566 | 13 | 13 |
| ***Microlobius*** | 3 | 1 | 12 | 253 | 2 | 1 |
| ***Mimosa*** | 832 | 589 | 621 | 36653 | 737 | 530 |
| ***Mimozyganthus*** | 1 | 1 | 1 | 60 | 1 | 1 |
| ***Neptunia*** | 17 | 11 | 50 | 2220 | 13 | 10 |
| ***Newtonia*** | 17 | 15 | 24 | 595 | 17 | 15 |
| ***Painteria*** | 3 | 3 | 14 | 139 | 3 | 3 |
| ***Parapiptadenia*** | 6 | 6 | 8 | 1409 | 6 | 6 |
| ***Pararchidendron*** | 5 | 1 | 13 | 506 | 4 | 1 |
| ***Parasenegalia*** | 11 | 11 | 40 | 267 | 10 | 10 |
| ***Paraserianthes*** | 3 | 1 | 17 | 228 | 2 | 1 |
| ***Parkia*** | 44 | 38 | 68 | 3323 | 39 | 33 |
| ***Pentaclethra*** | 3 | 3 | 4 | 1135 | 3 | 3 |
| ***Piptadenia*** | 33 | 33 | 11 | 2098 | 26 | 26 |
| ***Piptadeniastrum*** | 1 | 1 | 1 | 332 | 1 | 1 |
| ***Piptadeniopsis*** | 1 | 1 | 0 | 12 | 1 | 1 |
| ***Pithecellobium*** | 20 | 18 | 309 | 5346 | 20 | 18 |
| ***Pityrocarpa*** | 5 | 3 | 8 | 1311 | 4 | 3 |
| ***Plathymenia*** | 1 | 1 | 5 | 1517 | 1 | 1 |
| ***Prosopidastrum*** | 5 | 5 | 9 | 80 | 5 | 5 |
| ***Prosopis*** | 91 | 56 | 98 | 6410 | 67 | 47 |
| ***Pseudopiptadenia*** | 11 | 11 | 17 | 1333 | 10 | 10 |
| ***Pseudoprosopis*** | 9 | 7 | 9 | 353 | 8 | 7 |
| ***Pseudosamanea*** | 2 | 2 | 8 | 336 | 2 | 2 |
| ***Pseudosenegalia*** | 2 | 2 | 4 | 11 | 2 | 2 |
| ***Punjuba*** | 6 | 6 | 16 | 184 | 5 | 5 |
| ***Robrichia*** | 3 | 3 | 6 | 557 | 3 | 3 |
| ***Samanea*** | 3 | 3 | 20 | 839 | 3 | 3 |
| ***Sanjappa*** | 1 | 1 | 1 | 1 | 1 | 1 |
| ***Schleinitzia*** | 4 | 4 | 13 | 71 | 3 | 3 |
| ***Senegalia*** | 256 | 221 | 480 | 20636 | 214 | 190 |
| ***Serianthes*** | 27 | 18 | 19 | 168 | 21 | 17 |
| ***Sphinga*** | 3 | 3 | 15 | 563 | 3 | 3 |
| ***Stryphnodendron*** | 38 | 36 | 24 | 2480 | 28 | 26 |
| ***Sympetalandra*** | 5 | 5 | 4 | 74 | 5 | 5 |
| ***Tetrapleura*** | 2 | 2 | 2 | 314 | 2 | 2 |
| ***Thailentadopsis*** | 3 | 3 | 24 | 23 | 3 | 3 |

| | | | | | | |
|---|---|---|---|---|---|---|
| ***Vachellia*** | 233 | 169 | 556 | 24434 | 200 | 146 |
| ***Viguieranthus*** | 24 | 18 | 11 | 334 | 19 | 18 |
| ***Wallaceodendron*** | 1 | 1 | 4 | 33 | 1 | 1 |
| ***Xerocladia*** | 1 | 1 | 2 | 34 | 1 | 1 |
| ***Xylia*** | 11 | 9 | 13 | 488 | 10 | 9 |
| ***Zapoteca*** | 35 | 23 | 116 | 4031 | 31 | 20 |
| ***Zygia*** | 66 | 58 | 384 | 5238 | 61 | 56 |
| **Total** | 4149 | 3469 | 6155 | 424333 | 3755 | 3233 |

**Table S15.** Phylogenetic turnover variation partitioning results obtained with the metachronogram per region. The relative contribution of each predictor variable is shown, as well as the total deviance explained by the full model.

| | **pantropics** | **North America** | **South America** | **Africa** | **Asia** | **Australia** |
|---|---|---|---|---|---|---|
| **Climatic distance** | 0.04 | 0.48 | 0.55 | 0.63 | 0.20 | 0.47 |
| **Interaction** | 0.31 | 0.51 | 0.40 | 0.30 | 0.39 | 0.47 |
| **Spatial distance** | 0.64 | 0.01 | 0.05 | 0.06 | 0.41 | 0.07 |
| **Percentage deviance explained** | 54.47 | 29.92 | 21.84 | 28.10 | 34.58 | 44.67 |
| **Precipitation** | 0.75 | 0.85 | 0.80 | 1.00 | 0.59 | 0.47 |
| **Interaction** | -0.01 | 0.11 | 0.08 | 0.00 | -0.00 | 0.14 |
| **Temperature** | 0.26 | 0.04 | 0.12 | 0.00 | 0.41 | 0.39 |
| **Percentage deviance explained** | 3.00 | 9.10 | 6.69 | 15.00 | 2.31 | 16.20 |
| **Precipitation** | 0.60 | 0.83 | 0.61 | 0.93 | 0.04 | 0.51 |
| **Interaction** | 0.04 | 0.08 | 0.31 | 0.07 | 0.17 | 0.19 |
| **Precipitation seasonality** | 0.36 | 0.09 | 0.08 | 0.00 | 0.79 | 0.29 |
| **Percentage deviance explained** | 3.48 | 9.58 | 6.41 | 15.02 | 6.37 | 13.93 |
| **Precipitation** | 0.05 | 0.28 | 0.38 | 0.40 | 0.00 | 0.19 |
| **Interaction** | 0.73 | 0.66 | 0.60 | 0.56 | 0.23 | 0.73 |
| **Dry season length** | 0.22 | 0.06 | 0.02 | 0.04 | 0.77 | 0.08 |
| **Percentage deviance explained** | 2.84 | 9.31 | 6.04 | 15.66 | 5.96 | 10.68 |
| **Precipitation** | 0.77 | 0.90 | 0.91 | 0.81 | 0.40 | 0.49 |
| **Interaction** | -0.07 | -0.04 | -0.04 | 0.16 | 0.02 | 0.20 |
| **SD of cloud cover** | 0.29 | 0.14 | 0.14 | 0.03 | 0.58 | 0.31 |
| **Percentage deviance explained** | 3.14 | 10.13 | 6.84 | 15.51 | 3.24 | 14.19 |

**Table S16.** Phylogenetic turnover variation partitioning results obtained with the genus-level Mimosoid tree per region. The relative contribution of each predictor variable is shown, as well as the total deviance explained by the full model.

| | pantropics | North America | South America | Africa | Asia | Australia |
|---|---|---|---|---|---|---|
| **Climatic distance** | 0.11 | 0.43 | 0.41 | 0.42 | 0.09 | 0.41 |
| **Interaction** | 0.33 | 0.53 | 0.49 | 0.46 | 0.45 | 0.50 |
| **Spatial distance** | 0.56 | 0.04 | 0.09 | 0.12 | 0.47 | 0.08 |
| **Percentage deviance explained** | 51.98 | 36.74 | 31.86 | 41.65 | 38.63 | 53.41 |

| | | | | | | |
|---|---|---|---|---|---|---|
| **Precipitation** | 0.72 | 0.83 | 0.75 | 0.99 | 0.61 | 0.54 |
| **Interaction** | 0.01 | 0.07 | 0.07 | -0.00 | 0.08 | 0.16 |
| **Temperature** | 0.27 | 0.09 | 0.19 | 0.01 | 0.31 | 0.30 |
| **Percentage deviance explained** | 5.45 | 10.88 | 7.03 | 18.13 | 1.57 | 16.90 |

| | | | | | | |
|---|---|---|---|---|---|---|
| **Precipitation** | 0.64 | 0.89 | 0.77 | 0.92 | 0.06 | 0.41 |
| **Interaction** | 0.04 | 0.06 | 0.18 | 0.07 | 0.23 | 0.27 |
| **Precipitation seasonality** | 0.33 | 0.05 | 0.05 | 0.00 | 0.71 | 0.32 |
| **Percentage deviance explained** | 5.89 | 10.34 | 6.06 | 18.07 | 3.76 | 17.51 |

| | | | | | | |
|---|---|---|---|---|---|---|
| **Precipitation** | 0.13 | 0.32 | 0.33 | 0.44 | -0.00 | 0.05 |
| **Interaction** | 0.80 | 0.54 | 0.62 | 0.56 | 0.26 | 0.78 |
| **Dry season length** | 0.07 | 0.14 | 0.05 | 0.00 | 0.74 | 0.17 |
| **Percentage deviance explained** | 4.25 | 11.50 | 6.02 | 18.00 | 4.10 | 14.32 |

| | | | | | | |
|---|---|---|---|---|---|---|
| **Precipitation** | 0.84 | 0.89 | 0.93 | 0.73 | 0.39 | 0.38 |
| **Interaction** | -0.04 | -0.05 | -0.04 | 0.21 | -0.02 | 0.29 |
| **SD of cloud cover** | 0.20 | 0.16 | 0.11 | 0.06 | 0.63 | 0.34 |
| **Percentage deviance explained** | 4.94 | 11.70 | 6.43 | 19.14 | 2.91 | 17.87 |

**Table S17.** Fractions of trans-oceanic dispersal events per splitting event in the phylogeny through time, measured using eight different models and definitions of trans-oceanic dispersal, as well as the total deviance explained by the full model.

| Time period | DEC+j_7_ regions | DEC+j_3_ regions | DEC_7_ regions | DEC_3_ regions | BAY_7_ regions | BAY_3_ regions | DIVA_7_ regions | DIVA_3_ regions | Average |
|---|---|---|---|---|---|---|---|---|---|
| 0-5 | 0.02 | 0.00 | 0.02 | 0.00 | 0.02 | 0.00 | 0.02 | 0.00 | 0.01 |
| 5-10 | 0.02 | 0.01 | 0.02 | 0.01 | 0.02 | 0.01 | 0.02 | 0.01 | 0.02 |
| 10-15 | 0.03 | 0.02 | 0.04 | 0.02 | 0.03 | 0.02 | 0.04 | 0.03 | 0.03 |
| 15-20 | 0.03 | 0.02 | 0.03 | 0.02 | 0.02 | 0.02 | 0.03 | 0.03 | 0.03 |
| 20-25 | 0.09 | 0.06 | 0.09 | 0.05 | 0.09 | 0.06 | 0.11 | 0.06 | 0.08 |
| 25-30 | 0.05 | 0.03 | 0.05 | 0.05 | 0.03 | 0.00 | 0.11 | 0.05 | 0.05 |
| 30-35 | 0.09 | 0.05 | 0.14 | 0.09 | 0.05 | 0.05 | 0.14 | 0.09 | 0.09 |
| 35-40 | 0.00 | 0.00 | 0.14 | 0.14 | 0.00 | 0.00 | 0.00 | 0.00 | 0.04 |

### 6. Supplementary figures

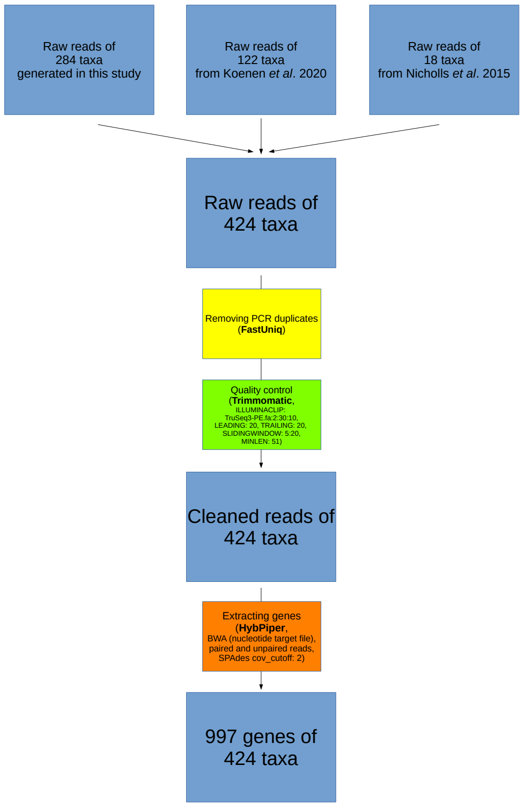

**Figure S1.** Overview of the data cleaning and target assembly workflow.

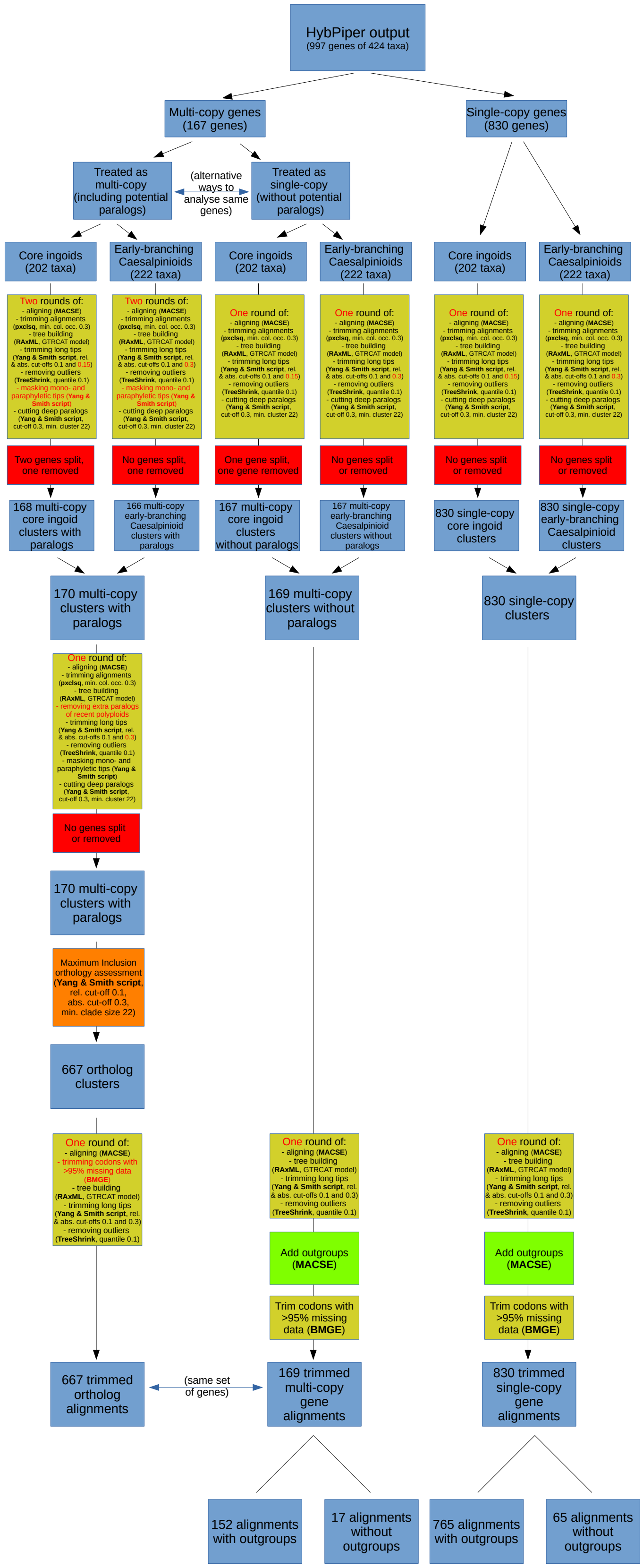

**Figure S2.** Overview and results of the orthology assessment workflow.

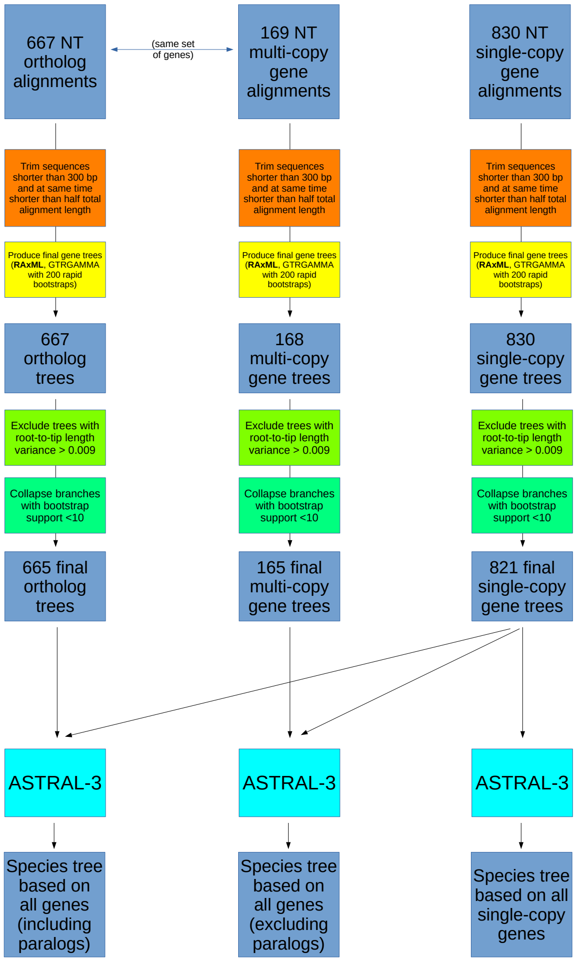

**Figure S3.** Overview of species tree inference workflow with ASTRAL-3.

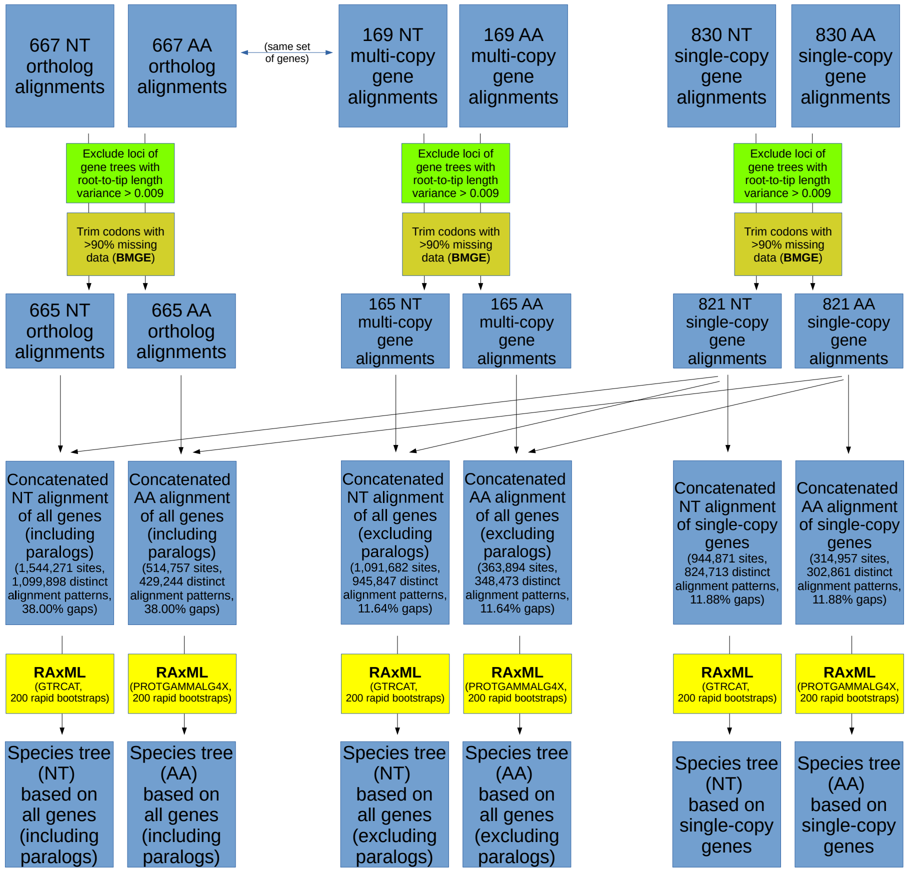

**Figure S4.** Overview of species tree inference workflow with RAxML.

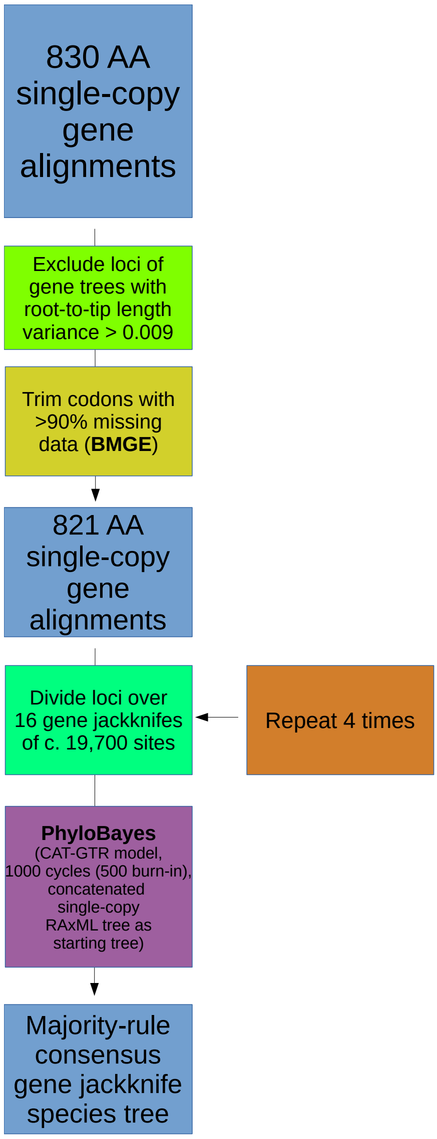

**Figure S5.** Overview of species tree inference workflow with PhyloBayes.

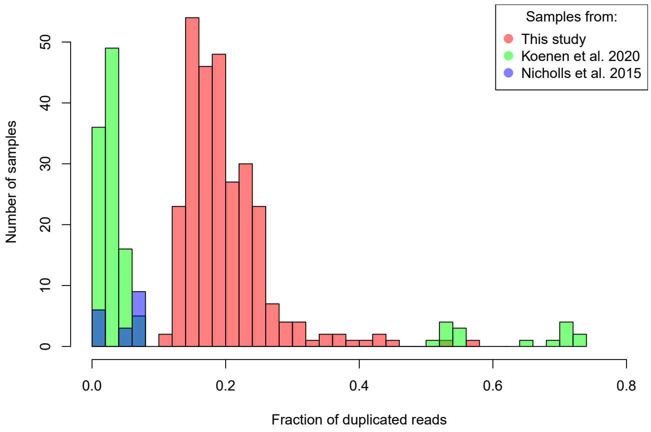

**Figure S6 (left).** Fractions of duplicated reads per sample.

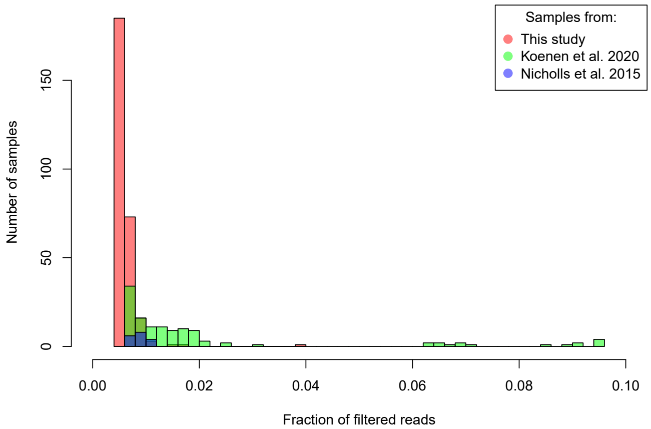

**Figure S7 (left).** Fractions of filtered reads per sample.

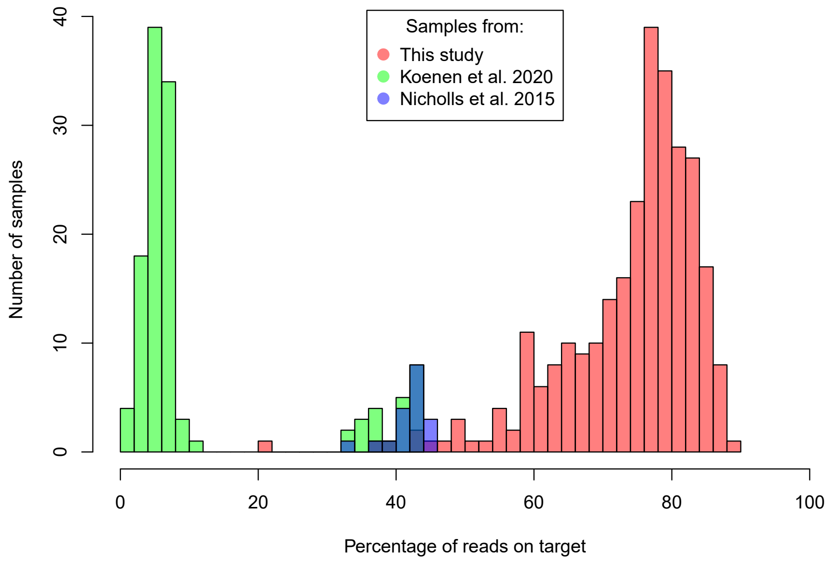

**Figure S8 (left).** Percentages of reads on target per sample.

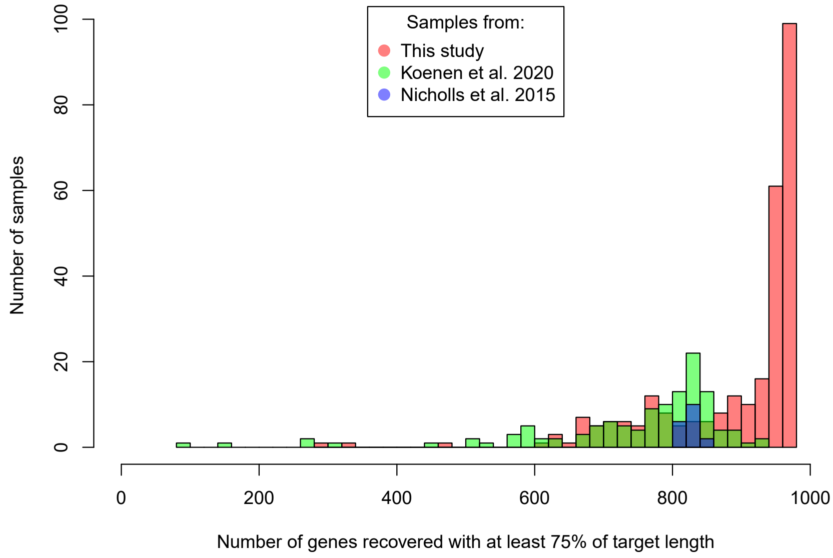

**Figure S9 (left).** Numbers of genes recovered with at least 75% of the target length per sample.

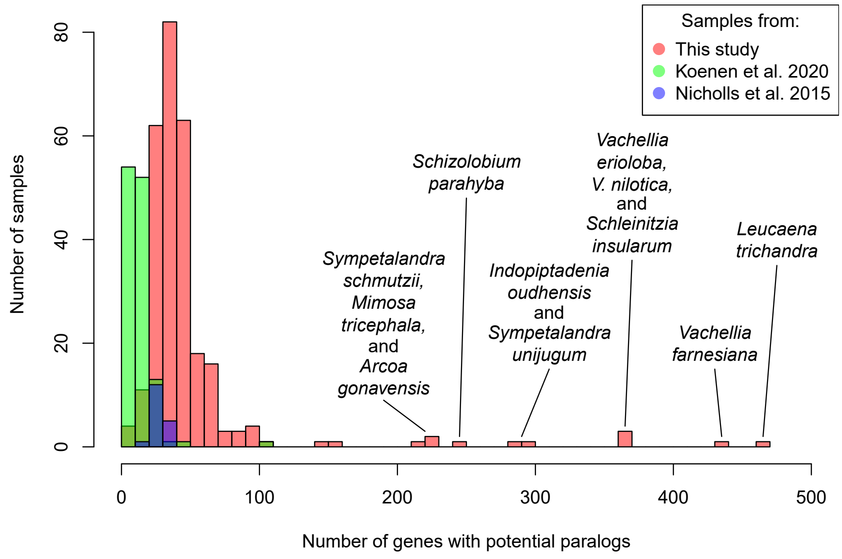

**Figure S10 (left).** Numbers of genes with potential paralogs per sample. The eleven samples with the most potential paralogs are labelled.

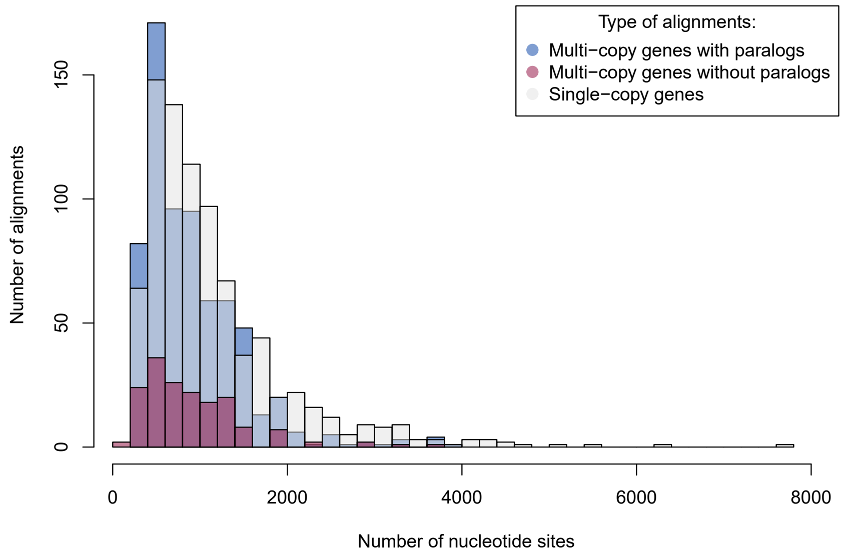

**Figure S11 (left).** Numbers of nucleotide sites per alignment.

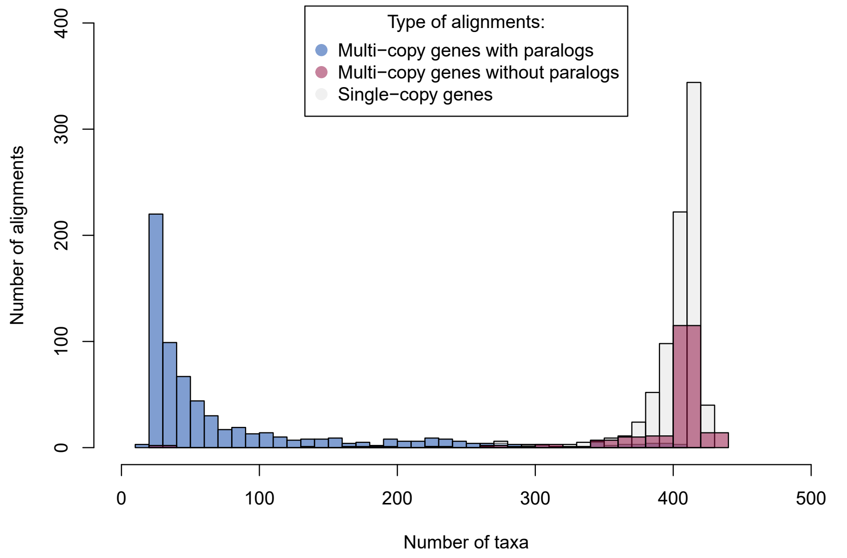

**Figure S12 (left).** Numbers of taxa per alignment.

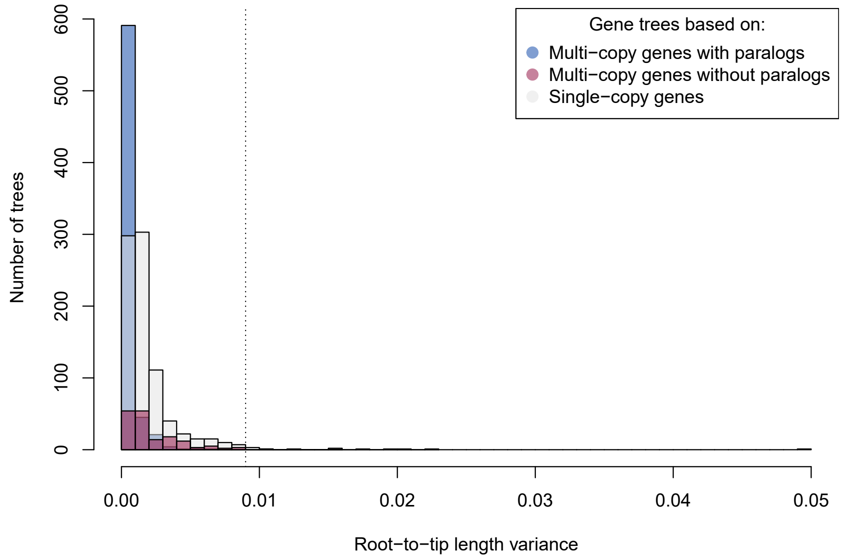

**Figure S13 (left).** Root-to-tip length variances per gene tree. The cut-off of 0.009 is indicated with a dashed vertical line.

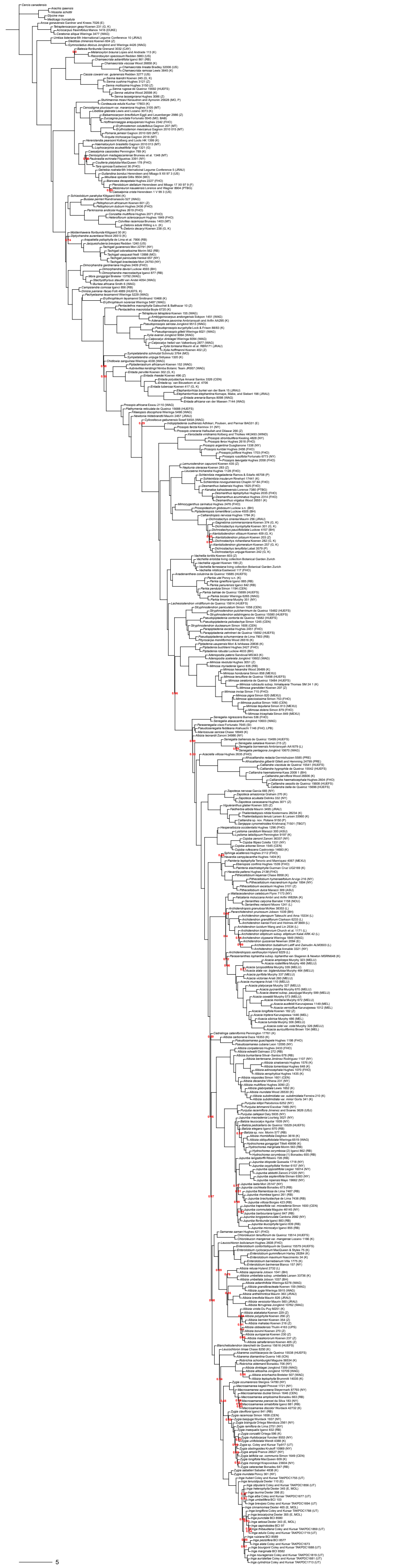

**Figure S14.** Phylogeny of Caesalpinioideae. ASTRAL species tree based on the 821 single-copy gene trees. Local posterior probability support values are only shown for nodes with a local posterior probability < 1. Branch lengths are expressed in coalescent units. Terminal branches were assigned an arbitrary uniform length for visual clarity.

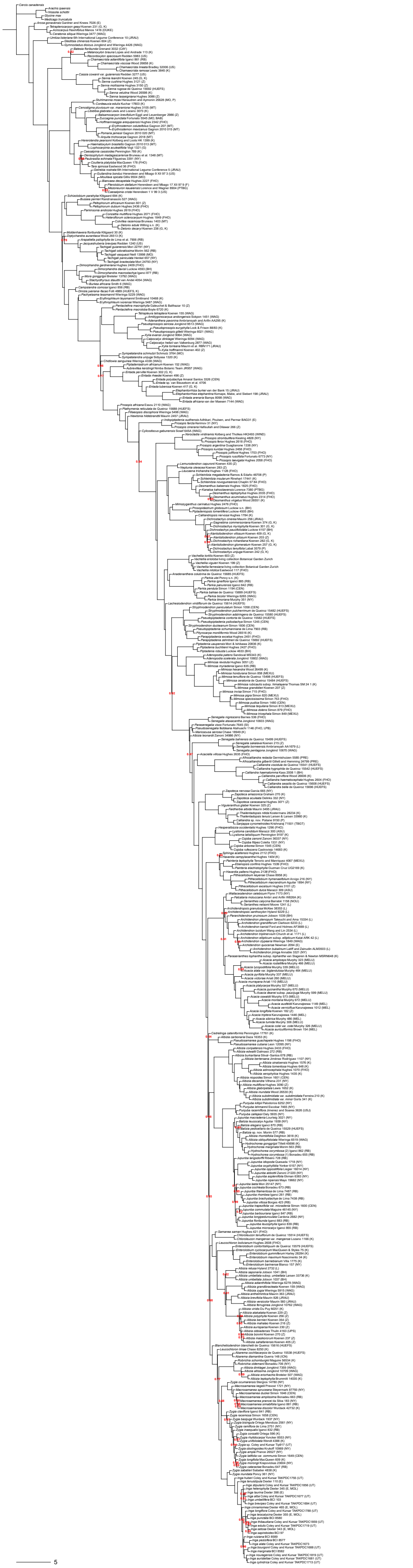

**Figure S15.** Phylogeny of Caesalpinioideae. ASTRAL species tree based on the 821 single-copy gene trees and the 165 multi-copy gene trees without orthology assessment. Local posterior probability support values are only shown for nodes with a local posterior probability < 1. Branch lengths are expressed in coalescent units. Terminal branches were assigned an arbitrary uniform length for visual clarity.

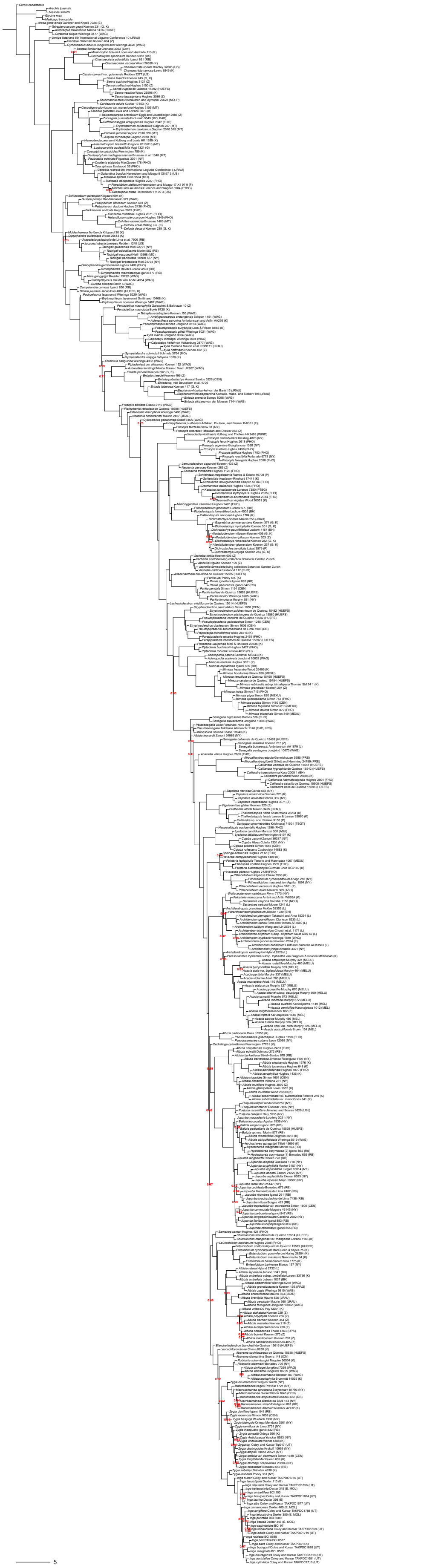

**Figure S16.** Phylogeny of Caesalpinioideae. ASTRAL species tree based on the 821 single-copy gene trees and the 665 ortholog trees resulting from orthology assessment. Local posterior probability support values are only shown for nodes with a local posterior probability < 1. Branch lengths are expressed in coalescent units. Terminal branches were assigned an arbitrary uniform length for visual clarity.

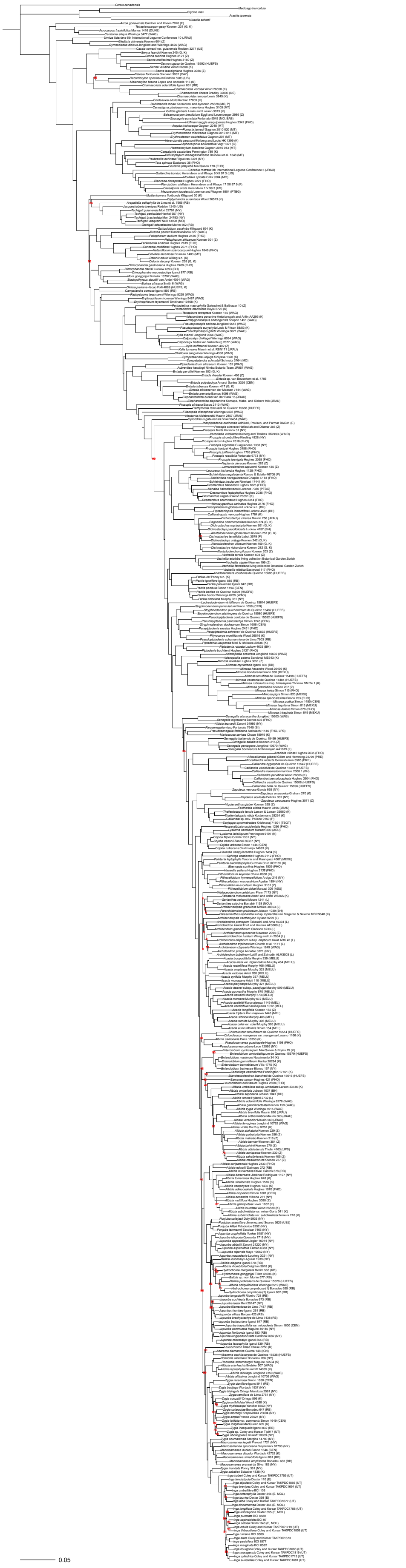

**Figure S17.** Phylogeny of Caesalpinioideae. RAxML species tree based on the nucleotide single-copy genes alignment. Bootstrap support values are only shown for nodes with < 100% bootstrap support.

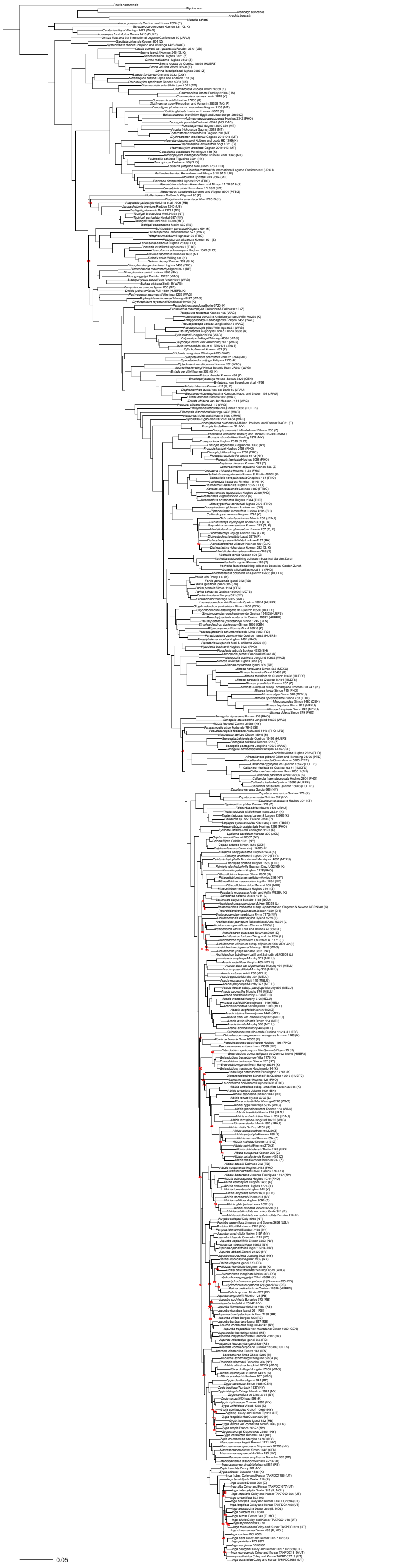

**Figure S18.** Phylogeny of Caesalpinioideae. RAxML species tree based on the nucleotide alignment of all genes without orthology assessment. Bootstrap support values are only shown for nodes with < 100% bootstrap support.

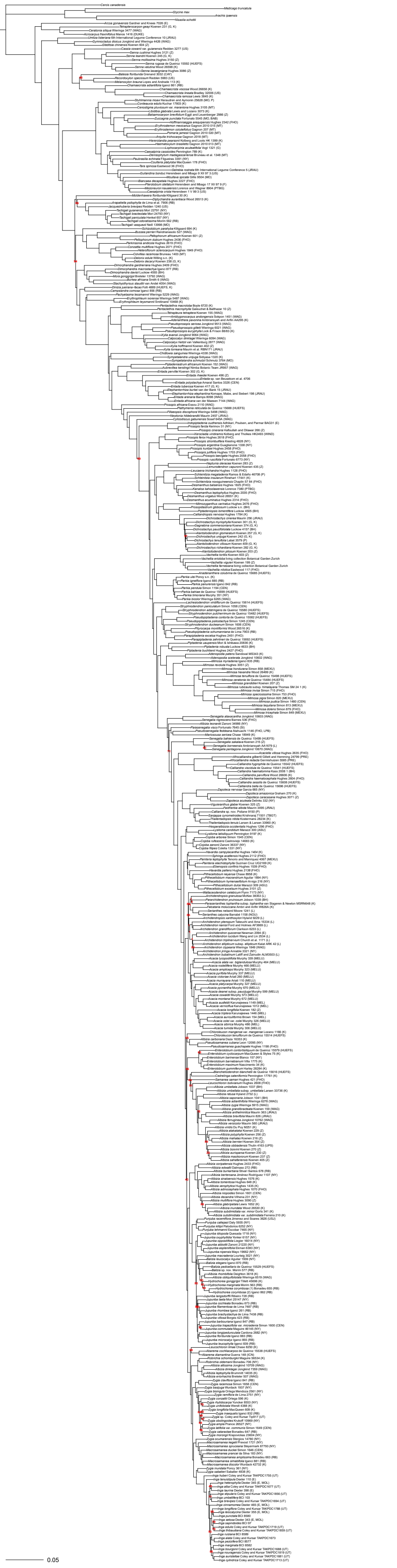

**Figure S19.** Phylogeny of Caesalpinioideae. RAxML species tree based on the nucleotide alignment of all genes with orthology assessment. Bootstrap support values are only shown for nodes with < 100% bootstrap support.

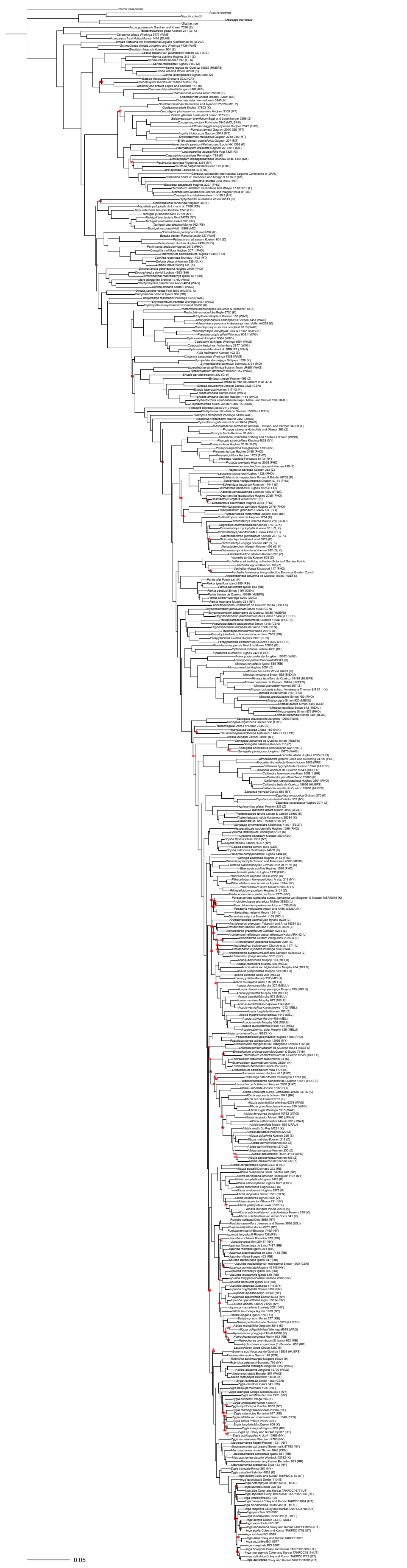

**Figure S20.** Phylogeny of Caesalpinioideae. RAxML species tree based on the amino acid single-copy genes alignment. Bootstrap support values are only shown for nodes with < 100% bootstrap support.

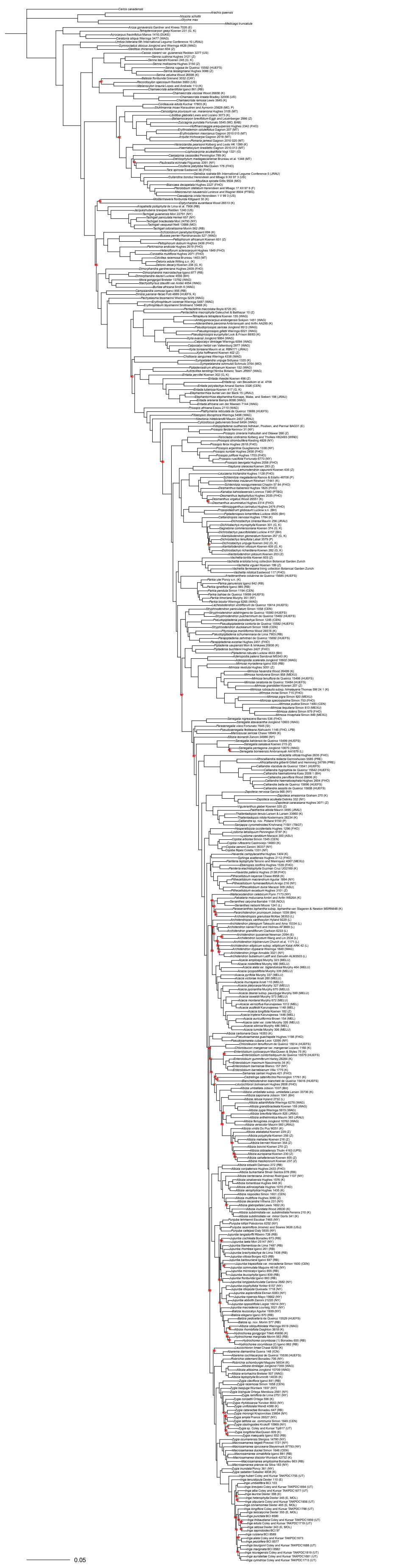

**Figure S21.** Phylogeny of Caesalpinioideae. RAxML species tree based on the amino acid alignment of all genes without orthology assessment. Bootstrap support values are only shown for nodes with < 100% bootstrap support.

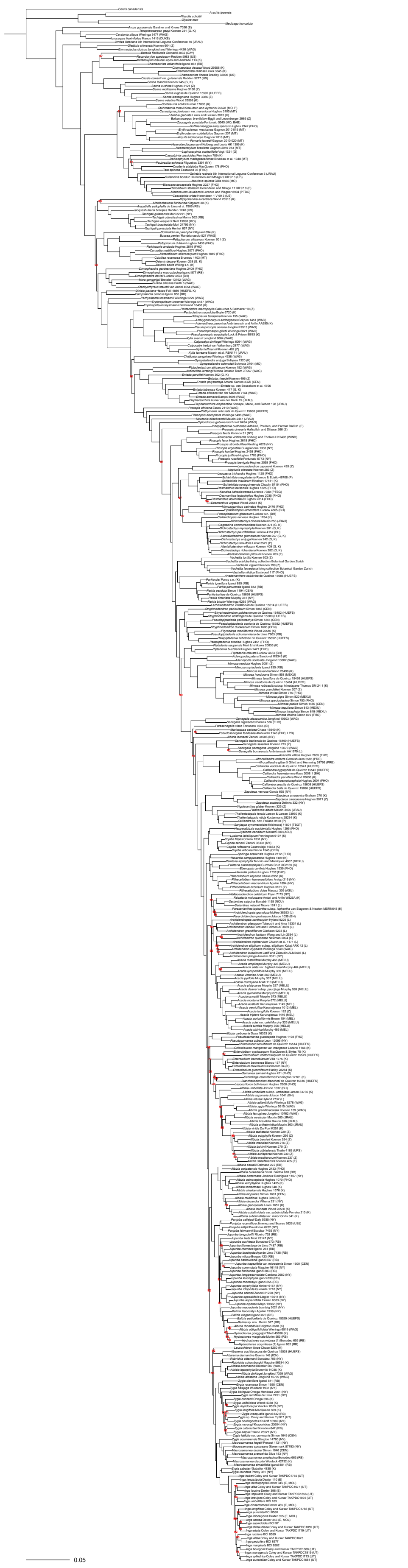

**Figure S22.** Phylogeny of Caesalpinioideae. RAxML species tree based on the amino acid alignment of all genes with orthology assessment. Bootstrap support values are only shown for nodes with < 100% bootstrap support.

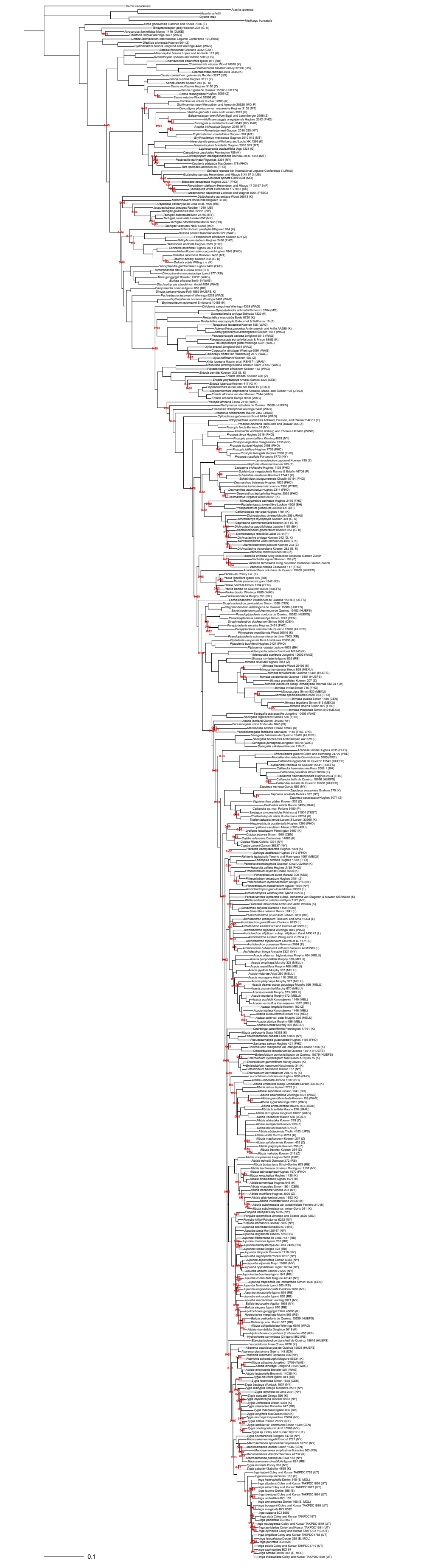

**Figure S23.** Phylogeny of Caesalpinioideae. PhyloBayes species tree. Posterior probability support values are only shown for nodes with a posterior probability < 1.

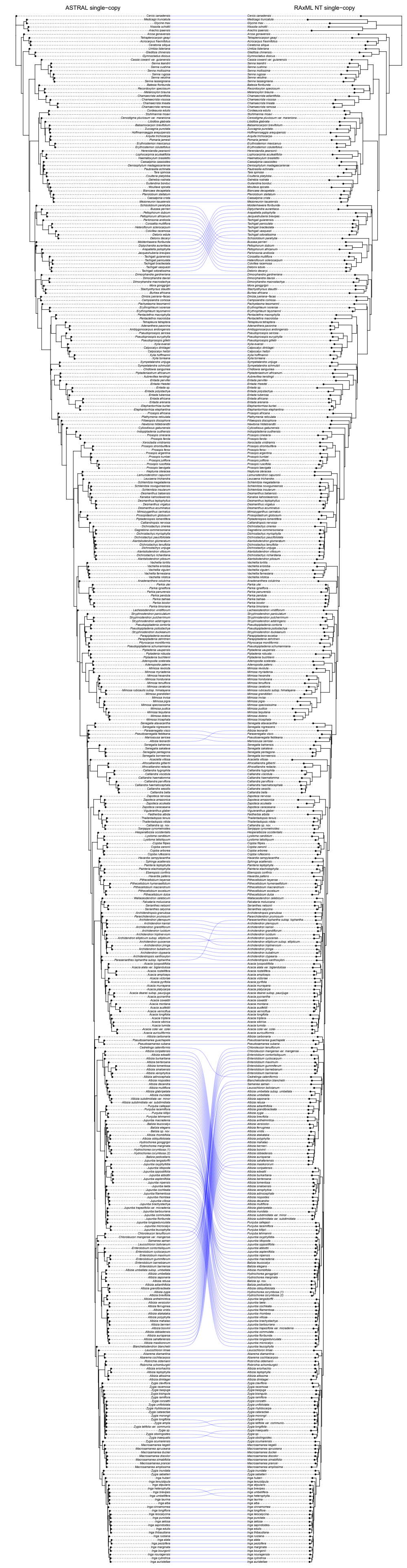

**Figure S24.** Tanglegram comparing the ASTRAL single-copy genes phylogeny (Figure S14) with the RAxML nucleotide single-copy genes phylogeny (Figure S17).

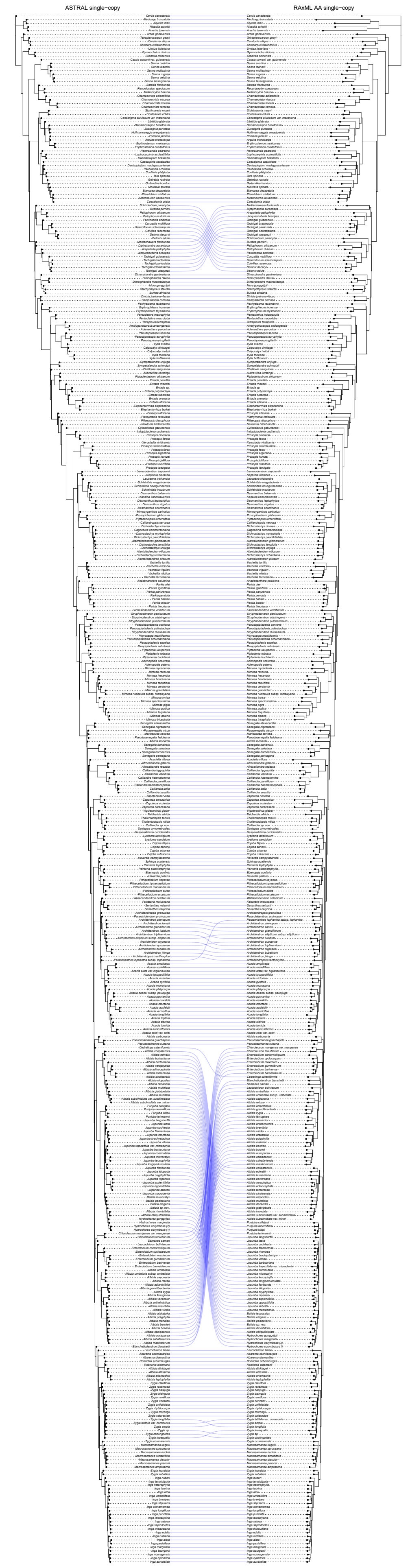

**Figure S25.** Tanglegram comparing the ASTRAL single-copy genes phylogeny (Figure S14) with the RAxML amino acid single-copy genes phylogeny (Figure S20).

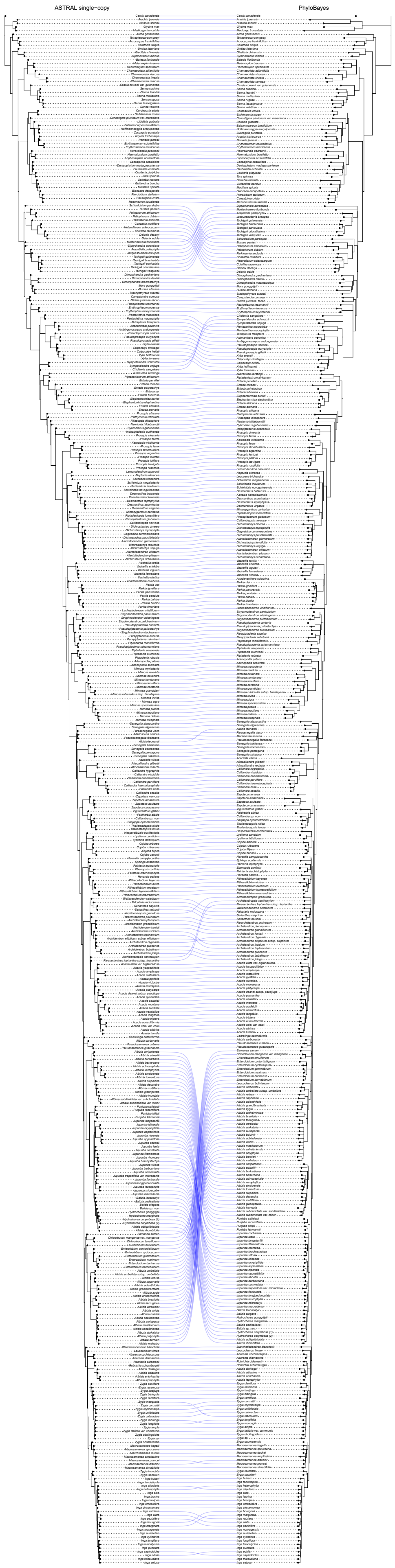

**Figure S26.** Tanglegram comparing the ASTRAL single-copy genes phylogeny (Figure S14) with the PhyloBayes phylogeny (Figure S23).

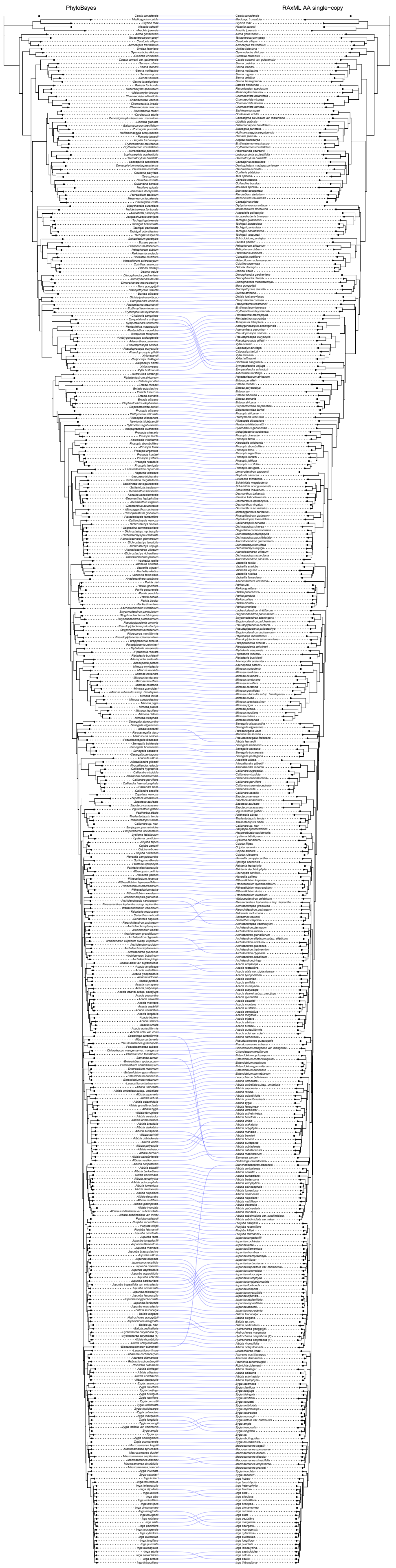

**Figure S27.** Tanglegram comparing the PhyloBayes phylogeny (Figure S23) with the RAxML amino acid single-copy genes phylogeny (Figure S20).

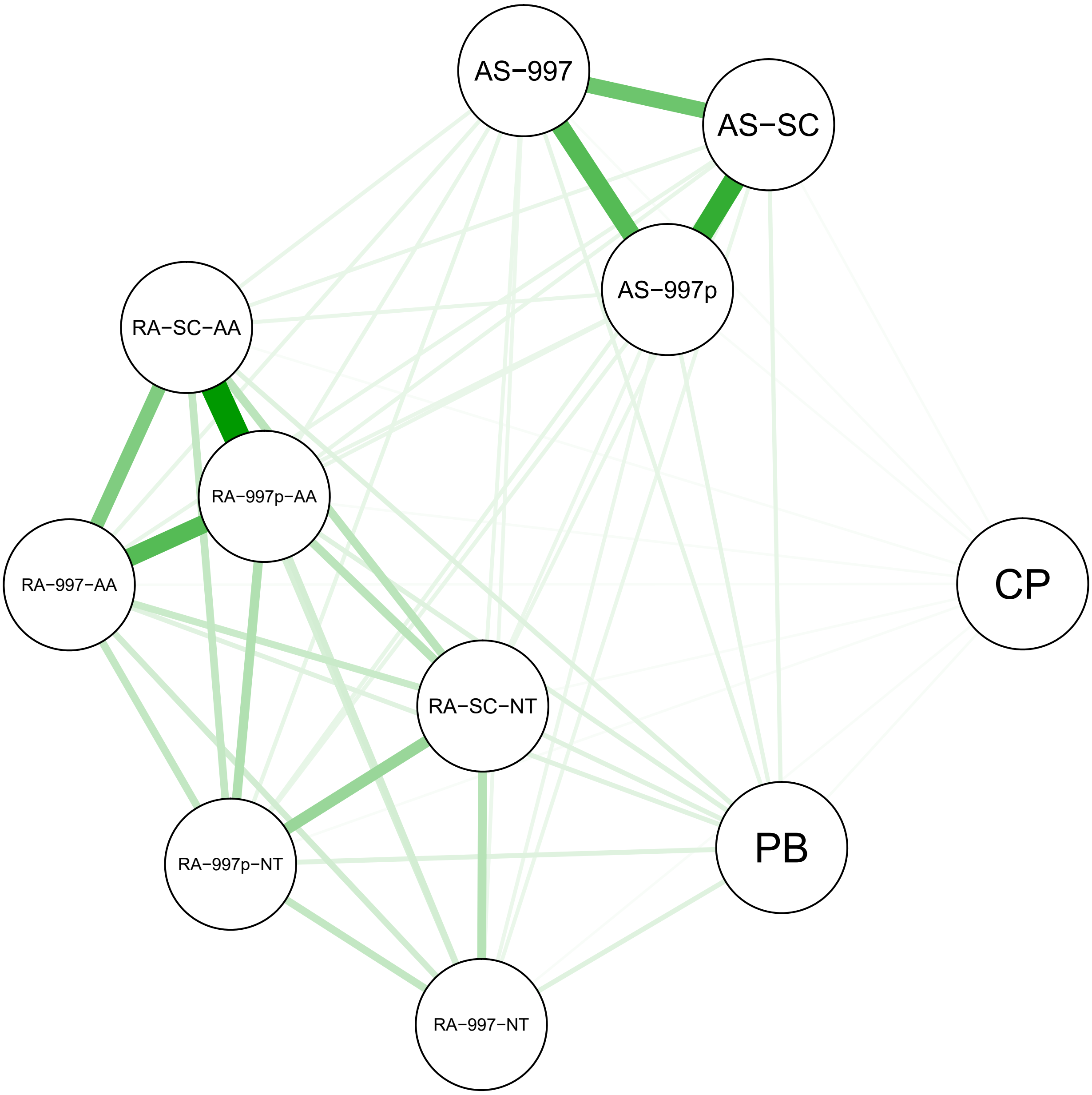

**Figure S28.** Levels of topological congruence between phylogenies generated in different ways, estimated as Robinson-Foulds (RF) distances between species trees. Exact values are in Table S12. Thickness of the connecting lines reflects RF distance. Abbreviations are as follows 'AS' = ASTRAL-3; 'RA' = RAxML; '997' = all genes without paralogs, '997p'= all genes with paralogs, 'SC' = single-copy genes; 'NT' = nucleotide alignment; 'AA' = amino acid alignment; 'PB' = PhyloBayes phylogeny; 'CP' = chloroplast phylogeny.

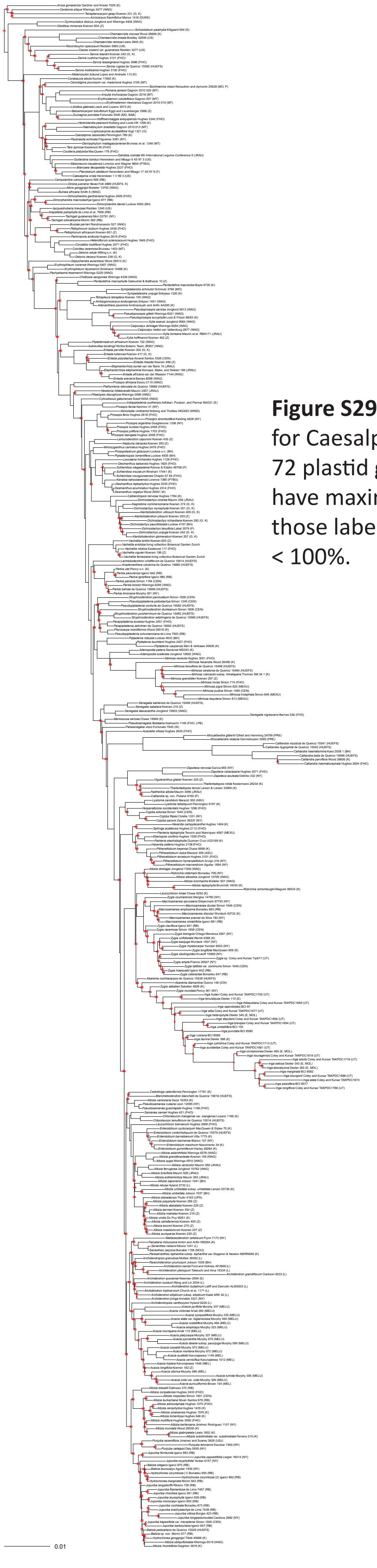

**Figure S29 (left).** Chloroplast gene tree for Caesalpinioideae based on analysis of 72 plastid genes using RAxML. All nodes have maximal bootstrap support except those labelled with actual support values < 100%.

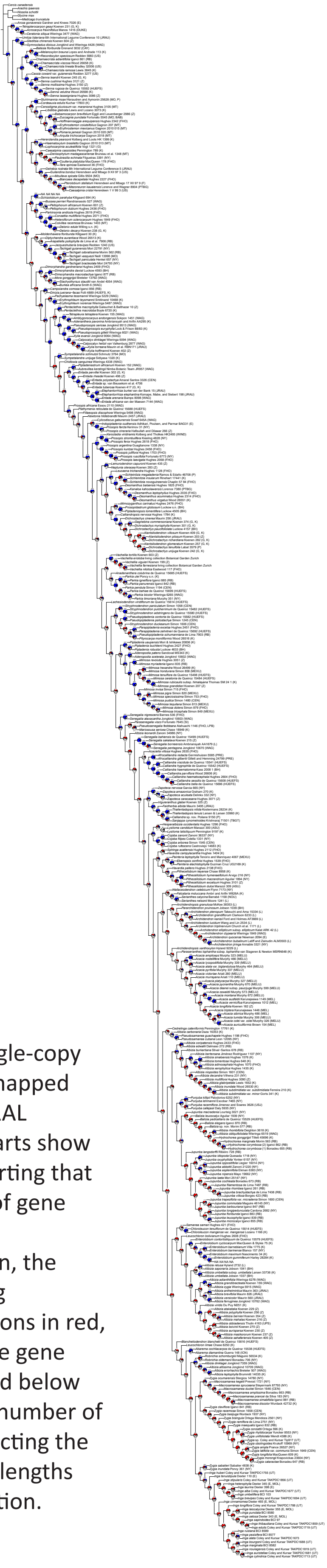

**Figure S30 (right).** Conflict and concordance among the 821 single-copy gene trees for each bipartition mapped onto the single-copy genes ASTRAL species tree (Figure S14). Pie charts show the fraction of gene trees supporting that bipartition in blue, the fraction of gene trees supporting the most likely alternative configuration in green, the fraction of gene trees supporting additional conflicting configurations in red, and the fraction of uninformative gene trees in grey. Numbers above and below the pie charts indicate the total number of gene trees supporting and conflicting the bipartition, respectively. Branch lengths are set equal for easier visualisation.

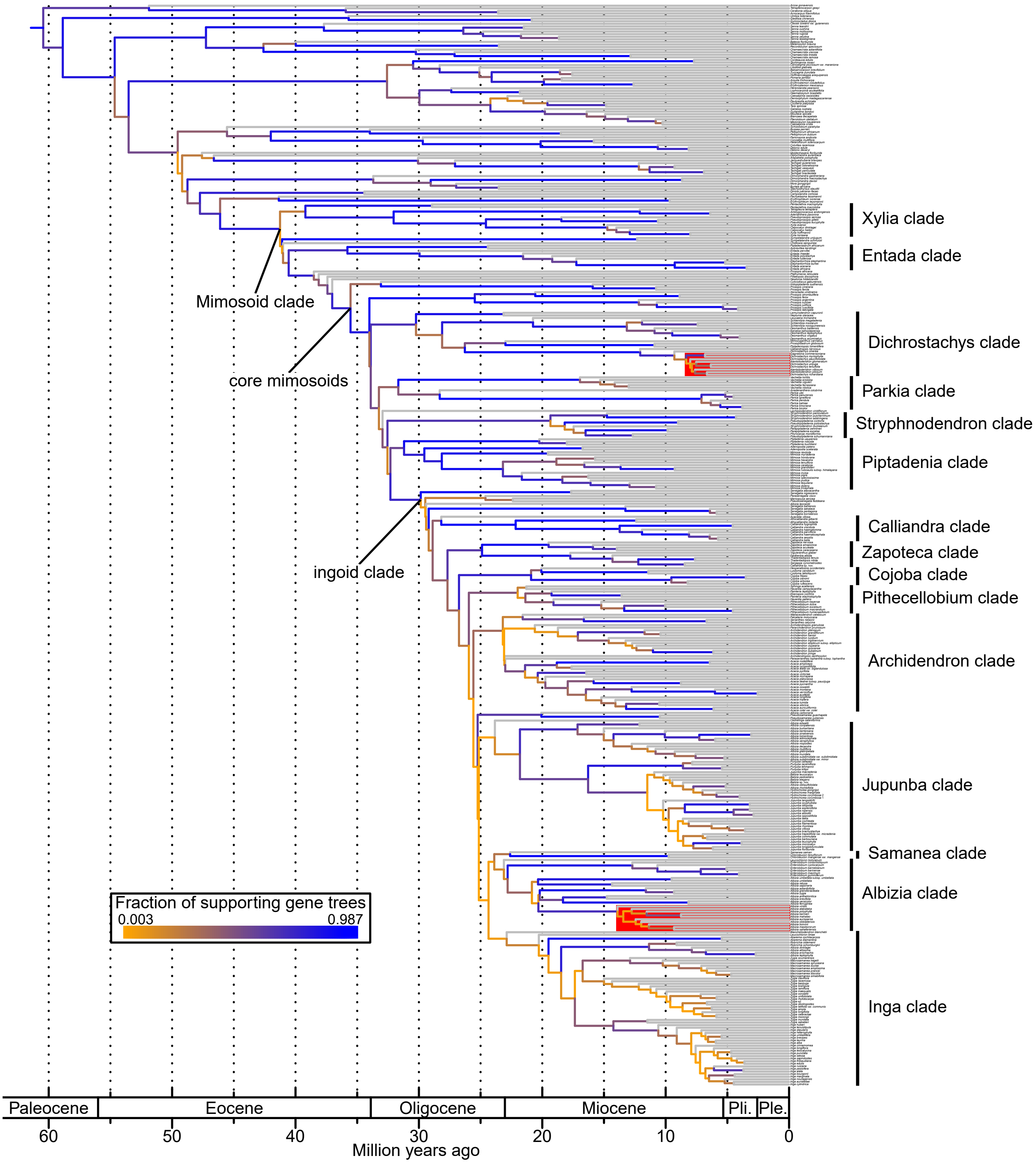

**Figure S31.** Gene tree incongruence mapped onto the time-calibrated version of the phylogenomic backbone of Caesalpinioideae. Each branch is coloured to reflect the ratio of total supporting versus total conflicting gene trees as determined by PhyParts. Clades named by Koenen et al. (*4*) are labelled. Two recent radiations in Madagascar, one in the Dichrostachys clade and one in Albizia, are highlighted.

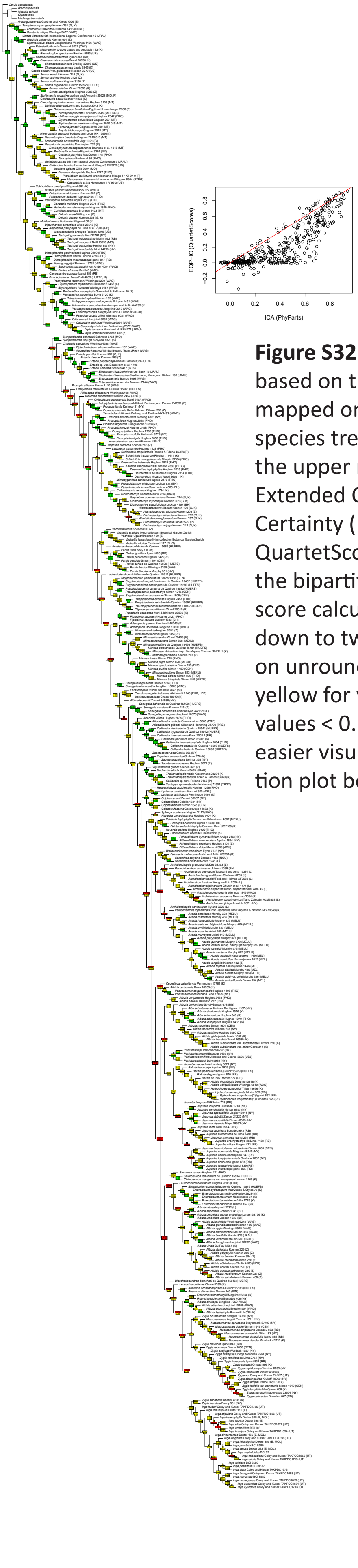

**Figure S32 (left).** Internode certainty values based on the 821 single-copy gene trees mapped onto the single-copy genes ASTRAL species tree (Figure S14). For each node, the upper number shows the quartet-based Extended Quadripartition Internode Certainty (EQP-IC) score calculated with QuartetScores, and the lower number shows the bipartition-based Internode Certainty All score calculated with PhyParts, both rounded down to two digits. Boxes are coloured based on unrounded values: green for values ≥ 0.5, yellow for values ≥ 0 and < 0.5, and red for values < 0. Branch lengths are set equal for easier visualisation. Inset depicts a correlation plot between the two measures.

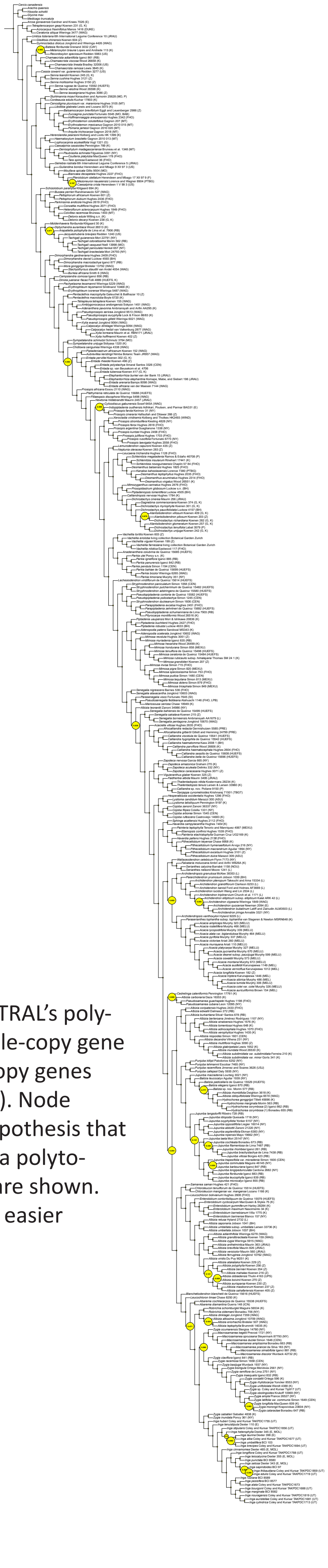

**Figure S33 (right).** Results of ASTRAL's polytomy test based on the 821 single-copy gene trees mapped onto the single-copy genes ASTRAL species tree (Figure S14). Node numbers are tests of the null hypothesis that a branch should be replaced by a polytomy. Only node numbers > 0.05 are shown. Branch lengths are set equal for easier visualisation.

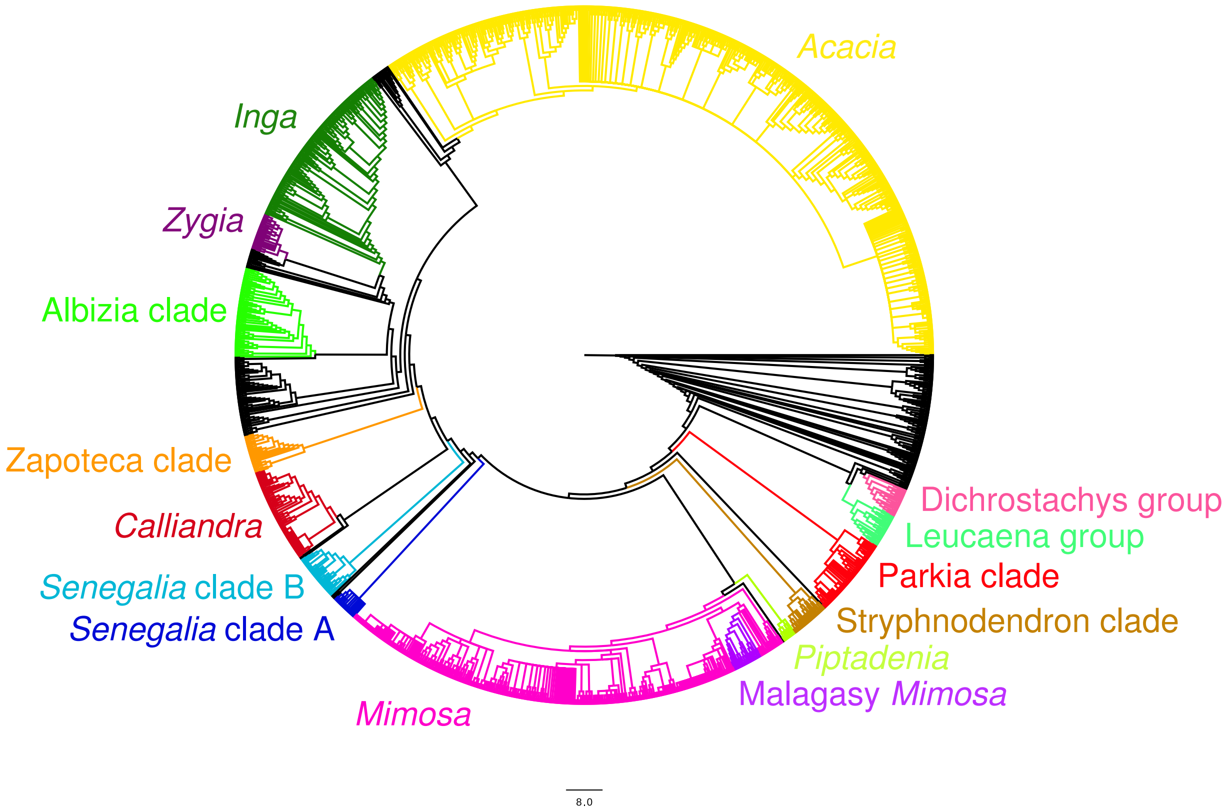

**Figure S34.** Metachronogram with the names and locations of all subtrees that were grafted onto the phylogenomic backbone.

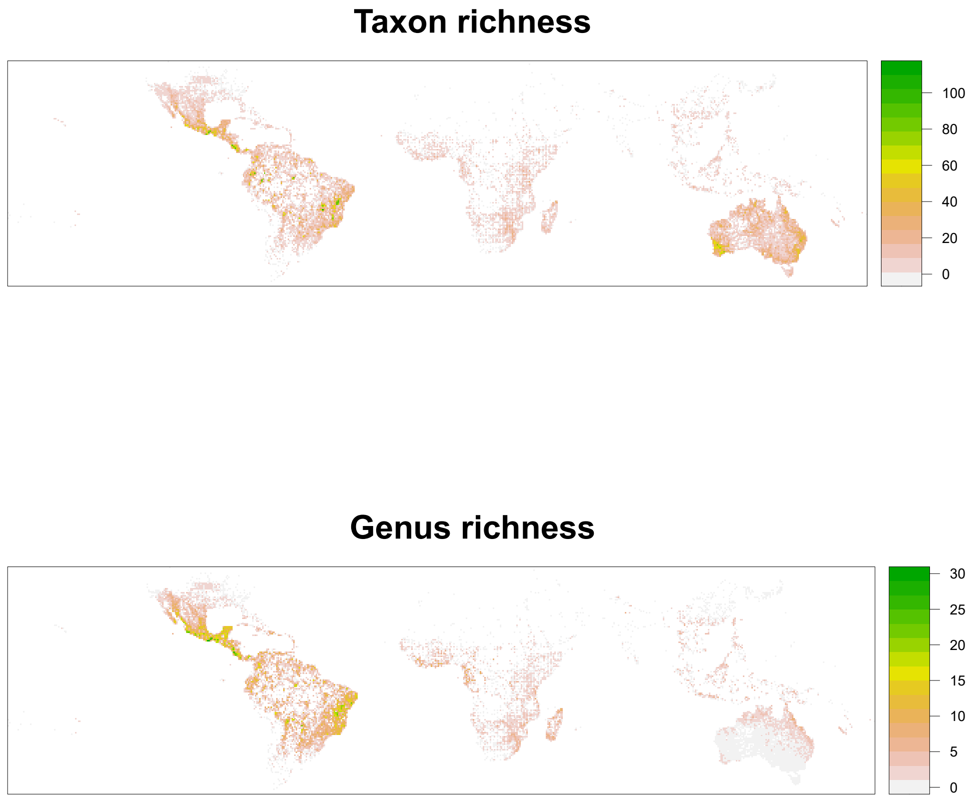

**Figure S35.** Mimosoid taxon and genus richness per half degree latitude/longitude grid cell.

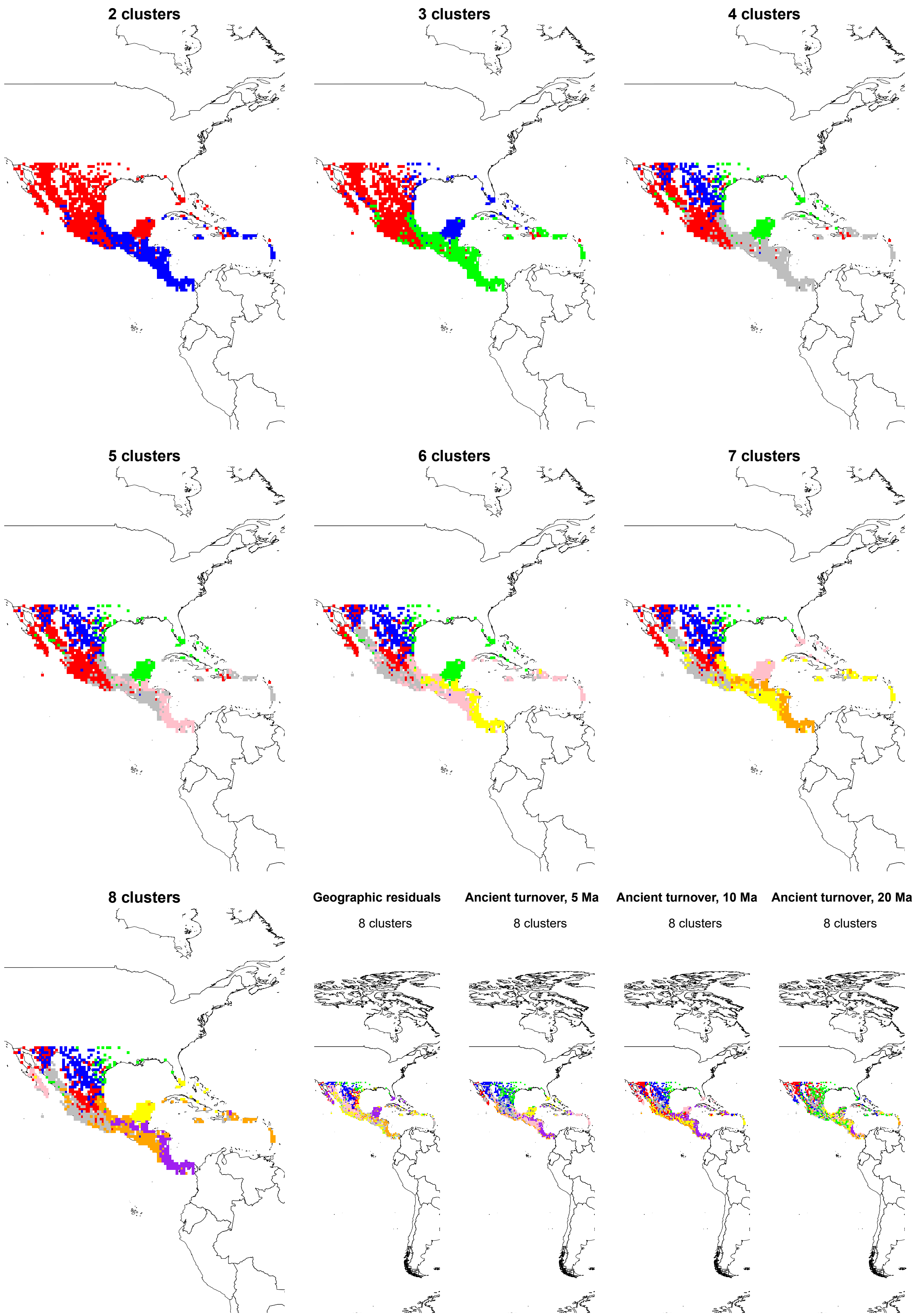

**Figure S36.** Phyloregionalization of North America using the metachronogram. Subfigures show clustering results with two to eight phyloregions, as well as the results of phyloregionalization analyses using the geographic residuals of phylogenetic turnover, and ancient phylogenetic turnover with a cut-off of 5, 10, and 20 million years.

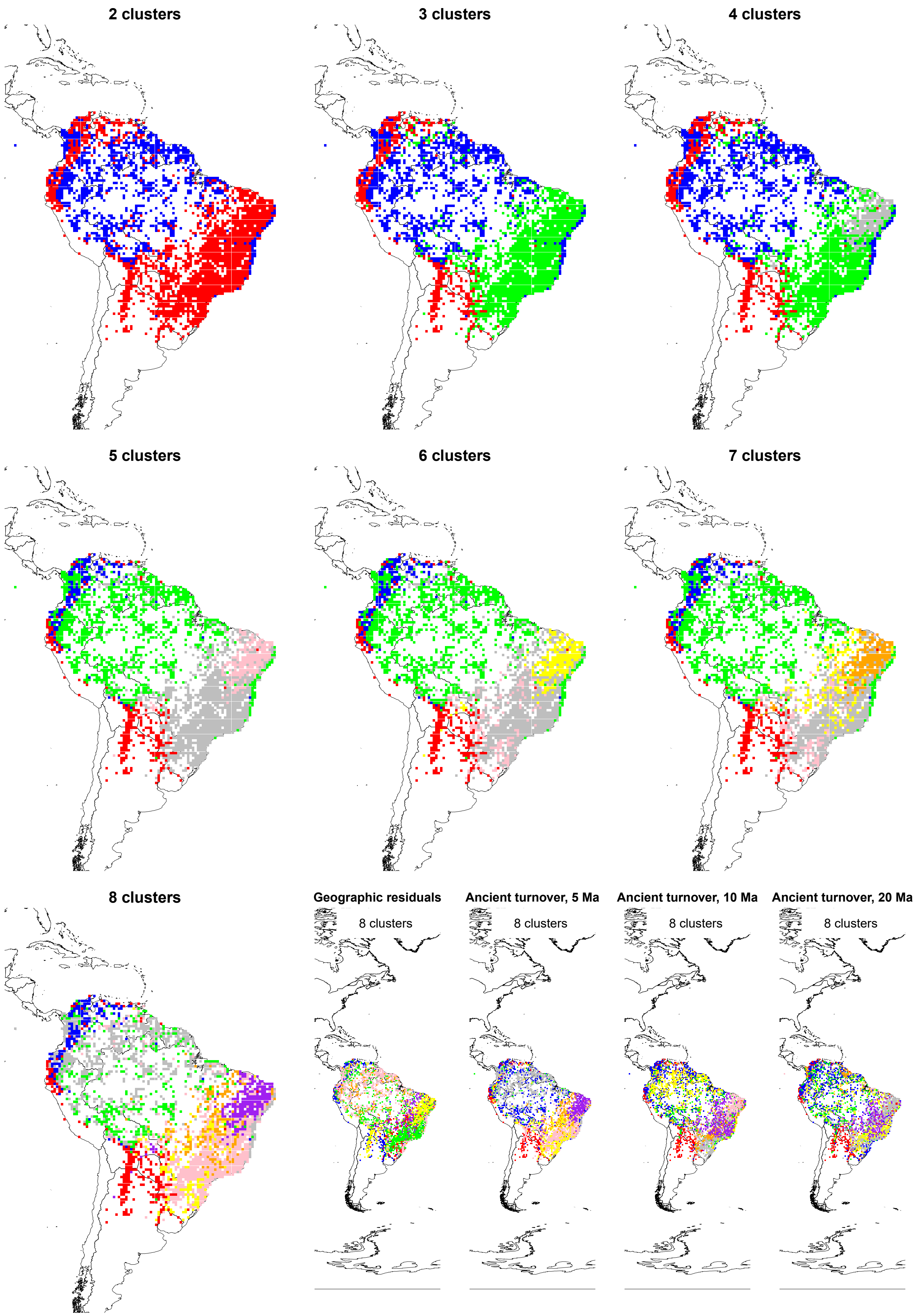

**Figure S37.** Phyloregionalization of South America using the metachronogram. Subfigures show clustering results with two to eight phyloregions, as well as the results of phyloregionalization analyses using the geographic residuals of phylogenetic turnover, and ancient phylogenetic turnover with a cut-off of 5, 10, and 20 million years.

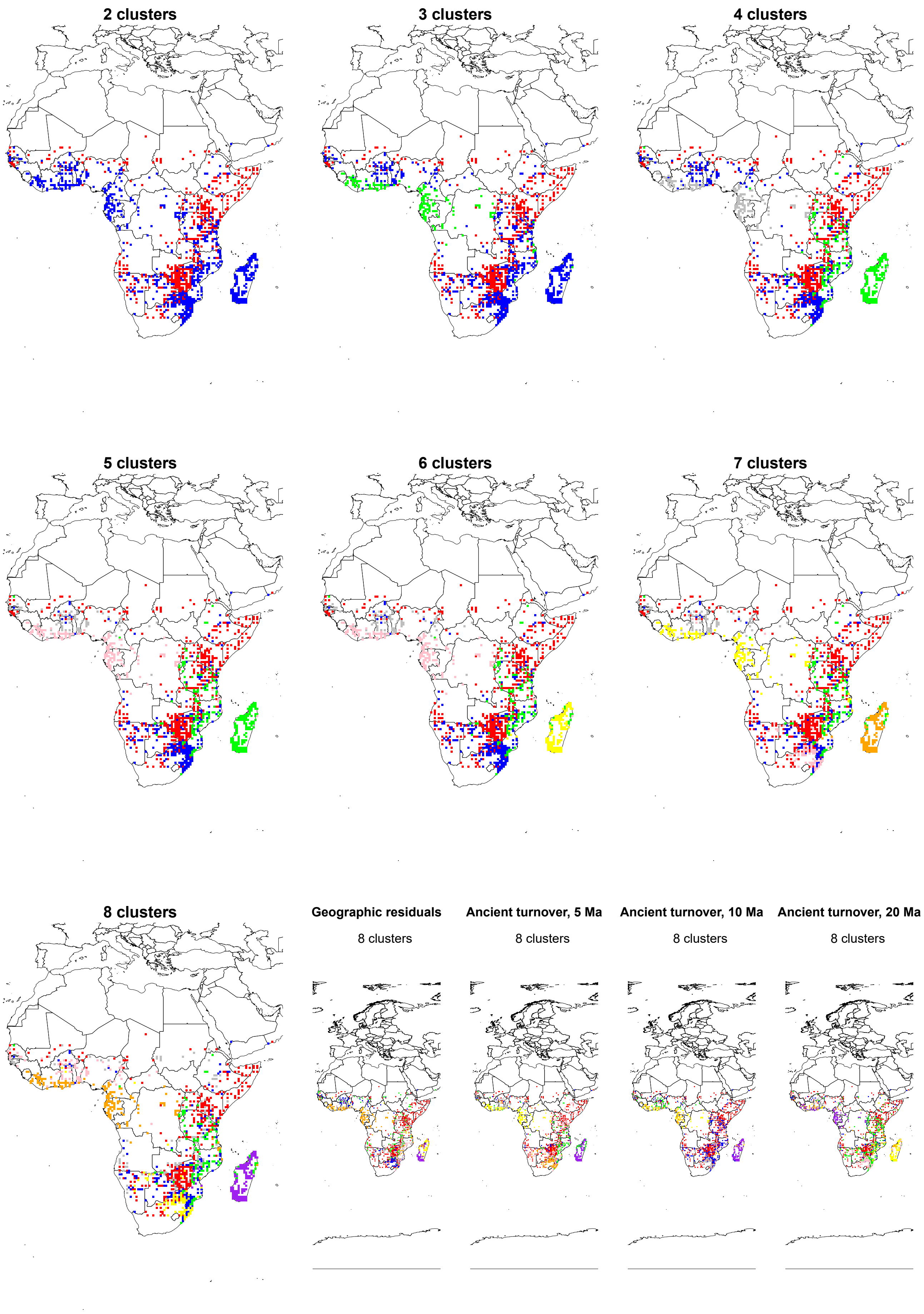

**Figure S38.** Phyloregionalization of Africa using the metachronogram. Subfigures show clustering results with two to eight phyloregions, as well as the results of phyloregionalization analyses using the geographic residuals of phylogenetic turnover, and ancient phylogenetic turnover with a cut-off of 5, 10, and 20 million years.

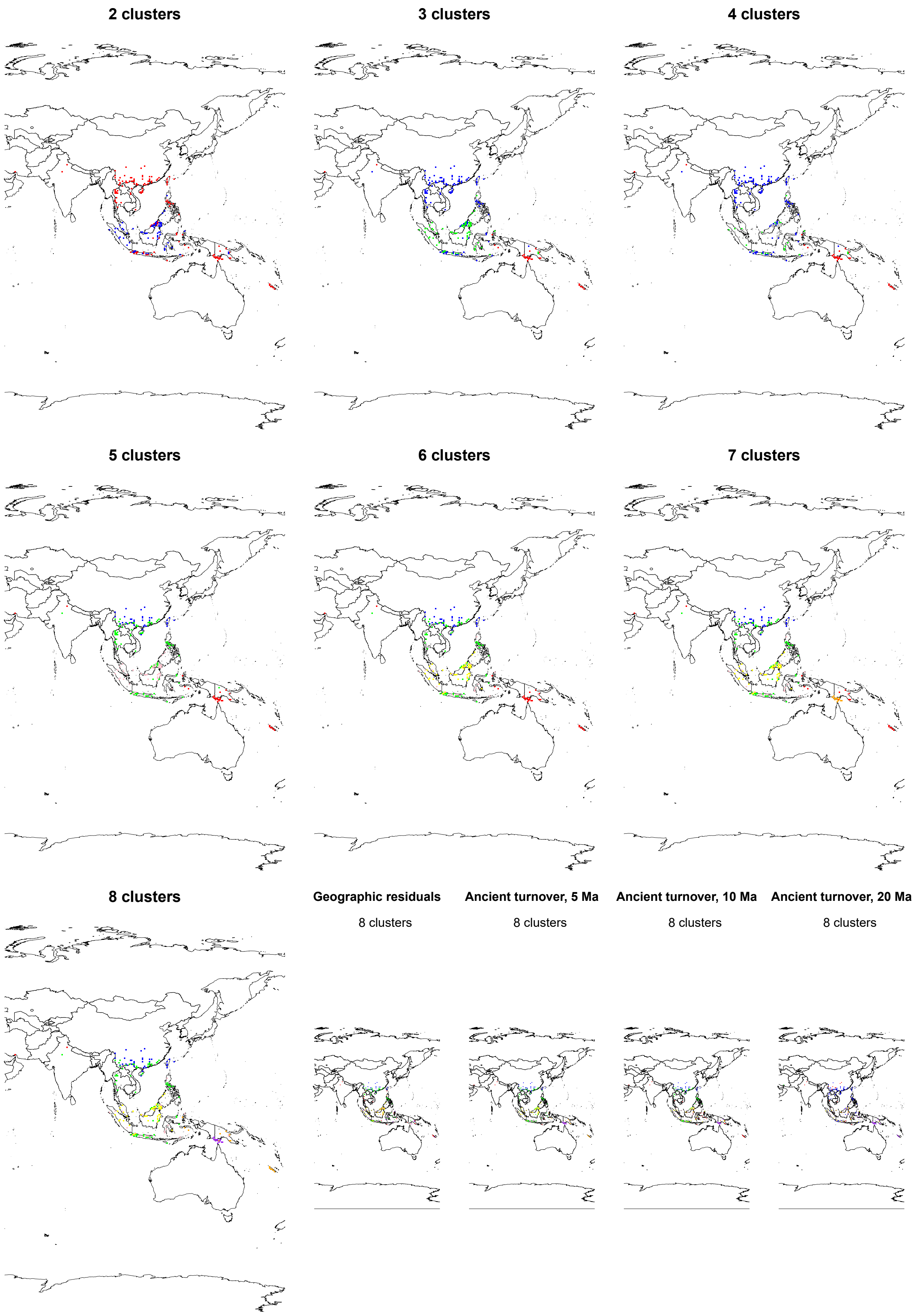

**Figure S39.** Phyloregionalization of Asia using the metachronogram. Subfigures show clustering results with two to eight phyloregions, as well as the results of phyloregionalization analyses using the geographic residuals of tphylogenetic turnover, and ancient phylogenetic turnover with a cut-off of 5, 10, and 20 million years.

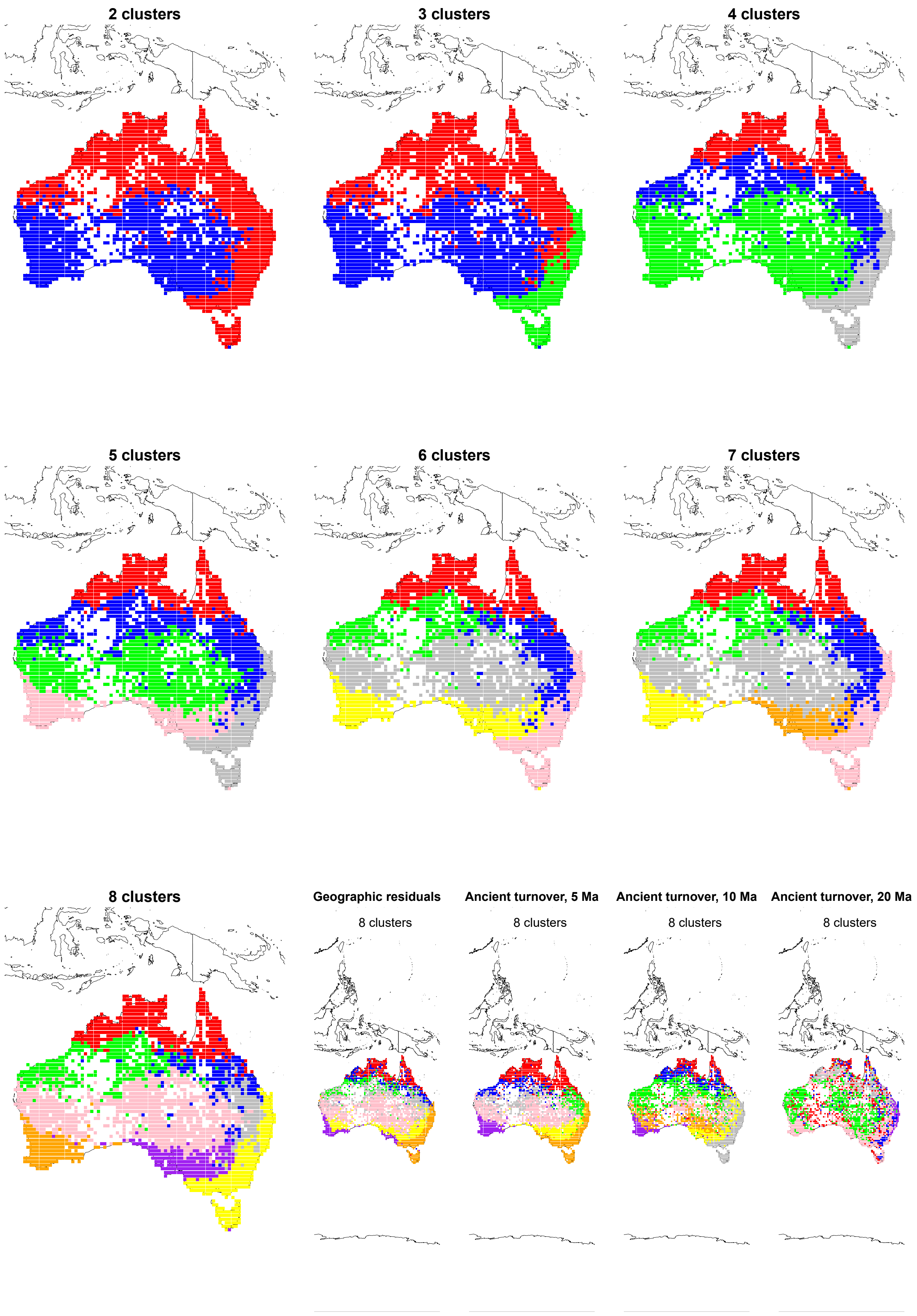

**Figure S40.** Phyloregionalization of Australia using the metachronogram. Subfigures show clustering results with two to eight phyloregions, as well as the results of phyloregionalization analyses using the geographic residuals of phylogenetic turnover, and ancient phylogenetic turnover with a cut-off of 5, 10, and 20 million years.

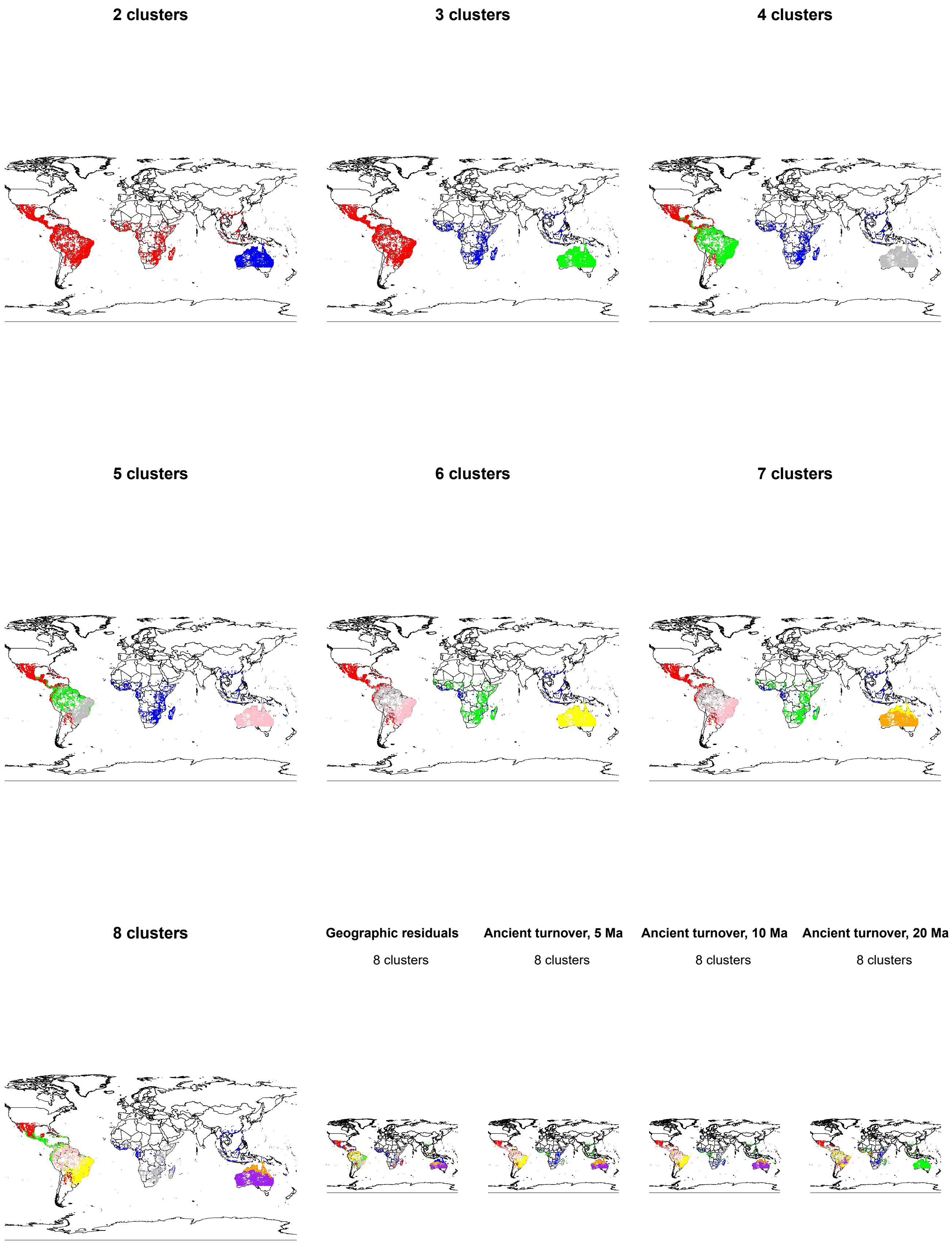

**Figure S41.** Phyloregionalization of the global tropics using the metachronogram. Subfigures show clustering results with two to eight phyloregions, as well as the results of phyloregionalization analyses using the geographic residuals of phylogenetic turnover, and ancient phylogenetic turnover with a cut-off of 5, 10, and 20 million years.

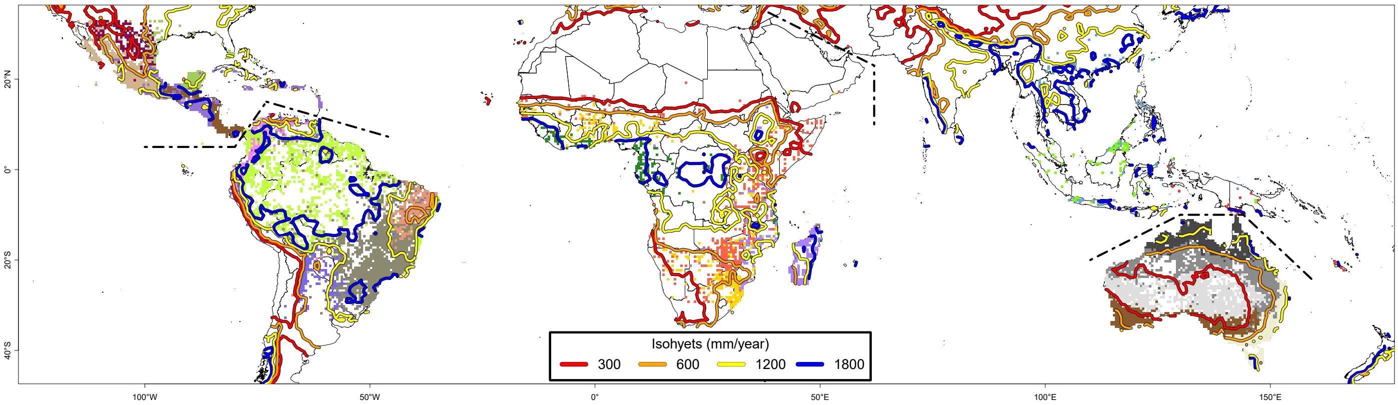

**Figure S42.** Phyloregionalization per continent using the metachronogram, showing global distribution of isohyets. Caption otherwise as for Figure 3.

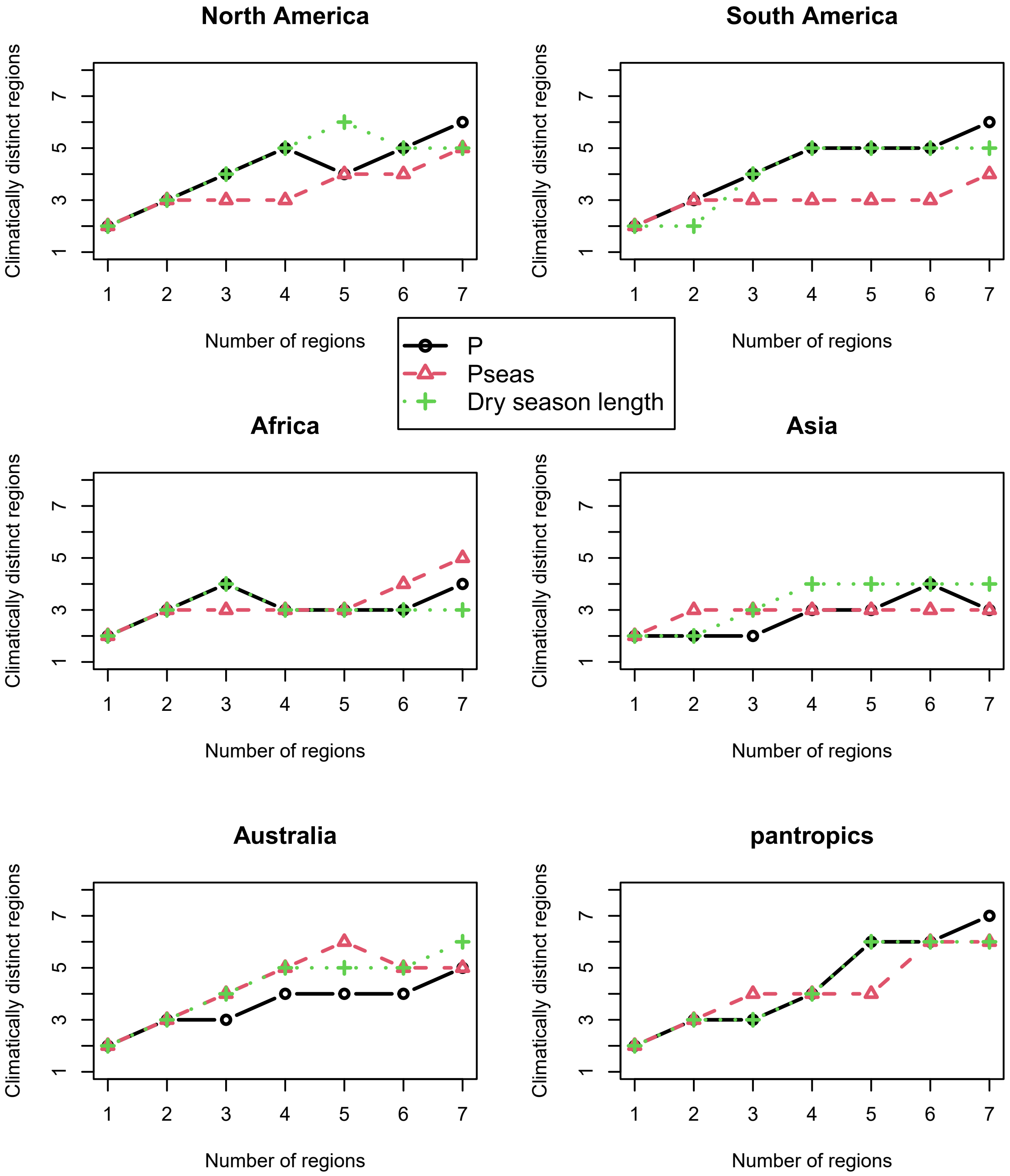

**Figure S43.** Climatic distinctiveness of phyloregions. Each subplot shows, for phyloregionalization analyses with two to eight phyloregions performed using the metachronogram, how many of the resulting phyloregions have statistically significant different climatic values from the other regions based on mean annual precipitation (P), precipitation seasonality (Pseas), and dry season length.

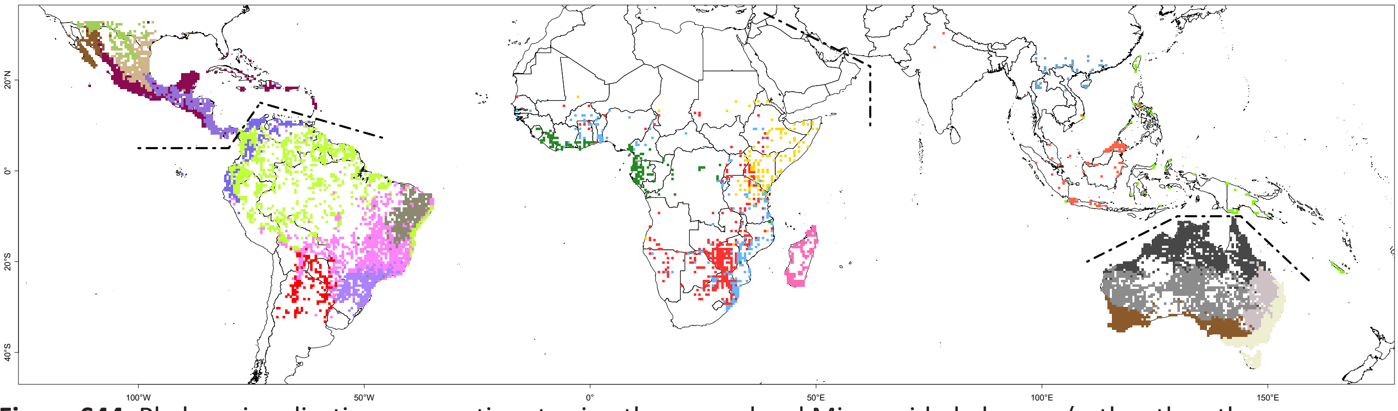

**Figure S44.** Phyloregionalization per continent using the genus-level Mimosoid phylogeny (rather than the metachronogram). Caption otherwise as for Figure 3.

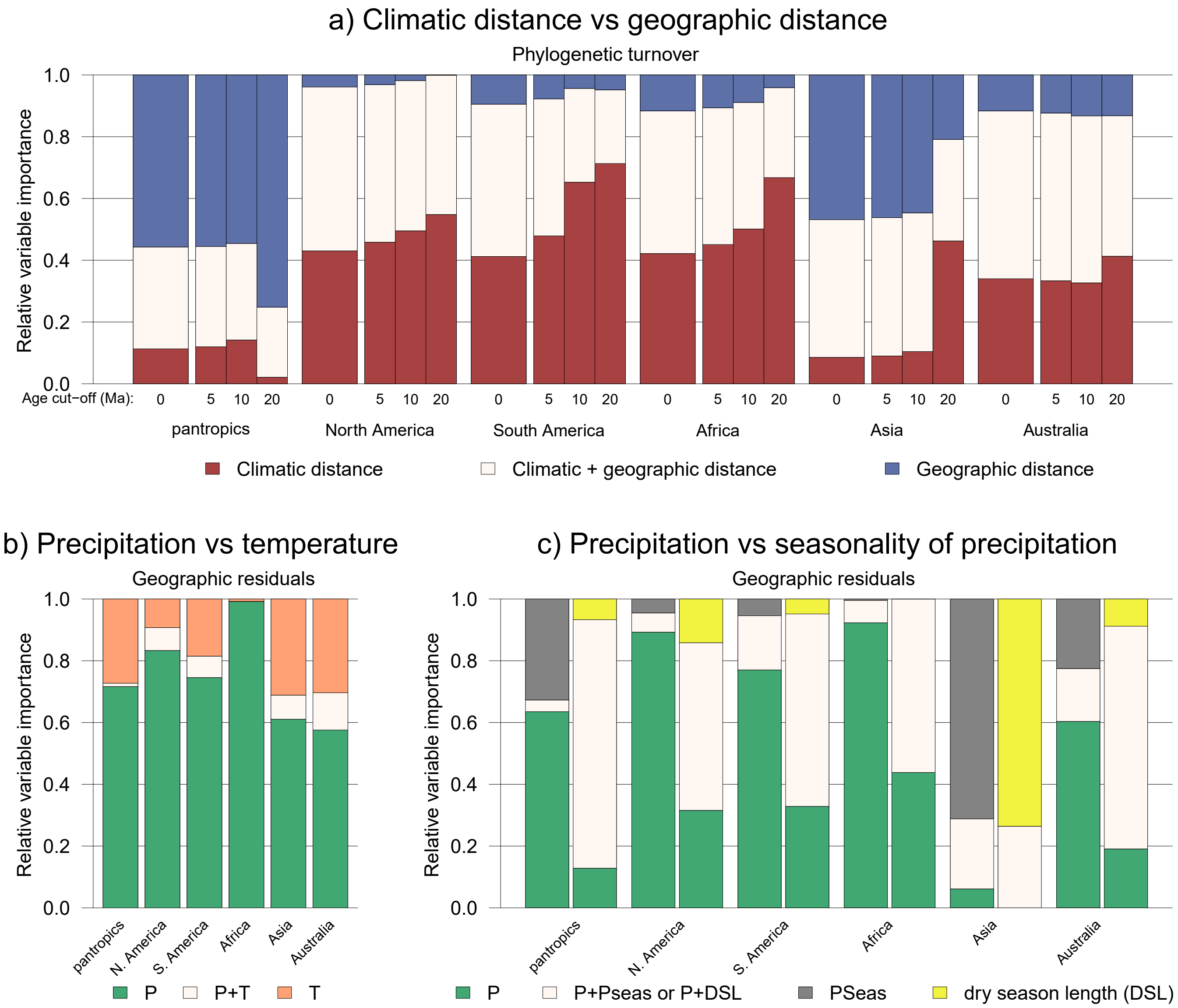

**Figure S45.** Variation partitioning results obtained using the genus-level Mimosoid phylogeny (rather than the metachronogram). See caption Figure 2 for explanation.

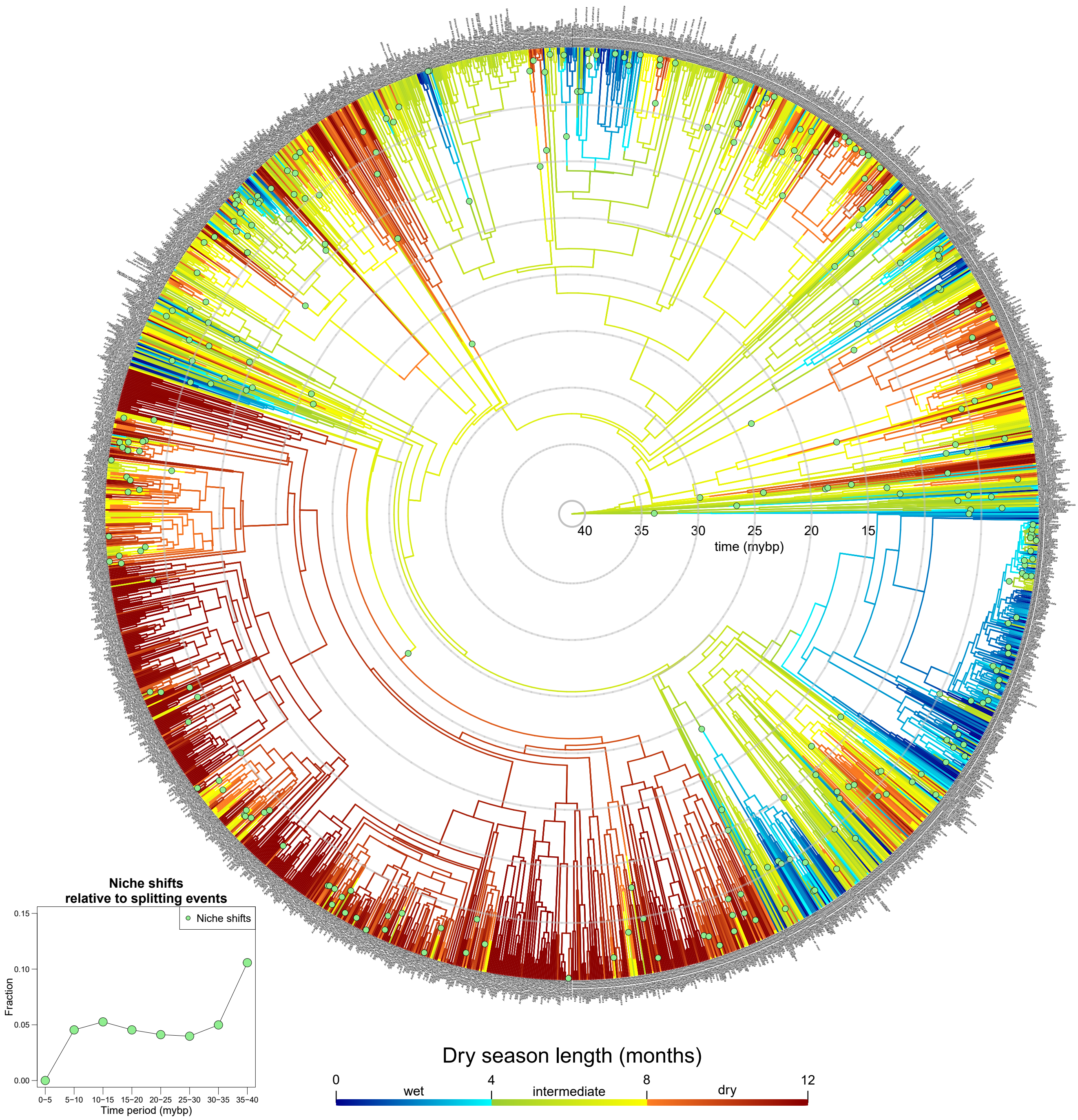

**Figure S46.** Optimisation of dry season length across the Mimosoid phylogeny. Inset shows the fraction of dry season length niche shifts per speciation event through time. See caption Figure 1 for explanation.

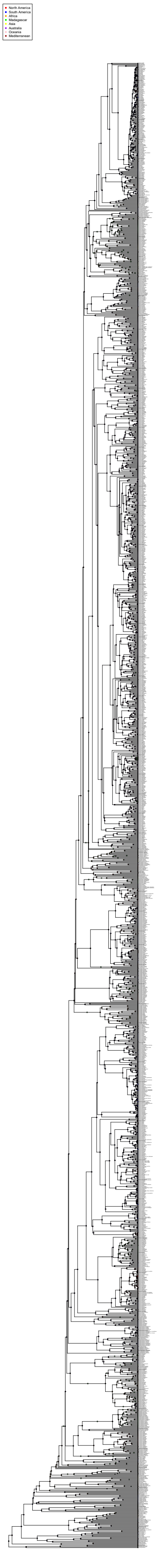

**Figure S47.** Ancestral range estimation of Caesalpinioideae, performed using BioGeoBEARS with the best-fitting model (i.e., DEC+J). Trans-oceanic dispersal events in the Mimosoid clade, based on a model with seven regions, are indicated with numbered green circles.

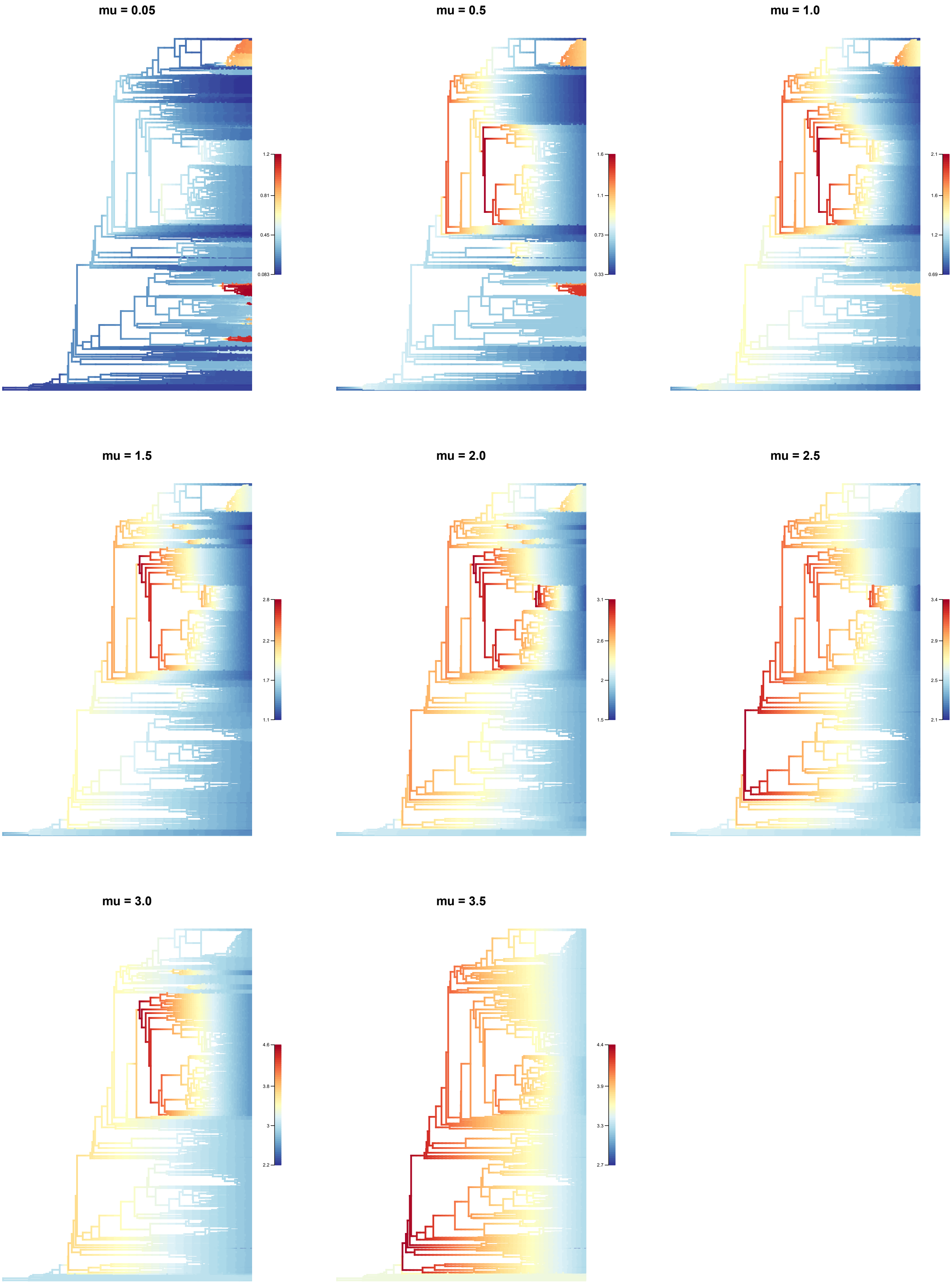

**Figure S48.** Speciation rates estimated across the Caesalpinioideae metachronogram under eight scenarios with different fixed extinction rates. Extinction rates are shown above each subfigure, while speciation rates are indicated by branch colours.

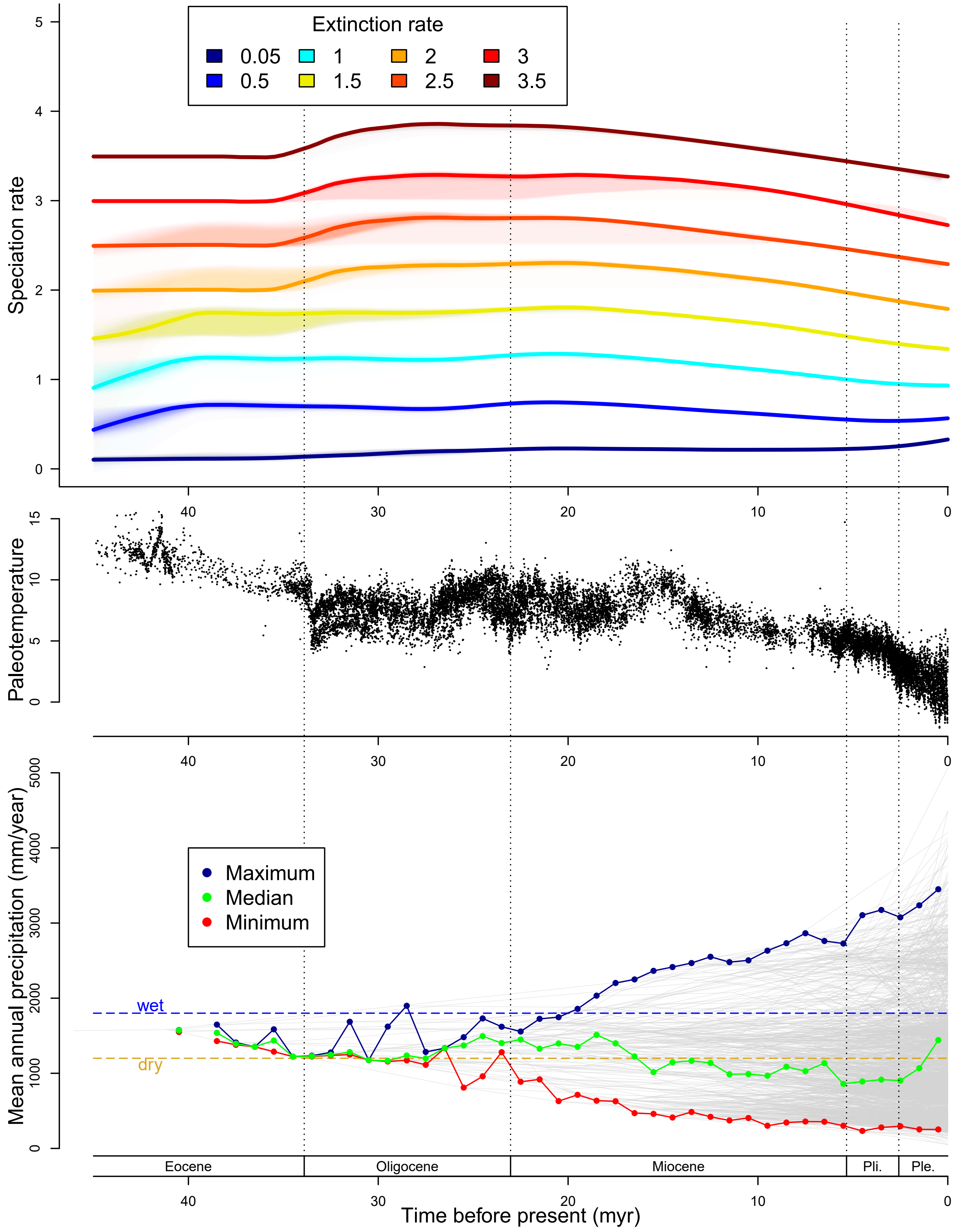

**Figure S49.** Top: Speciation rates in the Mimosoid clade through time, estimated under different extinction rate scenarios using BAMM. Middle: Paleotemperature inferred from delta O18 measurements, using data from Zachos et al. (*125*). Bottom: Phenogram of mean annual precipitation in the Mimosoid clade through time. Coloured lines with dots show the median, wettest, and driest reconstructed rainfall niche of all nodes in the phylogeny per time bin of one million years.

### Appendix 1

#### Information about the Mimosoid checklist and occurrence dataset

This appendix lists all the references used for building the Mimosoid checklist and assembling and quality controlling the occurrence dataset, as well as the sources of occurrence data, the GBIF DOIs, and any additional notes. The appendix is organised in the same way as the occurrence dataset was assembled, i.e. mainly per genus, but sometimes per clade, monograph, or geographic region. For example, the first entry reflects all the Neotropical genera discussed by Barneby & Grimes (1996, 1997) and Barneby (1998).

Notes for the occurrence dataset of the non-Mimosoid Caesalpinioideae are provided at the end of this appendix.

---

**_Abarema_, _Albizia_ (New World), _Balizia_, _Blanchetiodendron_, _Calliandra_, _Cedrelinga_, _Chloroleucon_, _Cojoba_, _Ebenopsis_, _Enterolobium_, _Falcataria_, _Havardia_, _Hesperalbizia_, _Hydrochorea_, _Jupunba_, _Leucochloron_, _Lysiloma_, _Macrosamanea_, _Painteria_, _Paraserianthes_, _Pithecellobium_, _Pseudosamanea_, _Punjuba_, _Robrichia_, _Samanea_, _Spinga_ and _Zygia_**

**References used for taxonomy and distribution**

- Barneby, RC & Grimes, JW (1996) Silk Tree, Guanacaste, Monkey's Earring: A Generic System for the Synandrous Mimosaceae of the Americas. Part I. *Abarema*, *Albizia*, and Allies. Memoirs of the New York Botanical Garden vol. 74, Bronx, New York.
- Barneby, RC & Grimes, JW (1997) Silk Tree, Guanacaste, Monkey's Earring: A Generic System for the Synandrous Mimosaceae of the Americas. Part 2. *Pithecellobium*, *Cojoba*, and *Zygia*. Memoirs of the New York Botanical Garden vol. 74, Bronx, New York.
- Barneby, RC (1998) Silk Tree, Guanacaste, Monkey's Earring: A Generic System for the Synandrous Mimosaceae of the Americas. Part 3. *Calliandra*. Memoirs of the New York Botanical Garden vol. 74, Bronx, New York.
- Bässler, M (1998) Flora de la República de Cuba. Seria A – Plantas Vasculares. Fascículo 2: Mimosaceae. Koeltz Scientific Books, Koenigstein, Germany.
- Brown, GK, Murphy, DJ & Ladiges, PY (2011) Relationships of the Australo-Malesian genus *Paraserianthes* (Mimosoideae: Leguminosae) identifies the sister group of *Acacia sensu stricto* and two biogeographical tracks. Cladistics 27: 380-390.
- Da Silva e Silva, WL, Morim, MP, Iganci, JRV & Moreira Dos Santos, JU (2016) New combination in *Macrosamanea* (Leguminosae-Mimosoideae). Phytotaxa 288(2): 187-192.
- Forero, E (2009) Estudios in leguminosas Colombianas II. Universidad Nacional de Colombia, Instituto de Ciencias Naturales, Facultad de Ciencas Bogota, Colombia.
- Gale, SW & Pennington, TD (2004) *Lysiloma* (Leguminosae: Mimosoideae) in Mesoamerica. Kew Bulletin 59: 453-467.
- García, R & Peguero, B (2005) *Cojoba urbanii* (Alain) R. García & B. Peguero (Mimosaceae),

nueva combinación. Moscosoa 14: 6-9.
- Guerra, E, Morim, MP & Iganci, JRV (2016) A new species of *Abarema* (Fabaceae) from Brazil. Phytotaxa 289(1): 77-82.
- Hammel, BE, Grayum, MH, Herrera, C & Zamora, N (eds.) (2010) Manual de Plantas de Costa Rica. Volumen V: Dicotiledóneas (Clusiaceae – Gunneraceae). Monographs in Systematic Botany from the Missouri Botanical Garden, Volume 119, St. Louis, Missouri.
- Hernández, HM (2008) *Calliandra dolichopoda* and *C. cualensis* (Leguminosae, Mimosoideae), two new species from Mexico. Brittonia 60(3): 245-251.
- Hernández, HM, de Stefano, RD, Gutiérrez, C, Carnevali-Fernández-Concha, G, Can, LL & Pool, E (2014) *Calliandra ricoana* (Leguminosae, Mimosoideae, Ingeae): A new and critically endangered species from Chiapas, Mexico. Botanical Sciences 92(2): 177-181.
- Hernández, HM & Gómez-Hinostrosa, C (2017) *Calliandra mayana* (Leguminosae, Mimosoideae), a new narrowly endemic species from Campeche, Mexico. Phytotaxa 307(4): 278-284.
- Hernández, HM & Gómez-Hinostrosa, C (2019) A narrowly endemic species of *Calliandra* series *Racemosae* (Fabaceae) from Sinaloa, Mexico. Phytotaxa 401(1): 49-54.
- Hughes, CE & Atahuachi, B (2006) A new species of *Leucochloron* (Leguminosae: Mimosoideae) endemic to Bolivia. Kew Bulletin 61(4): 559-563.
- Iganci, JRV & Morim, MP (2009) Three new species of Abarema (Leguminosae, Mimosoideae) from south-eastern Brazil. Kew Bulletin 64: 271-277.
- Iganci, JRV & Morim, MP (2012) *Abarema* (Fabaceae, Mimosoideae) in the Atlantic Domain, Brazil. Botanical Journal of the Linnean Society 168(4): 473-486.
- Linares, J (2005) Especie nueva de *Albizia* (Leguminosae: Mimosoideae) de Centroamérica. Revista Mexicana de Biodiversidad 76(1): 7-10.
- Nielsen, I, Guinet, Ph. & Baretta-Kuipers, T (1983) Studies in the Malesian, Australian and Pacific Ingeae (Leguminosae – Mimosoideae): the genera *Archidendropsis*, *Wallaceodendron*, *Paraserianthes*, *Pararchidendron* and *Serianthes*. Adansonia 5: 303-360.
- Mesquita, AdL (1993) *Enterolobium glaziovii* (Benth.) Mesquita, comb. nov. et "status novum" para as regiões Sudeste e Nordeste do Brasil. Acta Botanica Brasilica 7(2): 17-23.
- Ortiz-Rodriguez, AE, Hernández, HM & Perez-Farrera, MA (2015) *Calliandra bifoliolata* (Leguminosae, Mimosoideae), a new species from Chiapas, Mexico, with notes on *C. brenesii*, *C. grandifolia*, and *C. laevis*. Brittonia 67(3): 175-179.
- de Queiroz, LP (2009) Leguminosas da Caatinga. Universidade Estadual de Feira de Santana.
- Rico Arce, MdL (1999) New Combinations in Mimosaceae. Novon 9: 554-556.
- Rico Arce, MdL, Gale, SL & Maxted, N (2008) A taxonomic study of *Albizia* (Leguminosae: Mimosoideae: Ingeae) in Mexico and Central America. Anales del Jardín Botánico de Madrid 65(2): 255-305.
- Silverstone-Sopkin, P (2015) A new species of *Chloroleucon* (Leguminosae, Mimosoideae) from the Cauca Valley, Colombia. Novon 24(1): 50-54.
- Soares, MVB, Guerra, E, Morim, MP & Iganci, JRV (2021) Reinstatement and recircumscription of *Jupunba* and *Punjuba* (Fabaceae) based on phylogenetic evidence. Botanical Journal of the Linnean Society.
- de Souza, ÉR & Queiroz, LP (2004) Duas novas espécies de *Calliandra* Benth. (Leguminosae – Mimosoideae) da Chapada Diamantina, Bahia, Brasil. Revista Brasil. Bot. 27(4): 615-619.
- de Souza, ÉR (2010) *Calliandra paganuccii* (Leguminosae – Mimosoideae), a new species from the Chapada Diamantina, Bahil, Brazil. Neodiversity 5: 7-10.

- de Souza, ÉR, Lewis, GP, Forest, F, Schnadelbach, AS, van den Berg, C & Queiroz, LP (2013) Phylogeny of *Calliandra* (Leguminosae: Mimosoideae) based on nuclear and plastic molecular markers. Taxon 62(6): 1200-1219.
- de Souza, ÉR, Ferreira Lima, AV, Ribeiro dos Santos, FdA & Queiroz, LP (2014) Three new species of *Calliandra* in section *Monticola* (Leguminosae, Mimosoideae) from Chapada Diamantina, Bahia, Brazil. Phytotaxa 164(2): 104-114.
- Souza, ÉR de, Monteiro Luz, AR, Rocha, L & Lewis, GP (2022) Molecular and Morphological Analysis Supports the Separation of *Robrichia* as a Genus Distinct from *Enterolobium* (Leguminosae: Caesalpinioideae: Mimosoid Clade). Systematic Botany 47(1): 268-277.
- Stahl, B, Rico-Arce, MdL & Lewis, GP (2010) *Zygia nubigena* sp. nov. (Leguminosae-Mimosoideae) from a submontane cloud forest in western Ecuador. Nordic Journal of Botany 28: 453-456.
- Turner, BL (2000) The Texas species of *Calliandra* (Leguminosae, Mimosoideae). Lundellia 3: 13-18.

**Sources of occurrence data**

GBIF, DryFlor, SEINet, Barneby & Grimes 1996.

GBIF DOIs: https://doi.org/10.15468/dl.qovicd, https://doi.org/10.15468/dl.oivieb, https://doi.org/10.15468/dl.v2r37r, and https://doi.org/10.15468/dl.ic5opx.

**Notes**

After the recircumscription of Soares *et al*. 2021, *Abarema* only consists of two species (*A. cochliacarpos* and *A. diamantina*), while the remaining species in this genus were transferred to *Punjuba* and *Jupunba*. However, the generic affinities of five other *Abarema* species are still poorly known: *A. maestrensis*, *A. levelii*, *A. acreana*, *A. ricoae*, and *A. agropecuaria* (the latter two were suggested, but not formally published, by Barneby & Grimes 1996). These five species are unlikely to belong to *Abarema* s.s. and will be transferred in the near future (J. Iganci, pers. comm.). Until then, they are kept as *Abarema* in our taxonomic checklist and occurrence dataset.

This section of the dataset only contains *Albizia* species native to the New World.

*Albizia carbonaria*: following Barneby & Grimes 1996, all occurrences outside Colombia, Venezuela, and Panama are considered introduced, and were removed from the dataset.

*Pithecellobium dulce*: following Barneby & Grimes 1997, occurrences in the Caribbean, Florida, and eastern Brazil are considered introduced, and were removed.

*Samanea saman*: following Barneby & Grimes 1996, removed outside Venezuela, Colombia, and Central America.

Following Rico Arce 1999 and Iganci *et al*. 2016, *Hydrochorea acreana* is included in the dataset as *Abarema acreana.* However, *Albizia pedicellaris* and *A. elegans* are included as *Balizia pedicellaris* and *B. elegans*, respectively, following Barneby & Grimes 1996.

The varieties of *Zygia coccinea* are only included at the species level, not separately.

*Chloroleucon acacioides* does not occur in the Caatinga according to de Queiroz 2009, and its occurrences there were removed.

---

##### *Acacia*

**References used for taxonomy and distribution**

- Bell, SAJ & Driscoll, C (2017) *Acacia wollarensis* (Fabaceae, Mimosoideae sect. *Botrycephalae*), a distinctive new species endemic to the Hunter Valley of New South Wales, Australia. Telopea 20: 125-136.
- Bull, JP, Dillon, SJ & Brearley, DR (2019) *Acacia corusca* (Fabaceae: Mimosoideae), a new species from the Pilbara bioregion in north-western Australia. Nuytsia 30: 19-22.
- Cuff, NJ & Cowie, ID (2017) *Acacia nicholsonensis* (Fabaceae), a new 'Minni Ritchi'-barked species of *Acacia* sect. *Juliflorae* from the Gulf of Carpentaria region of Northern Australia. Nuytsia 28: 147-158.
- Du Puy, DJ, Labat, J-N, Rabevohitra, R, Villiers, J-F, Bosser, J & Moat, J (2002) The Leguminosae of Madagascar. Kew Publishing.
- González-Orozco, CE, Laffan, SW, Knerr, N & Miller, JT (2013) A biogeographical regionalization of Australian *Acacia* species. Journal of Biogeography 40(11): 2156-2166.
- Kodela, PG (2015) *Acacia yalwarensis* (Fabaceae, Mimosoideae sect. *Botrycephalae*), a new species from the South Coast of New South Wales, Australia. Telopea 18: 27-31.
- Le Roux, JJ, Strasberg, D, Rouget, M, Morden, CW, Koordom, M & Richardson, DM (2014) Relatedness defies biogeography: the tale of two island endemics (*Acacia heterophylla* and *A. koa*). New Phytologist 204: 230-242.
- Lewington, MA & Maslin, BR (2009) Three new species of *Acacia* (Leguminosae: Mimosoideae) from the Kimberley Region, Western Australia. Nuytsia 19(1): 63-75.
- Maslin, BR (2013) *Acacia gibsonii*, a distinctive, rare new species of *Acacia* sect. *Juliflorae* (Fabaceae: Mimosoideae) from south-west Western Australia. Nuytsia 23: 277-281.
- Maslin, BR (2014a) Miscellaneous new species of *Acacia* (Fabaceae: Mimosoideae) from south-west Western Australia. Nuytsia 24: 139-159.
- Maslin, BR (2014b) Two new species of *Acacia* (Fabaceae: Mimosoideae) with conservation significance from Banded Iron Formation ranges in the vicinity of Koolyanobbing, Western Australia. Nuytsia 24: 131-138.
- Maslin, BR (2014c) Four new species of *Acacia* (Fabaceae: Mimosoideae) with fasciculate phyllodes from south-west Western Australia. Nuytsia 24: 161-175.
- Maslin, BR (2014d) Four new species of *Acacia* section *Juliflorae* (Fabaceae: Mimosoideae) from the arid zone in Western Australia. Nuytsia 24: 193-205.
- Maslin, BR (2015) Synoptic overview of *Acacia* sensu lato (Leguminosae: Mimosoideae) in East and Southeast Asia. Gardens' Bulletin Singapore 67(1): 231-250.
- Maslin, BR & Buscumb, C (2007a) Two new species of *Acacia* (Leguminosae: Mimosoideae) from the Koolanooka Hills in the northern wheatbelt region of south-west Western Australia. Nuytsia 17: 253-262.
- Maslin, BR & Buscumb, C (2007b) Two new *Acacia* species (Leguminosae: Mimosoideae) from banded ironstone ranges in the Midwest region of south-west Western Australia. Nuytsia 17: 263-272.
- Maslin, BR & Murphy, DJ (2009) A taxonomic revision of *Acacia verniciflua* and *A. leprosa* (Leguminosae: Mimosoideae) in Australia. Muelleria 27(2): 183-223.
- Maslin, BR & Reid, JE (2012a) A taxonomic revision of Mulga (*Acacia aneura* and its

close relatives: Fabaceae) in Western Australia. Nuytsia 22(4): 129-267.
- Maslin, BR & Reid, JE (2012b) *Acacia bartlei* (Fabaceae: Mimosoideae), a new species from near Esperance, Western Australia. Nuytsia 22(2): 51-56.
- Maslin, BR, Barrett, MD & Barrett, RL (2013) A baker's dozen of new wattles highlights significant *Acacia* (Fabaceae: Mimosoideae) diversity and endemism in the north-west Kimberley region of Western Australia. Nuytsia 23: 543-587.
- Maslin, BR & Barrett, RL (2014) *Acacia mackenziei*, a new species of *Acacia* section *Lycopodiifoliae* (Fabaceae: Mimosoideae) with conservation significance from the east Kimberley region in northern Western Australia. Nuytsia 24: 187-192.
- Maslin, BR & Cowie, ID (2014) *Acacia equisetifolia*, a rare, new species of *Acacia* sect. *Lycopodiifoliae* (Fabaceae: Mimosoideae) from the Top End of the Northern Territory. Nuytsia 24: 1-5.
- Maslin, BR, Thomson, L & Mabberley, DJ (2019a) *Acacia x mangiiformis* hybrida nova (Leguminosae: Mimosoideae), a wattle of commercial importance in Asia. Telopea 22: 161-167.
- Maslin, BR, Ho, BC, Sun, H & Bai, L (2019b) Revision of *Senegalia* in China, and notes on introduced species of *Acacia*, *Acaciella*, *Senegalia*, and *Vachellia* (Leguminosae: Mimosoideae).
- Nielsen, IC (1992) Mimosaceae (Leguminosae – Mimosoideae). In: Flora Malesiana, Series I – Spermatophyta, Volume 11 – part 1. Rijksherbarium / Hortus Botanicus, Leiden University, Leiden, the Netherlands.
- Orchard, AE & Wilson, AJG (eds.) (2001) Flora of Australia. Volume 11A: Mimosaceae, Acacia part 1. Melbourne: ABRS/CSIRO Publishing.
- Pedley, L (1975) Revision of the extra-Australian species of *Acacia* subg. *Heterophyllum*. Contributions from the Queensland Herbarium 18: 1-24.
- Pedley, L (2006a) Notes on *Acacia* Mill. (Leguminosae: Mimosoideae), chiefly from Queensland, 5. Austrobaileya 7(2): 347-356.
- Pedley, L (2006b) Nomenclatural notes on *Acacia* Mill. (Leguminosae – Mimosoideae), consequential to the conservation of its name. Austrobaileya 7(2): 381-382.
- Pedley, L (2019) Notes on *Acacia* Mill. (Leguminosae: Mimosoideae), chiefly from Queensland, 6. Austrobaileya 10(3): 297-320.
- Pelser, PB, Barcelona, JF & Nickrent, DL (eds.) (2011 onwards) Co's Digital Flora of the Philippines (www.philippineplants.org) [visited on 18-11-2021]
- St. John, H (1979) Classification of *Acacia Koa* and relatives (Leguminosae). Hawaiian Plant Studies 93. Pacific Science 33(4): 357-367.
- Turner, IM (2014) Names of extant angiosperm species that are illegitimate homonyms of fossils. Annales Botanici Fennici 51(5): 305-317.
- WorldWideWattle ver. 2. Published on the Internet at: www.worldwidewattle.com [Accessed 30/10/2019].

#### Sources of occurrence data

GBIF, González-Orozco *et al.* 2013.

GBIF DOIs: https://doi.org/10.15468/dl.vkkonk, https://doi.org/10.15468/dl.ohfef1, https://doi.org/10.15468/dl.qmejj8, https://doi.org/10.15468/dl.sw7dgi, and https://doi.org/10.15468/dl.u6slab.

#### Notes

González-Orozco *et al*. (2013) assembled an occurrence dataset of 1020 Australian species of *Acacia* (available from http://dx.doi.org/10.5061/dryad.33kn3). We edited this dataset by adding, where possible, author names to species names using information from WorldWideWattle and IPNI, and in various species-specific ways (see below for full details). Subsequently, a species checklist was made of all species that are listed on WorldWideWattle but that are not present in the González-Orozco *et al*. (2013) dataset, either because they do not occur in Australia or have been described since 2013. GBIF was queried using this species checklist, resulting in 73 additional species being added to the dataset.

The following changes were made to the dataset of González-Orozco *et al*. (2013). For decisions on how to spell species names, we follow WorldWideWattle.

- all records of *A. spirorbis* were assigned to *A. spirorbis* subsp. *solandri* (the dataset of González-Orozco *et* al. (2013) does not contain infraspecific taxa)
- *A. brachycarpa* was changed to *A. neobrachycarpa* following Turner 2014
- *A. calcigera* was removed, as according to Orchard & Wilson 2001 it is a synonym of *A. valida*, which is *Vachellia valida* according to WorldWideWattle
- *A. caleyi* was changed to *A. decora* following Bruce Maslin (pers. comm.)
- *A. cometes* was changed to *A. lachnophylla* following Orchard & Wilson 2001
- *A. crassifructa* was changed to *A. pachycarpa* following Bruce Maslin (pers. comm.)
- *A. cunninghamii* was removed. According to Orchard & Wilson 2001, it is a synonym of either *A. trinervata*, *A. cretata*, *A. crassa*, *A. concurrens*, or *A. leiocalyx*, and it is impossible to tell which one without knowing the author
- *A. curvinervia* was changed to *A. julifera subsp. curvinervia* following WorldWideWattle
- *A. diffusa* was changed to *A. genistifolia* following Orchard & Wilson 2001
- *A. diphylla* was changed to *A. blakei subsp. diphylla* following WorldWideWattle
- *A. eborensis* is an unpublished manuscript name, and was changed to *Acacia sp. Small Red-leaved Wattle (J.B.Williams 95033)* following Bruce Maslin (pers. comm.)
- *A. eglandulosa* was changed to *A. cyclops* following Orchard & Wilson 2001
- *A. exilis* was changed to *A. exigua* following Turner 2014
- *A. gracillima* was changed to *A. minniritchi* following Turner 2014
- *A. hunteriana* was removed. According to Orchard & Wilson 2001 it is a synonym of *A. boormannii*, but the distributions did not match
- *A. jutsonii* was changed to *A. heteroneura var. jutsonii* following WorldWideWattle
- *A. karina* was changed to *A. karinae* following WorldWideWattle
- *A. microcephala* was removed, as it is *Vachellia leucophloea var. microcephala* according to WorldWideWattle
- *A. newmanii* is an unpublished manuscript name, and was changed to *Acacia sp. Kununurra (Lullfitz 6195)* following Bruce Maslin (pers. comm.)
- *A. nova-anglica* is an unpublished manuscript name, and was changed to *Acacia sp. New England (J.B.Williams 97011)* following Bruce Maslin (pers. comm.)
- *A. omalophylla* was changed to *A. homalophylla* following WorldWideWattle
- *A. pennata* was removed, as it is *Senegalia pennata* according to WorldWideWattle
- *A. rigida* was changed to *A. neorigida* following Turner 2014
- *A. robinae* was changed to *A. robiniae* following WorldWideWattle
- *A. stowardii* was changed to *A. sibirica* based on its distribution and following Bruce Maslin (pers. comm.)
- *A. tindaleae* was changed to *A. mariae* following Pedley 2006a

- *A. verricula* was changed to *A. verriculum* following WorldWideWattle

Although Le Roux *et al*. (2014) show that *A. heterophylla* and *A. koa* should be considered to belong to the same species, this taxonomic change has not officially been made yet, so here we keep them as two distinct taxa.

Pedley (1975) considered *A. koaia* to be a synonym of *A. koa*, but here we follow St. John (1979) and treat them as two distinct species.

---

#### *Acaciella*

##### References used for taxonomy and distribution

- Isely, D (1969) Legumes of the United States: I. Native *Acacia*. Sida 3(6): 365-386.
- Isely, D (1973) Leguminosae of the United States: 1. Subfamily Mimosoideae. Memoirs of the New York Botanical Garden 25(1), New York, USA.
- Turner, BL (2015) New names for the Texas taxa of *Acacia* (Fabaceae). Phytologia 97(2): 120-122.
- Rico Arce, M de L & Bachman, S (2006) A taxonomic revision of *Acaciella* (Leguminosae, Mimosoideae). Anales del Jardín Botánico de Madrid 63(2): 189-244.

##### Sources of occurrence data

GBIF, DryFlor, SEINet.

GBIF DOI: https://doi.org/10.15468/dl.olcr3l.

##### Notes

Following Rico Arce & Bachman 2006 and Isely 1973, occurrences of *A. villosa* in northern Mexico and the USA were removed.

---

#### *Adenanthera*

##### References used for taxonomy and distribution

- Du Puy, DJ, Labat, J-N, Rabevohitra, R, Villiers, J-F, Bosser, J & Moat, J (2002) The Leguminosae of Madagascar. Kew Publishing.
- Nielsen, IC (1992) Flora Malesiana. Series I – Spermatophyta. Flowering Plants. Volume 11, part 1. Mimosaceae (Leguminosae – Mimosoideae). Rijksherbarium/Hortus Botanicus, Leiden University, Leiden, the Netherlands.
- Nielsen, I & Guinet, Ph (1992) Synopsis of *Adenanthera* (Leguminosae-Mimosoideae). Nordic J. Bot. 12: 85-114.

##### Sources of occurrence data

GBIF.

GBIF DOI: https://doi.org/10.15468/dl.ahjxrw.

**Notes**

Following Nielsen 1992 and Nielsen & Guinet 1992, all occurrences of *A. pavonina* outside the area between the Malay Peninsula and northern Australia were considered cultivated and removed (occurrences in the Philippines were also removed).

---

##### ***Adenopodia*, *Microlobius*, *Parapiptadenia*, *Piptadenia*, *Pityrocarpa*, *Pseudopiptadenia* and *Stryphnodendron***

**References used for taxonomy and distribution**

- Barneby R. C. & Grimes J. W. 1984. Two new mimosaceous trees from the American tropics. Brittonia 36, 236-240.
- Barneby R. C. & Grimes J. W. 1984. Two new leguminous forest trees new to the flora of French Guiana. Brittonia 36, 45-48.
- Barneby R. C. 1986. A contribution to the taxonomy of *Piptadenia* (Mimosaceae) in South America. Brittonia 38, 222-229.
- Barneby R. C. in Berry P. E. *et al*. 2001. Flora of the Venezuelan Guiana. Vol 6. Liliaceae-Myrsinaceae. Missouri Botanical Garden pp. 803.
- Brenan J. P. M. 1955. Notes on Mimosoideae: 1. Kew Bulletin 10, 161-192.
- Brenan J. P. M. 1986. The genus *Adenopodia* (Leguminosae). Kew Bulletin 41, 73-90.
- Grimes J. W. 1993. *Calliandra anthoniae*, new species (Leguminosae: Mimosoideae: Ingeae) and a new combination in *Pseudopiptadenia* Rauschert (Leguminosae: Mimosoideae, Mimoseae). Brittonia 45, 25-27.
- Guinet P. & Caccavari M. A. 1992. Pollen morphology of the genus *Stryphnodendron* (Leguminosae: Mimosoideae) in relation to its taxonomy. Grana 31, 101-112.
- Jobson R. W. & Luckow M. 2007. Phylogenetic study of the genus *Piptadenia* (Mimosoideae: Leguminosae) using plastid trnL-F and trnK/matK sequenc data. Systematic Botany 32, 569-575.
- Lewis G. P. 1991. A new combination in *Pseudopiptadenia* (Leguminosae: Mimosoideae). Kew Bulletin 46, 118-119.
- Lewis G. P. 1991. Another new combination in *Pseudopiptadenia*. Kew Bulletin 46, 358.
- Lewis G. P. 1991. Five new taxa of *Piptadenia* from Brazil. Kew Bulletin 46, 159-168.
- Lewis G. P. & Lima L. P. M. 1991. *Pseudopiptadenia* no Brasil (Leguminosae: Mimosoideae) Arch. Jard. Bot. Rio de Janeiro 30, 43-67.
- Lewis G. P. 1994. A new species of *Parapiptadenia* (Leguminosae: Mimosoideae) from Brazil. Kew Bulletin 49, 99-101.
- Lewis G. P., Schrire B., Mackinder B., Lock M. (2005) *Legumes of the world*. Kew, UK: Royal Botanic Gardens.
- de Lima M. P. M. & Lima H. C. 1984. *Parapiptadenia* Brenam (Leguminosae: Mimosoideae). Estudo taxonomico das especies brasileiras. Rodriguesia 36, 23-30.
- Macbride, J.F. (1919) Notes on certain Leguminosae. Contributions from the Gray Herbarium of Harvard University 59, 1-27.
- Neill D. A. & Occhioni-Martins E. M. 1989. A new species of *Stryphnodendron* (Fabaceae: Mimosoideae) from Amazonian Ecuador. Annals Missouri Bot. Gard. 76, 357-359.

- Sousa S. M. & Andrade M. G. 1992. Identidad de *Microlobius* y *Goldmania* (Mimosoideae: Mimoseae) y nuevas combinaciones. Anales Inst. Biol. Univ. Nac. Auton. Mexico, Bot. 63, 101-107.
- Occhioni Martins E. M. & Martins A. G. 1972. *Stryphnodendron* Mart. (Leguminosae-Mimosoideae) as especies da Amazonia Brasileira. Leandra 2(2), 11-40.
- Occhioni Martins E. M. 1974. *Stryphnodendron* Mart. (Leguminosae-Mimosoideae): as especies do nordeste, sudeste e sul do Brasil II. Leandra 4-5, 53-66.
- Occhioni Martins E. M. 1981. *Stryphnodendron* Mart. (Leguminosae-Mimosoideae) com especial referencia aos taxa Amazonicos. Leandra 10-11, 3-100.
- Occhioni E. M. L. 1990. Consideracoes taxonomicas no genero *Stryphnodendron* Mart. (Leguminosae-Mimosoideae) e distribuicao geografica das especies. Acta Bot. Brasil. 4, 153-158.
- Occhioni Martins E. M. in Berry P. E. et al. 2001. Flora of the Venezuelan Guiana. Vol 6. Liliaceae-Myrsinaceae. Missouri Botanical Garden pp. 803.
- de Queiroz, LP (2009) Leguminosas da Caatinga. Universidade Estadual de Feira de Santana.
- Rizzini C. T. & Heringer E. P. 1987. As especies anas de *Stryphnodendro*n Mart. (Leguminosae-Mimosoideae). Rev. Brasil. Biol. 47, 447-454.
- Scalon V. R. 2007. Revisao taxonomica do genero *Stryphnodendron* Mart. (Leguminosae-Mimosoideae). PhD Thesis. Universidade de Sao Paulo.

**Sources of occurrence data**

GBIF, DryFlor, SEINet.

GBIF DOI: https://doi.org/10.15468/dl.glqwcj.

**Notes**

*Piptadenia flava* does not occur in the Caatinga according to de Queiroz 2009, and its occurrences there were removed.

---

***Afrocalliandra*, *Amblygonocarpus*, *Aubrevillea*, *Calpocalyx*, *Chidlowia*, *Cylicodiscus*, *Elephantorrhiza*, *Faidherbia*, *Fillaeopsis*, *Newtonia*, *Piptadeniastrum*, *Pseudoprosopis* and *Tetrapleura***

**References used for taxonomy and distribution**

- African Plant Database (version 3.4.0). Conservatoire et Jardin botaniques de la Ville de Genève and South African National Biodiversity Institute, Pretoria, Retrieved October 2019 from http://www.ville-ge.ch/musinfo/bd/cjb/africa/.
- Barnes, RD & Fagg, CW (2003) *Faidherbia albida*. Monograph and annotated bibliography. Oxford Forestry Institute, Department of Plant Sciences, University of Oxford, United Kingdom.
- Brenan, JPM (1959) Flora of Tropical East Africa. Leguminosae subfamily Mimosoideae.
- Brenan, JPM (1984) A new record and a new taxon in the genus *Pseudoprosopis* (Leguminosae) from Africa. Kew Bulletin 39(3): 657-658.
- Burkill, HM (1995) The Useful Plants of West Tropical Africa, vol. 3, families J-L. Royal Botanic Gardens Kew.
- de Souza, ER, Lewis, GP, Forest, F, Schnadelbach, AS, van den Berg, C & de Queiroz, LP (2013) Phylogeny of *Calliandra* (Leguminosae: Mimosoideae) based on nuclear and plastid molecular markers. Taxon 62(6): 1201-1220.

- Greve, M, Lykke, AM, Fagg, CW, Bogaert, J, Friis, I, Marchant, R, Marshall, AR, Ndayishimiye, J, Sandel, BS, Sandom, C, Schmidt, M, Timberlake, JR, Wieringa, JJ, Zizka, G & Svenning, J-C (2012) Continental-scale variability in browser diversity is a major driver of diversity patterns in acacias across Africa. Journal of Ecology 100: 1093-1104.
- Hoyle, AC (1932) *Chidlowia*, a new tree genus of Caesalpiniaceae from West tropical Africa. Bull. Misc. Inform. Kew 1932(2): 101.
- Lewis, G, Schrire, B, Mackinder, B & Lock, M (2005) Legumes of the World. Royal Botanic Gardens, Kew.
- Mackinder, B & Cheek, M (2003) A new species of *Newtonia* (Leguminosae-Mimosoideae) from Cameroon. Kew Bulletin 58(2): 447-452.
- Ross, JH (1974) The genus *Elephantorrhiza*. Bothalia 11(3): 247-257.
- Ross, JH (1975) Mimosoideae. Flora of Southern Africa 16(1). Pretoria, Botanical Research Institute.
- Villiers, J-F (1983) Le genre *Pseudoprosopis* Harms (Mimosaceae) en Afrique. Bulletin du Jardin botanique National de Belgique 53(3/4): 417-436.
- Villiers, J-F (1984) Le genre *Calpocalyx* (Leguminosae, Mimosoideae) en Afrique. Bull. Mus. Natn. Hist. Nat., Paris, 4e sér., 6, section B, Adansonia, no 3: 297-311.
- Villiers, J-F (1989) Flore du Gabon 31: Leguminosae, Mimosoideae.
- Villiers, J-F (1990) Contribution à l'étude du genre Newtonia Baillon (Leguminosae-Mimosoideae) en Afrique. Bulletin du Jardin botanique National de Belgique 60(1/2): 119-138.

**Sources of occurrence data**

GBIF, Greve *et al*. 2012.

GBIF DOI: https://doi.org/10.15468/dl.tsdtum.

---

***Alantsilodendron*, *Calliandropsis*, *Desmanthus*, *Dichrostachys*, *Gagnebina*, *Kanaloa*, *Lemurodendron*, *Leucaena*, *Mimozyganthus*, *Neptunia*, *Piptadeniopsis*, *Prosopidastrum* and *Schleinitzia***

**References used for taxonomy and distribution:**

- Aebli, Anahita (2015) Assesmbly of the Madagascan biota by replicated adaptive radiations: Case studies in Leguminosae – Mimosoideae. Master Thesis, Institute of Systematic Botany, University of Zürich.
- Du Puy, DJ, Labat, J-N, Rabevohitra, R, Villiers, J-F, Bosser, J & Moat, J (2002) The Leguminosae of Madagascar. Kew Publishing.
- Govindarajulu, R, Hughes, CE, Alexander, PJ & Bailey, CD (2011) The complex evolutionary dynamics of ancient and recent polyploidy in *Leucaena* (Leguminosae; Mimosoideae). American Journal of Botany 98(12): 2064-2076.
- Hughes, CE (1998) Monograph of *Leucaena* (Leguminosae: Mimosoideae). Systematic Botany Monographs 55: 1-244.
- Hughes, CE & Bailey, D. The genus *Leucaena*. BRAHMS Online database. https://herbaria.plants.ox.ac.uk/bol/leucaena. Last accessed 05/11/2018.
- Lewis, GP & Guinet, Ph (1986) Notes on *Gagnebina* (Leguminosae : Mimosoideae) in Madagascar and Neighbouring Islands. Kew Bulleting 41(2): 463-470.
- Lorence, D.H. & K.R. Wood (1994) *Kanaloa*, a New Genus of Fabaceae (Mimosoideae) from Hawaii. Novon, vol. 4, no. 2: 137-145.

- Luckow, M (1993) Monograph of *Desmanthus* (Leguminosae – Mimosoideae). Systematic Botany Monographs 38: 1-166.
- Luckow, M (1995) A phylogenetic analysis of the *Dichrostachys* group (Mimosoideae: Mimoseae). In: Crisp, M & Doyle, JJ (editors) Advances in Legume Systematics 7: Phylogeny, pp. 63-76. Royal Botanic Gardens, Kew.
- Luckow, M, Fortunato, RH, Sede, S & Livshultz, T (2005) The phylogenetic affinities of two mysterious monotypic mimosoids from southern South America. Systematic Botany 30(3): 585-602.
- Nielsen, IC (1992) Flora Malesiana. Series I – Spermatophyta. Volume 11, part 1. Mimosaceae (Leguminosae – Mimosoideae). Rijksherbarium, Leiden University, The Netherlands.
- Orchard, AE & McCarthy, PM (1998) Flora of Australia - Volume 12: Mimosaceae (excl. Acacia), Caesalpiniaceae. Melbourne: CSIRO Australia.
- Thulin, M (1983) Leguminosae of Ethiopia. Opera Botanica 68: 1-223.
- Thulin, M (1989) New or noteworthy species of Leguminosae in NE tropical Africa. Nordic Journal of Botany 8(5): 457-488.
- Varjâo Româo, M.C. & Mansano, V.F. (2018) A new combination in *Parkinsonia* (Caesalpinioideae/Fabaceae): *Parkinsonia andicola*. Phytotaxa 344: 295-296.

**Sources of occurrence data**

GBIF, Aebli 2015, Lorence & Wood 1994, Hughes & Bailey (Leucaena database), DryFlor, SEINet, J.L. Contreras (pers. comm.).

GBIF DOIs: http://doi.org/10.15468/dl.95gzq9, http://doi.org/10.15468/dl.cwclrm, http://doi.org/10.15468/dl.cynmca, http://doi.org/10.15468/dl.e7gb4q, http://doi.org/10.15468/dl.fun5sf, http://doi.org/10.15468/dl.ilgxuv, http://doi.org/10.15468/dl.l45f4x, http://doi.org/10.15468/dl.q8dlpd, http://doi.org/10.15468/dl.qgncrw, http://doi.org/10.15468/dl.uccakm, http://doi.org/10.15468/dl.vqsaxm, and http://doi.org/10.15468/dl.pmef26.

**Notes**

Additional records of *Dichrostachys dehiscens* were obtained from Alan Forrest.

Following Lewis & Guinet (1986) and Luckow (1995), *Gagnebina bernieriana*, *G. pervilleana*, and *G. myriophylla* are all treated as species of *Gagnebina* rather than *Dichrostachys*.

The infraspecific taxa of *Dichrostachys cinerea* are here lumped together at the species level.

As *Leucaena leucocephala* is widely invasive and cultivated throughout the tropics and its origin is not well-known (Govindarajulu *et al*. 2011), it was removed from the dataset.

---

#### *Albizia* (Old World)

**References used for taxonomy and distribution**

- African Plant Database (version 3.4.0). Conservatoire et Jardin botaniques de la Ville de Genève and South African National Biodiversity Institute, Pretoria, Retrieved October 2019 from http://www.ville-ge.ch/musinfo/bd/cjb/africa/.
- Cowan, RS (1998) Mimosaceae (excl. *Acacia*), Caesalpiniaceae. In: Orchard, AE (ed.), Flora of Australia, Volume 12. CSIRO Publishing, Collingwood, Australia.
- Du Puy, DJ, Labat, J-N, Rabevohitra, R, Villiers, J-F, Bosser, J & Moat, J (2002) The Leguminosae of Madagascar. Kew Publishing.
- Editorial Committee of the Flora of Taiwan (1993) Flora of Taiwan. Second Edition. Volume three: Angiosperms, Dicotyledons, Hamamelidaceae – Umbelliferae. Taipei, Taiwan.
- Nielsen, IC (1979) Notes on the genus *Albizia* Durazz. (Leguminosae – Mimosoideae) in mainland S.E. Asia. Adansonia, sèr. 2, 19(2): 199-229.
- Nielsen, IC (1985) The Malesian species of *Acacia* and *Albizia* (Leguminosae – Mimosoideae). Opera Botanica 81: 5-50.
- Nielsen, IC (1992) Mimosaceae (Leguminosae – Mimosoideae). In: Flora Malesiana, Series I – Spermatophyta, Volume 11 – part 1. Rijksherbarium / Hortus Botanicus, Leiden University, Leiden, the Netherlands.
- Wu, D & Nielsen, IC (2010) Flora of China. Volume 10, part 7. Tribe Ingeae.

**Sources of occurrence data**

GBIF.

GBIF DOIs: https://doi.org/10.15468/dl.sgxk8x, https://doi.org/10.15468/dl.m2v18r, https://doi.org/10.15468/dl.bli6ug, and https://doi.org/10.15468/dl.3ko5dm.

---

#### ***Anadenanthera*, *Lachesiodendron* and *Plathymenia***

**References used for taxonomy and distribution**

- Altschul, SvR (1965) A taxonomic study of the genus *Anadenanthera*. Contributions from the Gray Herbarium of Harvard University 193: 1-65.
- Little, Jr., EL & Wadsworth, FH (1964) Common trees of Puerto Rico and the Virgin Islands. U.S. Department of Agriculture, Washington D.C., United States of America.
- Ribeiro, PG, Luckow, M, Lewis, GP, Simon, MF, Cardoso, D, de Souza, ER, Silva, APC, Jesus, MC, dos Santos, FAR, Azevedo, V, de Queiroz, LP (2018) *Lachesiodendron*, a new monospecific genus segregated from *Piptadenia* (Leguminosae: Caesalpinioideae: mimosoid clade): Evidence from morphology and molecules. Taxon 67(1): 37-54.
- Warwick, MC & Lewis, GP (2003) Revision of *Plathymenia* (Leguminosae – Mimosoideae). Edinburgh Journal of Botany 60(2): 111-119.

**Sources of occurrence data**

GBIF, DryFlor, SEINet.

GBIF DOI: https://doi.org/10.15468/dl.mf9qjm.

---

#### *Archidendron*

##### References used for taxonomy and distribution

- Cowan, RS (1998) Mimosaceae (excl. *Acacia*), Caesalpiniaceae. In: Orchard, AE (ed.), Flora of Australia, Volume 12. CSIRO Publishing, Collingwood, Australia.
- Dash, SS & Sanjappa, M (2011) Two new species and a new distributional record of *Archidendron* F. Muell. (Leguminosae: Mimosoideae) from India. Nelumbo 53: 7-16.
- Gangopadhyay, M & Chakrabarty, T (1993) The genus *Archidendron* F. v. Muell. (Mimosaceae) in India. Journal of Economic and Taxonomic Botany 17(3): 683-691.
- Nielsen, IC (1979) Notes on the genera *Archidendron* F. v. Mueller and *Pithecellobium* Martius in mainland S.E. Asia. Adansonia, sér. 2, 19(1): 3-37.
- Nielsen, IC, Baretta-Kuipers, T & Guinet, Ph (1984) The genus *Archidendron* (Leguminosae – Mimosoideae). Opera Botanica 76: 1-120.
- Nielsen, IC (1992a) Mimosaceae (Leguminosae – Mimosoideae). In: Flora Malesiana, Series I – Spermatophyta, Volume 11 – part 1. Rijksherbarium / Hortus Botanicus, Leiden University, Leiden, the Netherlands.
- Nielsen, IC (1992b) The Australian species of *Archidendron*. Nordic Journal of Botany 2(5): 479-490.
- Ramesh, BR, Swaminath, MH, Patil, SV, Dasappa, Pélissier, R, Venugopal, PD, Aravajy, S, Elouard, C & Ramalingam, S (2010) Forest stand structure and composition n 96 sites along environmental gradients in the central Western Ghats of India. Ecology 91(10): 3118-3118.
- Sanjappa, M (1991) Legumes of India. Bishen Singh Mahendra Pal Singh, Dehra Dun, India.

##### Sources of occurrence data

GBIF, Dash & Sanjappa 2011, Ramesh *et al*. 2010.

GBIF DOI: https://doi.org/10.15468/dl.87hdqa.

---

#### *Archidendropsis* and *Pararchidendron*

##### References used for taxonomy and distribution

- Cowan, RS (1998) Mimosaceae (excl. *Acacia*), Caesalpiniaceae. In: Orchard, AE (ed.), Flora of Australia, Volume 12. CSIRO Publishing, Collingwood, Australia.
- Nielsen, IC, Guinet, Ph & Baretta-Kuipers, T (1983) Studies in the Malesian, Australian and Pacific Ingeae (Leguminosae – Mimosoideae): the genera *Archidendropsis*, *Wallaceodendron*, *Paraserianthes*, *Pararchidendron* and *Serianthes* (part 1). Bull. Mus. natn. Hist. nat., Paris, 4e sér., 5, section B, Adansonia 3: 303-329.
- Nielsen, IC, Guinet, Ph & Baretta-Kuipers, T (1983) Studies in the Malesian, Australian and Pacific Ingeae (Leguminosae – Mimosoideae): the generea *Archidendropsis*, *Wallaceodendron*, *Paraserianthes*, *Pararchidendron* and *Serianthes* (part 2). Bull. Mus. natn. Hist. nat., Paris, 4e sér., 5, section B, Adansonia 4: 335-360.
- Nielsen, IC, Guinet, Ph & Baretta-Kuipers, T (1984) Studies in the Malesian, Australian and Pacific Ingeae (Leguminosae – Mimosoideae): the genera *Archidendropsis*, *Wallaceodendron*, *Paraserianthes*, *Pararchidendron* and *Serianthes* (part 3). Bull. Mus. natn. Hist. nat., Paris, 4e sér., 6, section B, Adansonia 1: 79-111.
- Nielsen, IC (1992) Mimosaceae (Leguminosae – Mimosoideae). In: Flora Malesiana, Series I –

Spermatophyta, Volume 11 – part 1. Rijksherbarium / Hortus Botanicus, Leiden University, Leiden, the Netherlands.

**Sources of occurrence data**

GBIF.

GBIF DOI: https://doi.org/10.15468/dl.yqc1jg.

---

#### *Calliandra umbrosa*

**References used for taxonomy and distribution**

- Barneby, RC (1998) Silk tree, guanacaste, monkey's earring: A generic system for the synandrous Mimosaceae of the Americas; *Calliandra*. Memoirs of the New York Botanical Garden 74(3): 1-223.
- Paul, SR (1979) The genus *Calliandra* (Mimosaceae) in the Indian subcontinent. Feddes Repertorium 90(3): 155-164.
- de Souza, ER, Lewis, GP, Forest, F, Schnadelbach, AS, van den Berg, C & de Queiroz, LP (2013) Phylogeny of *Calliandra* (Leguminosae: Mimosoideae) based on nuclear and plastid molecular markers. Taxon 62(6): 1200-1219.

**Notes**

*Calliandra umbrosa* was segregated from the New World species of *Calliandra* by Barneby (1998) and should probably be placed in a new genus (de Souza *et al*. 2013), but no new combination has been made yet.

There are no occurrence records with coordinates of *C. umbrosa* on GBIF or any other repository of occurrence data that we know of.

---

#### *Entada*

**References used for taxonomy and distribution**

- Barneby, RC (1996) Neotropical Fabales at NY: asides and oversights. Brittonia 48(2): 174-187.
- Brenan, JPM (1955) Notes on Mimosoideae: I. Kew Bulletin 10(2): 161-192.
- Brenan, JPM (1959) Flora of Tropical East Africa. Leguminosae subfamily Mimosoideae.
- Brenan, JPM (1966) The genus *Entada*, its subdivisions and a key to the African species. Kew Bulletin 20(3): 361-378.
- Brenan, JPM (1978) New species of *Entada* and *Acacia* (Leguminosae) from Africa. Notes on Mimosoideae: XIII. Kew Bulletin 32(3): 545-550.
- Du Puy, DJ, Labat, J-N, Rabevohitra, R, Villiers, J-F, Bosser, J & Moat, J (2002) The Leguminosae of Madagascar. Kew Publishing.
- Hutchinson, J, Dalziel, JM & Keay, RWJ (1954-1958) Flora of West Tropical Africa. Second edition,

volume I, part I & II. The Whitefriars Press, Ltd., London and Tunbridge, United Kingdom.
- Lewis, G, Schrire, B, Mackinder, B & Lock, M (2005) Legumes of the World. Royal Botanic Gardens, Kew.
- Nielsen, IC (1981) Flore du Cambodge, du Laos et du Viet-Nam. Vol 19: Légumineuses – Mimosoïdées. Muséum National d'Histoire Naturelle, Paris, France.
- Nielsen, IC (1992) Mimosaceae (Leguminosae-Mimosoideae). Flora Malesiana, series I – Spermatophyta, volume 11, part I.
- Ramesh, BR, Swaminath, MH, Patil, SV, Dasappa, Pélissier, R, Venugopal, PD, Aravajy, S, Elouard, C & Ramalingam, S (2010) Forest stand structure and composition n 96 sites along environmental gradients in the central Western Ghats of India. Ecology 91(10): 3118-3118.
- Rodrigues, RS & Flores, AS (2012) A new combination in *Entada* (Leguminosae) from Roraima, Brazil. Phytotaxa 39: 47-59.
- Ross, JH (1975) Flora of Southern Africa. Volume 16, part I. Government Printer, Pretoria, South Africa.
- Sanjappa, M (1991) Legumes of India. Bishen Singh Mahendra Pal Singh, Dehra Dun, India.
- Tateishi, Y, Wakita, N & Kajita, T (2008) Taxonomic revision of the genus *Entada* (Leguminosae) in the Ryukyu Islands, Japan. Acta Phytotaxonomica et Geobotanica 59(3): 194-210.
- Villiers, J-F (1982) Une nouvelle espèce du genre *Entada* Adans. (Leguminosae, Mimosoideae) en Afrique occidentale. Bull. Mus. Natn. Hist. Nat., Paris, 4e sèr., 4, section B, Adansonia, 3-4: 193-197.

**Sources of occurrence data**

GBIF, SEINet, Ramesh *et al*. 2010, Rodrigues & Flores 2012.

GBIF DOIs: https://doi.org/10.15468/dl.femx39, https://doi.org/10.15468/dl.jwrrz4, https://doi.org/10.15468/dl.peiayu, https://doi.org/10.15468/dl.vpl5ch, and https://doi.org/10.15468/dl.zhyxse.

**Notes**

As there has been no recent global revision of *Entada*, it is difficult to distinguish between accepted names and synonyms for some species. Here we follow Du Puy *et al*. 2002, Nielsen 1992, and Sanjappa 1991 by treating *E. pursaetha* as a synonym of *E. rheedii*, Barneby 1996 by treating *E. polyphylla* as a synonym of *E. polystachya var. polyphylla*, and Ross 1975 by treating *E. nana* as a synonym of *E. arenaria*.

---

#### *Indopiptadenia*

**References used for taxonomy and distribution**

- Bajpai, O, Srivastava, AK, Kushwaha, AK & Chaudhary, LB (2014) Taxonomy of a monotypic genus *Indopiptadenia* (Leguminosae-Mimosoideae). Phytotaxa 164(2): 61-78.

**Sources of occurrence data**

Bajpai et al. 2014, B. Adhikari (personal communication).

**Notes**

Coordinates of three collections of *Indopiptadenia oudhensis* were provided by B. Adhikari.

---

#### *Inga*

##### References used for taxonomy and distribution

- Acevedo-Rodríguez, P & Strong, MT (2012) Catalogue of Seed Plants of the West Indies. Smithsonian Contributions to Botany, number 98. Washington D.C., USA.
- Cardoso, DBOS, Marinho, LC & Amorim, AM (2017) A remarkable new bifoliolate species of *Inga* (Leguminosae) from the Brazilian Atlantic Forest. Systematic Botany 42(3): 516-521.
- Cornejo, X & Bonifaz, C (2005) *Inga colonchensis* (Fabaceae, Mimosoideae), una Nueva Endémica del Bosque Seco Tropical en Ecuador. Novon 15: 270-273.
- Dexter, KG, Pennington, TD & Cunningham, CW (2010) Using DNA to assess errors in tropical tree identifications: How often are ecologists wrong and when does it matter? Ecological Monographs 80(2): 267-286.
- Dexter, KG & Pennington, TD (2011) *Inga pitmanii* (Fabaceae), a New Species from Madre de Dios, Peru. Novon 21: 322-325.
- Guevara Andino, JE, Pitman, NCA, Hernández, C, Valencia, R, Coley, PD, Kursar, TA & Endara, M-J (2019) A common but overlooked new species in the hyper-diverse genus *Inga* Mill. from the northwestern Amazon. Systematic Botany 44(3): 536-547.
- Padilla-V., E., Cuevas-G., R & Solís-M., A. (2005) *Inga colimana* (Leguminosae) una especie nueva del occidente de México. Acta Botanica Mexicana 72: 33-38.
- Pennington, TD (1997) The genus Inga. The Royal Botanic Gardens, Kew.
- Pennington, TD (2019) Leguminosae – Ingeae, part 1: *Inga*. Flora of Ecuador, no. 96, parts 82-84.
- Poncy, O (1996) Trois nouvelles espèces de Inga (Mimosaceae) des Guyanes et du Brésil. Bulletin du Muséum National d'Histoire Naturelle Paris, 4e sér., Section B, Adansonia 17-18: 67-73.
- Poncy, O (2007) A new species of *Inga* Mill. (Fabaceae, Mimosoideae) from the Guianas. Adansonia, sér. 3 29(2): 249-254.
- Romero, C (2005) Revisión de las especies Colombianas de *Inga* sección *Pseudinga*. In: Forero, E & Romero, C (eds.) Estudios en Leguminosas Colombianas. Bogotá, D.C., Colombia.
- Sousa S., M. (2009) Adiciones al género *Inga* (Ingeae, Mimosoideae, Leguminosae) para la flora Mesoamericana. Acta Botanica Mexicana 89: 25-41.

##### Sources of occurrence data

GBIF, DryFlor, SEINet, Pennington 2019, Dexter *et al*. 2010, Guevara Andino *et al*. 2019, Kyle Dexter (pers. comm.).

GBIF DOIs: https://doi.org/10.15468/dl.qdqpul and https://doi.org/10.15468/dl.bdberq.

##### Notes

Based on unpublished phylogenetic data from Kyle Dexter, Toby Pennington *et al*., and personal communication with Kyle Dexter, the *Inga* occurrence dataset and taxonomic checklist were edited in various ways. Besides removal of outliers from numerous species distribution maps, the following changes were made:

- part of the records of *I. barbata* were split off as *I. aff. barbata*

- part of the records of *I. brevipes* were split off as *I. aff. brachystachys*

- the subspecies of *I. ciliata*, *subsp. ciliata* and *subsp. subcapitata*, are treated as distinct species

- part of the records of *I. cylindrica* were split off as *I. aff. cylindrica*

- all subspecies of *I. nobilis* were lumped together at the species level in the occurrence dataset

- part of the records of *I. pezizifera* were split off as *I. aff. pezizifera*

- the subspecies of *I. subnuda*, *subsp. subnuda* and *subsp. luschnathiana*, are treated as distinct species

- all subspecies of *I. thibaudiana* were lumped together at the species level in the occurrence dataset

- part of the records of *I. velutina* were split off as *I. aff. velutina*

- two of the subspecies of *I. vera*, *subsp. vera* and *subsp. affinis*, are treated as distinct species

- records for the following currently unpublished species were added: *I. LA41*, *I. tripa*, *I. aff. umbellifera*, and *I. mendozana*

- *Inga schinifolia* is treated as distinct from *I. tenuis*

- *Inga jenmanii* is treated as distinct from *I. sertulifera subsp. leptopus*

- *Inga aria* is treated as distinct from *I. cayennensis*

- *Inga brevialata* is treated as distinct from *I. umbratica*

- *Inga longipedunculata* is treated as distinct from *I. leiocalycina*

- *Inga klugii* is treated as distinct from *I. coruscans*.

Due to these changes, various species are included in the occurrence dataset that are currently not formally described/published, and these species are therefore not included in the taxonomic checklist.

---

#### ***Mariosousa***

##### References used for taxonomy and distribution

- Jawad, JT, Seigler, DS & Ebinger, JE (2000) A systematic treatment of *Acacia coulteri* (Fabaceae, Mimosoideae) and similar species in the New World. Annals of the Missouri Botanical Garden 87(4): 528-548.
- Rico-Arce, MdL (2007) American species of *Acacia*. Offset Rebosán, S.A. de C.V., Mexico.
- Seigler, DS, Ebinger, JE & Miller, JT (2006) *Mariosousa*, a new segregate genus from *Acacia* s.l.

(Fabaceae, Mimosoideae) from Central and North America. Novon 16(3): 413-420.
- Seigler, DS & Ebinger, JE (2018) New combinations in *Parasenegalia* and *Mariosousa* (Fabaceae: Mimosoideae). Phytologia 100(4): 256-259.

**Sources of occurrence data**

GBIF, DryFlor, SEINet.

GBIF DOI: https://doi.org/10.15468/dl.u4ppox.

---

***Mimosa***

**References used for taxonomy and distribution**

- Alam M. K., Yusof MD. 1992. The genus *Mimosa* Linn. from Bangladesh. Bangladesh J. Bot. 21(1), 53-58.
- Atahuachi M., Hughes C. E. 2006. Two new species of *Mimosa* (Fabaceae) endemic to Bolivia. Brittonia 58(1), 59-65.
- Atahuachi M., van der Bent L., Wood J. R. I., Lewis G. P., Hughes C. E. 2016. Bolivian *Mimosa* (Leguminosae, Mimosoideae): three new species and a species checklist. Phytotaxa 260(3). 201-222.
- Barneby R. C. 1991. Sensitivae censitae: a description of the genus *Mimosa* Linnaeus (Mimosaceae) in the New World. Mem. New York Bot. Gard. 65, 1-835.
- Barneby R. C. 1993. Increments to the Genus *Mimosa* (Mimosaceae) from South America. Brittonia 45(4), 328-332.
- Barneby R. C. 1997. Toward a Census of Genus *Mimosa* (Mimosaceae) in the Americas: A New Species from Mexico (Baja California Sur) and Two from Planaltine Brazil (Goias, Minas Gerais). Brittonia 49(4), 452-457.
- Borges L. M., Simon M. F., Pirani J. R. 2014. The census continues: Two new montane species of *Mimosa* (Leguminosae Mimosoideae) from Southeastern Brazil. Phytotaxa 177(1), 35-48.
- Borges L. M., Simon M. F., Pirani J. R. 2017. Less is more. Adjusting the taxonomy of the polytypic *Mimosa setosa* (Leguminosae, Mimosoid). Rodriguésia 68(2), 515-540.
- Brenan J. P. M. 1955. Notes on Mimosoideae: I. Kew Bulletin 10(2), 161-192.
- Dutra V. F., Garcia F. C. P. 2013a. Two new species and one new variety of *Mimosa* sect. Habbasia (Leguminosae: Mimosoideae) from Central Brazil. Kew Bulletin 68(1), 163-171.
- Dutra V. F., Garcia F.C. 2013b. Three new species of *Mimosa* sect. Mimosa (Leguminosae, Mimosoideae) from the campos rupestres of Minas Gerais, Brazil. Brittonia 66(1), 33-41.
- Dutra V. F., Garcia F.C. 2013c. Three New Species of *Mimosa* (Leguminosae) from Minas Gerais, Brazil. Systematic Botany 38(2), 398-405.
- Fortunato R. H., Palese R. 1999. Una especie nueva del genéro *Mimosa* L. (Fabaceae-Mimoseae) para el Chaco boliviano: *M. craspedisetosa* Fortunato & Palese. Contribución al estudio de la flora y vegetación del Chaco. XIII. Candollea 54(1), 83–87.
- Gamble J. S. 1920. The Indian Species of *Mimosa*. Bulletin of Miscellaneous Information (Royal Botanic Gardens, Kew) 1920(1), 1-6.
- Glazier D., Mackinder B. 1997. Nomenclatural Notes on South American *Mimosa* (Leguminosae-Mimosoideae). Kew Bulletin 52(2), 459-463.
- Grether R., Martinez-Bernal A. 1996. *Mimosa tejupilcana*, a New Species of Series Plurijugae (Leguminosae) from the State of Mexico, Mexico. Systematic Botany 21(4), 617-621.
- Grether R. 2000. Nomenclatural Changes in the Genus *Mimosa* (Fabaceae, Mimosoideae) in

Southern Mexico and Central America. Novon. 10(1), 29-37.
- Grether R., Steinmann V. 2014. *Mimosa sotoi* (Leguminosae), a new species from Michoacán, Mexico. Brittonia 67(1), 5-10.
- Grings M., Ribas O. S. 2013. *Mimosa sobralii* (Fabaceae, Mimosoideae), a new tree species endemic to the southern Brazilian highland slopes. Phytotaxa 131(1), 23-28.
- Harms H. 1913. Leguminosae africanae. VI. Bot. Jahrb. Syst. 49: 419.
- Izaguirre P., Beyhaut R. 2002. Dos nuevas especies afines a *Mimosa* Sprengelii (Mimosoideae-Leguminosae) en el distrito Uruguayense de la región neotropical. Bol. Soc. Argent. Bot. 37(1-2), 107-114.
- Izaguirre P., Beyhaut R. 2009. Nuevas especies del género *Mimosa* de las subseries axillares y reptantes (Mimosoideae-Leguminosae) en el distrito Uruguayense de la región neotropical. Bol. Soc. Argent. Bot. 44(3-4), 351-359.
- Izaguirre P., Beyhaut R. 2003. Las Leguminosas en Uruguay y Regiones Vecinas. Parte 2: Caesalpinioideae, Parte 3: Mimosoideae. Ed. Hemisferio Sur, Buenos Aires, 174 – 177.
- Jordao L. S. B., Morim M. P., Baumgratz J. F. A. 2014. A new species of *Mimosa* (Leguminosae) from Brazil. Phytotaxa 184(3), 131-138.
- Jordao L. S. B., Morim M. P., Baumgratz J. F. A., Simon M. F. 2017. A new species of *Mimosa* (Leguminosae) endemic to the Brazilian Cerrado. Phytotaxa 312(2), 237-246.
- Lamarck J. B., Poiret J. L. M. 1783. Encyclopédie méthodique. Botanique, 20.
- Lefèvre G., Labat J.-N. 2006. A New Species of *Mimosa* (Fabaceae, Mimosoideae) from Madagascar. Novon. 16(1), 74-77.
- Lewis G. P., Hughes C. E., Daza Yomona A., Solange Sotuyo J., Simon M. F. 2010. Three new legumes endemic to the Marañón Valley, Perú. Kew Bulletin 65(2), 209-220.
- Morales M., Fortunato R. H. 2010. Novedades Taxonómicas y Nomenclaturales en *Mimosa* L. subser. Mimosa (Leguminosae) Para Sudamérica Austral. Candollea 65(1), 169-184.
- Morales M., Ribas O. S., Santos-Silva J. 2012. A New Polyploid Species of *Mimosa* (Leguminosae, Mimosoideae) from the Highlands of Southern Brazil. Systematic Botany 37(2), 399-403.
- Morales M., Fortunato R. H. 2013. A new species of *Mimosa* (Mimosoideae, Leguminosae) from the inter-Andean dry valleys. Phytotaxa 114(1), 33-41.
- Morales M., Santos-Silva J., Ribas O. S. 2013. A new species of *Mimosa* sect. Mimosa (Leguminosae, Mimosoideae) from Southern Brazil. Brittonia 65(2), 148-153.
- Morales M., Fortunato R. H. 2016. A new xerophytic species of *Mimosa* (Mimosoideae, Leguminosae) from Madagascar. Phytotaxa 270(4), 277-285.
- Queiroz L. P. de, Lewis G. P. 1999. A New Species of *Mimosa* L. (Leguminosae: Mimosoideae) Endemic to the Chapada Diamantina, Bahia, Brazil. Kew Bulletin 54(4), 983-986.
- Rico M. de L., Grether R. 2000. A New Name for *Mimosa diptera* Barneby (Leguminosae: Mimosoideae). Kew Bulletin 55(1), 224.
- Santos-Silva J., Azevedo Tozzi A. M. G. de. 2012. *Mimosa foreroana* (Leguminosae, Mimosoideae), a New Species from Nariño, Colombia. Systematic Botany 37(2), 437-441.
- Santos-Silva J., Simon M. F., Azevedo Tozzi A. M. G. de. 2013. A New Species of “Jurema” (*Mimosa* ser. Leiocarpae Benth.) from Bahia, Brazil. Systematic Botany 38(1), 127-131.
- Santos-Silva J., Simon M. F., Azevedo Tozzi A. M. G. de. 2015. Revisão taxonômica das espécies de *Mimosa* ser. Leiocarpae sensu lato (Leguminosae - Mimosoideae). Rodriguésia 66(1), 95-154.
- Särkinen T.E., Peña J. L. M., Daza Yomona A., Simon M. F.,Pennington R. T., Hughes C. E. 2011. Underestimated endemic species diversity in the dry inter-Andean valley of the Río Marañón, northern Peru: An example from *Mimosa* (Leguminosae, Mimosoideae). Taxon 60(1), 139-150.
- Savassi-Coutinho A. P., Lewis G. P., Souza V. C. 2012. *Mimosa roseoalba* (Leguminosae: Mimosoideae), a new species from Mato Grosso do Sul, Brazil. Kew Bulletin 67(4), 827-831.
- Schmidt-Silveira F., Bordignon S. A. L., Miotto S. T. 2016. A New Endemic *Mimosa* (Leguminosae, Mimosoideae) from Pampa Biome, Brazil. Phytotaxa 245(3), 197–206.
- Silva A. S. L. da, Secco R. de S. 2000. *Mimosa dasilvae*, uma nova Mimosaceae da Amazônia

Brasileira. *Acta Amazonica 30*(3), 449.
- Silva R. R., Azevedo Tozzi A. M. G. de. 2011. Uma nova espécie de *Mimosa* L. (Leguminosae, Mimosoideae) do Centro-Oeste do Brasil. Hoehnea 38, 143-146.
- Simon M. F., Hughes C. E., Harris S. A. 2010. Four New Species of *Mimosa* (Leguminosae) from the Central Highlands of Brazil. Systematic Botany 35(2), 277-288.
- Turner B. L. 1994a. Texas species of *Schrankia* (Mimosaceae) transferred to the genus *Mimosa*. Phytologia 76(5), 412-420.
- Turner B. L. 1994b. Mimosa rupertiana B. L. Turner, a new name for *M. occidentalis* (Wooton & Standley) B. L. Turner, not M. occidentalis Britton & Rose. Phytologia 77(2), 81-82.
- Villarreal Q. J. A. 1992. Dos nuevos taxa del género *Mimosa* (Leguminosae: Mimosoideae) para el norte de México. Acta Botanica Mexicana 20, 45-51.
- Villiers J.-F. 2002. Tribe Mimoseae. In D. J. DuPuy, J.-N. Labat, R. Rabevohitra, J.-F. Villiers, J. Bosser & J. Moat. The Leguminosae of Madagascar. Royal Botanic Gardens, Kew, 159-223.
- Vincent M. A., Zarucchi J. L., Gandhi K. N. 2018. A new varietal combination in *Mimosa pigra* (Fabaceae). Phytoneuron 70, 1-2.
- Weakley A. S., Poindexter D. B., LeBlond R. J., Sorrie B. A., Karlsson C. H., Williams P. J., Bridges E. L., Orzell S. L., Keener B. R., Weeks A., Noyes R. D., Flores-Cruz M., Diggs J. T., Gann G. D., Floden A. J. 2017. New combinations, rank changes, and nomenclatural and taxonomic comments in the vascular flora of the southeastern United States. II. Journal of the Botanical Research Institute of Texas 11, 291-325.

**Sources of occurrence data**

GBIF, DryFlor, SEINet.

GBIF DOIs: https://doi.org/10.15468/dl.ioo9mw, https://doi.org/10.15468/dl.fcnbjl, and https://doi.org/10.15468/dl.glqwcj.

**Notes**

Several widely invasive species, for which the native ranges are unknown, were removed from the dataset: *Mimosa bimucronata* (including var. *bimucronata*), *M. diplotricha* (including vars. *diplotricha* and *inermis*), *M. pigra*, and *M. pudica* (including vars. *unijuga*, *tetrandra*, and *hispida*). (Note that *M. diplotricha* var. *odibilis* and *M. pigra* vars. *asperata*, *dehiscens*, and *hispida* are still in the dataset, as the native ranges of these varieties are well-known (see Glazier & Mackinder 1997 for *M. pigra* vars. *dehiscens* and *hispida*, Vincent *et al*. 2018 for *M. pigra* var. *asperata*, and Barneby 1991 for *M. diplotricha* var. *odibilis*).)

---

#### ***Parasenegalia* and *Pseudosenegalia***

**References used for taxonomy and distribution**

- Barros, MJF & Morim, MP (2014) *Senegalia* (Leguminosae, Mimosoideae) from the Atlantic Domain, Brazil. Systematic Botany 39(2): 452-477.
- Seigler, DS, Ebinger, JE, Riggins, CW, Terra, V & Miller, JT (2017) *Parasenegalia* and *Pseudosenegalia* (Fabaceae): New genera of the Mimosoideae. Novon 25(2): 180-205.
- Seigler, DS & Ebinger, JE (2018) New combinations in *Parasenegalia* and *Mariosousa* (Fabaceae: Mimosoideae). Phytologia 100(4): 256-259.

#### Sources of occurrence data

GBIF, DryFlor, SEINet.

GBIF DOI: https://doi.org/10.15468/dl.xzkqek.

#### Notes

Following Seigler *et al*. 2017, all occurrences of *Pseudosenegalia visco* outside of northern Argentina, Bolivia, northern Chile, and Peru were considered cultivated, and were therefore removed.

Following Barros & Morim 2014, all occurrences of *Parasenegalia miersii* outside of Rio de Janeiro were removed.

---

### *Parkia*

#### References used for taxonomy and distribution

- Hopkins, HCF (1983) The taxonomy, reproductive biology and economic potential of *Parkia* (Leguminosae: Mimosoideae) in Africa and Madagascar. Botanical Journal of the Linnean Society 87(2): 135-167.
- Hopkins, HCF (1986) *Parkia* (Leguminosae: Mimosoideae). Flora Neotropica Vol. 43. New York Botanical Garden Press, New York, USA.
- Hopkins, HCF (1994) The Indo-Pacific species of *Parkia* (Leguminosae: Mimosoideae). Kew Bulletin 49(2): 181-234.
- Hopkins, HCF (2000a) *Parkia paya* (Leguminosae: Mimosoideae), a new species from swamp forest and notes on variation in *Parkia speciosa sensu lato* in Malesia. Kew Bulletin 55(1): 123-132.
- Hopkins, HCF (2000b) *Parkia lutea* (Leguminosae, Mimosoideae), a new species from Amazonian Brazil. Adansonia 22(1): 139-144.
- Hopkins, HCF (2000c) *Parkia barnebyana* (Leguminosae: Mimosoideae), a new species from Venezuelan Guyana. Kew Bulletin 55(1): 133-136.
- Neill, DA (2009) *Parkia nana* (Leguminosae, Mimosoideae), a new species from the sub-Andean sandstone cordilleras of Peru. Novon 19: 204-208.
- Nielsen, I (1980) Notes on Indo-Chinese Mimosaceae. Adansonia, sér. 2, 19(3): 339-363.

#### Sources of occurrence data

GBIF, DryFlor.

GBIF DOI: https://doi.org/10.15468/dl.uedddg.

#### Notes

According to Hopkins 1994, *P. biglandulosa* is only known from cultivation, and its native distribution is unknown. It was therefore removed from this dataset.

---

***Pentaclethra***

**References used for taxonomy and distribution**

- Burkill, HM (1995) The Useful Plants of West Tropical Africa, vol. 3, families J-L. Royal Botanic Gardens Kew, United Kingdom.
- Steyermark, JA, Berry, PE, Yatskievych, K & Holst, BK (eds.) (2001) Flora of the Venezuelan Guyana, vol. 6: Lileaceae – Myrsinaceae. Missouri Botanical Garden Press, St. Louis, United States of America.
- Villiers, J-F (1989) Flore du Gabon 31: Leguminosae, Mimosoideae. Museum national d'histoire naturelle, Paris, France.

**Sources of occurrence data**

GBIF, SEINet.

GBIF DOI: https://doi.org/10.15468/dl.lqpri8.

---

***Prosopis* and *Xerocladia***

**References used for taxonomy and distributions**

- African Plant Database (version 3.4.0). Conservatoire et Jardin botaniques de la Ville de Genève and South African National Biodiversity Institute, Pretoria, Retrieved November 2019 from http://www.ville-ge.ch/musinfo/bd/cjb/africa/.
- Burkart, A (1976) A monograph of the genus *Prosopis* (Leguminosae subfam. Mimosoideae). Journal of the Arnold Arboretum, volume 57, pages 219-249 & 450-525.
- Catalano, SA, Vilardi, JC, Tosto, D & Saidman, BO (2008) Molecular phylogeny and diversification history of *Prosopis* (Fabaceae: Mimosoideae). Biological Journal of the Linnean Society 93: 621-640.
- De Mera, AG, Perea, EL, Quino, JM & Orellana, JAV (2019) *Prosopis andicola* (Algarobia, Caesalpinioideae, Leguminosae), a new combination and rank, and *P. calderensis*, a new species for mesquite populations from Southern Peru. Phytotaxa 414(1): 48-54.
- Johnston, MC (1962) The North American Mesquites: *Prosopis* sect. Algorobia (Leguminosae). Brittonia 14, pages 72-90.
- Léonard, J (1986) Une variété nouvelle de *Prosopis koelziana* Burkart (Mimosacée) de la péninsule arabique. Bulletin du Jardin botanique National de Belgique 56 (3-4): 483-485.
- Núñez, LV, Puicón, JE & Mera, AH (2009) Cinco especies peruanas de *Prosopis* nuevas para la ciencia - Five species of *Prosopis* from Peru news for science. Sciéndo 12(1): 68-87.
- Núñez, LV, Puicón, JE & Mera, AH (2010) Los Algarrobos del Perú. INCAGRO-UNPRG, Lambayeque, Peru.
- Palacios, RA (2006) Los Mezquites Mexicanos: biodiversidad y distribución geográfica. Bol. Soc. Argent. Bot. 41 (1-2), pages 99-121.
- Roig, FA (1987) Arboles y arbustos en *Prosopis flexuosa* y *P. alpataco* (Leguminosae). Parodiana 5(1): 49-64.
- Ross, JH (Ed.) (1975) Mimosoideae. Flora of Southern Africa 16(1), pages 130-131.
- Whaley, OQ, Orellana-Garcia, A & Pecho-Quispe, JO (2019) An annotated checklist to vascular flora of the Ica region, Peru – with notes on endemic species, habitat, climate and agrobiodiversity.

Phytotaxa 389(1): 1-125.
- Zöllner, O & Olivares, M (2001) *Prosopis reptans* Benth. var. *chilensis* (Mimosaceae), una nueva variedad para Chile. Noticiario Mensual, Museo Nacional de Historia Natural. Santiago de Chile 344: 9-14.

**Sources of occurrence data**

GBIF, DryFlor, SEINet, Léonard 1986, Núñez *et al*. 2009, De Mera *et al*. 2019.

GBIF DOIs: https://doi.org/10.15468/dl.8ljqpj, https://doi.org/10.15468/dl.3ymuni, https://doi.org/10.15468/dl.oiol1q, https://doi.org/10.15468/dl.mueoxb, https://doi.org/10.15468/dl.nyrn3i, https://doi.org/10.15468/dl.rv7rtq, and https://doi.org/10.15468/dl.ylttjv.

**Notes**

*Prosopis juliflora* was removed from inland Mexico, as Burkart (1976) mentions it is a coastal species. It was also removed from the Caribbean, since Burkart strongly suspects it has been introduced there.

*Prosopis juliflora* varieties *inermis* and *juliflora* were included at the species level.

Following Whaley *et al*. 2019, *Prosopis limensis* is considered distinct from *P. pallida*. (However, no herbarium records with coordinates are available for *P. limensis*.)

---

#### ***Sanjappa***

**References used for taxonomy and distribution**

- de Souza, ER, Krishnaraj, MV & de Queiroz, LP (2016) *Sanjappa*, a new genus in the tribe Ingeae (Leguminosae: Mimosoideae) from India. Rheedea 26(1): 1-12.

**Sources of occurrence data**

de Souza *et al*. 2016.

---

#### ***Senegalia***

**References used for taxonomy and distribution**

- African Plant Database (version 3.4.0). Conservatoire et Jardin botaniques de la Ville de Genève and South African National Biodiversity Institute, Pretoria, Retrieved July 2019 from http://www.ville-ge.ch/musinfo/bd/cjb/africa/.
- Ali, SI (2014) The genus *Acacia s.l.* in Pakistan. Pak. J. Bot. 46(1): 1-4.
- Barros, MJF & Morim, MP (2014) *Senegalia* (Leguminosae, Mimosoideae) from the Atlantic

Domain, Brazil. Systematic Botany 39(2): 452-477.
- Boatwright, JS, Maurin, O & van der Bank, M (2015) Phylogenetic position of Madagascan species of *Acacia s.l.* And new combinations in *Senegalia* and *Vachellia* (Fabaceae, Mimosoideae, Acacieae). Botanical Journal of the Linnean Society 179(2): 288-294.
- Clarke, HD, Seigler, DS & Ebinger, JE (1990) *Acacia constricta* (Fabaceae: Mimosoideae) and related species from the southwestern U.S. and Mexico. American Journal of Botany 77(3): 305-315.
- Deshpande, AS, Krishnan, S, Janarthanam, MK & Maslin, BR (2019) Annotated checklist of *Senegalia* and *Vachellia* (Fabaceae: Mimosoideae) for the Indian subcontinent. Nordic Journal of Botany 37(4): 1-20.
- Du Puy, DJ, Labat, J-N, Rabevohitra, R, Villiers, J-F, Bosser, J & Moat, J (2002) The Leguminosae of Madagascar. Kew Publishing.
- Ebinger, JE, Seigler, DS & Clarke, HD (2000) Taxonomic revision of South American species of the genus *Acacia* subgenus *Acacia* (Fabaceae: Mimosoideae). Systematic Botany 25(4): 588-617.
- Ebinger, JE, Seigler, DS & Clarke, HD (2002) Notes on the segregates of *Acacia farnesiana* (L.) Willd. (Fabaceae: Mimosoideae) and related species in North America. The Southwestern Naturalist 47(1): 86-147.
- Ebinger, JE (2017) A new *Senegalia* (*S. seigleri*, Fabaceae: Mimosoideae) from Bahia, Brazil. Phytologia 99(2): 126-129.
- Glass, CE & Seigler, DS (2006) A new combination in *Senegalia* and typification of six New World *Acacia* names. Taxon 55(4): 993-995.
- Greve, M, Lykke, AM, Fagg, CW, Bogaert, J, Friis, I, Marchant, R, Marshall, AR, Ndayishimiye, J, Sandel, BS, Sandom, C, Schmidt, M, Timberlake, JR, Wieringa, JJ, Zizka, G & Svenning, J-C (2012) Continental-scale variability in browser diversity is a major driver of diversity patterns in acacias across Africa. Journal of Ecology 100: 1093-1104.
- Hahn, N (2013) *Senegalia lotterii* (Fabaceae) a new species endemic to the Barberton Centre of Endemism, South Africa. Phytotaxa 119(1): 51-54.
- Hahn, N (2016) *Senegalia montis-salinarum*, a new species of Fabaceae: Mimosoideae endemic to the Soutpansberg, South Africa. Phytotaxa 244(2): 174-180.
- Kyalangalilwa, B, Boatwright, JS, Daru, BH, Maurin, O & van der Bank, M (2013) Phylogenetic position and revised classification of *Acacia s.l.* (Fabaceae: Mimosoideae) in Africa, including new combinations in *Vachellia* and *Senegalia*. Botanical Journal of the Linnean Society 172(4): 500-523.
- Lee, YS, Seigler, DS & Ebinger, JE (1989) *Acacia rigidula* (Fabaceae) and related species in Mexico and Texas. Systematic Botany 14(1): 91-100.
- Maslin, BR (2012) New combinations in *Senegalia* (Fabaceae: Mimosoideae) for Australia. Nuytsia 22(6): 465-468.
- Maslin, BR, Seigler, DS & Ebinger, J (2013) New combinations in *Senegalia* and *Vachellia* (Leguminosae: Mimosoideae) for Southeast Asia and China. Blumea 58: 39-44.
- Maslin, BR (2015) Synoptic overview of *Acacia* sensu lato (Leguminosae: Mimosoideae) in East and Southeast Asia. Gardens' Bulletin Singapore 67(1): 231-250.
- Pedley, L (1986) Derivation and dispersal of *Acacia* (Leguminosae), with particular reference to Australia, and the recognition of *Senegalia* and *Racosperma*. Botanical Journal of the Linnaean Society 92(3): 219-254.
- Pedley, L (2014) New combinations for *Senegalia* Raf. and *Vachellia* Wight & Arn. species (Mimosaceae) that occur in Australia. Austrobaileya 9(2): 314-315.
- de Queiroz, LP (2009) Leguminosas da Caatinga. Universade Estadual de Feira de Santana.
- Ragupathy, S, Seigler, DS, Ebinger, JE & Maslin, BR (2014) New combinations in *Vachellia* and *Senegalia* (Leguminosae: Mimosoideae) for south and west Asia. Phytotaxa 162(3): 174-180.
- Rico-Arce, MdL (2007) American species of *Acacia*. Offset Rebosán, S.A. de C.V., Mexico.
- Seigler, DS & Ebinger, JE (1988) *Acacia macracantha*, *A. pennatula*, and *A. cochliacantha* (Fabaceae: Mimosoideae) species complexes in Mexico. Systematic Botany 13(1): 7-15.
- Seigler, DS, Ebinger, JE & Miller, JT (2006) The genus *Senegalia* (Fabaceae: Mimosoideae) from the

New World. Phytologia 88(1): 38-93.
- Seigler, DS & Ebinger, JE (2009) New combinations in the genus *Senegalia* (Fabaceae: Mimosoideae). Phytologia 91(1): 26-30.
- Seigler, DS & Ebinger, JE (2010) New combinations in *Senegalia* and *Vachellia* (Fabaceae: Mimosoideae). Phytologia 92(1): 92-95.
- Seigler, DS & Ebinger, JE (2012) *Senegalia aristeguietana* (L. Cardénas) Seigler & Ebinger (Fabaceae: Mimosoideae): an uncommon species of Central America and northern South America. Phytologia 94(2): 276-279.
- Seigler, DS, Ebinger, JE & Glass, C (2012a) *Senegalia berlandieri*, *S. greggii* and *S. wrightii* hybrids (Fabaceae: Mimosoideae) in Texas and adjacent Mexico. Phytologia 94(3): 439-455.
- Seigler, DS, Ebinger, JE & Ribeiro, PG (2012b) A previously unrecognized species of *Senegalia* (Fabaceae) from northeastern Brazil. Journal of the Botanical Research Institute of Texas 6(2): 397-401.
- Seigler, DS, Morim, MP, Barros, M & Ebinger, JE (2013) A new species of *Senegalia* (Fabaceae) from Brazil. Phytotaxa 132(1): 59-63.
- Seigler, DS (2014) A new *Senegalia* (Fabaceae, Mimosoideae) from Southern Peru. Novon 23(1): 90-93.
- Seigler, DS & Ebinger, JE (2014) A new species of *Senegalia* (Fabaceae, Mimosoideae) from Central America and Colombia. Novon 23(1): 94-97.
- Seigler, DS, Ebinger, JE, Ribeiro, PG & De Queiroz, LP (2014) Three new species of *Senegalia (Fabaceae)* from Brazil. J. Bot. Res. Inst. Texas 8(1): 61-69.
- Seigler, DS & Ebinger, JE (2015) New species of Senegalia (Fabaceae) from South America. J. Bot. Res. Inst. Texas 9(2): 335-343.
- Seigler, DS, Ebinger, JE & Glass, C (2015a) Validation of the names *Senegalia x turneri*, *S. x zamudii*, and *Vachellia x ziggyi*. Phytologia 97(3): 224-225.
- Seigler, DS, Ebinger, JE & Glass, C (2015b) A second attempt for validation of the names *Senegalia x turneri*, *S. x zamudii*, and *Vachellia x ziggyi*. Phytologia 97(4): 291-292.
- Seigler, DS & Ebinger, JE (2017) A new *Senegalia*, (*S. alexae*, Fabaceae: Mimosoideae) from Panama, Brazil, and Peru. Phytologia 99(3): 221-225.
- Seigler, DS, Ebinger, JE, Riggins, CW, Terra, V & Miller, JT (2017) *Parasenegalia* and *Pseudosenegalia* (Fabaceae): New genera of the Mimosoideae. Novon 25(2): 180-205.
- Seigler, DS & Ebinger, JE (2018) New combinations in *Parasenegalia* and *Mariosousa* (Fabaceae: Mimosoideae). Phytologia 100(4): 256-259.
- Terra, V & Garcia, FCP (2016) A new species of *Senegalia* (Leguminosae-Mimosoideae) from the Caatinga Domain, Brazil. Phytotaxa 288(2): 181-186.
- Terra, V & Garcia, FCP (2019) Three new species of Senegalia (Leguminosae-Mimosoideae) from the Atlantic Forest domain, Brazil. Phytotaxa 408(1): 30-40.
- WorldWideWattle ver. 2. Published on the Internet at: www.worldwidewattle.com [Accessed 09/10/2019].
- Zhu, X (2015) Nomenclatural novelties and new synonyms of Leguminosae in China. Biodiversity Science 23(2): 247-251.

**Sources of occurrence data**

GBIF, DryFlor, SEINet, Greve *et al*. 2012.

GBIF DOIs: https://doi.org/10.15468/dl.9zjyif, https://doi.org/10.15468/dl.jhbryb, https://doi.org/10.15468/dl.dgx4di, https://doi.org/10.15468/dl.j83m4m.

**Notes**

Maslin *et al*. (2013) consider *Senegalia pennata* to have four subspecies (*pennata*, *kerrii*, *hainanensis*, and *insuavis*). Here we follow Pedley 2014 and Zhu 2015, who treat the latter two as species (*S. hainanensis* and *S. insuavis*, respectively).

Following Hahn 2016, *S. schlechteri* is considered to be a synonym of *S. burkei*.

Following Seigler *et al*. 2017, we treat *S. rurrenabaqueana*, *S. santosii*, *S. skleroxyla*, *S. visco*, *S. muricata*, and *S. vogeliana* as species of *Parasenegalia*, and *S. feddeana* and *S. riograndensis* of *Pseudosenegalia*.

Following Seigler & Ebinger 2018, we treat *S. amorimii*, *S. grazielae*, *S. incerta*, and *S. miersii* as species of *Parasenegalia*.

---

#### *Serianthes*

##### References used for taxonomy and distribution

- Nielsen, IC (1992) Mimosaceae (Leguminosae – Mimosoideae). In: Flora Malesiana, Series I – Spermatophyta, Volume 11 – part 1. Rijksherbarium / Hortus Botanicus, Leiden University, Leiden, the Netherlands.
- Nielsen, IC, Guinet, Ph & Baretta-Kuipers, T (1984) Studies in the Malesian, Australian and Pacific Ingeae (Leguminosae – Mimosoideae): the genera *Archidendropsis*, *Wallaceodendron*, *Paraserianthes*, *Pararchidendron* and *Serianthes* (part 3). Bull. Mus. natn. Hist. nat., Paris, 4e sér., 6, section B, Adansonia 1: 79-111.

##### Sources of occurrence data

GBIF.

GBIF DOIs: https://doi.org/10.15468/dl.drw11w and https://doi.org/10.15468/dl.ru9fym.

##### Notes

*S. vitiensis* is listed among the 'insufficiently known species' by Nielsen *et al*. 1984, but it is included in our dataset.

---

#### *Sympetalandra*, *Thailentadopsis*, *Wallaceodendron*

##### References used for taxonomy and distribution

- Lewis, GP & Schrire, BD (2003) *Thailentadopsis* Kostermans (Leguminosae: Mimosoideae: Ingeae) resurrected. Kew Bulleting 58: 491-494.
- Nielsen, IC (1992) Flora Malesiana. Series I – Spermatophyta. Flowering Plants. Volume 11, part 1. Mimosaceae (Leguminosae – Mimosoideae). Rijksherbarium/Hortus Botanicus, Leiden University,

Leiden, the Netherlands.
- van Steenis, CGGJ (1975) A review of the genus *Sympetalandra* Stapf and its position in Caesalpinioideae. Blumea 22: 159-167.

**Sources of occurrence data**

GBIF.

GBIF DOI: https://doi.org/10.15468/dl.t6ttxa.

---

***Vachellia***

**References used for taxonomy and distribution**

- African Plant Database (version 3.4.0). Conservatoire et Jardin botaniques de la Ville de Genève and South African National Biodiversity Institute, Pretoria, Retrieved July 2019 from http://www.ville-ge.ch/musinfo/bd/cjb/africa/.
- Ali, SI (2014) The genus *Acacia s.l.* in Pakistan. Pak. J. Bot. 46(1): 1-4.
- Boatwright, JS, Maurin, O & van der Bank, M (2015) Phylogenetic position of Madagascan species of *Acacia s.l.* And new combinations in *Senegalia* and *Vachellia* (Fabaceae, Mimosoideae, Acacieae). Botanical Journal of the Linnean Society 179(2): 288-294.
- Clarke, HD, Seigler, DS & Ebinger, JE (1989) *Acacia farnesiana* (Fabaceae: Mimosoideae) and related species from Mexico, the Southwestern U.S., and the Caribbean. Systematic Botany 14(4): 549-564.
- Clarke, HD, Seigler, DS & Ebinger, JE (1990) *Acacia constricta* (Fabaceae: Mimosoideae) and related species from the southwestern U.S. and Mexico. American Journal of Botany 77(3): 305-315.
- Clarke, HD, Seigler, DS & Ebinger, JE (2009) Taxonomic revision of the *Vachellia acuifera* species group (Fabaceae: Mimosoideae) in the Carribean. Systematic Botany 34(1): 84-101.
- Deble, LP & Marchiori, JNC (2010) A new combination in *Vachellia* Wight & Arn. (Mimosaceae). Balduinia 20: 31-34.
- Deshpande, AS, Krishnan, S, Janarthanam, MK & Maslin, BR (2019) Annotated checklist of *Senegalia* and *Vachellia* (Fabaceae: Mimosoideae) for the Indian subcontinent. Nordic Journal of Botany 37(4): 1-20.
- Du Puy, DJ, Labat, J-N, Rabevohitra, R, Villiers, J-F, Bosser, J & Moat, J (2002) The Leguminosae of Madagascar. Kew Publishing.
- Ebinger, JE, Seigler, DS & Clarke, HD (2000) Taxonomic revision of South American species of the genus *Acacia* subgenus *Acacia* (Fabaceae: Mimosoideae). Systematic Botany 25(4): 588-617.
- Ebinger, JE, Seigler, DS & Clarke, HD (2002) Notes on the segregates of *Acacia farnesiana* (L.) Willd. (Fabaceae: Mimosoideae) and related species in North America. The Southwestern Naturalist 47(1): 86-147.
- Ebinger, JE & Seigler, DS (2012) *Vachellia x cedilloi* (L. Rico) Seigler & Ebinger (Fabaceae: Mimosoideae), a probable hybrid of two species of ant-*Acacias*. The Southwestern Naturalist 57(3): 349-353.
- García, R, Clase, T, Seigler, DS & Ebinger, JE (2014) A new species of *Vachellia* (Fabaceae, Mimosoideae) from the Dominican Republic. Novon 23(3): 278-280.
- Greve, M, Lykke, AM, Fagg, CW, Bogaert, J, Friis, I, Marchant, R, Marshall, AR, Ndayishimiye, J, Sandel, BS, Sandom, C, Schmidt, M, Timberlake, JR, Wieringa, JJ, Zizka, G & Svenning, J-C (2012) Continental-scale variability in browser diversity is a major driver of diversity patterns in acacias across Africa. Journal of Ecology 100: 1093-1104.

- Kodela, PG & Wilson, PG (2006) New combinations in the genus *Vachellia* (Fabaceae: Mimosoideae) from Australia. Telopea 11(2): 233-244.
- Kyalangalilwa, B, Boatwright, JS, Daru, BH, Maurin, O & van der Bank, M (2013) Phylogenetic position and revised classification of *Acacia s.l.* (Fabaceae: Mimosoideae) in Africa, including new combinations in *Vachellia* and *Senegalia*. Botanical Journal of the Linnean Society 172(4): 500-523.
- Lee, YS, Seigler, DS & Ebinger, JE (1989) *Acacia rigidula* (Fabaceae) and related species in Mexico and Texas. Systematic Botany 14(1): 91-100.
- Maslin, BR, Seigler, DS & Ebinger, J (2013) New combinations in *Senegalia* and *Vachellia* (Leguminosae: Mimosoideae) for Southeast Asia and China. Blumea 58: 39-44.
- Pedley, L (2014) New combinations for *Senegalia* Raf. and *Vachellia* Wight & Arn. species (Mimosaceae) that occur in Australia. Austrobaileya 9(2): 314-315.
- Ragupathy, S, Seigler, DS, Ebinger, JE & Maslin, BR (2014) New combinations in *Vachellia* and *Senegalia* (Leguminosae: Mimosoideae) for south and west Asia. Phytotaxa 162(3): 174-180.
- Rico-Arce, MdL (2007) American species of *Acacia*. Offset Rebosán, S.A. de C.V., Mexico.
- Seigler, DS & Ebinger, JE (1988) *Acacia macracantha*, *A. pennatula*, and *A. cochliacantha* (Fabaceae: Mimosoideae) species complexes in Mexico. Systematic Botany 13(1): 7-15.
- Seigler, DS & Ebinger, JE (2005a) New combinations in the genus *Vachellia* (Faceae: Mimosoideae) from the New World. Phytologia 87(3): 139-178.
- Seigler, DS & Ebinger, JE (2005b) Taxonomic revision of the ant-*Acacias* (Fabaceae, Mimosoideae, *Acacia*, series Gummiferae) of the New World. Annals of the Missouri Botanical Garden 82(1): 117-138.
- Seigler, DS, Mejía, M, García, R & Ebinger, JE (2012) A new species of *Vachellia* (Fabaceae: Mimosoideae) from Haiti. Journal of the Botanical Research Institute of Texas 6(1): 45-47.
- Seigler, DS & Ebinger, JE (2013) Hybridization between *Vachellia collinsii* and *V. pennatula* (Fabaceae: Mimosoideae) in the New World tropics. Phytologia 95(4): 296-301.
- Seigler, DS & Ebinger, JE (2015) *Vachellia x ruthvenii* (*V. bravoensis* x *V. rigidula*) (Fabaceae: Mimosoideae) in Texas. Phytologia 97(3): 170-174.
- Seigler, DS, Ebinger, JE & Glass, C (2015) Validation of the names *Senegalia x turneri*, *S. x zamudii*, and *Vachellia x ziggyi*. Phytologia 97(3): 224-225.
- Seigler, DS, Ebinger, JE & Glass, C (2015) A second attempt for validation of the names *Senegalia x turneri*, *S. x zamudii*, and *Vachellia x ziggyi*. Phytologia 97(4): 291-292.
- WorldWideWattle ver. 2. Published on the Internet at: www.worldwidewattle.com [Accessed 10/07/2019].

**Sources of occurrence data**

GBIF, DryFlor, SEINet, Greve *et al*. 2012, www.anbg.gov.au/jmiller/factsheets/Vachellia/polypyrigenes.htm.

GBIF DOIs: https://doi.org/10.15468/dl.vbuglm, https://doi.org/10.15468/dl.4k4zhq, https://doi.org/10.15468/dl.gxxret, and https://doi.org/10.15468/dl.bg8uil.

**Notes**

Following Pedley 2014, *Vachellia pallidifolia* (Tindale) Kodela is treated as a synonym of *V. turbata* (Pedley) Pedley.

---

***Viguieranthus***

#### References used for taxonomy and distribution

- Du Puy, DJ, Labat, J-N, Rabevohitra, R, Villiers, J-F, Bosser, J & Moat, J (2002) The Leguminosae of Madagascar. Kew Publishing.

#### Sources of occurrence data

GBIF.

GBIF DOI: https://doi.org/10.15468/dl.kj4rzc.

#### Notes

Varieties of *V. densinervus* and *V. perrieri* are here only included at the species level.

---

### *Xylia*

#### References used for taxonomy and distribution

- Brenan, JPM (1959) Flora of Tropical East Africa. Leguminosae subfamily Mimosoideae.
- Brenan, JPM (1970) Flora Zambesiaca. Volume three: part one. Leguminosae.
- Du Puy, DJ, Labat, J-N, Rabevohitra, R, Villiers, J-F, Bosser, J & Moat, J (2002) The Leguminosae of Madagascar. Kew Publishing.
- Hutchinson, J, Dalziel, JM & Keay, RWJ (1954) Flora of West Tropical Africa. Second edition. Volume 1, part 1.
- Ramesh, BR, Swaminath, MH, Patil, SV, Dasappa, Pélissier, R, Venugopal, PD, Aravajy, S, Elouard, C & Ramalingam, S (2010) Forest stand structure and composition n 96 sites along environmental gradients in the central Western Ghats of India. Ecology 91(10): 3118-3118.
- Robyns, W (ed.) (1952) Flore du Congo Belge et du Ruanda-Urundi. Spermatophytes. Volume 3. Brussels, Belgium.
- Sanjappa, M (1991) Legumes of India. Bishen Singh Mahendra Pal Singh, Dehra Dun, India.

#### Sources of occurrence data

GBIF, Ramesh *et al.* 2010.

GBIF DOI: https://doi.org/10.15468/dl.hdmafv.

---

### *Zapoteca*

#### References used for taxonomy and distribution

- Bässler, M (1998) Flora de la República de Cuba. Seria A: Plantas Vasculares. Fascículo 2: Mimosaceae. Koeltz Scientific Books, Koenigstein, Germany.
- Hernández, HM (1986) *Zapoteca*: a new genus of Neotropical Mimosoideae. Annals of the Missouri Botanical Garden 73(4): 755-763.
- Hernández, HM (1989) Systematics of *Zapoteca* (Leguminosae). Annals of the Missouri Botanical Garden 76(3): 781-862.
- Hernández, HM (1990) A New Subgenus and a New Species of *Zapoteca* (Leguminosae). Systematic Botany 15(2): 226-230.
- Hernández, HM & Campos V, A (1994) A New Species of *Zapoteca* (Leguminosae, Mimosoideae) from Mexico. Novon 4(1): 32-34.
- Hernández, HM & Hanan-Alipi, AM (1998) *Zapoteca quichoi* (Leguminosae, Mimosoideae), a new species from southeastern Mexico. Brittonia 50(2): 211-213.
- Hernández, HM (2015) New taxa of *Zapoteca* (Leguminosae, Mimosoideae) from Mexico. Phytotaxa 239(3): 223-241.
- Levin, GA & Moran, G (1989) The vascular flora of Isla Socorro, Mexico. Memoirs of the San Diego Society of Natural History 16: 1-71.

**Sources of occurrence data**

GBIF, DryFlor, SEINet.

GBIF DOI: https://doi.org/10.15468/dl.0ttcmt.

**Notes**

Following Bässler 1998, *Z. formosa* subsp. *gracilis* is included here as *Z. gracilis*.

---

**Non-Mimosoid Caesalpinioideae**

The occurrence dataset of the non-Mimosoid Caesalpinioideae only contains data for the non-Mimosoid Caesalpinioideae species present in the phylogenomic backbone.

**References used for taxonomy and distribution**

- Genève and South African National Biodiversity Institute, Pretoria, “Retrieved September 2021”, from <http://africanplantdatabase.ch>.
- Barneby, R. C. (1996). Neotropical Fabales at NY: Asides and oversights. Brittonia, 48(2), 174–187. https://doi.org/10.2307/2807811
- Bortoluzzi, R. L. C., Lima, A. G., Souza, V. C., Rosignoli-Oliveira, L. G., & Conceição, A. S. (2020). *Senna*. Flora do Brasil 2020. Jardim Botânico do Rio de Janeiro.Disponível em: <http://floradobrasil.jbrj.gov.br/reflora/floradobrasil/FB83716>. Accessed on: 22 September 2021.
- Chudnoff, M. (1984). Tropical Timbers of the World. Agriculture Handbook Number 607. USDA Forest Service.
- Cota, M. M. T. (2020a). *Batesia*. Flora do Brasil 2020. Jardim Botânico do Rio de Janeiro.Disponível em: <http://floradobrasil.jbrj.gov.br/reflora/floradobrasil/FB22810>. Acessed on: 27 September 2021.
- Cota, M. M. T. (2020b). *Campsiandra*. Flora do Brasil 2020. Jardim Botânico do Rio de Janeiro.Available at: <http://floradobrasil.jbrj.gov.br/reflora/floradobrasil/FB78539>. Accessed on:

01 Oct. 2021.
- Cota, M. M. T. (2020c). *Recordoxylon*. Flora do Brasil 2020. Jardim Botânico do Rio de Janeiro.Available at: <http://floradobrasil.jbrj.gov.br/reflora/floradobrasil/FB109405>. Accessed on: 22 September 2021.
- da Silva, M. F. (1986). *Dimorphandra* (Caesalpiniaceae). Flora Neotropica, 44, 1–127.
- Du Puy, D. J., Labat, J.-N., Rabevohitra, R., Villiers, J.-F., Bosser, J., & Moat, J. (2002). The Leguminosae of Madagascar. Kew, United Kingdom: Royal Botanic Gardens.
- Dwyer, J. D. (1954). The Tropical American Genus *Tachigalia* Aubl. (Caesalpiniaceae). Annals of the Missouri Botanical Garden, 41(2), 223–260.
- Gagnon, E., Ringelberg, J. J., Bruneau, A., Lewis, G. P., & Hughes, C. E. (2019). Global Succulent Biome phylogenetic conservatism across the pantropical Caesalpinia Group (Leguminosae). New Phytologist, 222(4), 1994–2008. https://doi.org/10.1111/nph.15633
- Irwin, H. S., & Barneby, R. C. (1978). Monographic studies in *Cassia* (Leguminosae-Caesalpinioideae) III. Sections Absus and Grimaldia. Memoirs of the New York Botanical Garden, 30, 1–277.
- Irwin, H. S., & Barneby, R. C. (1982). The American Cassiinae. A synoptical revision of Leguminosae, tribe Cassieae, subtribe Cassiinae in New World. Memoirs of the New York Botanical Garden, 35(1–2), 1–918.
- Irwin, H. S., & Rogers, D. J. (1967). Monographic studies in *Cassia* (Leguminosae-Caesalpinioideae). II A Taximetric study of Section Apoucouita. Memoirs of the New York Botanical Garden, 16, 1–120.
- Kuntz, J., & Lima, A. G. (2020). *Diptychandra*. Flora do Brasil 2020. Jardim Botânico do Rio de Janeiro.Disponível em: <http://floradobrasil.jbrj.gov.br/reflora/floradobrasil/FB83135>. Acesso em: 27 set. 2021.
- Lee, Y.-T. (1976). The genus *Gymnocladus* and its tropical affinity. Journal of the Arnold Arboretum, 57, 91–112.
- Lewis, G. P., Schrire, B., Mackinder, B., & Lock, M. (2005). Legumes of the world. Kew, United Kingdom: Royal Botanic Gardens.
- Lewis, G. P., Siqueira, G. S., Banks, H., & Bruneau, A. (2017). The majestic canopy-emergent genus *Dinizia* (Leguminosae: Caesalpinioideae), including a new species endemic to the Brazilian state of Espírito Santo. Kew Bulletin, 72, 48. https://doi.org/10.1007/S12225-017-9720-7
- Lima, A. G., & Kuntz, J. (2020). *Arapatiella*. Flora do Brasil 2020. Jardim Botânico do Rio de Janeiro.Available at: <http://floradobrasil.jbrj.gov.br/reflora/floradobrasil/FB78506>. Accessed on: 01 Oct. 2021.
- Pipoly III, J. J. (1995). A new *Tachigali* (Fabaceae: Caesalpinioideae) from western Amazonia. Sida, 16(3), 407–411.
- POWO. (2021). Plants of the World Online. Facilitated by the Royal Botanic Gardens, Kew. Published on the Internet; http://www.plantsoftheworldonline.org/ Retrieved 01 October 2021.
- Queiroz, L. P. (2009). Leguminosas da Caatinga. Universidade Estadual de Feira de Santana.
- Rando, J. G., Carvalho, D. A. S., & Silva, T. S. (2020). *Melanoxylon*. Flora do Brasil 2020. Jardim Botânico do Rio de Janeiro.Available at: <http://floradobrasil.jbrj.gov.br/reflora/floradobrasil/FB78737>. Accessed on: 22 September 2021.
- Rando, J. G., Cota, M. M. T., Conceição, A. S., Barbosa, A. R., & Barros, T. L. A. (2020). *Chamaecrista*. Flora do Brasil 2020. Jardim Botânico do Rio de Janeiro. Available at: <http://floradobrasil.jbrj.gov.br/reflora/floradobrasil/FB82881>. Accessed on: 22 September 2021.
- Ringelberg, J. J., Zimmermann, N. E., Weeks, A., Lavin, M., & Hughes, C. E. (2020). Biomes as evolutionary arenas: Convergence and conservatism in the trans-continental succulent biome. Global Ecology and Biogeography, 29(7), 1100–1113. https://doi.org/10.1111/geb.13089
- Souza, V. C., & Lima, A. G. (2020). *Dimorphandra*. Flora do Brasil 2020. Jardim Botânico do Rio de Janeiro.Disponível em: <http://floradobrasil.jbrj.gov.br/reflora/floradobrasil/FB78675>. Acesso em: 27 set. 2021.
- Stergios, B. (1996). Contributions to South American Caesalpiniaecae. II. A taxonomic update of *Campsiandra* (Caesalpinieae). Novon, 6(4), 434–459.

- van der Werff, H. (2008). A synopsis of the genus *Tachigali* (Leguminosae: Caesalpinioideae) in northern South America. Annals of the Missouri Botanical Garden, 95(4), 618–660. https://doi.org/10.3417/2007159
- Vivas, C. V., & Queiroz, L. P. (2020). *Moldenhawera*. Flora do Brasil 2020. Jardim Botânico do Rio de Janeiro.Available at: <http://floradobrasil.jbrj.gov.br/reflora/floradobrasil/FB28148>. Accessed on: 01 Oct. 2021.

**Sources of occurrence data**

See below for data sources.

GBIF DOIs: https://doi.org/10.15468/dl.fkhywf, https://doi.org/10.15468/dl.jtk2kr, https://doi.org/10.15468/dl.m8gxhd, https://doi.org/10.15468/dl.wgqs87, https://doi.org/10.15468/dl.acsyq7, https://doi.org/10.15468/dl.m3pxsx, https://doi.org/10.15468/dl.5gxx8t, https://doi.org/10.15468/dl.ktjgw9, https://doi.org/10.15468/dl.h8bsbs, and http://doi.org/10.15468/dl.ckr5lw.

**References and notes per species**

The resources used to check and correct distribution ranges are listed for each species, as well as specific notes. Unless specified otherwise, all data were newly downloaded from GBIF.

***Acrocarpus fraxinifolius***: Lewis et al. (2005).

***Arapatiella sp.***: Lima & Kuntz (2020). As the species is not specified, data from both species in this genus (i.e., *A. emarginata* and *A. psilophylla*) were pooled to represent this taxon.
***Arcoa gonavensis***: Lewis et al. (2005).

***Arquita trichocarpa***: data from Gagnon et al. (2019).

***Balsamocarpon brevifolium***: data from Gagnon et al. (2019).

***Batesia floribunda***: Cota (2020a).

***Biancaea decapetala***: data from Gagnon et al. (2019). As no occurrence data of *B. decapetala* were available, the data of other *Biancaea* species in the dataset of Gagnon et al. (2019) were pooled to represent this taxon.

***Burkea africana***: African Plant Database (2021).

***Bussea perrieri***: Du Puy et al. (2002).

***Caesalpinia cassioides***: data from Gagnon et al. (2019).

***Caesalpinia crista***: data from Gagnon et al. (2019).

***Campsiandra comosa***: Stergios (1996), POWO (2021), and Cota (2020b). Stergios (1996) and POWO (2021) do not list this species as occurring outside the Guianas, and it is not mentioned by Cota (2020b) in the Flora of Brazil, so all occurrences outside the Guianas were removed.

***Cassia cowanii var. guianensis***: Irwin & Barneby (1982). Collection Irwin, H.S. et al. 47820, from Brazil, was determined by Irwin as *C. cowanii var. guianensis*, but the specimen is without flowers to confirm the determination, so it was excluded, as was Pires et al. 50730, collected in Amapá, Brazil.

***Cenostigma pluviosum var. maraniona***: data from Gagnon et al. (2019).

***Ceratonia siliqua***: Lewis et al. (2005).

***Chamaecrista adiantifolia***: Irwin & Rogers (1967) and Rando, Cota et al. (2020).

***Chamaecrista lineata***: Irwin & Barneby (1982).

***Chamaecrista ramosa***: Irwin & Barneby (1982) and Rando, Cota et al. (2020). Although the specimen from Belize (Ratter, J.A.; Bisset, H.A.; Bridgewater, S.G.M. 6626, K herbarium) is a geographical outlier and does not have an available image, it was determined by an expert, and therefore kept in the dataset.

***Chamaecrista viscosa***: Irwin & Barneby (1978) and Rando, Cota et al. (2020).

***Colvillea racemosa***: data from Ringelberg et al. (2020).

***Conzattia multiflora***: data from Ringelberg et al. (2020).

***Cordeauxia edulis***: data from Gagnon et al. (2019).

***Coulteria platyloba***: data from Gagnon et al. (2019).

***Delonix decaryi***: data from Ringelberg et al. (2020).

***Delonix edule***: data from Ringelberg et al. (2020). Formerly known as *Lemuropisum edule*.

***Denisophytum madagascariense***: data from Gagnon et al. (2019).

***Dimorphandra davisii***: da Silva (1986) and Souza & Lima (2020).

***Dimorphandra gardneriana***: da Silva (1986) and Souza & Lima (2020).

***Dimorphandra macrostachya***: da Silva (1986) and Souza & Lima (2020).

***Dinizia jueirana-facao***: Lewis et al. (2017).

***Diptychandra aurantiaca***: Kuntz & Lima (2020) and Queiroz (2009).

***Erythrophleum ivorense***: African Plant Database (2021).

***Erythrophleum teysmannii***: POWO (2021).

***Erythrostemon coluteifolus***: data from Gagnon et al. (2019).

***Erythrostemon mexicanus***: data from Gagnon et al. (2019).

***Gelrebia rostrata***: data from Gagnon et al. (2019).

***Gleditsia sinensis***: Flora of China, Volume 10: Fabaceae (http://www.efloras.org/florataxon.aspx?flora_id=2&taxon_id=200012140).

***Guilandina bonduc***: data from Gagnon et al. (2019).

***Gymnocladus dioica***: Lee (1976).

***Haematoxylum brasiletto***: data from Gagnon et al. (2019).

***Heteroflorum sclerocarpum***: data from Ringelberg et al. (2020).

***Hererolandia pearsonii***: data from Gagnon et al. (2019).

***Hoffmannseggia arequipensis***: data from Gagnon et al. (2019). As no occurrence data of *H. arequipensis* were available, the data of other *Hoffmannseggia* species in the dataset of Gagnon et al. (2019) were pooled to represent this taxon.

***Jacqueshuberia brevipes***: POWO (2021).

***Libidibia glabrata***: data from Gagnon et al. (2019).

***Lophocarpinia aculeatifolia***: data from Gagnon et al. (2019).

***Melanoxylon brauna***: Rando, Carvalho, et al. (2020).

***Mezoneuron kauaiense***: data from Gagnon et al. (2019).

***Moldenhawera floribunda***: Vivas & Queiroz (2020).

***Mora gonggrijpii***: Chudnoff (1984).

***Moullava spicata***: data from Gagnon et al. (2019). As no occurrence data of *M. spicata* were available, the data of other *Moullava* species in the dataset of Gagnon et al. (2019) were pooled to represent this taxon.

***Pachyelasma tessmannii***: African Plant Database (2021).

***Parkinsonia andicola***: data from Ringelberg et al. (2020).

***Paubrasilia echinata***: data from Gagnon et al. (2019).

***Peltophorum africanum***: African Plant Database (2021).

***Peltophorum dubium***: Barneby (1996). All points in Florida were excluded, as these likely represent cultivated plants.

***Pomaria jamesii***: data from Gagnon et al. (2019).

***Pterolobium stellatum***: data from Gagnon et al. (2019).

***Recordoxylon speciosum:*** Cota (2020c).

***Schizolobium parahyba***: Barneby (1996).

***Senna cushina***: Irwin and Barneby (1982).

***Senna lasseigniana***: POWO (2021). The collection of Oldeman 2133 from French Guiana is not *S. lasseigniana*, and was therefore excluded.

***Senna leandrii***: Du Puy et al. (2002).

***Senna mollissima***: Irwin and Barneby (1982). Points in Brazil were excluded.

***Senna rugosa***: Irwin and Barneby (1982) and Bortoluzzi et al. (2020).

***Senna velutina:*** Irwin and Barneby (1982) and Bortoluzzi et al. (2020).

***Stachyothyrsus staudtii***: African Plant Database (2021).

***Stuhlmannia moavi***: data from Gagnon et al. (2019).

***Tachigali bracteolata***: Dwyer (1954) and Van der Werff (2008). Van der Werff (2008) considers this species a synonym of *T. richardiana*, but here it is kept as a separate entity.

***Tachigali guianensis***: Dwyer (1954) and Van der Werff (2008).

***Tachigali odoratissima***: Dwyer (1954) and Van der Werff (2008).

***Tachigali paniculata***: Dwyer (1954) and Van der Werff (2008).

***Tachigali vasquezii***: Pipoly III (1995).

***Tara spinosa***: data from Gagnon et al. (2019).

***Tetrapterocarpon geayi***: Du Puy et al. (2002).

***Umtiza listeriana***: Lewis et al. (2005).

***Zuccagnia punctata***: data from Gagnon et al. (2019).

**Notes**

Except for taxa specifically identified at the infraspecific level in the phylogeny (e.g., *Cassia cowanii* var. *guianensis*), occurrence records identified at the infraspecific level were included at the specific level.
